# GEO-BERT: A Retrieval augmented LLM Framework for Intelligent Dataset Retrieval from the Gene Expression Omnibus

**DOI:** 10.64898/2026.10.15.759430

**Authors:** Maruf Islam Shuvo, Monjura Afrin Rumi, Naznin Akhter, Ewy A. Mathé, Shafiul Alam, Mahfuza Sharmin

## Abstract

The Gene Expression Omnibus (GEO) is a commonly used data repository of high throughput gene expression and other functional genomics data. This platform is globally used by researchers for submitting experimental datasets, accessing these datasets for hypothesis validation, reusing them in data-driven research, generating ideas for new research directions. Despite global usage, searching and identifying relevant and usable datasets in GEO remains challenging with an underlying keyword based search system needing manual intervention for the retrieval of desired datasets. GEO’s inconsistent metadata and varying data annotation practices across submissions, variable data quality are contributing to its challenge for building structured query tools. With the recent advancement of Large Language Models (LLMs), it is timely to revisit the GEO datasets and its query system. Here, we present an AI chat-bot, GEO-BERT, leveraging an underlying retrieval augmented generator framework on the GEO metadata with a novel hierarchical hybrid field aware retrieval system to mine the datasets. GEO-BERT is locally installable, available for programmers and project developers (https://github.com/mrumi/GEOBERTtest) and readily usable by regular users on subset of test data. We demonstrated that GEO-BERT enhances the semantic search performance not currently available by any of the leading existing AI-powered chatbots, while improving the performance of GEO’s traditional search performance for project development. Given the global usage of GEO and the latest advancement of the large language models, GEO-BERT fills the gap for context-aware extraction of GEO datasets.

## Introduction

The Gene Expression Omnibus (GEO) is a public database of high-throughput gene expression and other functional genomics data —such as hybridization arrays, microarrays, and expression chips—submitted by the biomedical research community ^1,2^. The research community widely reuse these datasets to validate hypotheses, generate new research questions, perform independent data validation across studies. In addition, GEO serves as an invaluable resource for data-driven research enabling scientists to mine through hundreds or even thousands of samples to uncover new hypotheses and detect meaningful patterns and behaviors in biological data. However, navigating GEO’s vast collection, comprising millions of samples across thousands of studies, is extremely challenging ^3–5^. Identifying the most relevant and high-quality datasets for a specific research question often requires significant time and manual effort. Several factors contribute to the difficulty of finding relevant datasets – a) keyword-based search system, b) inconsistency of the metadata upon which the search system is built on, and c) the metadata collection process. Each GEO dataset typically includes metadata describing the study design, sample characteristics, experimental conditions and data matrices representing the measurements from the high-throughput experiment. All metadata of thousands of studies are indexed and made searchable by NCBI’s powerful engine. The basis of the GEO search system is keywords and Boolean logic of filters and field-specific queries through structured metadata. This search interface is limited due to lack of support for semantic search, synonyms, related concepts or context-aware querying. The GEO search system is further limited by inconsistent metadata, stemming from its collections strategy and experimental complexity. During GEO data submission, the researchers often describe their experiments using free text without enforced standardization, controlled vocabularies or ontologies, and use different terms for the same concepts (e.g., “liver” vs. “hepatic tissue”). This results in researcher-specific language, jargon, biological nuance, ambiguity, unstructured descriptions in the metadata. These challenges are further extended by the complexity of experimental contexts – there is no universal guideline for grouping samples, labeling samples, ordering information leading to lack of clarity. Additional sources of inconsistency are introduced by variability in platform, data format, annotation styles, multi-factor experimental designs and levels of detail.

To facilitate dataset navigation, GEO offers a GUI-based web search, in addition, several tools have also been developed for programmatic access. The NCBI E-Utilities (Entrez API) enables automated searches without using a web browser. The GEOquery ^6^, a Bioconductor package, allows retrieval of datasets using accession numbers but does not support search functionality. The GEOParse ^7^, a python-based tool, supports keyword-based search and data retrieval. The GEOMetadb ^8^, another Bioconductor/R package provides a searchable SQLite database with download options. Biopython ^9^ is another python package offering lightweight GEO searches. Despite these capabilities, all these tools share common limitations - reliance on keyword-based queries, no support for complex or semantic search and inconsistent metadata handling.

Numerous secondary websites have been developed containing transcriptomics data from GEO, serving diverse purposes such as exploratory data analysis, differential expression analysis, expression pattern visualization, access to uniformly processed datasets, LLM-powered querying of single-cell data, gene signature exploration, and co-expression analysis ^10–14^. However, these platforms all share a common limitation: they rely on the keyword-based search system of GEO, inheriting its associated drawbacks.

To improve dataset discoverability, researchers have developed several resources aimed at standardizing metadata. ReGEO focuses on restructuring metadata from GEO’s functional genomics data ^3^, MetaSRA curates and maps metadata for human samples in SRA ^15^, and BioSample stores structured metadata for biological samples linked to sequence data in repositories like GEO and SRA ^16^. However, these tools still rely on keyword-based, non-semantic search mechanisms. Additional tools such as GEOMetaCuration are used to harmonize metadata across datasets ^17^, TidyGEO helps clean and simplify complex metadata tables ^18^, and CEDAR supports the creation of structured ^19^, ontology-linked metadata during data submission. More recently, two large language model (LLM)-based approaches, have shown promising potential for automating metadata curation and annotating GEO datasets: CistromeMeta ^20^ and ChiP-GPT ^21^ focusing on ChIPSeq and Epigenetic metadata. None of these resources allow researchers to directly query the datasets of interest in natural language.

Natural language processing is tailored for understanding free text, vocabullaries, concepts with different terms; and repurposing various advances in Large Language Models (LLMs) should be the solution for replacing ancient keyword based search with context-aware search from GEO datasets ^22^. In this article, we developed GEO-BERT, a retrieval augmented generator (RAG) framework to query GEO datasets in natural language. The framework combines a GEO specific context-aware data preprocessing, a fusioned field aware semantic indexing by biomedical specific embedding model - PubMedBERT ^23^ and SPLADE ^24^, layered storages (vocabulary index, multi-stage vector database) for scalable retrieval, hierarchical filed-aware hybrid retrieval, dataset-grounded query interpretation and answer generation to avoid hallucinations (Figure 1A). We evaluated GEO-BERT across 10 queries using seven RAG evaluation metrics, constructing a manually curated reference dataset for each query to support performance assessment. We further benchmarked GEO-BERT against Gemini, a widely used AI-powered chatbot, and the conventional GEO search system, using the same queries spanning both keyword-based and semantic search tasks. Finally, we conducted a case study comparing GEO and GEO-BERT for data-driven resource construction, demonstrating how improved dataset discovery can facilitate downstream research. Overall, GEO-BERT consistently outperformed the comparison systems across both keyword-based and semantic retrieval tasks.

**Figure 1.**
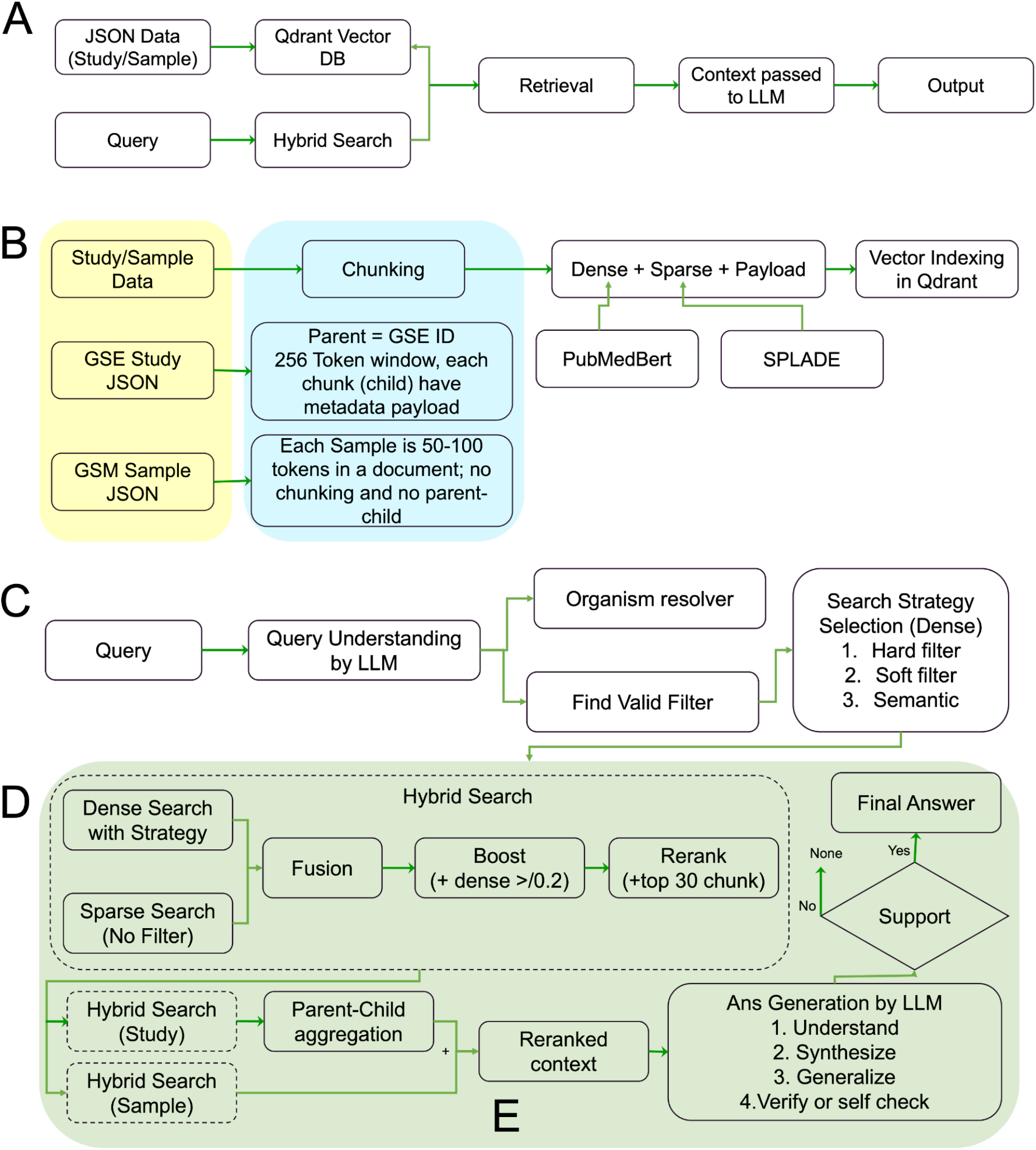
A) GEO-BERT Pipeline, B) Data Ingestion and Indexing, C) Query Understanding and Retrieval Preparation, D) Hybrid Search, E) Evidence Grounded Answer Generation.

## Results

### GEO-BERT workflow

GEO-BERT optimizes the semantic search system following a Retrieval Augmented Generation (RAG) framework where we propose a hybrid retrieval algorithm on a multi-level storage consisting of two index vectors. The whole system is divided into four layers (**Figure 1B-E, details are in Methods**): 1) Data Ingestion and Indexing, 2) Query Understanding and Retrieval Preparation, 3) Hybrid Retrieval with confidence aware reranking and 4) Evidence grounded Answer Generation. In the 1st step, we preprocessed the metadata extracted from the GEO portal into separate study and sample level json files to reduce the data redundancy of each sample; optionally chunked the study json files maintaining parent-child relationship for each GSE id for better tracking during retrieval; generated three component index vectors (dense, sparse and payload obtained by pubmedbert, SPLADE and Qdrant ^25^) for each chunk or json files; and finally stored the index vectors in Qdrant vector DB. In the 2nd step, upon query arrival, we generated a structured seven-field json file by the LLM inference of the query to represent its intent, type etc where the query type describes the main nature of the query the user wants; if necessary identified constraints needed by the query and normalized these constraints and organism name against NCBI Taxonomy Database ^26^ ; decided a query-specific retrieval strategy based on the number of constraints validated against payload indexes; and finally converted the query json into similar three component vector (dense+sparse+payload) where the payload consists of the constraints obtained from the query. In the 3rd step, during the hybrid retrieval, we performed both dense and sparse retrieval; fusioned the retrieval score; boosted the retrieval score based on query constraints; reranked the retrieved vectors with an additional cross-encoder, aggregated any existing parent-child chunk for the study vectors, separately retrieved the samples in similar fashion; and built contexts for both study and samples. In the 4th step, we combined the retrieved studies and samples to format the answer, designed the prompts with query intents, study and and sample contexts and verification of factual claims against the context to reduce hallucination.

To evaluate our framework, we assembled a test dataset comprising approximately 6,000 GEO Series (GSE) accessions, selected to provide a combination of random sampling, broad representation, and enrichment for retinal studies as a focused biological use case (Supplementary Table 1). We benchmarked GEO-BERT using 10 queries (Supplementary Table 2). For each query, candidate GSE accessions retrieved by GEO-BERT, conventional GEO search, and Gemini were pooled and manually reviewed to construct a reference dataset for relevance check (Supplementary Table 3). The framework can be run locally on a desktop using this test dataset, enabling researchers and developers to perform semantic dataset retrieval and identify relevant processed data for downstream analysis.

### GEO-BERT Provides Improved Dataset Retrieval and Facilitates Data-Resource Construction

To evaluate GEO-BERT’s utility for dataset retrieval, we compared its results with those returned by GEO. For each query, GEO-BERT provides an answer summary, study- and sample-level statistics, matching GSE IDs with brief explanations and relevance scores, an overall assessment of match quality, and downloadable session history (Figure 2A). Using 10 queries and their query-specific Ground Truth datasets (Supplementary Table 3), GEO-BERT outperformed or performed comparably to GEO for 9 of 10 queries (Figure 2B). Notably, GEO-BERT missed fewer true-positive datasets across all queries; this advantage is not fully reflected in Figure 2B because missing results were treated as negative predictions (Figure 2D).

**Figure 2.**
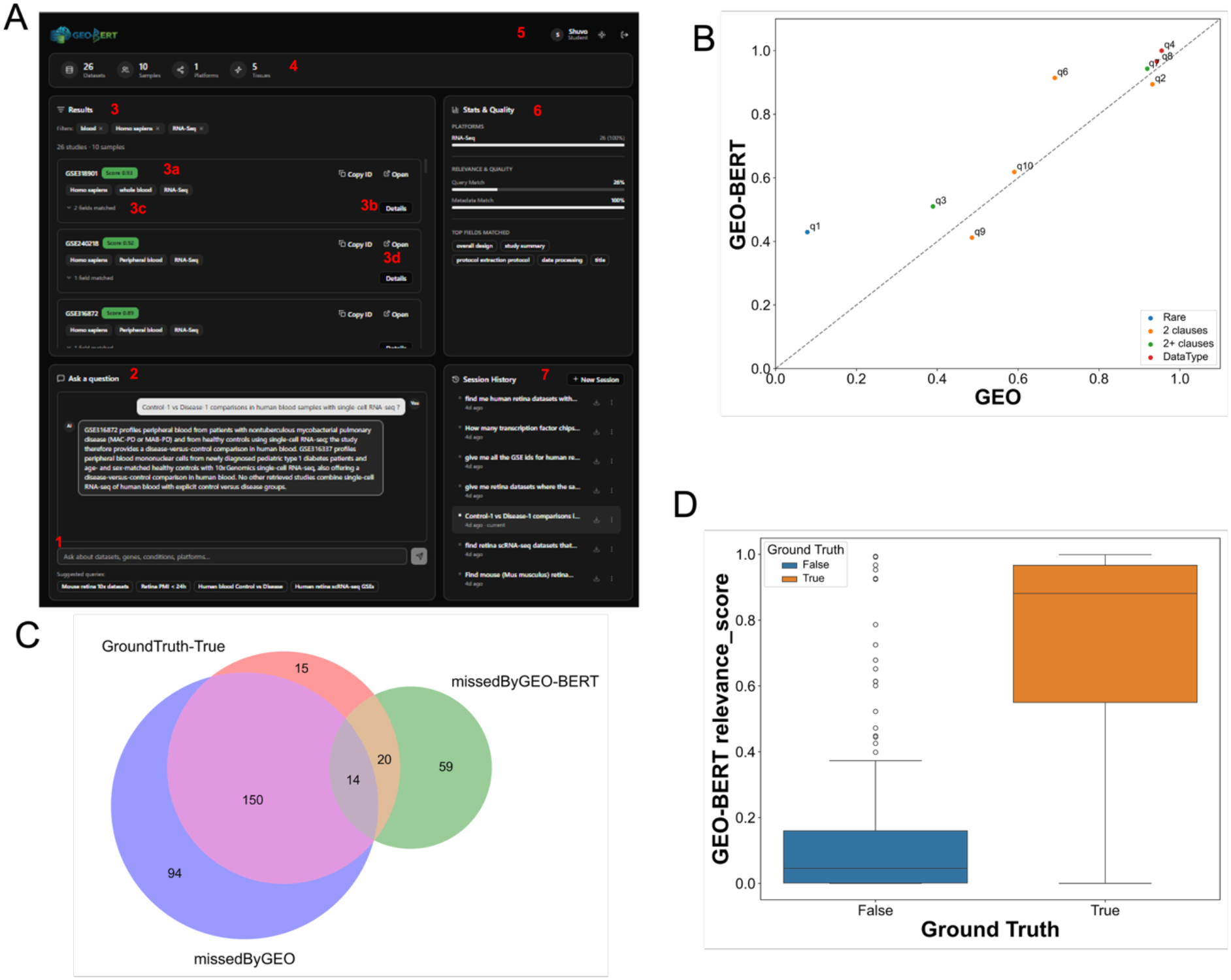
A) Application FrontEnd (1. Query Input Panel, 2. Overall summary, 3. Search Results (a. Total Results, b.View snapshot details, c. Matched Fields), 4. View in GEO, 5. Result statistics, 6. User and Role, 7. Overall Quality and Matches, 8. Downloadable Session History); B) Performance comparison between GEO-BERT and GEO, C) GroundTruth-True missed by GEO and GEO-BERT, D) Relevance scores of the retrieved data of GEO-BERT.

Manual examination revealed several limitations of keyword-based retrieval. Query terms frequently appeared in study descriptions without the corresponding data being deposited in GEO. For example, a proteomics query (q1) retrieved records mentioning *protein, proteostasis*, or *proteomics* even when the deposited data consisted of RNA-seq, ChIP-seq, CRISPR screens, or spatial transcriptomics. In other cases, proteomics data were deposited elsewhere, obtained from external datasets, or mentioned only in metadata, including the journal name. Conversely, relevant datasets could be missed when metadata used semantically related terms rather than the exact query term; for retina-related queries (q2, q6, and q9), for example, relevant records sometimes used terms such as *eye, eye disease*, or *ocular disease* rather than *retina*. Queries containing specific datatype requirements also resulted in missed datasets in GEO (q4 and q8). Both GEO and GEO-BERT performed well for queries targeting nonexistent data (q5, not shown in the plot). GEO-BERT retrieval is currently constrained by the number of top-ranked vectors that can be passed to the LLM within the available token limit, which may restrict recall for queries with many relevant datasets.

Because GEO-BERT uses retrieval-augmented generation (RAG) to retrieve top-ranked candidates, we next examined whether its relevance scores could help users distinguish highly relevant datasets from partial matches. GEO-BERT assigned significantly higher relevance scores to Ground Truth–positive than Ground Truth–negative datasets (mannwhitney pvalue 6.18e-34), indicating that the scores effectively discriminate datasets according to query relevance (Figure 2C). We further evaluated the practical utility of this feature by simulating dataset retrieval for a hypothetical portal containing human-retina scRNA-seq and ChIP-seq datasets. Because GEO-BERT provides a semantic explanation for each match, candidate datasets could be evaluated rapidly against the query. In our manual comparison restricted to the approximately 6,000 datasets used in this analysis, reviewing GEO-BERT results required approximately 5 minutes, compared with approximately 2 hours to cross-check comparable GEO search results.

Taken together, these results indicate that GEO-BERT can improve the precision and interpretability of GEO dataset retrieval, incorporate additional metadata into retrieval decisions, and substantially reduce the effort required for downstream manual review.

### GEO-BERT Enables Semantic Retrieval of Relevant GEO Datasets

We first examined how samples in our test dataset were represented in the embedding space. To this end, we randomly selected 1,000 sample vectors from the Qdrant database and visualized their embeddings, focusing on blood- and retina-related samples represented by heterogeneous metadata annotations. Blood-associated samples included labels ranging from “blood” and “whole blood” to “peripheral blood mononuclear cells” and “blood plasma,” while retinal samples encompassed diverse annotations such as “retina,” “RPE,” “retinal organoid,” “retinal explant,” “optic nerve,” and “whole eye” (Supplementary Table 4). Despite substantial variation in terminology, biologically related samples tended to localize near one another in the embedding space, indicating that the vector representations capture biological similarity beyond exact lexical matches (Figure 3A). Within the retina-associated samples, we further observed the structure consistent with biological context: annotations corresponding to retina and RPE tended to be more closely related, whereas fetal retinal and retinal-organoid samples occupied more distinct regions of the embedding space (Figure 3B).

**Figure 3.**
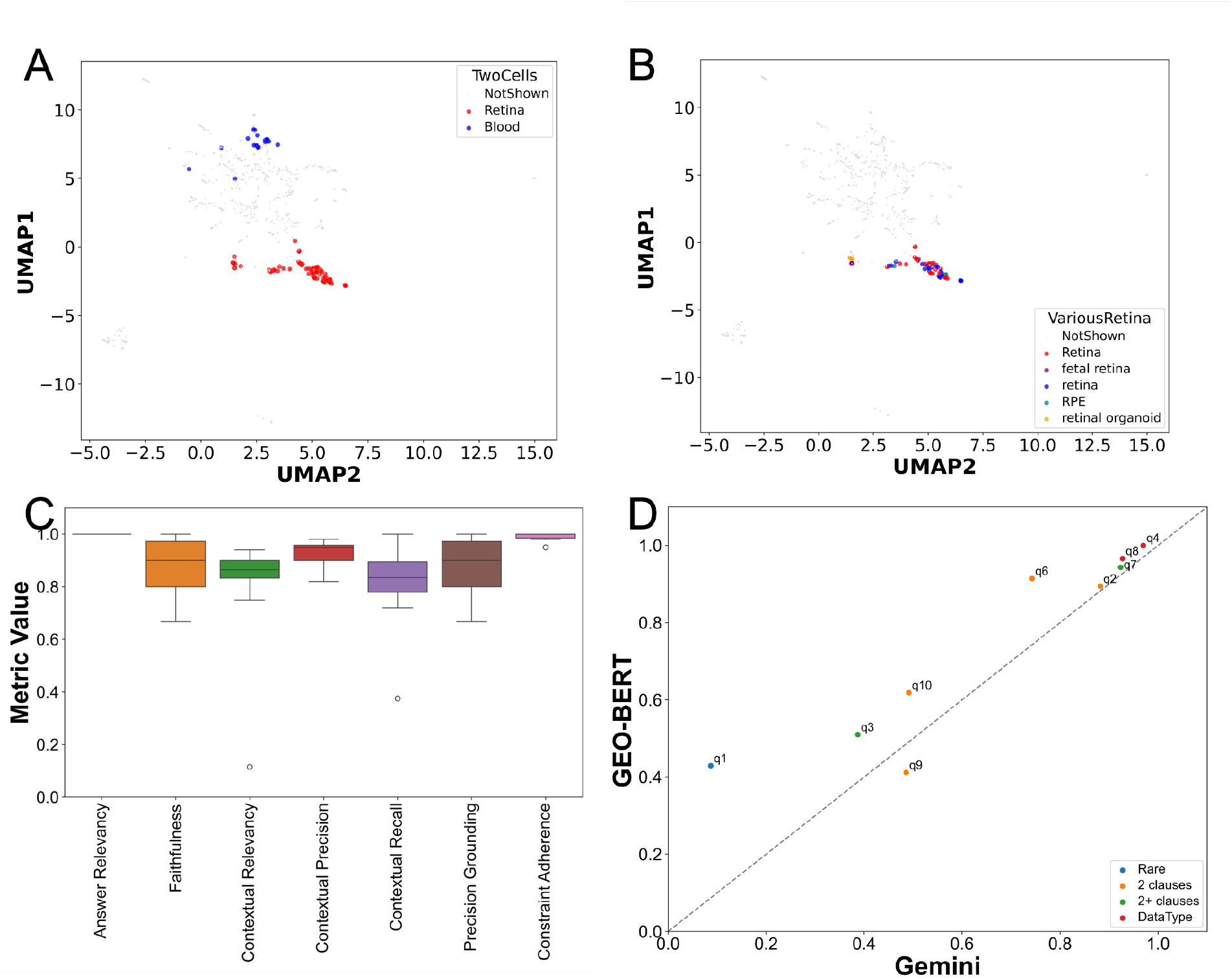
A-B) Semantic relations of the samples with similar tissues, C) Performance measure of GEO-BERT using RAG evaluation metrics, D) Performance comparison between GEO-BERT and existing AI-powered chatbot, Gemini.

We next evaluated GEO-BERT using seven RAG evaluation metrics for assessing *Answer Quality, Retrieval Context Quality, Grounding & Constraint Compliance* (Supplementary Table 5). Across the evaluated queries, the RAG evaluation metrics showed consistently strong performance across the 10 queries (Figure 3C). For Answer Quality, Answer Relevancy was uniformly high (1.0 for all queries), indicating that the generated responses consistently addressed the intended questions, while Faithfulness had a median score of 0.90, showing that the answers were generally supported by the retrieved evidence. For Retrieval Context Quality, Contextual Precision had a median of 0.95, indicating that relevant evidence was generally prioritized within the retrieved context. Contextual Recall had a median of 0.84, while Contextual Relevancy had a median of 0.87, with greater variation across queries, suggesting that the coverage and relevance of retrieved information depended more on the specific query. For Grounding and Constraint Compliance, Precision Grounding had a median score of 0.90 and Constraint Adherence remained consistently high (median = 1.00), indicating that GEO-BERT generally grounded its responses in the retrieved evidence while following query-specific requirements. Together, these results show that GEO-BERT produced relevant and evidence-supported answers, with most of the variation occurring in the retrieval of supporting information. We additionally evaluated the same 10 queries using general-purpose LLM systems: Gemini as a representative example. Gemini, ChatGPT etc typically returned substantially fewer GSE accessions than GEO-BERT. A systematic comparison against Gemini further showed that GEO-BERT achieved better retrieval performance on our manually curated benchmark, supporting the value of a domain-specific retrieval framework for semantic discovery of GEO datasets (Figure 3D).

Overall, GEO-BERT demonstrates the ability to retrieve semantically relevant datasets and outperforms general-purpose systems as a domain-specific retrieval framework.

## Methods

### Data Collection

GEO Series (GSE) data was collected from the NCBI Gene Expression Omnibus (GEO) database randomly taking 4331, 371, 273, 310, 323, 238, 203 GSEIDs from 2026 to 2020 respectively, yielding 6049 GSEIDs from GSE accessions in total. For each discovered GSE, we downloaded its SOFT metadata file directly from GEO over HTTPS, then parsed the file using GEOparse [] to extract study-level metadata (title, summary, organism, platform, design) and sample-level (GSM) metadata (characteristics, protocols, library/sequencing details). After indexing, a total of 42223 study vectors (1.5 GB) and 174219 sample vectors (2.1 GB) have been stored in Qdrant DB.

### Overview of GEO-BERT

GEO-BERT is a Retrieval-Augmented Generation (RAG) system designed for natural language querying and discovering genomics datasets from the Gene Expression Omnibus (GEO) repository. The system is organized into four sequential stages:

1. Data ingestion and indexing,
2. Query understanding and retrieval preparation,
3. Hybrid retrieval, and
4. Evidence grounded answer generation.

### Data Ingestion and Vector Indexing

GEO-BERT operates on structured JSON representations of GEO records exported from NCBI’s Gene Expression Omnibus. Each GEO Series (GSE) study is represented as a JSON file containing descriptive sections having a title, study summary, experimental design description, laboratory protocols, and data processing steps alongside structured metadata fields such as organism, tissue, assay type, library strategy, and sequencing platform. Each GEO Sample (GSM) is represented as a corresponding JSON file containing per-sample biological metadata (tissue, organism, disease state, genotype, cell line, cell type, strain) and technical metadata along with experimental condition descriptors and extra metadata fields captured from GEO’s raw submission records.

The two document types are processed and indexed separately into dedicated vector collections: one for study-level documents and one for sample-level documents. This separation reflects their fundamentally different information types. GEO-BERT uses Qdrant as its vector store. Each point in both collections carries three components: a) a dense vector for semantic similarity retrieval, b) a sparse vector for lexical retrieval, and c) a structured payload for metadata filtering. a) The dense vector is a 768-dimensional embedding produced by neuml/pubmedbert-base-embeddings, a PubMedBERT model fine-tuned on biomedical literature. PubMedBERT was selected over general-purpose embedding models because its domain-specific pre-training on PubMed abstracts and full-text articles produces substantially better representations for biomedical terminology, assay names, and biological processes (ref1). Embeddings are cosine-normalized before storage, enabling cosine similarity queries via dot product. b) The sparse vector is produced by prithivida/Splade_PP_en_v1, a SPLADE model running through the FastEmbed library. SPLADE (SParse Lexical AnD Expansion) is a learned sparse retrieval model that assigns non-zero weights to semantically informative token positions in a high-dimensional vocabulary space (ref2). Unlike classical BM25, which computes relevance from raw term frequency statistics, SPLADE learns term importance from training data and performs implicit query and document expansion. This enables it to capture keyword-level signals for specific technical terms like gene names, assay types, chemical compounds, sequencing platforms that dense semantic embeddings may not reliably distinguish at the lexical level. c) Payload indexes are applied to all metadata fields to enable efficient pre-filtering before vector search. Identity fields generated by the indexing pipeline itself (gse_id, gsm_id, parent_id, child_index, level, field) are indexed as exact-match KEYWORD types. Biological and technical content fields originating from GEO’s metadata (organism, tissue, disease, library_strategy, assay_type, instrument_model, cell_type, strain, and others) are indexed using a word-level tokenizer with lowercase normalization.

### Study Documents Ingestion

GSE records can contain long textual sections that exceed the input length of the embedding model. Therefore, each GSE is converted into a study document composed of multiple semantic sections rather than treating the complete study as a single text. Each major textual section which we call a field, such as the study title, summary, overall design, experimental protocols, and data-processing information, is first treated as a separate parent section. Each parent section is then divided into smaller overlapping child chunks using the PubMedBERT tokenizer. A chunk size of 256 tokens with a 50-token overlap has been used. This preserves the relationship between a retrieved chunk and its original study section while allowing long study records to be searched efficiently. The resulting child chunks are embedded using both the dense and sparse models and stored as individual points in the study collection. The payload associated with each chunk preserves the GSE identifier, parent-section identifier, child position, section type, and relevant study metadata. This structure allows individual chunks to be retrieved during search and later reconstructed into study-level information.

### Sample Documents Ingestion

GEO sample records are indexed independently of study records. Each sample contains its own biological metadata. Since sample records are typically short, no chunking has been applied; each GSM record is stored as a single vector point. Dynamic metadata fields, including experimental conditions and additional submission-specific attributes, are converted into a standardized key-value text representation to preserve relevant experimental context.

### Query Understanding and Retrieval Preparation

#### Query Understanding

When a user submits a natural-language query, GeoBERT first performs **query understanding** using a single LLM call through the Groq API. The model used for this stage is openai/gpt-oss-120b with temperature 0 and low reasoning effort. The LLM converts the natural-language query into a structured seven-field JSON representation:

1. **Intent**: Primary user goal e.g. find, compare, explain, identify, list, rank, summarize, describe.
2. **Operations**: All actions implied by the query.
3. **Query type**: One of the eight categories-SEMANTIC, METADATA, MULTI CONSTRAINT, STUDY DESIGN, EXPERIMENTAL PROTOCOL, DATA PROCESSING, MODALITY, PLATFORM.
4. **Concepts**: Key biomedical terms for semantic retrieval.
5. **Semantic focus fields**: GEO text sections that are most relevant to the query.
6. **Logical operators**: Used to combine multiple filters (e.g. AND, OR, NOT).
7. **Filters**: Structured metadata constraints.

In the abode structured query JSON, the query type describes the main nature of the information the user is looking for. SEMANTIC represents a general biological query without specific metadata constraints, whereas METADATA focuses primarily on GEO metadata, MULTI CONSTRAINT represents queries containing multiple specific constraints, STUDY DESIGN, EXPERIMENTAL PROTOCOL, and DATA PROCESSING represent queries concerning experimental design, laboratory procedures, and computational processing, respectively, MODALITY represents sequencing or assay technologies, and PLATFORM represents specific sequencing platforms or instruments. Logical operators and filters govern Qdrant metadata retrieval. Semantic focus fields inform field-aware score boosting during candidate ranking. Intent, operations, and concepts are embedded in the answer generation prompt to align the response format and vocabulary with the user’s information need.

The extracted filters are restricted to predefined GEO metadata fields, including organism, tissue, disease, assay type, library strategy, library source, library selection, molecule, instrument model, genotype, cell line, cell type, and status. After extracting structured query filters, GEO-BERT resolves organism terms against the NCBI Taxonomy database to map user-provided expressions to canonical organism names used in GEO metadata. This step is intended to improve metadata-based retrieval and reduce the candidate search space before semantic retrieval. GEO metadata follows SRA controlled vocabulary conventions (NCBI SRA, 2012) that differ systematically from how researchers express the same concepts informally. The query understanding prompt therefore embeds explicit vocabulary mappings. For example, normalizing ‘scRNA-seq’ and ‘single-cell RNA sequencing’ to the SRA-standard value ‘RNA-Seq’ ensures extracted constraints match the values stored in the collection before retrieval is executed.

#### NCBI Taxonomy Organism Resolution

Organism names extracted by the LLM are resolved to canonical scientific names using the NCBI Taxonomy database. This resolution step is necessary because GEO stores organism names in standard scientific names (“Homo sapiens”, “Mus musculus”) while users commonly type informal names (“human”, “mouse”, “norway rat”). Organism resolution reduces filter retrieval failures caused by differences between informal user terminology and canonical organism names stored in GEO metadata. Resolution proceeds through two NCBI Entrez API calls: esearch.fcgi accepts the raw term and returns a taxonomy identifier, and esummary.fcgi accepts the identifier and returns the canonical scientific name and common name. All resolved organism names are cached in a SQLite database that persists across application restarts. After the first successful resolution, every future lookup for the same organism term is served from the local cache with zero network latency. When organism resolution succeeds, the raw user term in the filter dictionary is replaced by the canonical scientific name before the filter is used in retrieval.

#### Retrieval strategy selection

Following query understanding, filter extraction and organism resolution, the extracted filters are validated against the indexed collection. Each filter is checked individually, and a filter that does not match any indexed record is removed rather than causing the entire query to fail. The number of valid filters determines how metadata constraints participate in dense retrieval. HARD FILTER (two or more filters) applies Qdrant metadata filters as hard constraints during dense retrieval. SOFT FILTER (one filter) runs retrieval without hard filtering but rewards matching documents through post-retrieval score adjustments; SEMANTIC ONLY (zero filters) performs pure vector search with no metadata involvement. Along with one of these three dense retrieval, GEO-BERT also performs sparse retrieval and combines dense and sparse retrieval to obtain complementary candidates as described in the hybrid retrieval in the following.

### Hybrid Retrieval

#### Hybrid (Dense-Sparse) Retrieval and fusion

For each query, GEO-BERT generates two query representations. A dense PubMedBERT embedding captures semantic similarity between the query and indexed text, while a SPLADE representation captures lexical and vocabulary-level signals. Dense and sparse retrieval are performed separately for both the study and sample collections. When a retrieval strategy provides valid metadata constraints, the constraints are applied to the dense search according to the selected strategy. Sparse retrieval is performed globally without metadata filtering. The dense and sparse retrieval processes produce separate ranked candidate sets. These sets are merged according to the union of their document identifiers. Because the raw scores produced by dense and sparse retrieval are not directly comparable, each score distribution is independently normalized to the range [0,1] using min-max normalization over the merged candidate set. The resulting hybrid score is calculated as a weighted linear combination of the normalized dense and sparse scores: s_hybrid = α_d x s_dense + α_s x s_sparse, where s_dense and s_sparse denote the normalized scores, and α_d=0.65 and α_ds=0.35 are the dense and sparse fusion weights, respectively.

#### Score Boosting and candidates dropping

After hybrid score calculation, three additional score adjustments are applied. First, a 0.15 organism-match boost is applied when the stored organism matches the normalized organism filter. Second, a 0.10 field-match boost is applied for each matching non-organism metadata filter. Third, a 0.15 semantic-focus boost is applied when the retrieved study chunk belongs to a field identified by the query-understanding stage as particularly relevant to the query. After these adjustments, candidates are filtered using a dense similarity threshold of 0.20. Candidates whose dense similarity is below this threshold are removed before reranking. This step prevents candidates with primarily lexical matches but weak semantic similarity from proceeding to the more expensive reranking stage.

#### Cross-Encoder Reranking

The bi-encoder retrieval architecture encodes queries and documents independently, preventing fine-grained interaction modeling between query terms and document terms. The cross-encoder, ms-marco-MiniLM-L-6-v2, processes each query and candidate chunk jointly and produces a relevance score based on their combined representation. Reranking is applied separately to study chunks and sample chunks. The raw output is an unbounded real-valued logit where positive values indicate relevance and negative values indicate irrelevance. After scoring, candidates are sorted by logit descending and the top-k study and sample chunks are retained. The improved interaction modeling comes at a higher computational cost, so cross-encoder scoring is applied only after the candidate set has been reduced through hybrid retrieval, score adjustment, and threshold filtering.

#### Parent-Child Aggregation with Neighbor Expansion for study retrieval

The flat ranked list of top-k study child chunks spans multiple GSE studies and multiple field sections within each study. An aggregation step reconstructs study-level context from child-level retrieval results. Chunks are first grouped by their `gse id` payload field. Within each study group, the child chunk with the highest cross-encoder score for each field type is selected as the representative for that field. This per-field selection eliminates redundancy: when multiple overlapping chunks from the same long field survive retrieval and reranking, only the most informative one contributes to the study’s context block. For each selected best child chunk, the system optionally fetches the immediately adjacent chunks: the predecessor and successor through direct Qdrant scroll queries filtered by `parent id` and `child index`. This neighbor expansion (window size = 1) addresses the boundary fragmentation problem inherent in overlapping chunking: a sentence split across two consecutive chunks may be represented incompletely in the retrieved chunk. Fetching neighbors recovers the surrounding sentence context without requiring the system to rescore all chunks from that parent. The retrieved texts are assembled in reading order preceding neighbor, core chunk, following neighbor to form an expanded field representation. The final context for each study includes all field texts, the full structured metadata, series relationship flags, and a `relevance score` computed as σ(z_max) where z_max is the highest cross-encoder logit observed across all children of that study. Studies are sorted by `relevance_score` descending in the final output.

#### Sample Retrieval

Sample retrieval follows the same hybrid retrieval with fusion scoring, score boosting and candidate dropping, and cross-encoder reranking process. However, sample retrieval is performed independently against the sample collection. Because each GSM is already stored as a single sample document, no parent-child aggregation is required. After cross-encoder reranking, the highest-ranked GSM documents directly constitute the final sample context.

### Answer Generation

#### Final Context Formatting

The retrieved study and sample-level context is provided to the answer-generation model. Study context is presented in a consistent field order, prioritizing the study title, summary, experimental design, protocols, and data-processing information. Selected study-level metadata, including organism, tissue, assay type, library strategy, instrument, and BioProject information, are also provided. Sample-level evidence is presented separately and grouped by their GSE identifier. Each retrieved GSM record is represented using concise metadata describing available attributes such as tissue, disease, genotype, cell line, and cell type.

#### Answer Generation Prompt Design

The answer-generation model uses the openai/gpt-oss-120b model through Groq with a temperature of 0.1 and a maximum generation budget of 1,800 tokens. The prompt separates the query, query interpretation, final study context, final sample context, and generation instructions using structured XML-style blocks. The generation process follows four prompt-guided steps: understanding the information needed, synthesizing the final study and sample contexts, generating a concise answer, and verifying factual claims against the final contexts.

The generator is instructed not to invent GSE/GSM identifiers, sample counts, experimental details, or other information not supported by the final contexts. When retrieval validation removes one or more filters, these unresolved filters are provided to the generator so that the response can distinguish exact matches from the closest available matches. In addition, when the highest-ranked study has a normalized relevance score below 0.50, the generator is instructed to communicate the possibility of partial matching rather than presenting the retrieved studies with unwarranted certainty. The resulting response is therefore generated from the final study context, final sample context, structured query interpretation, and unresolved retrieval constraints, providing an evidence-grounded answer to the user’s GEO dataset query.

### Manually curated reference Datasets

For each query, we collected the GSE accessions retrieved by GEO, GEO-BERT, Gemini, and combined them into a nonredundant candidate set. Each candidate GSE was then manually reviewed using its GEO record and associated metadata/JSON files and assigned a relevant (TRUE) or non-relevant (FALSE) label for the corresponding query (Supplementary Table 3). Because the four systems retrieved substantially non-overlapping sets of GSE accessions, pooling their results produced a diverse collection of relevant and non-relevant candidates for evaluation. In the absence of a comprehensive ground-truth list of all relevant GSE accessions for each query, this pooled-retrieval and manual-curation strategy provided a practical reference set for comparing retrieval performance across systems.

### GRO-BERT app

The frontend is implemented in Next.js 16 with React 19, TypeScript, and Tailwind CSS. The choice of Next.js provides server-side rendering capability, file-based routing, and a mature production build pipeline. shadcn/ui components were used for accessible, theme-consistent UI primitives.

The dashboard follows an information hierarchy matching the retrieval architecture: aggregate statistics (dataset count, sample count, platform distribution) appear at the top, ranked study cards populate the central panel, and the conversational answer occupies the bottom section. This layout ensures a researcher can assess retrieval quality (stat bar, quality panel) while reading the generated answer simultaneously.

Dark and light themes are supported via next-themes with system preference as the default. JWT authentication tokens are stored in localStorage with client-side guard on the main page via GET /auth/me verification on mount, redirecting to login on expiry. Session history with result snapshot persistence enables users to navigate between past query results within a session without re-executing the pipeline.

## Discussion

Identifying datasets that satisfy specific scientific requirements remains a substantial challenge for data-driven research. Although GEO provides access to a large collection of functional genomics datasets, its heterogeneous organization and metadata can make it difficult to identify datasets appropriate for secondary analysis, hypothesis testing, and independent data reuse. GEO-BERT addresses this challenge by enabling researchers to search GEO based on scientific intent rather than relying primarily on conventional keyword-based retrieval.

Considerable efforts have focused on improving the organization, standardization, and annotation of GEO metadata. GEO-BERT provides a complementary approach by facilitating dataset discovery within the existing metadata landscape, without requiring complete restructuring of the underlying resource. This is particularly relevant because GEO metadata are neither fully structured nor entirely free text; rather, they combine structured fields, keywords, short annotations, and lengthy free-text descriptions. Such heterogeneity makes dataset retrieval difficult when the biological requirements of a query are distributed across multiple metadata elements or expressed using terminology different from that used by the researcher.

Our results demonstrate that a retrieval-augmented generation (RAG) framework can effectively support semantic dataset discovery in this setting. By combining domain-specific semantic representations with retrieval and evidence-based response generation, GEO-BERT retrieved a broader set of relevant datasets than general-purpose LLM systems in our evaluation while providing information about how individual datasets matched the query. This evidence-grounded design also reduced unsupported responses relative to general-purpose chatbot searches, where we observed instances of incorrect or unverifiable dataset recommendations. Because systems such as Gemini and ChatGPT are not specifically designed for comprehensive GEO retrieval, they also tended to return relatively small numbers of candidate datasets. In contrast, GEO search frequently returned substantially larger result sets, but many datasets were only partially relevant to the intended query, requiring additional manual review to identify appropriate datasets. GEO-BERT provides a middle ground by returning a manageable number of top-ranked candidate datasets while also providing dataset-level relevance scores and information describing how each result matches the query. This additional context allows users to more readily assess the relevance of retrieved datasets and prioritize those most appropriate for further investigation.

The current implementation has several practical limitations. In particular, use of freely available LLM services imposes token and request constraints that can limit the number of retrieved datasets processed during a single query. These limitations are primarily implementation-dependent and could be reduced through access to models with larger context and usage limits or through alternative local or hosted inference strategies. More broadly, retrieval performance remains dependent on the quality and completeness of the underlying GEO metadata, and the manually curated reference sets used in our evaluation may not capture every relevant dataset.

The successful application of GEO-BERT also suggests opportunities beyond GEO. Rapidly growing resources for proteomics, CRISPR screens, and other high-throughput molecular data may face related challenges involving heterogeneous metadata, free text description of the experimental design and protocols, and increasing dataset volume etc. More broadly, the GEO-BERT architecture could be adapted to other biomedical data repositories, including those containing proteomics, CRISPR-screening, and other high-throughput molecular datasets, where semantic dataset discovery could facilitate data reuse. Domain-specific retrieval systems that combine semantic search with evidence-grounded language models could provide a general strategy for improving discovery and reuse across such biomedical data resources.

## Declaration of interests

The authors declare no competing interests.

## Funding

This work was supported in part by the intramural program, Informatics Research (*ZIC TR000410*) at the National Center for Advancing Translational Sciences, part of the National Institutes of Health.

## Disclaimer

The contributions of the NIH author(s) were made as part of their official duties as NIH federal employees, are in compliance with agency policy requirements, and are considered Works of the United States Government. However, the findings and conclusions presented in this paper are those of the author(s) and do not necessarily reflect the views or policies of the Department of Health and Human Services, nor does mention of trade names, commercial products, or organizations imply endorsement by the U.S. Government.

## Contributions

Conceptualization and Supervision – MS; Data curation, Formal analysis, Investigation – MIS,MAR,NA,MS; Methodology – MS,MIS; Resources – EM,SA; Software – MIS,MAR,NA,MS; Validation – MS,MIS,MAR; Visualization – MAR,MS; Writing - original draft– MIS,MAR,NA,MS; Writing – review & editing - All authors.

## Notes

### Competing Interest Statement

The authors have declared no competing interest.

