## Supplemental table1-5 for "GEO-BERT: A Retrieval augmented LLM Framework for Intelligent Dataset Retrieval from the Gene Expression Omnibus"

Table S1: List of GSEIDS loaded for benchmarking

| GSE ID | Organism | Public on | Bioproject | Tissue |
|---|---|---|---|---|
| GSE302018 | Mus musculus | Jan 06 2026 | | lung |
| GSE298118 | Homo sapiens | Jan 26 2026 | PRJNA1268393 | |
| GSE249173 | Mus musculus | Feb 18 2026 | PRJNA1047597 | ureter |
| GSE319173 | Mus musculus | Feb 15 2026 | PRJNA1422503 | hippocampus |
| GSE277320 | Arabidopsis thaliana | Feb 18 2026 | PRJNA1161709 | Leaf |
| GSE196097 | Homo sapiens | Jul 13 2022 | PRJNA803241 | retina |
| GSE262835 | Homo sapiens | Feb 23 2026 | PRJNA1093453 | |
| GSE299179 | Mus musculus | Jan 23 2026 | PRJNA1273027 | Bone marrow |
| GSE307765 | Mus musculus | Jan 28 2026 | PRJNA1327287 | Lung |
| GSE236027 | Homo sapiens | Aug 04 2023 | PRJNA988462 | |
| GSE161735 | Mus musculus | Feb 26 2021 | PRJNA679270 | |
| GSE253551 | Homo sapiens;Mus musculus | Jan 18 2026 | PRJNA1066259 | Adult Heart |
| GSE273676 | Mus musculus | Feb 28 2026 | PRJNA1142601 | E14.5 interneuron |
| GSE282025 | Mus musculus | Feb 09 2026 | PRJNA1186672 | Lung |
| GSE310171 | Homo sapiens | Jan 20 2026 | PRJNA1365228 | lung cancer |
| GSE198650 | Homo sapiens | Mar 18 2022 | PRJNA816334 | |
| GSE153053 | Mus musculus | Jun 24 2020 | PRJNA641357 | Retina |
| GSE312958 | Mus musculus | Jan 13 2026 | PRJNA1377658 | FL-BMDC |
| GSE291367 | Homo sapiens | Jan 14 2026 | PRJNA1233251 | |
| GSE298271 | Rattus norvegicus | Jan 14 2026 | PRJNA1268779 | brain |
| GSE299572 | Arabidopsis thaliana | Jan 18 2026 | PRJNA1274995 | Plant leaves |
| GSE176035 | Homo sapiens | Aug 01 2022 | PRJNA734617 | |
| GSE315552 | Mus musculus | Feb 28 2026 | PRJNA1398252 | Mammary tissue |
| GSE317864 | Mus musculus | Feb 12 2026 | PRJNA1415732 | small intestine |
| GSE309910 | Canis lupus familiaris | Jan 03 2026 | PRJNA1363980 | |
| GSE273528 | Homo sapiens | Jan 26 2026 | PRJNA1142049 | lung adenocarcinoma |
| GSE286178 | Homo sapiens | Jan 07 2026 | PRJNA1208064 | |
| GSE221045 | Homo sapiens | Aug 11 2023 | PRJNA912662 | |
| GSE303327 | Mus musculus | Jan 01 2026 | PRJNA1294836 | Bone marrow |
| GSE240218 | Homo sapiens | Feb 24 2026 | PRJNA1002995 | Peripheral blood |
| GSE281295 | Homo sapiens | Feb 24 2026 | PRJNA1183338 | |
| GSE269377 | Mus musculus | Jan 08 2026 | PRJNA1121300 | mouse spinal cord |
| GSE256145 | Homo sapiens | Feb 20 2026 | PRJNA1078324 | |
| GSE298759 | Mus musculus | Feb 18 2026 | PRJNA1271060 | colon |
| GSE292846 | Homo sapiens | Feb 12 2026 | PRJNA1241731 | |
| GSE282131 | Homo sapiens | Jan 02 2026 | PRJNA1187257 | |
| GSE141019 | Gallus gallus | Apr 17 2020 | PRJNA591782 | retina |
| GSE314779 | Saccharomyces cerevisiae | Jan 08 2026 | PRJNA1392577 | |
| GSE297859 | Mus musculus | Feb 13 2026 | PRJNA1266606 | colon tissue |
| GSE186152 | Arabidopsis thaliana | Dec 12 2022 | PRJNA772746 | Seedling |
| GSE307707 | Mus musculus | Feb 10 2026 | PRJNA1327130 | Lung |
| GSE314638 | Mus musculus | Jan 20 2026 | PRJNA1392356 | |
| GSE292907 | Homo sapiens | Feb 12 2026 | PRJNA1241844 | |
| GSE317912 | Homo sapiens | Jan 31 2026 | PRJNA1415799 | melanoma |
| GSE270909 | Mus musculus | Jan 12 2026 | PRJNA1128817 | spleen |
| GSE316780 | Homo sapiens | Jan 21 2026 | PRJNA1404189 | |
| GSE307065 | Caenorhabditis elegans | Feb 25 2026 | PRJNA1314203 | whole animal |
| GSE315346 | Rattus | Jan 04 2026 | PRJNA1395856 | Dorsal root ganglia |
| GSE276441 | Homo sapiens | Dec 24 2024 | PRJNA1157381 | |
| GSE286039 | Homo sapiens | Jan 06 2026 | PRJNA1207533 | peripheral blood |
| GSE282725 | Mus musculus | Feb 21 2026 | PRJNA1190076 | Prostate |
| GSE215957 | Mus musculus | Feb 16 2026 | PRJNA891342 | lung |
| GSE102485 | Homo sapiens | Aug 10 2020 | PRJNA397857 | anch retinal vein occlusion |
| GSE246944 | Homo sapiens | Sep 30 2025 | PRJNA1035136 | |
| GSE134871 | Mus musculus | Mar 10 2020 | PRJNA556669 | Rod photoreceptors |
| GSE287058 | Homo sapiens | Jan 15 2026 | PRJNA1211075 | Cell Culture |
| GSE283744 | Homo sapiens | Jan 15 2026 | PRJNA1195124 | Blood |
| GSE310752 | Mus musculus | Jan 09 2026 | PRJNA1367028 | erior dorsal cerebellar vermis |
| GSE294726 | Homo sapiens | Jan 14 2026 | PRJNA1251514 | ed human broncial epithelium |
| GSE283894 | Mus musculus | Feb 12 2026 | PRJNA1195980 | |
| GSE311051 | Homo sapiens | Feb 13 2026 | PRJNA1368605 | |

| | | | | |
|---|---|---|---|---|
| GSE293872 | Homo sapiens | Feb 19 2026 | PRJNA1246633 | Bone Marrow |
| GSE316867 | Homo sapiens | Jan 31 2026 | PRJNA1405455 | glioma |
| GSE293681 | Mus musculus | Jan 21 2026 | PRJNA1245684 | Brown adipose tissue |
| GSE297482 | Homo sapiens | Feb 11 2026 | PRJNA1264984 | |
| GSE277503 | Homo sapiens | Sep 15 2025 | PRJNA1162532 | retina |
| GSE272003 | Homo sapiens | Jan 20 2026 | PRJNA1134608 | |
| GSE319750 | Homo sapiens | Feb 20 2026 | PRJNA1425046 | diffuse gastric cancer |
| GSE294073 | Danio rerio | Jan 21 2026 | PRJNA1247998 | Heart ventricle |
| GSE160140 | Danio rerio | Oct 27 2020 | PRJNA672101 | |
| GSE306127 | Homo sapiens | Feb 24 2026 | PRJNA1309493 | |
| GSE273700 | Homo sapiens | Jan 28 2026 | PRJNA1142876 | PBMC |
| GSE315989 | Polypterus senegalus | Jan 15 2026 | PRJNA1400066 | pectoral fin (intact) |
| GSE287453 | Glycine max | Jan 19 2026 | PRJNA1213028 | hairy roots |
| GSE295012 | Homo sapiens | Jan 07 2026 | PRJNA1252451 | skin |
| GSE300783 | Candidozyma auris | Jan 06 2026 | PRJNA1282106 | |
| GSE286432 | Homo sapiens | Feb 07 2026 | PRJNA1209136 | |
| GSE291211 | Mus musculus | Feb 11 2026 | PRJNA1232205 | Breast tumor |
| GSE318731 | Mus musculus | Feb 16 2026 | PRJNA1419852 | Mouse cervical spinal cord |
| GSE165322 | Homo sapiens | Mar 05 2021 | PRJNA694104 | |
| GSE309256 | coccus elongatus PCC 7942 = FA | Feb 23 2026 | PRJNA1334990 | |
| GSE259256 | Homo sapiens;Mus musculus | Feb 10 2025 | PRJNA1080533 | cell |
| GSE305990 | Mus musculus | Jan 12 2026 | PRJNA1308851 | |
| GSE317322 | Homo sapiens | Jan 22 2026 | PRJNA1406736 | human |
| GSE293356 | Drosophila melanogaster | Feb 12 2026 | PRJNA1244244 | First tarsal segment |
| GSE165831 | Mus musculus | Dec 22 2021 | PRJNA697989 | retina |
| GSE287822 | Rattus norvegicus | Jan 23 2026 | PRJNA1214774 | hippocampal |
| GSE308555 | Homo sapiens | Jan 01 2026 | PRJNA1330874 | cell |
| GSE299956 | Mus musculus | Jan 20 2026 | PRJNA1277721 | telencephalon |
| GSE193429 | Homo sapiens | Jan 14 2026 | PRJNA796204 | |
| GSE146641 | Homo sapiens | Sep 05 2022 | PRJNA611569 | retina organoid |
| GSE197148 | Homo sapiens | Feb 25 2022 | PRJNA809208 | |
| GSE278881 | Mus musculus | Aug 18 2025 | PRJNA1169351 | retina |
| GSE317440 | Homo sapiens | Jan 28 2026 | PRJNA1412785 | bone marrow |
| GSE280985 | Mus musculus | Feb 17 2026 | PRJNA1181780 | liver |
| GSE261696 | Homo sapiens | Jan 31 2026 | PRJNA1088438 | circulating blood |
| GSE299792 | Homo sapiens | Jan 09 2026 | PRJNA1276530 | |
| GSE319287 | Homo sapiens | Feb 22 2026 | PRJNA1422777 | adrenal cortex |
| GSE281042 | Danio rerio | Jun 24 2025 | PRJNA1181875 | eye |
| GSE192602 | Mus musculus | Apr 27 2022 | PRJNA792267 | |
| GSE161394 | Mus musculus | Dec 30 2020 | PRJNA678013 | |
| GSE143669 | Homo sapiens | Jan 15 2020 | PRJNA601209 | |
| GSE247315 | Mus musculus | Dec 22 2023 | PRJNA1037099 | |
| GSE260927 | Homo sapiens | Jan 27 2026 | PRJNA1084015 | |
| GSE316257 | Mus musculus | Jan 25 2026 | PRJNA1401891 | nor-infiltrating lymphocytes |
| GSE239456 | Mus musculus | Aug 01 2023 | PRJNA999329 | neuro retina |
| GSE263470 | Homo sapiens | Jun 30 2026 | PRJNA1097596 | retina |
| GSE242229 | Macaca fascicularis | Sep 06 2023 | PRJNA1012311 | retina |
| GSE315862 | Homo sapiens | Feb 12 2026 | PRJNA1399455 | |
| GSE269083 | Homo sapiens | Feb 25 2026 | PRJNA1120074 | kidney |
| GSE312736 | Porphyridium purpureum | Jan 14 2026 | PRJNA1375806 | |
| GSE279065 | Homo sapiens | Feb 12 2026 | PRJNA1170278 | |
| GSE278279 | Arabidopsis thaliana | Jan 05 2026 | PRJNA1166215 | Shoot |
| GSE210744 | Drosophila melanogaster | Apr 18 2023 | PRJNA867354 | phila third instar larvae eye disc |
| GSE293275 | Mus musculus | Feb 05 2026 | PRJNA1243467 | Pancreatic cancer |
| GSE297569 | Mus musculus | Feb 17 2026 | PRJNA1265204 | small intestine |
| GSE304621 | Macaca mulatta | Mar 30 2026 | PRJNA1301929 | Retina |
| GSE237291 | Mus musculus | Aug 15 2023 | PRJNA994535 | etinal pigment epithelium |
| GSE299875 | Homo sapiens | Jan 21 2026 | PRJNA1277213 | e-etinal pigment epithelium |
| GSE302914 | Mus musculus | Jan 01 2026 | PRJNA1292817 | testis |
| GSE309899 | Homo sapiens | Feb 20 2026 | PRJNA1363972 | |
| GSE296017 | Mus musculus | Feb 05 2026 | PRJNA1257321 | skeletal muscles |
| GSE314834 | Homo sapiens | Jan 09 2026 | PRJNA1392905 | |
| GSE302085 | Homo sapiens | Jan 14 2026 | PRJNA1288495 | Intestine |

| | | | | |
|---|---|---|---|---|
| GSE305701 | | Jan 08 2026 | | |
| GSE178312 | Mus musculus | Jun 17 2021 | PRJNA738469 | retina |
| GSE293149 | Caenorhabditis elegans | Feb 19 2026 | PRJNA1242888 | bulk |
| GSE316321 | Homo sapiens | Jan 28 2026 | PRJNA1402299 | Uterus, Cervix |
| GSE223657 | Mus musculus | Nov 08 2023 | PRJNA927051 | Retina |
| GSE314786 | Homo sapiens | Jan 30 2026 | PRJNA1392604 | |
| GSE287840 | Homo sapiens | Jan 23 2026 | PRJNA1214799 | |
| GSE150703 | Mus musculus | Aug 20 2020 | PRJNA633178 | |
| GSE279113 | Homo sapiens | Dec 31 2024 | PRJNA1170898 | retina |
| GSE301260 | Rattus norvegicus | Feb 18 2026 | PRJNA1284349 | brain |
| GSE309203 | Homo sapiens | Feb 26 2026 | PRJNA1334328 | lung |
| GSE285223 | Mus musculus | Jan 09 2026 | PRJNA1202066 | |
| GSE235022 | Danio rerio | Aug 01 2023 | PRJNA984201 | retina |
| GSE288777 | Mus musculus | Jan 21 2026 | PRJNA1219240 | Lung |
| GSE317377 | Mus musculus | Jan 26 2026 | | inguinal white adipose tissue (iWAT) |
| GSE305054 | Mus musculus | Feb 25 2026 | PRJNA1304372 | |
| GSE334496 | Homo sapiens | Jun 08 2026 | PRJNA1475347 | |
| GSE301748 | Macaca mulatta | Feb 26 2026 | PRJNA1287516 | Bone marrow |
| GSE228590 | Mus musculus | Apr 07 2023 | PRJNA950412 | |
| GSE292400 | Mus musculus | Feb 26 2026 | PRJNA1238318 | Bone marrow |
| GSE234721 | Drosophila melanogaster | Feb 24 2026 | PRJNA982815 | Ovary |
| GSE279646 | Homo sapiens | Feb 25 2026 | PRJNA1173760 | |
| GSE318284 | Mus musculus | Feb 05 2026 | PRJNA1418527 | Bone Marrow |
| GSE317415 | Mus musculus | Jan 23 2026 | PRJNA1412566 | Placenta |
| GSE278145 | Homo sapiens | Jan 26 2026 | PRJNA1165764 | Nasal mucosa |
| GSE266031 | Homo sapiens | Sep 25 2025 | PRJNA1105369 | 3D cerebral organoid |
| GSE146176 | Mus musculus | Mar 03 2020 | PRJNA609651 | Retina |
| GSE289716 | Arabidopsis thaliana | Feb 15 2026 | PRJNA1224247 | Root |
| GSE300168 | Mus musculus | Jan 02 2026 | PRJNA1279166 | Heart |
| GSE276696 | Homo sapiens | Mar 10 2025 | PRJNA1158714 | eye |
| GSE293220 | Mus musculus | Jan 01 2026 | PRJNA1243370 | lung |
| GSE279007 | Mus musculus | Feb 20 2026 | PRJNA1169907 | Heart ventricle |
| GSE315191 | Mus musculus | Jan 14 2026 | PRJNA1395102 | lung |
| GSE168582 | Mus musculus | Feb 27 2026 | PRJNA708325 | |
| GSE242754 | Homo sapiens | Feb 08 2024 | | |
| GSE316557 | Homo sapiens | Jan 20 2026 | PRJNA1403378 | liver |
| GSE299820 | Homo sapiens | Jan 20 2026 | PRJNA1276569 | Whole Blood |
| GSE203619 | Gallus gallus | May 19 2025 | PRJNA841600 | |
| GSE137935 | Mus musculus | Mar 10 2020 | PRJNA573908 | |
| GSE315800 | Histoplasma ohiense | Jan 26 2026 | PRJNA1399334 | |
| GSE285462 | Homo sapiens | Jan 01 2026 | PRJNA1203672 | PB |
| GSE268080 | Homo sapiens | Jan 01 2026 | PRJNA1114624 | |
| GSE317029 | Mus musculus | Feb 25 2026 | PRJNA1405688 | Spleen |
| GSE175752 | Homo sapiens | Sep 27 2021 | PRJNA733482 | |
| GSE289602 | Mus musculus | Feb 08 2026 | | bone marrow |
| GSE283593 | Homo sapiens | Feb 17 2026 | PRJNA1194590 | Osteosarcoma |
| GSE288890 | Mus musculus | Feb 05 2026 | PRJNA1219754 | dorsal root ganglion |
| GSE308376 | Homo sapiens | Jan 29 2026 | PRJNA1330304 | |
| GSE298876 | Homo sapiens | Feb 19 2026 | PRJNA1271543 | |
| GSE192509 | Mus musculus | Aug 16 2022 | PRJNA791672 | Retina |
| GSE304121 | Homo sapiens | Feb 11 2026 | PRJNA1299299 | |
| GSE214512 | Homo sapiens | Oct 04 2022 | PRJNA885786 | |
| GSE256099 | Mus musculus | Feb 18 2026 | PRJNA1078247 | |
| GSE255140 | Mus musculus | Jan 28 2026 | PRJNA1073604 | spleen |
| GSE308414 | Mus musculus | Jan 22 2026 | PRJNA1330372 | |
| GSE267144 | Mus musculus | Jan 18 2026 | PRJNA1109847 | |
| GSE280637 | Mus musculus | Feb 12 2026 | PRJNA1179862 | ovary |
| GSE303624 | Mus musculus | Feb 13 2026 | PRJNA1295943 | ankle synovium |
| GSE272200 | Homo sapiens | Feb 19 2026 | PRJNA1135666 | Thalamus |
| GSE216208 | Mus musculus | Oct 25 2022 | PRJNA892591 | retina |
| GSE318532 | Mus musculus | Feb 10 2026 | PRJNA1419183 | lung |
| GSE315066 | Mus musculus | Feb 18 2026 | PRJNA1394952 | cell line |
| GSE168475 | Macaca mulatta | Mar 09 2021 | PRJNA707516 | retina |

| | | | | |
|---|---|---|---|---|
| GSE315579 | Homo sapiens | Jan 09 2026 | PRJNA1398286 | F2-mutated schwannoma |
| GSE289097 | Mus musculus | Feb 09 2026 | PRJNA1221532 | TG |
| GSE216720 | Homo sapiens | Sep 18 2024 | PRJNA895054 | colon |
| GSE212523 | Mus musculus | Aug 18 2024 | PRJNA875897 | Retina |
| GSE233896 | Danio rerio | Aug 28 2023 | PRJNA978532 | whole retina |
| GSE286319 | Homo sapiens | Feb 07 2026 | PRJNA1208669 | |
| GSE290239 | Mus musculus | Feb 26 2025 | | retina |
| GSE261617 | Mus musculus | Mar 31 2025 | PRJNA1088121 | Skeletal muscle |
| GSE298981 | Homo sapiens | Jan 20 2026 | PRJNA1272051 | |
| GSE303613 | Mus musculus | Jan 28 2026 | PRJNA1295930 | colon |
| GSE315733 | Homo sapiens | Jan 30 2026 | PRJNA1398858 | |
| GSE216603 | Mycobacterium tuberculosis | Nov 30 2022 | PRJNA894495 | |
| GSE255194 | Homo sapiens | Feb 06 2026 | PRJNA1073925 | |
| GSE304388 | Mus musculus | Feb 17 2026 | PRJNA1301236 | E9.5 placenta |
| GSE305194 | Homo sapiens | Nov 02 2025 | PRJNA1304574 | Retinal Organoid |
| GSE169262 | Mus musculus | Nov 16 2021 | PRJNA715918 | |
| GSE208777 | Mus musculus | Nov 02 2022 | PRJNA860983 | |
| GSE161236 | Homo sapiens | Sep 30 2021 | PRJNA675959 | Endothelial cells |
| GSE294119 | Mus musculus | Jan 20 2026 | | skin |
| GSE283547 | Mus musculus | Jan 08 2026 | PRJNA1194507 | macrophages |
| GSE244882 | Homo sapiens | Jul 19 2024 | PRJNA1025270 | |
| GSE315610 | Mus musculus | Jan 09 2026 | PRJNA1398327 | Spleen |
| GSE291664 | Mus musculus | Mar 02 2026 | PRJNA1234616 | retina |
| GSE290478 | Homo sapiens | Feb 11 2026 | PRJNA1228142 | |
| GSE279786 | Mus musculus | Feb 24 2026 | PRJNA1174679 | |
| GSE302995 | Homo sapiens | Sep 22 2025 | PRJNA1293324 | Human retinal organoid |
| GSE296907 | Mus musculus | May 01 2026 | | Retina |
| GSE302004 | Homo sapiens | Jan 01 2026 | PRJNA1288269 | |
| GSE237210 | Rhabdomys pumilio | Oct 01 2023 | PRJNA994172 | Retina |
| GSE201986 | Macaca mulatta | Feb 01 2025 | PRJNA833937 | adrenal gland |
| GSE268229 | Homo sapiens | Jan 16 2026 | PRJNA1115210 | PDA Cell Line |
| GSE208740 | Homo sapiens | Jan 26 2026 | PRJNA860925 | |
| GSE277000 | Mus musculus | Jan 01 2026 | PRJNA1160202 | bulk bone marrow |
| GSE234797 | Mus musculus | Jun 06 2024 | PRJNA983249 | |
| GSE319253 | Homo sapiens | Feb 11 2026 | PRJNA1422690 | Brain |
| GSE289953 | Mus musculus | Feb 18 2026 | PRJNA1225629 | ell-line-derived Xenograft |
| GSE262968 | Mus musculus | Jan 01 2026 | | kidney_cortex |
| GSE291070 | Homo sapiens | Jan 14 2026 | PRJNA1231501 | prostate |
| GSE318550 | Homo sapiens | Feb 09 2026 | PRJNA1419216 | Brain |
| GSE288932 | Homo sapiens | Feb 11 2026 | PRJNA1220050 | Differentiating cells |
| GSE266598 | Mus musculus | Feb 05 2026 | PRJNA1107812 | Calvaria |
| GSE209660 | Homo sapiens | Jul 28 2022 | PRJNA862071 | |
| GSE269309 | Mus musculus | Jan 06 2026 | PRJNA1120936 | |
| GSE173701 | Mus musculus | Jan 11 2026 | PRJNA726801 | rtic arch and thoracic aorta |
| GSE315766 | Caenorhabditis elegans | Jan 07 2026 | | whole animal |
| GSE153426 | Solenopsis invicta | Jun 30 2020 | PRJNA642634 | Whole body |
| GSE310251 | Mus musculus | Jan 06 2026 | PRJNA1365518 | T cells |
| GSE317494 | Homo sapiens | Jan 28 2026 | PRJNA1413920 | |
| GSE290539 | Mus musculus | Feb 28 2026 | PRJNA1228588 | Hippocampus |
| GSE164884 | Homo sapiens | Jun 14 2022 | PRJNA692366 | Retinal Organoid |
| GSE286897 | Mus musculus | Feb 25 2026 | PRJNA1210514 | |
| GSE228273 | Chlorocebus sabaeus | Aug 30 2023 | PRJNA949184 | retina/choroid |
| GSE269621 | Homo sapiens | Feb 11 2026 | PRJNA1122780 | xenografts tumor |
| GSE313697 | Mus musculus | Feb 16 2026 | PRJNA1381358 | Cecum |
| GSE309282 | Mus musculus | Feb 09 2026 | PRJNA1335070 | m of first molar extraction socket, gingiva removed |
| GSE241230 | Mus musculus | Feb 01 2025 | PRJNA1006760 | Retina |
| GSE314730 | Homo sapiens | Feb 20 2026 | PRJNA1392511 | liver |
| GSE211187 | Homo sapiens | Aug 14 2023 | PRJNA869448 | |
| GSE272537 | Homo sapiens | Jan 21 2026 | PRJNA1137309 | Breast |
| GSE319264 | Chlamydomonas reinhardtii | Feb 12 2026 | | cell |
| GSE318326 | Homo sapiens | Feb 09 2026 | PRJNA1418555 | Tonsil |
| GSE306252 | Homo sapiens | Jan 21 2026 | | blood |
| GSE268990 | Homo sapiens | Jun 04 2024 | PRJNA1119636 | retinal organoid |

| | | | | |
|---|---|---|---|---|
| GSE168503 | Mus musculus | Mar 15 2021 | PRJNA707576 | retinal cones |
| GSE313947 | Mus musculus | Jan 31 2026 | PRJNA1387407 | 71+ EB-derived erythroid cells, originally derived from ESC |
| GSE319658 | Escherichia coli K-12 | Feb 24 2026 | PRJNA1424934 | strain |
| GSE310410 | Mus musculus | Jan 09 2026 | PRJNA1365904 | Small intestine |
| GSE282544 | Mus musculus | Nov 01 2025 | | retina |
| GSE319347 | Homo sapiens | Feb 16 2026 | | |
| GSE160048 | Homo sapiens;Mus musculus | Nov 21 2020 | PRJNA671700 | Kidney glomerulus |
| GSE303996 | Arabidopsis thaliana | Feb 25 2026 | PRJNA1298601 | Whole root |
| GSE143281 | Mus musculus | Aug 07 2020 | PRJNA599586 | Retina |
| GSE310427 | Streptomyces venezuelae | Feb 11 2026 | PRJNA1365920 | |
| GSE293296 | Mus musculus | Jan 26 2026 | PRJNA1243950 | brain |
| GSE299107 | Homo sapiens | Feb 06 2026 | PRJNA1272592 | |
| GSE168749 | Danio rerio | Oct 06 2021 | PRJNA713817 | |
| GSE312911 | Mus musculus | Jan 08 2026 | PRJNA1377587 | brain |
| GSE319412 | Tabanus nipponicus | Feb 20 2026 | PRJNA1423256 | midgut |
| GSE313783 | Mus musculus | Jan 15 2026 | PRJNA1381590 | Bone marrow |
| GSE271298 | Mus musculus | Jan 13 2026 | PRJNA1130881 | FL-BMDC |
| GSE289683 | Homo sapiens | Feb 18 2026 | PRJNA1223934 | |
| GSE165702 | Mus musculus | Jan 28 2024 | PRJNA695504 | ıoroidal-retinal flat mounts |
| GSE302579 | Drosophila melanogaster | Feb 11 2026 | | head |
| GSE298479 | Homo sapiens | Jan 26 2026 | PRJNA1269531 | kidney tumor |
| GSE317702 | Mus musculus | Jan 26 2026 | PRJNA1414659 | spleen |
| GSE275219 | Mus musculus | Feb 06 2026 | PRJNA1149828 | colonic stem cell |
| GSE153787 | Homo sapiens | Aug 20 2020 | PRJNA658093 | Osteosarcoma cells |
| GSE279551 | Canis lupus familiaris | Jan 20 2026 | PRJNA1173236 | ıe from control dogs_Spiked_0T_0W |
| GSE283790 | Mus musculus | Feb 12 2026 | PRJNA1195586 | mPFC |
| GSE270206 | Homo sapiens | Jan 01 2026 | PRJNA1125409 | PCA34 tumor |
| GSE286290 | Homo sapiens;Mus musculus | Feb 17 2026 | PRJNA1208536 | |
| GSE318802 | Mus musculus | Feb 11 2026 | PRJNA1420637 | Bone-marrow derived |
| GSE307599 | Mus musculus | Feb 18 2026 | PRJNA1322363 | Brown Adipose Tissue |
| GSE306785 | Mus musculus | Feb 11 2026 | PRJNA1312104 | Retina |
| GSE317060 | Rattus norvegicus | Feb 18 2026 | PRJNA1405715 | ileum |
| GSE143992 | Mus musculus | Jan 11 2026 | PRJNA602461 | |
| GSE284637 | Homo sapiens | Dec 16 2025 | PRJNA1199978 | |
| GSE316754 | Mus musculus | Jan 16 2026 | PRJNA1404142 | prostate |
| GSE292720 | Homo sapiens | Mar 28 2025 | PRJNA1241146 | Lung |
| GSE147728 | Shiraia bambusicola | Mar 30 2023 | PRJNA616262 | hypha |
| GSE317548 | Homo sapiens | Jan 30 2026 | PRJNA1414444 | |
| GSE300674 | Mus musculus | Jan 05 2026 | PRJNA1281652 | Liver WT, aged |
| GSE277396 | Mus musculus | Jan 13 2026 | PRJNA1162106 | spinal cord |
| GSE307920 | Homo sapiens | Feb 26 2026 | PRJNA1327817 | |
| GSE232949 | Homo sapiens;Pan troglodytes | May 23 2023 | PRJNA974465 | |
| GSE204933 | Mus musculus | Feb 19 2026 | PRJNA842744 | |
| GSE290387 | Homo sapiens | Feb 25 2026 | PRJNA1227868 | nasal |
| GSE316529 | Homo sapiens | Jan 21 2026 | PRJNA1403280 | |
| GSE294184 | Mus musculus | Jan 22 2026 | PRJNA1248525 | lung |
| GSE309641 | Pseudomonas putida | Feb 26 2026 | PRJNA1336665 | |
| GSE178994 | Homo sapiens | Jul 21 2024 | PRJNA741807 | |
| GSE308703 | Mus musculus | Jan 12 2026 | PRJNA1332566 | thymus |
| GSE305257 | Schizosaccharomyces pombe | Jan 01 2026 | PRJNA1304945 | |
| GSE202725 | Mus musculus | Feb 15 2023 | PRJNA837182 | retina |
| GSE313668 | Homo sapiens | Jan 05 2026 | PRJNA1380504 | |
| GSE149799 | sophila melanogaster;Homo sap | Feb 12 2021 | PRJNA630171 | cortical brain organoids |
| GSE289768 | Mus musculus | Jan 23 2026 | PRJNA1224753 | primary lung tumor |
| GSE311923 | Homo sapiens | Jan 01 2026 | PRJNA1371965 | Synovial tissue |
| GSE267253 | Homo sapiens | Feb 23 2026 | PRJNA1110757 | Spleen |
| GSE297461 | Homo sapiens | Jan 07 2026 | PRJNA1263890 | |
| GSE286384 | Homo sapiens;Mus musculus | Jan 01 2026 | PRJNA1209036 | |
| GSE170172 | Homo sapiens | Mar 29 2021 | | |
| GSE316115 | Mus musculus | Feb 27 2026 | PRJNA1400631 | Lung |
| GSE301229 | Mus musculus | Jan 29 2026 | PRJNA1284302 | Liver |
| GSE287487 | Mus musculus | Jan 25 2025 | PRJNA1213083 | organoids |
| GSE296355 | Homo sapiens | Feb 10 2026 | PRJNA1258972 | |

| | | | | |
|---|---|---|---|---|
| GSE159257 | Mus musculus | Oct 08 2023 | PRJNA668143 | |
| GSE245286 | Homo sapiens | Jan 03 2024 | PRJNA1027764 | |
| GSE315799 | Mus musculus | Jan 19 2026 | PRJNA1399335 | |
| GSE234454 | Homo sapiens | Jan 12 2026 | PRJNA981470 | |
| GSE145836 | Homo sapiens | Oct 23 2025 | PRJNA608488 | man retinal Mueller glial cell |
| GSE313201 | Homo sapiens | Feb 18 2026 | PRJNA1378788 | |
| GSE230348 | Homo sapiens | Jan 18 2024 | PRJNA960647 | Macula Retina |
| GSE223277 | Danio rerio | May 15 2023 | PRJNA925508 | retina |
| GSE285156 | Mus musculus | Jan 26 2026 | PRJNA1201318 | |
| GSE262625 | Canis lupus familiaris | May 15 2024 | PRJNA1092667 | Retina |
| GSE274705 | Homo sapiens | Jan 29 2026 | PRJNA1147681 | Buffy coat |
| GSE319801 | Danio rerio | Feb 23 2026 | PRJNA1425107 | retina |
| GSE300143 | Mus musculus | Feb 10 2026 | PRJNA1279123 | liver |
| GSE270264 | Mus musculus | Mar 05 2025 | PRJNA1125904 | optic nerve |
| GSE292986 | Homo sapiens | Jan 08 2026 | PRJNA1242296 | |
| GSE253585 | Rattus norvegicus | Feb 26 2026 | | Normal mammary tissues |
| GSE223522 | Mus musculus | Jan 28 2023 | PRJNA926634 | Retina |
| GSE118011 | Mus musculus | Jul 29 2022 | PRJNA484001 | retinal ganglion cells |
| GSE180163 | Mus musculus | Jul 15 2022 | PRJNA746811 | neural retinal |
| GSE144860 | Mus musculus | Jan 25 2026 | PRJNA605159 | Liver |
| GSE304377 | Mus musculus | Feb 06 2026 | PRJNA1301136 | Brain |
| GSE274050 | Rattus norvegicus | Feb 10 2026 | PRJNA1144809 | demyelinated neurons |
| GSE301368 | Homo sapiens | Feb 17 2026 | PRJNA1284855 | |
| GSE317248 | Homo sapiens | Jan 27 2026 | PRJNA1406621 | kidney |
| GSE283936 | Mus musculus | Jan 08 2026 | PRJNA1196353 | Colon tumor |
| GSE316229 | hallii var. filipes;Panicum hallii v | Jan 16 2026 | | Leaf |
| GSE317035 | Mus musculus | Feb 20 2026 | PRJNA1405698 | Lung, medLN, Spleen |
| GSE155469 | Solenopsis japonica | Aug 01 2020 | PRJNA649847 | Whole body |
| GSE301884 | Homo sapiens | Jan 24 2026 | PRJNA1287686 | uman embryonic stem cell |
| GSE312374 | Danio rerio | Jan 14 2026 | PRJNA1373192 | heart |
| GSE313168 | Mus musculus | Jan 28 2026 | PRJNA1378732 | Tumor |
| GSE251754 | Homo sapiens | Jul 13 2024 | PRJNA1055164 | |
| GSE316468 | Mus musculus | Jan 21 2026 | | tumor |
| GSE308003 | Mus musculus | Jan 13 2026 | PRJNA1328501 | testis |
| GSE298292 | Mus musculus | Jan 21 2026 | PRJNA1268901 | thymus |
| GSE307692 | Homo sapiens | Jan 05 2026 | PRJNA1322536 | connective tissue |
| GSE312416 | Mus musculus | Feb 22 2026 | PRJNA1373268 | Colon |
| GSE249915 | Homo sapiens | Jan 30 2026 | PRJNA1051131 | primary human breast |
| GSE301036 | Homo sapiens | Feb 09 2026 | PRJNA1283110 | |
| GSE213933 | Homo sapiens | Nov 02 2022 | PRJNA883213 | |
| GSE318298 | Mus musculus | Feb 05 2026 | PRJNA1418551 | bone marrow |
| GSE243315 | Homo sapiens | Dec 31 2024 | PRJNA1017802 | retina |
| GSE202747 | Homo sapiens | Jun 08 2023 | PRJNA837204 | neural retina |
| GSE220792 | Homo sapiens | Aug 08 2023 | PRJNA911333 | Retinal Organoids |
| GSE222457 | Homo sapiens | Jan 01 2026 | PRJNA922137 | ediatric high grade glioma |
| GSE213095 | Homo sapiens | Feb 22 2026 | PRJNA879087 | Peripheral blood |
| GSE159235 | Homo sapiens | Dec 29 2020 | PRJNA668115 | |
| GSE185215 | Mus musculus | Jan 10 2024 | PRJNA768277 | Cortex |
| GSE141798 | Homo sapiens | Apr 14 2021 | PRJNA594743 | |
| GSE315099 | Homo sapiens | Jan 01 2026 | PRJNA1394976 | |
| GSE318943 | Mus musculus | Feb 18 2026 | PRJNA1420849 | |
| GSE202012 | Mus musculus | Jun 15 2022 | PRJNA834042 | Retina |
| GSE315586 | Homo sapiens | Jan 05 2026 | PRJNA1398292 | DLPFC_BA46 |
| GSE210331 | Homo sapiens | Aug 05 2022 | PRJNA865289 | |
| GSE275358 | Mus musculus | Feb 26 2026 | PRJNA1150536 | Muscle |
| GSE285717 | Mus musculus | Jan 01 2026 | PRJNA1205701 | Kidney |
| GSE267564 | Homo sapiens | Jan 24 2026 | PRJNA1111867 | ICM |
| GSE270658 | Mus musculus | Nov 13 2025 | PRJNA1127641 | RPE-choroidal complex |
| GSE281778 | Homo sapiens;Mus musculus | Feb 09 2026 | PRJNA1185697 | hematopoietic |
| GSE304510 | Mus musculus | Jan 26 2026 | PRJNA1301402 | /A tumor-bearing mice at 7 days after tumor inoculation |
| GSE274637 | Homo sapiens | Jan 14 2026 | PRJNA1147321 | |
| GSE312430 | Mus musculus | Feb 12 2026 | PRJNA1373289 | Ankle Joint Synovium |
| GSE244978 | Homo sapiens | Feb 25 2026 | PRJNA1026557 | whole blood |

| | | | | |
|---|---|---|---|---|
| GSE202761 | Homo sapiens | Jan 13 2026 | PRJNA837229 | |
| GSE254272 | Mus musculus | Dec 05 2024 | PRJNA1069114 | |
| GSE272902 | Mus musculus | Jan 16 2026 | PRJNA1139209 | mammary gland |
| GSE253839 | | Feb 22 2024 | | |
| GSE284994 | Oryctolagus cuniculus | Feb 11 2026 | PRJNA1200759 | lung |
| GSE310097 | Streptomyces venezuelae | Feb 11 2026 | PRJNA1365127 | |
| GSE288252 | Homo sapiens | Jan 27 2026 | PRJNA1216629 | |
| GSE314294 | Homo sapiens | Jan 05 2026 | PRJNA1390673 | breast |
| GSE275656 | Mus musculus | Feb 11 2026 | PRJNA1152513 | Retina |
| GSE243451 | Drosophila melanogaster | Feb 27 2026 | PRJNA1018520 | embryo |
| GSE310588 | Methylosinus trichosporium OB3 | Jan 03 2026 | PRJNA1366542 | Bacterial cell |
| GSE316151 | Homo sapiens | Jan 12 2026 | PRJNA1401730 | |
| GSE289551 | Rattus norvegicus | Jan 01 2026 | PRJNA1223418 | |
| GSE310931 | Homo sapiens | Feb 24 2026 | PRJNA1367564 | |
| GSE289566 | Mus musculus | Jan 28 2026 | PRJNA1223449 | |
| GSE298283 | Homo sapiens;Mus musculus | Feb 09 2026 | PRJNA1268878 | e marrow hematopoietic cells |
| GSE152872 | Mus musculus | Dec 31 2023 | PRJNA640673 | |
| GSE303480 | Mus musculus | Jan 05 2026 | PRJNA1295498 | liver |
| GSE285731 | Mus musculus | Jan 01 2026 | PRJNA1205725 | |
| GSE304038 | Mus musculus | Jun 11 2026 | PRJNA1298811 | |
| GSE297370 | Mus musculus | May 17 2025 | PRJNA1263591 | Neural retina |
| GSE318289 | Drosophila melanogaster | Feb 09 2026 | PRJNA1418539 | whole body |
| GSE275103 | Mus musculus | Jan 01 2026 | | Kidney |
| GSE164199 | Homo sapiens | Dec 06 2022 | PRJNA689541 | |
| GSE249904 | Mus musculus | Jan 14 2026 | PRJNA1051114 | bone marrow |
| GSE312889 | Mus musculus | Jan 30 2026 | PRJNA1377555 | |
| GSE215175 | Homo sapiens | May 10 2023 | PRJNA889037 | Pigmented epithelium; RetinaÂ |
| GSE282696 | Homo sapiens | Jan 14 2026 | PRJNA1189813 | Colonic biopsy |
| GSE310812 | Mus musculus | Apr 30 2026 | PRJNA1367108 | RPE |
| GSE300165 | Homo sapiens | Jan 13 2026 | PRJNA1279160 | |
| GSE316045 | Mus musculus | Jan 16 2026 | PRJNA1400516 | Cornea |
| GSE301666 | Mus musculus | Jan 01 2026 | PRJNA1285946 | |
| GSE202886 | Homo sapiens | Jun 08 2023 | PRJNA837596 | neural retina |
| GSE315483 | Oncorhynchus mykiss | Jan 13 2026 | PRJNA1397220 | Visceral adipose tissue |
| GSE272498 | Homo sapiens | Jan 12 2026 | PRJNA1136976 | epithelium |
| GSE300607 | Mus musculus | Jul 16 2025 | PRJNA1281536 | Retina |
| GSE284673 | Mycobacterium tuberculosis | Feb 27 2026 | PRJNA1200020 | |
| GSE227052 | Mus musculus | May 19 2023 | PRJNA942963 | retina |
| GSE286938 | Rattus norvegicus | Jan 01 2026 | PRJNA1210577 | rat dorsal root ganglia |
| GSE185310 | Homo sapiens | Oct 12 2021 | PRJNA768489 | |
| GSE188644 | Homo sapiens | Mar 13 2023 | PRJNA779741 | |
| GSE319827 | riticum turgidum subsp. dicoccu | Feb 24 2026 | PRJNA1425155 | Leaf |
| GSE311844 | Homo sapiens | Jan 28 2026 | | |
| GSE288413 | Gallus gallus | Jan 31 2026 | PRJNA1217401 | |
| GSE312513 | Mus musculus | Feb 13 2026 | PRJNA1374083 | microglia |
| GSE317982 | Homo sapiens | Feb 12 2026 | PRJNA1416262 | |
| GSE281344 | Mus musculus | Jan 01 2026 | | left ventricular |
| GSE309478 | Mus musculus | Jan 21 2026 | PRJNA1336144 | hind limb muscles |
| GSE301651 | Homo sapiens | Feb 10 2026 | PRJNA1285922 | bladder cancer |
| GSE306094 | Homo sapiens | Feb 24 2026 | PRJNA1309114 | |
| GSE144846 | Mus musculus | Mar 10 2021 | PRJNA605144 | |
| GSE310825 | Mus musculus | Feb 25 2026 | PRJNA1367110 | embryonic limb tissue |
| GSE316072 | Mus musculus | Jan 09 2026 | PRJNA1400569 | Aorta |
| GSE289472 | Mus musculus | Feb 25 2026 | PRJNA1223185 | Mouse Testes |
| GSE313433 | Mus musculus | Jan 15 2026 | PRJNA1379491 | Bone marrow |
| GSE179603 | Homo sapiens | May 25 2022 | PRJNA744251 | PVR |
| GSE301710 | Mycobacterium avium | Jan 01 2026 | PRJNA1287462 | |
| GSE315788 | | Jan 07 2026 | | |
| GSE292625 | Homo sapiens | Jan 28 2026 | PRJNA1240323 | |
| GSE316651 | Mus musculus | Jan 15 2026 | PRJNA1403540 | PBMC |
| GSE300771 | Homo sapiens | Jan 04 2026 | PRJNA1282074 | Brain |
| GSE312452 | Mus musculus | Jan 14 2026 | PRJNA1373341 | |
| GSE307825 | Mus musculus | Feb 14 2026 | PRJNA1327377 | |

| | | | | |
|---|---|---|---|---|
| GSE270149 | Homo sapiens | Jun 26 2026 | PRJNA1125342 | Pigmented epithelium; Retina |
| GSE138533 | Homo sapiens | Jan 20 2020 | PRJNA576260 | |
| GSE310797 | Homo sapiens | Feb 26 2026 | | OSCC tumor |
| GSE304545 | Homo sapiens | Aug 12 2025 | PRJNA1301643 | |
| GSE306680 | Mus musculus | Jan 13 2026 | PRJNA1311616 | spleen |
| GSE276929 | Mus musculus | Feb 18 2026 | PRJNA1159754 | |
| GSE302483 | Mus musculus | Feb 06 2026 | PRJNA1291065 | Epidermis |
| GSE263041 | Mus musculus | Jan 06 2026 | PRJNA1095510 | lung |
| GSE202734 | Mus musculus | Feb 15 2023 | PRJNA837191 | retina |
| GSE316666 | Mus musculus | Feb 25 2026 | PRJNA1403556 | Aorta |
| GSE196882 | Zea mays | Jan 06 2026 | | |
| GSE319795 | icus;Bos indicus x Bos taurus;Bo: | Feb 17 2026 | | whole blood |
| GSE292170 | Mus musculus | Jan 01 2026 | PRJNA1237325 | TMJ condyles |
| GSE247318 | Mus musculus | Dec 22 2023 | PRJNA1037100 | |
| GSE288718 | Acinetobacter baumannii | Feb 10 2026 | PRJNA1219132 | |
| GSE310953 | Mus musculus | Jan 28 2026 | PRJNA1367595 | cerebral cortex |
| GSE224878 | Homo sapiens | Feb 26 2026 | PRJNA932841 | |
| GSE310683 | Homo sapiens | Jan 27 2026 | PRJNA1366936 | cells from a healthy donor |
| GSE145872 | Homo sapiens | Apr 06 2020 | PRJNA608562 | |
| GSE301604 | Mus musculus | Jan 19 2026 | PRJNA1285701 | intestinal tissue |
| GSE313038 | Canis lupus familiaris | Apr 01 2026 | PRJNA1378173 | skin |
| GSE290689 | occidioides posadasii;unidentifie | Feb 26 2026 | PRJNA1229574 | |
| GSE295696 | Mus musculus | Feb 12 2026 | PRJNA1255615 | heart |
| GSE132945 | Mus musculus | Feb 28 2020 | PRJNA549507 | |
| GSE270217 | Homo sapiens | Jan 26 2026 | PRJNA1125425 | |
| GSE293278 | Homo sapiens | Jan 13 2026 | PRJNA1243474 | |
| GSE282913 | Homo sapiens | Feb 28 2026 | PRJNA1190997 | |
| GSE308164 | Homo sapiens | Jan 19 2026 | PRJNA1329125 | Breast |
| GSE280658 | Mus musculus | Feb 19 2026 | PRJNA1179904 | Thymus |
| GSE147573 | Mus musculus | Apr 22 2020 | PRJNA615623 | Retina |
| GSE288313 | Danio rerio | May 14 2025 | PRJNA1216901 | eye |
| GSE228865 | Homo sapiens | Feb 26 2026 | PRJNA951753 | |
| GSE222989 | Homo sapiens | Jan 01 2026 | PRJNA924344 | exocrine pancreas |
| GSE300652 | Homo sapiens | Feb 11 2026 | PRJNA1281607 | cell line |
| GSE309894 | Homo sapiens | Jan 08 2026 | PRJNA1363954 | Lymph node |
| GSE182580 | Mus musculus | Mar 01 2022 | PRJNA756965 | RPE |
| GSE185345 | Mus musculus | Jul 01 2022 | PRJNA768801 | |
| GSE123064 | Mus musculus | Jan 30 2026 | PRJNA507432 | |
| GSE313961 | Homo sapiens | Feb 05 2026 | PRJNA1387435 | |
| GSE296298 | Homo sapiens | Jun 18 2026 | PRJNA1258875 | |
| GSE312091 | Mus musculus | Feb 14 2026 | PRJNA1372156 | |
| GSE201219 | Homo sapiens | Jun 01 2022 | PRJNA830445 | enerated human retinal organoids |
| GSE286003 | Mus musculus | Feb 08 2026 | PRJNA1207460 | Spinal cord |
| GSE268981 | Homo sapiens | Jun 08 2024 | PRJNA1119548 | brain organoid |
| GSE144300 | Homo sapiens | Mar 31 2023 | PRJNA603335 | retina |
| GSE287228 | Homo sapiens | Mar 03 2025 | | |
| GSE288837 | Mus musculus | Feb 26 2026 | | Testes |
| GSE315663 | Homo sapiens | Feb 09 2026 | PRJNA1398643 | |
| GSE221898 | Homo sapiens | Jan 01 2026 | PRJNA916708 | |
| GSE281802 | Mus musculus | Feb 23 2026 | PRJNA1185734 | Skin |
| GSE286249 | Mus musculus | Jan 13 2025 | PRJNA1208486 | |
| GSE303021 | Mus musculus | Feb 27 2026 | PRJNA1293366 | lung |
| GSE277894 | Homo sapiens | Jan 28 2026 | PRJNA1164720 | glioma tissue |
| GSE285390 | Homo sapiens | Feb 11 2026 | PRJNA1203267 | cell line |
| GSE316386 | Histoplasma ohiense | Jan 26 2026 | PRJNA1402408 | |
| GSE236316 | Homo sapiens | Jan 19 2026 | PRJNA989665 | |
| GSE314043 | Mus musculus | Dec 16 2025 | | retina |
| GSE310240 | Mus musculus | Feb 18 2026 | PRJNA1365514 | whole liver |
| GSE317485 | Homo sapiens | Jan 28 2026 | PRJNA1413412 | peripheral blood (NSG) |
| GSE271682 | Homo sapiens | Jan 07 2026 | PRJNA1132921 | |
| GSE312185 | Homo sapiens | Jan 12 2026 | | plasma |
| GSE294049 | Mus musculus | Apr 14 2025 | PRJNA1247953 | Mouse retina, C57BL/6 |
| GSE292490 | Mus musculus | Jan 01 2026 | PRJNA1238854 | Murine pancreatic cancer |

| | | | | |
|---|---|---|---|---|
| GSE279825 | Mus musculus | Dec 03 2024 | PRJNA1174746 | Retina |
| GSE319275 | Mus musculus | Feb 28 2026 | PRJNA1422726 | brain |
| GSE306402 | Danio rerio | Feb 18 2026 | PRJNA1310580 | Larval fish |
| GSE291056 | Drosophila melanogaster | Feb 18 2026 | PRJNA1231470 | adult ovary |
| GSE289942 | Arabidopsis thaliana | Jan 29 2026 | PRJNA1225620 | Seedlings |
| GSE287141 | Mus musculus | Jan 15 2026 | PRJNA1211175 | Liver |
| GSE297836 | Homo sapiens | Jan 13 2026 | PRJNA1266256 | |
| GSE252636 | Mus musculus | Feb 07 2026 | PRJNA1061546 | Spleen |
| GSE336907 | Homo sapiens | Jul 04 2026 | PRJNA1483545 | |
| GSE315268 | Streptomyces lividans | Jan 21 2026 | | |
| GSE290000 | Mus musculus | Feb 19 2026 | PRJNA1225722 | Whole heart tissue |
| GSE264212 | Homo sapiens | Sep 04 2024 | PRJNA1101468 | |
| GSE303209 | Homo sapiens | Feb 25 2026 | PRJNA1294342 | Small intestine |
| GSE225640 | Homo sapiens | Mar 11 2024 | PRJNA937023 | etinal pigment epithelium |
| GSE311435 | Homo sapiens | Jan 01 2026 | PRJNA1369457 | |
| GSE302984 | Homo sapiens | Jan 13 2026 | PRJNA1293305 | |
| GSE169039 | Mus musculus | Aug 22 2022 | PRJNA714949 | |
| GSE315636 | Mus musculus | Jan 15 2026 | PRJNA1398362 | Ovary |
| GSE124323 | Mus musculus | Jan 01 2020 | PRJNA511607 | |
| GSE303043 | Homo sapiens;Mus musculus | Feb 26 2026 | PRJNA1293407 | small cell lung cancer |
| GSE318128 | Homo sapiens | Feb 28 2026 | PRJNA1416773 | |
| GSE303721 | Homo sapiens | Feb 06 2026 | PRJNA1297156 | |
| GSE299621 | Saccharomyces cerevisiae | Jan 28 2026 | PRJNA1275684 | |
| GSE310301 | Mus musculus | Jan 14 2026 | PRJNA1365725 | 4T1 breast tumor |
| GSE304875 | Mus musculus | Jan 13 2026 | PRJNA1303126 | |
| GSE298122 | Mus musculus | Jul 18 2025 | PRJNA1268389 | Retina |
| GSE128143 | Homo sapiens | Apr 22 2020 | PRJNA526534 | nal Pigmented Epithelial Cells |
| GSE299060 | Homo sapiens | Jan 21 2026 | PRJNA1272490 | |
| GSE289599 | Mus musculus | Feb 08 2026 | | bone marrow |
| GSE315040 | Mus musculus | Jan 05 2026 | | spleen |
| GSE287885 | Mus musculus | Jan 01 2026 | PRJNA1214858 | Adipose |
| GSE313186 | Homo sapiens | Jan 27 2026 | PRJNA1378772 | Ethmoid sinus tissue |
| GSE209624 | Homo sapiens | Feb 15 2026 | PRJNA861848 | |
| GSE318276 | Mus musculus | Feb 05 2026 | PRJNA1418516 | |
| GSE294749 | Mus musculus | Jan 09 2026 | PRJNA1251547 | Pancreas Tumor |
| GSE296802 | Homo sapiens | Feb 19 2026 | PRJNA1261066 | |
| GSE287369 | Homo sapiens | Jan 31 2026 | PRJNA1212504 | |
| GSE149715 | Mus musculus | Jun 22 2020 | PRJNA629793 | Retina |
| GSE286175 | Mus musculus | Feb 18 2026 | PRJNA1208063 | cortex |
| GSE291249 | Homo sapiens | Feb 16 2026 | PRJNA1232661 | heart right atrium |
| GSE277711 | Homo sapiens | Jan 14 2026 | PRJNA1163422 | |
| GSE290062 | Homo sapiens | Feb 28 2026 | PRJNA1226129 | plasma exosomes |
| GSE294261 | Mus musculus | Feb 18 2026 | PRJNA1248894 | dorsal hippocampus |
| GSE311009 | Homo sapiens | Jan 26 2026 | PRJNA1368479 | |
| GSE314987 | Homo sapiens | Jan 06 2026 | PRJNA1394200 | colon |
| GSE311274 | Homo sapiens | Feb 18 2026 | PRJNA1368880 | colon |
| GSE149040 | Mus musculus | Nov 24 2020 | PRJNA627199 | Neural retina |
| GSE305091 | Mus musculus | Jan 05 2026 | | Pancreatic Islet |
| GSE289920 | Homo sapiens | Feb 06 2026 | PRJNA1225295 | |
| GSE304792 | Bos taurus;Homo sapiens | Feb 12 2026 | | GV and MII bovine oocytes |
| GSE315077 | Homo sapiens | Jan 08 2026 | PRJNA1394965 | |
| GSE200339 | Homo sapiens | Aug 26 2022 | PRJNA824114 | noid derived from human iPSCs |
| GSE200915 | Mus musculus | May 21 2022 | PRJNA827181 | |
| GSE312010 | Mus musculus | Jun 03 2026 | PRJNA1372054 | Retina |
| GSE311858 | Homo sapiens | Jan 01 2026 | PRJNA1371772 | |
| GSE289613 | Homo sapiens | Feb 13 2026 | PRJNA1223819 | |
| GSE316213 | Mus musculus | Jan 15 2026 | PRJNA1401817 | murine pancreatic cancer |
| GSE243713 | Mus musculus | Jan 12 2026 | PRJNA1019679 | |
| GSE197507 | Mus musculus | Nov 15 2022 | PRJNA810718 | |
| GSE297767 | Homo sapiens | Jan 07 2026 | PRJNA1266143 | skin |
| GSE147012 | Homo sapiens | Jul 15 2022 | PRJNA612796 | Retina |
| GSE306919 | Mus musculus | Feb 12 2026 | PRJNA1313918 | ovary |
| GSE226507 | Homo sapiens | Jan 14 2026 | PRJNA940453 | |

| | | | | |
|---|---|---|---|---|
| GSE298694 | Homo sapiens | Feb 15 2026 | PRJNA1270959 | |
| GSE315826 | Mus musculus | Feb 08 2026 | | Spleen |
| GSE280339 | Mus musculus | Jan 09 2026 | PRJNA1177879 | liver |
| GSE239347 | Mus musculus | Nov 16 2023 | PRJNA998873 | |
| GSE316546 | Human immunodeficiency virus | Jan 22 2026 | PRJNA1403292 | |
| GSE225867 | Mus musculus;synthetic construc | Mar 05 2023 | PRJNA937735 | whole retina |
| GSE147258 | Gallus gallus | Aug 01 2020 | PRJNA613618 | retina |
| GSE134311 | apiens;Mus musculus;synthetic c | Aug 24 2020 | PRJNA554684 | orsal root ganglion neuron |
| GSE316224 | Mus musculus | Feb 16 2026 | PRJNA1401830 | Control Mouse skin |
| GSE325523 | ;Macaca fascicularis;Mus muscu | Mar 25 2026 | | Heart - Atrium,Heart - Atrium |
| GSE301407 | Mus musculus | Jul 08 2025 | | retina |
| GSE294396 | Mus musculus | Feb 05 2026 | PRJNA1249381 | thyroid |
| GSE288127 | Arabidopsis | Jan 27 2026 | PRJNA1216242 | Callus |
| GSE314870 | Mus musculus | Jan 23 2026 | PRJNA1393043 | lung |
| GSE281031 | Mus musculus | Jun 23 2025 | PRJNA1181864 | Retina |
| GSE315932 | Histoplasma ohiense | Jan 26 2026 | PRJNA1399967 | |
| GSE181251 | Mus musculus | Aug 03 2021 | PRJNA751223 | |
| GSE158771 | Mus musculus | Jan 13 2022 | PRJNA666453 | liver |
| GSE302680 | Mus musculus | Feb 18 2026 | PRJNA1291718 | Mediastinal lymph nodes |
| GSE252680 | Drosophila melanogaster | Jan 30 2026 | PRJNA1061799 | head |
| GSE288279 | Mus musculus | Jan 21 2026 | PRJNA1216666 | m flash-frozen powderized whole pancreas |
| GSE318979 | Mus musculus | Feb 28 2026 | PRJNA1420916 | tumor stage |
| GSE299083 | Mus musculus | Jan 01 2026 | PRJNA1272543 | colon |
| GSE286196 | Mus musculus | Jan 08 2026 | PRJNA1208098 | Brain |
| GSE312895 | Mus musculus | Jan 30 2026 | PRJNA1377564 | |
| GSE280050 | Homo sapiens | Jan 29 2026 | PRJNA1176030 | |
| GSE300738 | Mus musculus | Feb 25 2026 | PRJNA1282003 | Stomach corpus |
| GSE298973 | Homo sapiens | Jun 04 2026 | PRJNA1272042 | |
| GSE192882 | Homo sapiens | Oct 28 2022 | PRJNA793671 | |
| GSE325128 | Mus musculus | Apr 09 2026 | PRJNA1438672 | Retina |
| GSE309547 | Mus musculus | Feb 26 2026 | PRJNA1336237 | embryo |
| GSE301513 | Homo sapiens | Jan 29 2026 | | bone |
| GSE317433 | Homo sapiens | Jan 28 2026 | PRJNA1412668 | bone |
| GSE288033 | Homo sapiens | Jan 23 2026 | PRJNA1215751 | heral blood mononuclear cells |
| GSE263517 | Mus musculus | Feb 17 2026 | PRJNA1097856 | cortex |
| GSE295581 | Caulobacter vibrioides | Feb 25 2026 | PRJNA1255209 | |
| GSE285078 | Mus musculus | Jan 20 2026 | PRJNA1201149 | |
| GSE281519 | Homo sapiens | Jan 05 2026 | PRJNA1184175 | L.Ovary |
| GSE315094 | Mus musculus | Jan 06 2026 | PRJNA1394984 | CD1 mouse fetal kidney |
| GSE309225 | Homo sapiens | Feb 23 2026 | PRJNA1334373 | |
| GSE327901 | Branchiostoma floridae | May 07 2026 | PRJNA1452963 | brain |
| GSE289113 | Homo sapiens | Feb 07 2026 | PRJNA1221568 | |
| GSE234646 | Danio rerio | Jun 13 2023 | PRJNA982182 | dissected eye |
| GSE287910 | Homo sapiens | Feb 23 2026 | PRJNA1215267 | Spleen |
| GSE318481 | Mus musculus | Feb 12 2026 | PRJNA1418823 | pancreas |
| GSE313771 | Homo sapiens | Jan 31 2026 | PRJNA1381577 | |
| GSE297705 | Methylorubrum extorquens | Jan 21 2026 | PRJNA1265739 | |
| GSE196779 | Mus musculus | Feb 11 2026 | PRJNA807299 | |
| GSE312072 | Mus musculus | Jan 13 2026 | PRJNA1372149 | |
| GSE263149 | Mus musculus | Feb 11 2026 | PRJNA1095981 | bone marrow |
| GSE267662 | Homo sapiens | May 19 2024 | PRJNA1112368 | umbilical cord |
| GSE185671 | Mus musculus | Feb 01 2022 | PRJNA770676 | |
| GSE304165 | Mus musculus | Jan 28 2026 | PRJNA1299300 | femoral bone |
| GSE244812 | Mus musculus | Aug 13 2024 | PRJNA1024933 | Eye |
| GSE314812 | Homo sapiens | Feb 23 2026 | PRJNA1392638 | GC Primary tumor |
| GSE292232 | Mus musculus | Feb 24 2026 | PRJNA1237695 | gastrocnemius |
| GSE281612 | Homo sapiens;Mus musculus | Feb 09 2026 | PRJNA1184798 | Bone |
| GSE312727 | Homo sapiens | Jan 31 2026 | PRJNA1375778 | |
| GSE280673 | Mus musculus | Feb 25 2026 | | |
| GSE294189 | Homo sapiens | Jan 01 2026 | PRJNA1248524 | |
| GSE301618 | Homo sapiens | Jan 11 2026 | PRJNA1285800 | |
| GSE304318 | Mus musculus | Jun 03 2026 | PRJNA1300372 | retina |
| GSE305104 | Mus musculus | Jan 23 2026 | PRJNA1304433 | corneal endothelium |

| | | | | |
|---|---|---|---|---|
| GSE303683 | Homo sapiens | Feb 25 2026 | PRJNA1297084 | GBM |
| GSE271661 | Mus musculus | Jan 29 2025 | PRJNA1132714 | retina |
| GSE202728 | Mus musculus | Feb 15 2023 | PRJNA837179 | retina |
| GSE299783 | Mus musculus | Jan 28 2026 | PRJNA1276509 | all intestinal lamina propria |
| GSE293412 | Homo sapiens;Mus musculus | Jan 12 2026 | PRJNA1244375 | aorta |
| GSE312479 | Mus musculus | Jan 26 2026 | PRJNA1374047 | l, spleen, cLN, mLN and sLN |
| GSE307480 | Mus musculus | Jan 04 2026 | PRJNA1321816 | osreosarcoma cells |
| GSE318595 | Homo sapiens | Mar 23 2026 | PRJNA1419282 | |
| GSE154544 | Homo sapiens | Oct 31 2020 | PRJNA646635 | |
| GSE150258 | Mus musculus | Dec 17 2022 | PRJNA631682 | retina |
| GSE305027 | Homo sapiens | Jan 30 2026 | PRJNA1303732 | Epithelial cell line |
| GSE289030 | Mus musculus | Feb 06 2026 | PRJNA1220669 | |
| GSE284564 | rymna berryi;Octopus bimaculo | Jan 01 2026 | PRJNA1199547 | Arm |
| GSE312633 | Homo sapiens | Jan 10 2026 | PRJNA1375552 | Blood |
| GSE163568 | Homo sapiens;Mus musculus | Jan 11 2022 | PRJNA686783 | Human Umbilical Vein Endothelial Cells) |
| GSE277963 | Mus musculus | Jan 14 2026 | PRJNA1164831 | spleen |
| GSE197446 | Homo sapiens | Feb 26 2022 | PRJNA810362 | iPSC-derived retinal tissue |
| GSE314097 | Mus musculus | Jan 19 2026 | PRJNA1390095 | |
| GSE289752 | Homo sapiens | Feb 10 2026 | PRJNA1224446 | Peripheral blood |
| GSE292326 | Homo sapiens | Jan 12 2026 | PRJNA1237939 | meningioma |
| GSE311919 | Danio rerio | Jan 19 2026 | PRJNA1371935 | caudal fin |
| GSE311597 | Mus musculus | Jan 26 2026 | PRJNA1369976 | 12.5 fetal liver |
| GSE298926 | Homo sapiens | Jan 21 2026 | PRJNA1271941 | |
| GSE310294 | Rattus norvegicus | Jan 07 2026 | PRJNA1365715 | bone |
| GSE285787 | Arabidopsis thaliana | Jan 06 2026 | PRJNA1206087 | seedlings |
| GSE223828 | Mus musculus | Jan 25 2026 | PRJNA928478 | liver |
| GSE87200 | Homo sapiens | Sep 02 2020 | PRJNA343848 | |
| GSE233440 | Mus musculus | May 31 2023 | PRJNA976271 | retina |
| GSE315009 | Caenorhabditis elegans | Jan 05 2026 | PRJNA1394779 | whole animal |
| GSE262965 | Mus musculus | Jan 06 2026 | PRJNA1095007 | lung |
| GSE319741 | Homo sapiens | Feb 17 2026 | PRJNA1425037 | colon |
| GSE302335 | Homo sapiens | Jan 16 2026 | PRJNA1290110 | |
| GSE216267 | Homo sapiens | Jan 28 2026 | PRJNA892898 | Tonsil |
| GSE310774 | Homo sapiens | Feb 20 2026 | PRJNA1367060 | cell line |
| GSE269304 | Mus musculus | Jan 06 2026 | PRJNA1120930 | |
| GSE314577 | Homo sapiens | Feb 18 2026 | PRJNA1391543 | plasma |
| GSE309954 | Mus musculus | Feb 12 2026 | PRJNA1364038 | Liver |
| GSE301937 | Mus musculus | Feb 24 2026 | PRJNA1287764 | |
| GSE270482 | Mus musculus | Jan 28 2025 | PRJNA1126727 | p6 neonatal murine brain |
| GSE317499 | Mus musculus | Jan 28 2026 | PRJNA1414267 | Brain tissue - CA2 region |
| GSE307160 | Euprymna scolopes | Feb 25 2026 | PRJNA1314669 | light organ |
| GSE310743 | Homo sapiens | Feb 24 2026 | PRJNA1367017 | Skeletal Muscle |
| GSE288586 | Mus musculus | Jan 14 2026 | PRJNA1218629 | |
| GSE227861 | Mus musculus | Feb 25 2024 | PRJNA947200 | Retina |
| GSE307602 | Mus musculus | Jan 21 2026 | PRJNA1322367 | Pancreas |
| GSE139099 | Homo sapiens | Jul 23 2020 | PRJNA578382 | |
| GSE282175 | Mus musculus | Jul 28 2025 | PRJNA1187531 | Retina |
| GSE256101 | Homo sapiens;Mus musculus | Feb 18 2026 | PRJNA1078236 | |
| GSE316876 | Mus musculus | Jan 24 2026 | PRJNA1405470 | colon |
| GSE315521 | Homo sapiens | Jan 08 2026 | PRJNA1398207 | cell culture |
| GSE309790 | Homo sapiens | Feb 16 2026 | PRJNA1363762 | cord clood |
| GSE319137 | nio albolineatus;Danio rerio x D | Feb 11 2026 | PRJNA1422161 | Whole skin |
| GSE285860 | Mus musculus | Jan 01 2026 | PRJNA1206297 | colon |
| GSE307043 | Homo sapiens | Jan 21 2026 | PRJNA1314167 | CSF |
| GSE304485 | | Feb 12 2026 | | |
| GSE248449 | Mus musculus | Jan 26 2026 | PRJNA1044036 | brain |
| GSE318449 | Mus musculus | Feb 07 2026 | PRJNA1418746 | yonic day 14.5 Mullerian duct |
| GSE279594 | Homo sapiens;Mus musculus | Feb 27 2026 | PRJNA1173554 | bone marrow |
| GSE129443 | Mus musculus | Jan 24 2020 | PRJNA531379 | retina |
| GSE299422 | Mus musculus | Feb 06 2026 | PRJNA1274402 | |
| GSE315435 | Homo sapiens;Mus musculus | Jan 06 2026 | PRJNA1397166 | CD1 mouse fetal kidney |
| GSE252709 | Homo sapiens | Jan 01 2026 | PRJNA1062303 | glioblastoma |
| GSE316962 | Mus musculus | Jan 21 2026 | PRJNA1405600 | heart, testis |

| | | | | |
|---|---|---|---|---|
| GSE298609 | Mus musculus | Jan 22 2026 | PRJNA1270033 | incisor mesenchyme |
| GSE293784 | Drosophila melanogaster | Jan 23 2026 | PRJNA1246211 | Whole brain |
| GSE302216 | Homo sapiens | Jan 24 2026 | PRJNA1289442 | human 4C embryo |
| GSE297587 | Homo sapiens | Jan 22 2026 | PRJNA1265224 | Brain |
| GSE227975 | Homo sapiens | May 01 2023 | PRJNA947606 | derived retinal organoid day 40 |
| GSE310657 | Escherichia coli K-12 | Feb 08 2026 | PRJNA1366827 | |
| GSE306777 | Mus musculus | Jan 13 2026 | PRJNA1312087 | sdLNs and thymus |
| GSE313291 | Homo sapiens | Feb 26 2026 | PRJNA1378879 | Peripheral blood |
| GSE311154 | Mus musculus | Feb 23 2026 | PRJNA1368752 | imary visual cortex nucleus |
| GSE140128 | Mus musculus | Sep 30 2020 | PRJNA588391 | |
| GSE303629 | Ictidomys tridecemlineatus | Jul 28 2025 | PRJNA1295946 | E18 retina |
| GSE286441 | Homo sapiens | Jan 25 2025 | PRJNA1209360 | |
| GSE179871 | Homo sapiens | Feb 09 2026 | PRJNA745443 | |
| GSE315216 | Homo sapiens | Jan 07 2026 | PRJNA1395137 | |
| GSE277570 | Mus musculus | Sep 18 2025 | PRJNA1162833 | Retina |
| GSE184286 | Mus musculus | Oct 19 2021 | PRJNA763893 | n vitro primary Muller glia |
| GSE306154 | Homo sapiens;Mus musculus | Feb 26 2026 | PRJNA1309594 | colon lamina propria |
| GSE316823 | Homo sapiens | Jan 21 2026 | PRJNA1404292 | Pancreas |
| GSE301791 | Rattus norvegicus | Jan 06 2026 | PRJNA1287571 | Dissociated brain |
| GSE241268 | Mus musculus | Oct 01 2023 | PRJNA1007621 | mouse retina |
| GSE303454 | Mus musculus | Jan 22 2026 | PRJNA1295464 | liver |
| GSE181299 | Drosophila melanogaster | Jan 17 2023 | PRJNA751507 | Pupal retina |
| GSE292860 | Homo sapiens | Feb 12 2026 | PRJNA1241777 | |
| GSE151948 | Gallus gallus | Apr 19 2021 | PRJNA637867 | |
| GSE303463 | Homo sapiens | Jan 14 2026 | PRJNA1295470 | Placenta |
| GSE245543 | Mus musculus | Jun 06 2024 | PRJNA1028974 | retina |
| GSE289585 | Mus musculus | Jan 14 2026 | PRJNA1223471 | brain |
| GSE316185 | Histoplasma ohiense | Jan 26 2026 | PRJNA1403597 | |
| GSE204880 | Homo sapiens;Mus musculus | Aug 15 2023 | PRJNA842665 | kidney |
| GSE312699 | Homo sapiens | Jan 16 2026 | PRJNA1375739 | primary chordoma |
| GSE237309 | Mus musculus | Jul 13 2023 | PRJNA994552 | etinal pigment epithelium |
| GSE186309 | Homo sapiens | Jan 29 2026 | PRJNA773376 | |
| GSE286169 | Mus musculus | Jan 01 2026 | PRJNA1208044 | Colon |
| GSE307102 | Homo sapiens | Jan 28 2026 | PRJNA1314264 | whole blood |
| GSE311022 | Mus musculus | Feb 07 2026 | PRJNA1368475 | Spleen |
| GSE309901 | Trypanosoma cruzi | Feb 13 2026 | PRJNA1363973 | unicellular organism |
| GSE287534 | Mus musculus | Jan 01 2026 | PRJNA1213687 | |
| GSE262182 | Mus musculus | Jan 27 2026 | PRJNA1090523 | Bone marrow |
| GSE315460 | Mus musculus | Feb 19 2026 | PRJNA1397185 | istal part of small intestine |
| GSE251685 | Mus musculus | Jan 01 2024 | PRJNA1055063 | Retina |
| GSE310160 | Homo sapiens | Jan 20 2026 | PRJNA1365218 | lung cancer |
| GSE294114 | Mus musculus | Feb 06 2026 | PRJNA1248409 | Aorta |
| GSE273378 | Homo sapiens | Jan 26 2026 | PRJNA1141507 | lung adenocarcinoma |
| GSE315302 | Homo sapiens | Feb 18 2026 | PRJNA1395604 | |
| GSE260525 | Mus musculus | Feb 18 2026 | PRJNA1081945 | |
| GSE261739 | Homo sapiens | Feb 12 2026 | PRJNA1088525 | Whole Blood |
| GSE319155 | Homo sapiens | Feb 11 2026 | PRJNA1422218 | Lymph node |
| GSE252521 | Homo sapiens | Jan 14 2026 | PRJNA1061090 | Prostate |
| GSE190615 | Homo sapiens | Dec 31 2022 | PRJNA787705 | |
| GSE252516 | Mus musculus | Jan 26 2024 | PRJNA1061076 | |
| GSE297798 | Mus musculus | Feb 05 2026 | PRJNA1266198 | Pancreatic cancer |
| GSE287261 | Homo sapiens | Jan 17 2026 | PRJNA1211999 | |
| GSE226785 | Homo sapiens | Jan 01 2026 | PRJNA942050 | |
| GSE200206 | Homo sapiens | Feb 11 2023 | PRJNA823674 | |
| GSE272153 | Mus musculus | Jan 28 2026 | PRJNA1135219 | liver |
| GSE275289 | Mus musculus | Jan 01 2026 | PRJNA1150193 | |
| GSE315457 | Mus musculus | Feb 19 2026 | PRJNA1397174 | istal part of small intestine |
| GSE299477 | Homo sapiens | Jan 29 2026 | PRJNA1274514 | sigmoid colon organoids |
| GSE300820 | Capra hircus | Feb 24 2026 | PRJNA1282149 | hair follicle stem cells from goat skin |
| GSE315981 | Homo sapiens | Jan 12 2026 | PRJNA1400059 | |
| GSE282444 | Homo sapiens | Feb 28 2026 | PRJNA1188591 | Colon |
| GSE290278 | Arabidopsis thaliana | Jan 29 2026 | PRJNA1227116 | Seedlings |
| GSE184080 | Homo sapiens | Sep 17 2021 | PRJNA763113 | |

| | | | | |
|---|---|---|---|---|
| GSE125564 | Homo sapiens | Feb 26 2020 | PRJNA516819 | |
| GSE314071 | Homo sapiens | Jan 01 2026 | PRJNA1389916 | PBMC |
| GSE286125 | Mus musculus | Jan 01 2026 | PRJNA1207974 | |
| GSE266902 | Chlamydomonas reinhardtii | Jan 26 2026 | PRJNA1108696 | cell |
| GSE299752 | Mus musculus | Feb 11 2026 | PRJNA1276435 | Tibialis anterior muscle |
| GSE315772 | Mus musculus | Jan 11 2026 | PRJNA1398935 | whole Liver |
| GSE310245 | Homo sapiens | Jun 10 2026 | PRJNA1365526 | Retina |
| GSE268136 | Homo sapiens | May 31 2024 | PRJNA1114744 | |
| GSE315745 | Mus musculus | Jan 06 2026 | PRJNA1398883 | lung |
| GSE253665 | 3ombyx mandarina;Bombyx mor | Jan 14 2026 | PRJNA1066739 | wing disc |
| GSE302479 | Mus musculus | Feb 15 2026 | PRJNA1291064 | spinal cord |
| GSE311546 | Mus musculus | Jan 10 2026 | PRJNA1369841 | |
| GSE290032 | Mus musculus | Jan 07 2026 | PRJNA1225843 | bone marrow |
| GSE319512 | Mus musculus | Feb 18 2026 | PRJNA1423651 | Colon |
| GSE256407 | Mus musculus | Apr 22 2025 | PRJNA1079361 | retina |
| GSE299007 | Homo sapiens | Feb 20 2026 | PRJNA1272097 | tumor of oral mucosa |
| GSE306118 | Sinorhizobium meliloti | Feb 07 2026 | PRJNA1309463 | |
| GSE189281 | Homo sapiens | Dec 16 2021 | PRJNA782342 | Pancreatic cancer |
| GSE245538 | Mus musculus | Jun 06 2024 | PRJNA1028959 | retina |
| GSE315538 | Mus musculus | Feb 28 2026 | PRJNA1398225 | dental epithelial cells |
| GSE302604 | Mus musculus | Jan 16 2026 | PRJNA1291490 | Liver |
| GSE319270 | Homo sapiens | Feb 16 2026 | PRJNA1422732 | Blood |
| GSE318332 | Homo sapiens | Feb 13 2026 | PRJNA1418590 | PBMC |
| GSE290111 | Homo sapiens | Feb 25 2026 | PRJNA1226218 | Serum |
| GSE319431 | Homo sapiens | Feb 17 2026 | PRJNA1423271 | |
| GSE302045 | Homo sapiens | Feb 11 2026 | PRJNA1288409 | PC metastasis |
| GSE298145 | Mus musculus | Jan 28 2026 | PRJNA1268420 | Tumor |
| GSE307545 | Mus musculus | Feb 21 2026 | PRJNA1322185 | Spleen |
| GSE273303 | Oryza sativa | Jan 19 2026 | PRJNA1141399 | stigma |
| GSE313953 | Mus musculus | Feb 26 2026 | PRJNA1387421 | spleen |
| GSE276603 | Homo sapiens | Nov 04 2025 | PRJNA1157900 | |
| GSE311618 | Homo sapiens | Jan 19 2026 | PRJNA1370009 | |
| GSE281196 | Mus musculus | Feb 17 2026 | PRJNA1182721 | liver |
| GSE302727 | Plasmodium falciparum | Jan 04 2026 | PRJNA1291806 | |
| GSE311307 | Mus musculus | Feb 18 2026 | PRJNA1368914 | kidney |
| GSE307227 | Danio rerio | Feb 26 2026 | PRJNA1314796 | eye |
| GSE149104 | Homo sapiens | Dec 02 2020 | PRJNA627391 | |
| GSE282078 | Mus musculus | Feb 06 2026 | PRJNA1186955 | |
| GSE298610 | Mus musculus | Jan 22 2026 | PRJNA1270034 | incisor mesenchyme |
| GSE291147 | Homo sapiens | Feb 06 2026 | | |
| GSE313964 | Mus musculus | Jan 06 2026 | PRJNA1387445 | Testis |
| GSE310433 | Homo sapiens | Jan 16 2026 | PRJNA1372807 | human pancreatic cancer |
| GSE306513 | Homo sapiens | Jan 12 2026 | PRJNA1311015 | |
| GSE293282 | Mus musculus | Jan 26 2026 | PRJNA1243699 | brain |
| GSE319888 | Homo sapiens | Feb 23 2026 | PRJNA1425482 | brain |
| GSE316297 | Mus musculus | Jan 21 2026 | PRJNA1402246 | |
| GSE318305 | Rattus norvegicus | Feb 06 2026 | PRJNA1418579 | heart |
| GSE289388 | Homo sapiens | Feb 11 2026 | PRJNA1222834 | peripheral blood |
| GSE206087 | Mus musculus | Jan 11 2023 | PRJNA849166 | lung |
| GSE298033 | Homo sapiens | Feb 25 2026 | PRJNA1267134 | bone marrow |
| GSE296830 | Homo sapiens | Jan 07 2026 | PRJNA1261788 | blood |
| GSE272214 | Homo sapiens | Jan 01 2026 | PRJNA1135702 | |
| GSE208677 | Homo sapiens | Jul 21 2022 | PRJNA860599 | Tumor |
| GSE298751 | Mus musculus | Feb 10 2026 | PRJNA1271055 | heart |
| GSE269102 | Mus musculus | Jul 04 2024 | PRJNA1120107 | Neural retina |
| GSE267936 | Mus musculus | Jan 23 2025 | PRJNA1113670 | retina |
| GSE256465 | Arabidopsis thaliana | Feb 22 2026 | PRJNA1079740 | 10-d-old seedlings |
| GSE317380 | Danio rerio | Apr 17 2026 | PRJNA1411997 | testes |
| GSE313812 | Solanum tuberosum | Jan 06 2026 | PRJNA1381632 | stolon |
| GSE268636 | Homo sapiens | Jan 07 2026 | PRJNA1117945 | prostate |
| GSE307404 | Homo sapiens | Feb 16 2026 | PRJNA1320703 | rsolateral prefrontal cortex |
| GSE306618 | Mus musculus | Jan 13 2026 | PRJNA1311488 | lung |
| GSE104884 | Mus musculus | Oct 01 2023 | PRJNA414090 | Retina |

| | | | | |
|---|---|---|---|---|
| GSE175686 | Homo sapiens | May 26 2022 | PRJNA733186 | hTERT forskin fibroblasts |
| GSE186310 | Homo sapiens | Jan 29 2026 | PRJNA773375 | |
| GSE312873 | Mus musculus | Jan 15 2026 | PRJNA1377542 | Lung macrophages |
| GSE294253 | Homo sapiens | Jan 04 2026 | PRJNA1248880 | |
| GSE136880 | Mus musculus | Jun 02 2020 | PRJNA563908 | Retina |
| GSE159826 | Homo sapiens | Oct 23 2020 | PRJNA670656 | etinal Pigment Epithelium |
| GSE290611 | Mus musculus | Feb 23 2026 | PRJNA1229003 | gastroceiumus muscle |
| GSE236530 | Homo sapiens;synthetic construc | Feb 15 2026 | PRJNA991538 | |
| GSE313931 | Homo sapiens | Feb 24 2026 | PRJNA1387384 | |
| GSE305811 | Homo sapiens | Aug 25 2025 | PRJNA1307877 | endometrium |
| GSE289831 | Mus musculus | Feb 17 2026 | | spleen |
| GSE312950 | Mus musculus | Feb 07 2026 | PRJNA1377649 | |
| GSE319453 | Mus musculus | Feb 20 2026 | PRJNA1423300 | Intestine |
| GSE288850 | Mus musculus | Feb 10 2026 | PRJNA1219605 | Ear skin |
| GSE276892 | Homo sapiens | Dec 06 2024 | PRJNA1159345 | hyalocytes |
| GSE221471 | Danio rerio | Jan 03 2023 | PRJNA914428 | |
| GSE165743 | Rattus norvegicus | Jan 29 2021 | PRJNA695565 | |
| GSE315604 | Homo sapiens | Jan 05 2026 | PRJNA1398318 | choalveolar lavage fluid (BALF) |
| GSE268077 | Homo sapiens | Jan 16 2026 | PRJNA1114462 | PDA Cell Line |
| GSE184867 | Homo sapiens | Feb 12 2025 | PRJNA766685 | |
| GSE322628 | Mus musculus | Jun 24 2026 | PRJNA1430759 | Retina |
| GSE299613 | Homo sapiens | Feb 23 2026 | PRJNA1275682 | |
| GSE293582 | Homo sapiens | Mar 22 2026 | PRJNA1245288 | retinal organoid |
| GSE317978 | Mus musculus | Feb 06 2026 | PRJNA1416249 | islet |
| GSE256213 | synthetic construct | Nov 22 2024 | PRJNA1078425 | Synthetic constructs |
| GSE287278 | Homo sapiens | Sep 04 2025 | | uterine endometrium |
| GSE308381 | Mus musculus | Feb 20 2026 | PRJNA1330306 | Pancreatic beta cells |
| GSE291125 | Homo sapiens | Jan 20 2026 | PRJNA1232049 | rectum |
| GSE237204 | Homo sapiens | Oct 01 2023 | PRJNA994169 | Retina |
| GSE315151 | Rattus norvegicus | Jan 29 2026 | PRJNA1395062 | tooth |
| GSE290339 | Homo sapiens | Feb 24 2026 | PRJNA1227511 | |
| GSE317294 | Sus scrofa | Feb 09 2026 | PRJNA1406696 | |
| GSE283719 | Homo sapiens | Jan 21 2026 | PRJNA1195089 | |
| GSE289806 | Homo sapiens | Feb 17 2026 | PRJNA1224834 | iPSC-derived cells |
| GSE310451 | Homo sapiens;Mus musculus | Feb 11 2026 | PRJNA1366362 | nal motor neuron organoid |
| GSE276656 | Mus musculus | Feb 10 2026 | PRJNA1158460 | mPFC |
| GSE311352 | Homo sapiens | Jan 26 2026 | PRJNA1369202 | |
| GSE318398 | Mus musculus | Feb 08 2026 | PRJNA1418682 | bone narrow cells |
| GSE214005 | Mus musculus | Jan 20 2023 | PRJNA883604 | hal pigment epithelium (RPE) |
| GSE264095 | Oryza sativa Japonica Group | Jan 24 2026 | PRJNA1100973 | Shoot |
| GSE316568 | Homo sapiens | Jan 21 2026 | PRJNA1403408 | |
| GSE198863 | Homo sapiens | Mar 18 2023 | PRJNA817206 | kidney, Embryo |
| GSE317774 | Homo sapiens | Jan 27 2026 | PRJNA1415218 | |
| GSE280320 | Mus musculus | Feb 18 2026 | PRJNA1177836 | liver |
| GSE304354 | Homo sapiens | Feb 16 2026 | PRJNA1300448 | Cortical organoids D60 |
| GSE242274 | Homo sapiens | Jan 07 2026 | PRJNA1013015 | skin and soft tissue |
| GSE270758 | Mus musculus | Jul 02 2024 | PRJNA1128127 | retina |
| GSE279510 | Mus musculus | Jan 29 2026 | PRJNA1173166 | Epidermis of back skin |
| GSE310499 | Homo sapiens | Jan 31 2026 | PRJNA1366442 | Prostate |
| GSE319822 | Mus musculus | Feb 24 2026 | | Spinal cord |
| GSE300602 | Homo sapiens | Jan 09 2026 | PRJNA1281535 | small intestine |
| GSE289322 | Mus musculus | Feb 11 2026 | PRJNA1222414 | RV |
| GSE255302 | Arabidopsis thaliana | Feb 25 2026 | PRJNA1074416 | leaves |
| GSE147400 | Mus musculus | Mar 24 2020 | PRJNA614479 | bar 4 and 5 dorsal root ganglia |
| GSE310934 | Homo sapiens | Feb 24 2026 | PRJNA1367572 | |
| GSE298286 | Homo sapiens | Feb 17 2026 | PRJNA1268887 | ne metastatic breast tumor |
| GSE319901 | Homo sapiens | Feb 20 2026 | PRJNA1425499 | Serum |
| GSE274605 | Mus musculus | Jan 13 2026 | PRJNA1147272 | Brain |
| GSE167045 | Homo sapiens | Sep 30 2021 | PRJNA702788 | |
| GSE266039 | Mus musculus | May 31 2025 | PRJNA1105560 | P0 retina |
| GSE317102 | Mus musculus | Feb 06 2026 | PRJNA1405990 | Brain |
| GSE280049 | Homo sapiens | Aug 11 2025 | PRJNA1176036 | |
| GSE286490 | Homo sapiens | Jan 01 2026 | PRJNA1209618 | |

| | | | | |
|---|---|---|---|---|
| GSE114413 | Danio rerio | May 31 2026 | PRJNA471313 | retina |
| GSE266526 | Homo sapiens | Sep 02 2024 | PRJNA1107509 | retinal organoid |
| GSE211344 | Mus musculus | Aug 19 2022 | PRJNA869926 | Retina |
| GSE275131 | Mus musculus | Jan 22 2026 | PRJNA1149661 | Brain |
| GSE297342 | Homo sapiens | May 20 2025 | PRJNA1263555 | Retina |
| GSE223340 | Danio rerio | Dec 08 2023 | PRJNA925841 | larvae |
| GSE300209 | Mus musculus | Feb 10 2026 | PRJNA1279336 | liver |
| GSE295598 | Mus musculus | Feb 26 2026 | PRJNA1255234 | lung |
| GSE312648 | Homo sapiens | Mar 09 2026 | PRJNA1375651 | |
| GSE294977 | Mus musculus | Jan 28 2026 | PRJNA1252126 | lymph node |
| GSE306823 | Homo sapiens | Jan 14 2026 | PRJNA1312461 | |
| GSE317348 | Mus musculus | Jan 22 2026 | PRJNA1406772 | Brain |
| GSE316154 | Homo sapiens | Jan 22 2026 | PRJNA1401729 | Brain |
| GSE267570 | Homo sapiens | Jan 24 2026 | PRJNA1111877 | ICM |
| GSE316984 | Homo sapiens | Jan 27 2026 | PRJNA1405633 | ma around benign epithelium |
| GSE317716 | Mus musculus | Jan 31 2026 | PRJNA1415091 | colon |
| GSE207158 | Homo sapiens | Oct 01 2022 | PRJNA853957 | |
| GSE206344 | Toxoplasma gondii | Jun 20 2022 | PRJNA850353 | |
| GSE270212 | Homo sapiens | Jan 26 2026 | PRJNA1125416 | |
| GSE266332 | Mus musculus | Jan 29 2025 | PRJNA1106775 | Retina |
| GSE319684 | Mus musculus | Feb 23 2026 | PRJNA1424955 | Lung |
| GSE318498 | Mus musculus | Feb 06 2026 | PRJNA1419041 | |
| GSE237299 | Homo sapiens | Jan 26 2026 | PRJNA994571 | Bone marrow |
| GSE312574 | Homo sapiens;Mus musculus | Feb 17 2026 | PRJNA1374195 | |
| GSE313277 | Danio rerio | Dec 15 2025 | PRJNA1378864 | Retina |
| GSE149386 | Mus musculus | Aug 26 2020 | PRJNA628602 | |
| GSE308603 | Arabidopsis thaliana | Jan 05 2026 | PRJNA1331691 | seeding |
| GSE273827 | Homo sapiens | Jan 30 2026 | PRJNA1143424 | |
| GSE263150 | Mus musculus | Feb 11 2026 | PRJNA1095980 | bone marrow |
| GSE222535 | Mus musculus | Jan 01 2026 | PRJNA922485 | 2-cell embryo |
| GSE259428 | Mus musculus | Feb 06 2026 | PRJNA1081458 | Mouse embryos |
| GSE221862 | Homo sapiens | Dec 28 2022 | PRJNA916507 | retina |
| GSE319840 | Homo sapiens | Feb 18 2026 | PRJNA1425300 | Breast cancer |
| GSE159357 | Homo sapiens | Oct 06 2021 | PRJNA668654 | Retina periphery |
| GSE230616 | Mus musculus | Jan 21 2026 | PRJNA962013 | Liver |
| GSE308156 | Mus musculus | Jan 23 2026 | PRJNA1329046 | |
| GSE71364 | Mus musculus | Jul 31 2020 | PRJNA291052 | retina |
| GSE158629 | Homo sapiens | Dec 25 2025 | PRJNA666014 | etinal pigment epithelium |
| GSE159435 | Homo sapiens | Jun 16 2021 | PRJNA668929 | |
| GSE153874 | Mus musculus | Jun 22 2021 | PRJNA644380 | retina |
| GSE302808 | Mus musculus | Feb 18 2026 | PRJNA1292202 | |
| GSE303385 | Mus musculus | Jan 01 2026 | PRJNA1294984 | lung |
| GSE223315 | Mus musculus | Feb 26 2026 | PRJNA925569 | Bone marrow |
| GSE309269 | Homo sapiens | Feb 18 2026 | PRJNA1334994 | oblastoma from primary adrenal mass |
| GSE131534 | Mus musculus | Jun 03 2020 | PRJNA543932 | retina |
| GSE297475 | Homo sapiens | Jan 21 2026 | PRJNA1264969 | |
| GSE180529 | Mus musculus | May 06 2025 | PRJNA748598 | |
| GSE202053 | Drosophila melanogaster | Dec 31 2022 | PRJNA834097 | |
| GSE284055 | Mus musculus | Jan 08 2026 | PRJNA1196914 | Colon tumor |
| GSE309776 | Mus musculus | Feb 23 2026 | PRJNA1363157 | liver |
| GSE253869 | Mus musculus | Jan 20 2025 | PRJNA1067590 | Liver peri-tumor tissue |
| GSE316429 | Homo sapiens | Feb 19 2026 | PRJNA1402850 | |
| GSE300987 | Homo sapiens | Feb 20 2026 | PRJNA1282959 | blood |
| GSE300509 | Toxoplasma gondii ME49 | Feb 25 2026 | PRJNA1281083 | es from an infected mouse brain |
| GSE308042 | Mus musculus | Jan 12 2026 | PRJNA1328629 | Retina |
| GSE255516 | Homo sapiens | Jan 26 2026 | PRJNA1075141 | Lung |
| GSE292142 | Homo sapiens | Jan 26 2026 | PRJNA1237271 | Primary Tumour |
| GSE312457 | Mus musculus | Jan 21 2026 | PRJNA1373345 | Fallopian tube |
| GSE319954 | Homo sapiens | Feb 20 2026 | | |
| GSE304577 | Homo sapiens | Feb 24 2026 | PRJNA1301822 | PBMC |
| GSE315292 | Homo sapiens | Feb 09 2026 | PRJNA1395600 | plasma |
| GSE279304 | Homo sapiens | Feb 10 2026 | PRJNA1171825 | |
| GSE293123 | Mus musculus | Sep 07 2025 | PRJNA1242849 | Retina |

| | | | | |
|---|---|---|---|---|
| GSE247157 | Homo sapiens | Nov 07 2023 | PRJNA1036530 | macula lutea |
| GSE301077 | Mus musculus | Mar 19 2026 | PRJNA1283228 | |
| GSE214339 | Homo sapiens | Sep 25 2025 | PRJNA885026 | |
| GSE311091 | Mus musculus | Feb 05 2026 | PRJNA1368652 | heart |
| GSE273328 | Gallus gallus | Jul 16 2025 | PRJNA1141430 | hypothalamus |
| GSE286267 | Mus musculus | Jan 01 2026 | PRJNA1208503 | thymus |
| GSE295358 | omys tridecemlineatus;Mus mus | Apr 28 2025 | PRJNA1254267 | P5 retina |
| GSE316967 | Glycine max | Jan 20 2026 | PRJNA1405618 | leaves |
| GSE136564 | Mus musculus | Jan 08 2020 | PRJNA562698 | retina |
| GSE220624 | Homo sapiens | Sep 15 2023 | PRJNA910552 | retinal organoid |
| GSE285852 | Mus musculus | Feb 21 2026 | | None |
| GSE295125 | Mus musculus | Feb 27 2026 | PRJNA1253603 | kidney |
| GSE298471 | Homo sapiens | Jan 14 2026 | PRJNA1269509 | |
| GSE237007 | Homo sapiens | Sep 26 2023 | PRJNA993408 | uman retinal organoid cells |
| GSE254839 | Mus musculus | Jul 10 2024 | PRJNA1072052 | |
| GSE249894 | Homo sapiens | Dec 01 2025 | PRJNA1051102 | retinas |
| GSE277649 | Mus musculus | Jan 16 2026 | PRJNA1163279 | splenocytes |
| GSE237030 | Mus musculus | Feb 26 2026 | PRJNA993573 | in vivo gonad |
| GSE315365 | Mus musculus | Jan 06 2026 | PRJNA1396595 | whole heart |
| GSE311479 | Mus musculus | Jan 20 2026 | PRJNA1369530 | ale Tibialis Anterior Muscle |
| GSE157739 | Mus musculus | Sep 06 2021 | PRJNA662598 | Kidney |
| GSE309845 | Homo sapiens | Jan 31 2026 | PRJNA1363882 | |
| GSE277361 | Homo sapiens | Jan 15 2025 | PRJNA1162098 | Macula |
| GSE287553 | Mus musculus | Jan 06 2026 | PRJNA1213730 | |
| GSE318153 | Mus musculus | Jan 31 2026 | PRJNA1416808 | Subcutaneous tumor |
| GSE286250 | Mus musculus | Jan 13 2025 | PRJNA1208474 | Brain organoid |
| GSE285389 | Homo sapiens | Feb 11 2026 | PRJNA1203262 | cell line |
| GSE310896 | Anableps anableps | Apr 08 2026 | PRJNA1367503 | Dorsal retina |
| GSE156758 | Rattus norvegicus | Aug 21 2021 | PRJNA658979 | |
| GSE298324 | Mus musculus | Jan 28 2026 | PRJNA1268967 | B16-F0 melanoma |
| GSE161893 | Homo sapiens;Mus musculus | Sep 30 2021 | PRJNA679792 | Endothelial cells |
| GSE283880 | Mus musculus | Dec 10 2024 | PRJNA1195953 | embryonic day 14 retina |
| GSE270198 | Mus musculus | Dec 31 2024 | PRJNA1125399 | Brain |
| GSE312483 | Mus musculus | Feb 11 2026 | PRJNA1374055 | |
| GSE128504 | Mus musculus | Sep 29 2020 | PRJNA527876 | |
| GSE161102 | Mus musculus | Apr 28 2021 | PRJNA675586 | |
| GSE298565 | Homo sapiens | Feb 06 2026 | PRJNA1269946 | |
| GSE293866 | Homo sapiens;Mus musculus | Jan 29 2026 | PRJNA1246627 | |
| GSE252465 | Homo sapiens | Feb 07 2026 | PRJNA1060696 | Lung BALF |
| GSE310024 | Homo sapiens | Feb 11 2026 | PRJNA1365026 | Lung; Pleural effusion |
| GSE298718 | Homo sapiens | Feb 05 2026 | PRJNA1271001 | |
| GSE291550 | Scophthalmus maximus | Dec 04 2025 | PRJNA1234131 | Eye |
| GSE315524 | Oryctolagus cuniculus | Jan 14 2026 | PRJNA1398213 | corneal-limbal |
| GSE287470 | Mus musculus | Jan 17 2026 | PRJNA1213060 | Brain |
| GSE298230 | Drosophila melanogaster | Jan 13 2026 | PRJNA1268542 | |
| GSE300964 | Mus musculus | Jan 10 2026 | PRJNA1282738 | hippocampus |
| GSE235583 | Homo sapiens | Feb 15 2024 | PRJNA986509 | organoid |
| GSE279485 | Mus musculus | Jan 20 2026 | PRJNA1173125 | Pancreatic islet |
| GSE318047 | Homo sapiens | Feb 16 2026 | PRJNA1416379 | rebral cortex - glioblastoma |
| GSE317825 | Homo sapiens | Feb 09 2026 | PRJNA1415369 | colon |
| GSE236627 | Mus musculus | Jan 25 2024 | PRJNA991920 | retina |
| GSE244572 | Homo sapiens | Jun 28 2025 | PRJNA1023589 | RPE |
| GSE294705 | Homo sapiens | Feb 07 2026 | PRJNA1251344 | Male |
| GSE309181 | Mus musculus | Feb 25 2026 | PRJNA1334288 | emisphere (without cerebellum) |
| GSE263840 | Homo sapiens | Feb 09 2026 | PRJNA1099581 | |
| GSE237065 | Drosophila melanogaster | Feb 05 2026 | PRJNA993655 | hemocyte |
| GSE226780 | Mus musculus | Jun 05 2024 | PRJNA941790 | Retina |
| GSE290158 | Arabidopsis thaliana | Jan 29 2026 | PRJNA1226490 | Seedlings |
| GSE182504 | Mus musculus | Aug 27 2021 | PRJNA756533 | |
| GSE283578 | Homo sapiens | Dec 09 2024 | PRJNA1194563 | retina |
| GSE307982 | Mus musculus | Feb 13 2026 | PRJNA1327918 | |
| GSE308682 | Homo sapiens | Feb 11 2026 | PRJNA1332459 | Blood |
| GSE225346 | Mus musculus | Feb 08 2026 | PRJNA935314 | embryo |

| | | | | |
|---|---|---|---|---|
| GSE253572 | Homo sapiens;Rattus norvegicus | Feb 26 2026 | | Breast invasive carcinoma |
| GSE315452 | Mus musculus | Jan 18 2026 | PRJNA1397170 | spleen |
| GSE147979 | Homo sapiens;Sus scrofa | Jul 29 2021 | PRJNA622707 | Choroid |
| GSE319605 | Homo sapiens | Feb 20 2026 | PRJNA1424716 | heart |
| GSE290325 | Mus musculus | Feb 09 2026 | PRJNA1227484 | Liver |
| GSE317288 | Mus musculus | Jan 27 2026 | PRJNA1406689 | |
| GSE307746 | Homo sapiens | Feb 19 2026 | PRJNA1327251 | ronchoalveolar lavage cells |
| GSE253545 | Homo sapiens | Jan 18 2026 | PRJNA1066272 | Adult Heart |
| GSE316932 | Caenorhabditis elegans | Jan 27 2026 | | Whole animal |
| GSE310165 | Homo sapiens | Jan 20 2026 | PRJNA1365226 | lung cancer |
| GSE309827 | Homo sapiens | Feb 25 2026 | PRJNA1363778 | |
| GSE303738 | Mus musculus | Jan 07 2026 | PRJNA1297192 | Tumor lymph node |
| GSE290470 | Mus musculus | Mar 14 2025 | | retina |
| GSE317953 | Arabidopsis thaliana | Feb 18 2026 | PRJNA1416082 | 14-day-old seedlings |
| GSE287253 | Mus musculus | Jan 21 2025 | PRJNA1211691 | organoids |
| GSE263877 | Mus musculus | Jun 06 2024 | PRJNA1099665 | retina |
| GSE200049 | Mus musculus | Apr 03 2025 | PRJNA822803 | RPE/choroid/sclera |
| GSE316811 | Homo sapiens;Mus musculus | Jan 22 2026 | PRJNA1404279 | liver |
| GSE245546 | Mus musculus | Jun 06 2024 | PRJNA1028976 | retina |
| GSE298989 | Homo sapiens | Feb 15 2026 | PRJNA1272069 | cell line |
| GSE222497 | Homo sapiens | Jan 01 2026 | PRJNA922207 | plasma in POI group |
| GSE297396 | Homo sapiens | Feb 20 2026 | PRJNA1263754 | |
| GSE227197 | Homo sapiens | Mar 17 2024 | PRJNA944112 | |
| GSE160663 | Danio rerio | Dec 01 2020 | PRJNA673887 | eyes |
| GSE215193 | Mus musculus | Oct 13 2022 | PRJNA889067 | retina |
| GSE137311 | Homo sapiens;Mus musculus | Mar 03 2020 | PRJNA573850 | |
| GSE277708 | Homo sapiens | Jan 14 2026 | PRJNA1163401 | |
| GSE287347 | Homo sapiens;Mus musculus | Feb 25 2026 | PRJNA1212168 | |
| GSE277217 | Homo sapiens | May 14 2025 | PRJNA1161386 | |
| GSE319044 | Homo sapiens | Feb 12 2026 | PRJNA1422011 | Lung |
| GSE282125 | Homo sapiens | Jan 20 2026 | PRJNA1187245 | Breast ( Breast Cancer) |
| GSE295219 | Homo sapiens | Jan 14 2026 | PRJNA1253718 | cell line |
| GSE300588 | Mus musculus | Jul 16 2025 | PRJNA1281515 | Retina |
| GSE308952 | Mus musculus | Jan 09 2026 | PRJNA1333483 | Regenerating skin wound |
| GSE303451 | Homo sapiens;Mus musculus | Feb 28 2026 | PRJNA1295446 | lung |
| GSE295706 | Homo sapiens | Feb 09 2026 | PRJNA1255643 | Skin |
| GSE286326 | Sus scrofa | Feb 27 2026 | PRJNA1208924 | Liver |
| GSE243839 | Mus musculus | Jan 12 2026 | PRJNA1020133 | |
| GSE290206 | Drosophila melanogaster | Jan 19 2026 | PRJNA1226571 | Brain |
| GSE269683 | Mus musculus | Feb 27 2025 | PRJNA1123222 | |
| GSE304509 | Canis lupus familiaris | Jan 08 2026 | PRJNA1301403 | |
| GSE290084 | Arabidopsis thaliana | Jan 29 2026 | PRJNA1226181 | Seedlings |
| GSE197421 | Mus musculus | Sep 23 2022 | PRJNA810185 | |
| GSE301009 | Drosophila melanogaster | Jan 09 2026 | PRJNA1283059 | proventriculus |
| GSE189555 | Mus musculus | Apr 13 2022 | PRJNA783568 | RPE-choroid |
| GSE279665 | Mus musculus | Feb 20 2026 | PRJNA1174055 | Brain |
| GSE302841 | Bos taurus | Jan 15 2026 | PRJNA1292253 | |
| GSE285562 | Homo sapiens | Sep 03 2025 | PRJNA1204363 | |
| GSE288036 | Mus musculus | Jan 26 2025 | PRJNA1215754 | retina+choroid |
| GSE235990 | Homo sapiens;Rattus norvegicus | Mar 13 2024 | PRJNA988130 | rat choroid |
| GSE279107 | Drosophila melanogaster | Jan 27 2026 | PRJNA1170876 | Head |
| GSE259220 | Homo sapiens;Mus musculus | Mar 01 2024 | | Human embryonic kidney cells |
| GSE310928 | Homo sapiens | Feb 24 2026 | PRJNA1367551 | |
| GSE293317 | Homo sapiens | Feb 16 2026 | PRJNA1244011 | cell line |
| GSE297114 | Homo sapiens | Feb 26 2026 | PRJNA1262733 | |
| GSE222615 | Mus musculus | Jan 01 2026 | PRJNA922877 | |
| GSE284749 | Mus musculus | Jan 05 2026 | PRJNA1200601 | testis |
| GSE317401 | Dermatophagoides farinae | Feb 15 2026 | PRJNA1412510 | whole organism |
| GSE200195 | Mus musculus | Nov 11 2022 | PRJNA823657 | Retina |
| GSE308276 | Mus musculus | Jan 14 2026 | PRJNA1330051 | gastrocnemius muscle |
| GSE288990 | Mus musculus | Jan 05 2026 | PRJNA1220174 | Kidney |
| GSE284108 | Mus musculus | Jan 15 2026 | PRJNA1197366 | muscle and myenteric plexus (ileum) |
| GSE267167 | Mus musculus | Feb 23 2026 | PRJNA1109991 | mammary gland |

| | | | | |
|---|---|---|---|---|
| GSE301360 | Mus musculus | Feb 17 2026 | PRJNA1284847 | |
| GSE314399 | Homo sapiens | Feb 24 2026 | PRJNA1390829 | xenograft tumor |
| GSE280614 | Mus musculus | Feb 18 2026 | PRJNA1179494 | |
| GSE308128 | Homo sapiens | Jan 04 2026 | PRJNA1328789 | |
| GSE314686 | Mus musculus | Feb 24 2026 | PRJNA1392462 | |
| GSE315814 | Homo sapiens | Jan 14 2026 | PRJNA1399347 | |
| GSE303934 | Homo sapiens | Jan 01 2026 | PRJNA1298256 | Tumor tissue |
| GSE285476 | Rattus norvegicus | Jan 14 2026 | PRJNA1203900 | liver |
| GSE57699 | Mus musculus | Dec 01 2024 | PRJNA247761 | liver |
| GSE304893 | Homo sapiens | Feb 26 2026 | PRJNA1303186 | male |
| GSE325526 | ns;Macaca fascicularis;Mus musc | Mar 25 2026 | PRJNA1440402 | Eye,Eye |
| GSE289616 | Rhizopus microsporus | Feb 28 2026 | PRJNA1223826 | Mycelium |
| GSE278045 | Homo sapiens | Jan 28 2026 | PRJNA1165271 | |
| GSE249809 | Homo sapiens | Feb 09 2026 | PRJNA1050774 | |
| GSE300629 | Mus musculus | Jan 05 2026 | | Cerebellum 1 |
| GSE283098 | Homo sapiens | Jan 27 2026 | PRJNA1191862 | ligament |
| GSE317515 | Mus musculus | Jan 30 2026 | PRJNA1414401 | liver |
| GSE319198 | Homo sapiens | Feb 18 2026 | PRJNA1422613 | skin |
| GSE301648 | Mus musculus | Feb 10 2026 | PRJNA1285918 | |
| GSE246574 | Mus musculus | Nov 03 2023 | PRJNA1033654 | Retinas |
| GSE299803 | Saccharomyces cerevisiae | Jan 01 2026 | PRJNA1276559 | S.cerevisiae |
| GSE313074 | Homo sapiens | Feb 09 2026 | | postmortem brain |
| GSE264596 | Homo sapiens | Feb 09 2026 | PRJNA1103207 | Bone marrow |
| GSE186585 | Homo sapiens | Mar 13 2023 | PRJNA774659 | |
| GSE248868 | Mus musculus | Dec 26 2023 | PRJNA1046136 | ched RGC from mouse retina |
| GSE206625 | Mus musculus | Aug 20 2022 | PRJNA851596 | retina |
| GSE269812 | Mus musculus | Jun 28 2024 | PRJNA1123731 | Optic Nerve |
| GSE304716 | Oncorhynchus mykiss | Jan 08 2026 | PRJNA1302392 | (freshly fertilized egg/zygote) |
| GSE245962 | Mus musculus | Jan 01 2026 | PRJNA1030758 | plasms of the mouse mammary gland |
| GSE290784 | Mus musculus | Feb 27 2026 | PRJNA1230107 | Placenta |
| GSE255268 | Homo sapiens | May 29 2024 | PRJNA1074306 | |
| GSE300768 | Homo sapiens | Jan 04 2026 | PRJNA1282069 | Pancreas; Duct |
| GSE319948 | Mus musculus | Feb 19 2026 | PRJNA1425579 | Heart |
| GSE314092 | Homo sapiens | Jan 19 2026 | PRJNA1390080 | |
| GSE303399 | Mus musculus | Jul 23 2025 | PRJNA1295347 | ganglion cell layer |
| GSE149791 | Mus musculus | Apr 30 2025 | PRJNA630164 | retina |
| GSE234035 | Homo sapiens | Nov 29 2023 | PRJNA979220 | Eye |
| GSE242163 | Mus musculus | Jan 21 2026 | PRJNA1011882 | brain meningeal layer |
| GSE304043 | Mus musculus | Jun 11 2026 | PRJNA1298819 | |
| GSE138002 | Homo sapiens | May 07 2020 | PRJNA574132 | issociated retinal organoid |
| GSE165812 | Escherichia coli;Mus musculus | Jan 30 2021 | PRJNA697959 | retina |
| GSE318590 | Mus musculus | Feb 11 2026 | PRJNA1419267 | |
| GSE274888 | Mus musculus | Jun 19 2025 | PRJNA1148409 | retina |
| GSE157816 | Danio rerio | Sep 12 2020 | PRJNA662909 | |
| GSE267026 | Homo sapiens | Feb 05 2026 | PRJNA1109362 | heral Blood Mononuclear Cells |
| GSE316243 | Mus musculus | Feb 24 2026 | PRJNA1401854 | Distal lung |
| GSE174538 | Danio rerio | Mar 01 2022 | PRJNA730498 | |
| GSE168789 | Mus musculus | Mar 24 2022 | PRJNA713995 | |
| GSE317540 | Larix kaempferi | Jan 27 2026 | PRJNA1414440 | plant tissue cells |
| GSE276182 | Mus musculus | Jan 20 2026 | PRJNA1155649 | brain cortex |
| GSE309434 | Lambdavirus lambda | Jan 12 2026 | PRJNA1336081 | |
| GSE328981 | Homo sapiens | Jun 01 2026 | PRJNA1457319 | |
| GSE202473 | Homo sapiens | Sep 09 2022 | PRJNA836267 | Adult Retina |
| GSE290690 | Homo sapiens | Feb 27 2026 | PRJNA1229573 | |
| GSE291802 | Homo sapiens | Jan 13 2026 | PRJNA1235270 | |
| GSE235177 | Homo sapiens | Jan 12 2026 | PRJNA984717 | Blood |
| GSE317222 | Homo sapiens | Jan 26 2026 | PRJNA1406288 | cell culture |
| GSE305890 | Mus musculus | Jan 21 2026 | PRJNA1308124 | lymph node |
| GSE173180 | Homo sapiens | May 17 2023 | PRJNA724040 | |
| GSE271012 | Mus musculus | Jan 12 2026 | PRJNA1129274 | spleens |
| GSE316009 | Homo sapiens | Feb 26 2026 | PRJNA1400450 | |
| GSE274512 | Mus musculus | Feb 15 2026 | PRJNA1146915 | |
| GSE316516 | Mus musculus | Jan 19 2026 | PRJNA1403048 | brain |

| | | | | |
|---|---|---|---|---|
| GSE319387 | Homo sapiens | Feb 14 2026 | PRJNA1423213 | |
| GSE279559 | Caenorhabditis elegans | Jan 12 2026 | PRJNA1173493 | Whole animal |
| GSE281142 | Mus musculus | Jan 10 2026 | PRJNA1182370 | Brain |
| GSE293503 | Mus musculus | Feb 23 2026 | PRJNA1244833 | Kidney |
| GSE317577 | Mus musculus | Jan 27 2026 | PRJNA1414493 | Lung |
| GSE296003 | Homo sapiens | Jan 21 2026 | PRJNA1257306 | Prostate |
| GSE309403 | Homo sapiens | Jan 27 2026 | PRJNA1336040 | CSF |
| GSE303792 | Macaca mulatta | Feb 25 2026 | PRJNA1297689 | Granuloma |
| GSE314820 | Oryctolagus cuniculus | Feb 25 2026 | PRJNA1392647 | Terminal ileum |
| GSE127761 | Homo sapiens | Apr 22 2020 | PRJNA525342 | vitreous |
| GSE160099 | Homo sapiens | Dec 22 2020 | PRJNA671828 | |
| GSE315850 | Drosophila melanogaster | Jan 15 2026 | PRJNA1399422 | whole gut |
| GSE263320 | Sus scrofa | Jan 25 2026 | PRJNA1096703 | dorsal root ganglion |
| GSE281153 | Mus musculus | Jan 12 2026 | PRJNA1182391 | Prostate |
| GSE245850 | Mus musculus | Jan 05 2026 | PRJNA1030083 | nadal white adipose tissue |
| GSE118542 | Rattus norvegicus | Aug 01 2021 | PRJNA485980 | |
| GSE309370 | Mus musculus | Jan 22 2026 | PRJNA1335251 | |
| GSE243996 | Rattus norvegicus | Sep 08 2024 | PRJNA1020857 | Reference |
| GSE286289 | Mus musculus | Feb 17 2026 | PRJNA1208537 | |
| GSE314831 | Mus musculus | Feb 18 2026 | PRJNA1392906 | bcutaneous adipose tissue |
| GSE222538 | Homo sapiens | Jan 05 2026 | PRJNA922483 | |
| GSE259425 | Mus musculus | Feb 06 2026 | PRJNA1081462 | Mouse embryos |
| GSE269599 | Mus musculus | Jan 01 2025 | PRJNA1122598 | eye cup |
| GSE222745 | Mus musculus | Jan 01 2026 | | |
| GSE303947 | Mus musculus | Feb 25 2026 | PRJNA1298285 | Retina |
| GSE173800 | Mus musculus | Feb 06 2026 | PRJNA727085 | Liver |
| GSE325169 | Mus musculus | Mar 22 2026 | PRJNA1438744 | retina |
| GSE313671 | Mus musculus | Jan 14 2026 | PRJNA1380503 | Lung |
| GSE298094 | Homo sapiens | Jan 09 2026 | PRJNA1268338 | Prostate |
| GSE213948 | Homo sapiens | May 30 2023 | PRJNA883248 | |
| GSE274899 | Mus musculus | Jan 31 2026 | PRJNA1148426 | Liver |
| GSE318581 | Homo sapiens | Feb 05 2026 | PRJNA1419252 | prostate cancer |
| GSE287181 | Mus musculus | Jan 15 2026 | PRJNA1211569 | bone marrow |
| GSE292178 | Homo sapiens | Feb 10 2026 | PRJNA1237340 | NPC generated from iPSC |
| GSE301230 | Gallus gallus | Jan 12 2026 | PRJNA1284303 | neural tube |
| GSE300533 | Mus musculus | Jan 12 2026 | PRJNA1281110 | Liver |
| GSE253853 | Mus musculus | Jan 30 2026 | PRJNA1067560 | |
| GSE315973 | Mus musculus | Jan 08 2026 | PRJNA1400045 | Whole Placenta |
| GSE272968 | Mus musculus | Jan 16 2026 | PRJNA1139725 | mammary gland |
| GSE141081 | Homo sapiens | Jun 11 2020 | PRJNA591952 | |
| GSE300504 | Danio rerio | Jul 02 2026 | PRJNA1281084 | otic vesicle |
| GSE318938 | Mus musculus | Feb 11 2026 | | spinal cord |
| GSE317327 | Homo sapiens | Jan 27 2026 | PRJNA1406746 | Achilles tendon |
| GSE301207 | Homo sapiens | Feb 09 2026 | PRJNA1284270 | |
| GSE313646 | Mus musculus | Feb 18 2026 | PRJNA1380458 | |
| GSE162993 | Mus musculus | Dec 24 2020 | PRJNA687474 | retina |
| GSE246346 | Homo sapiens;Mus musculus | Sep 05 2025 | | |
| GSE311583 | Mus musculus | Jan 27 2026 | PRJNA1369968 | brain ctx |
| GSE313359 | Mus musculus | Jan 27 2026 | PRJNA1379317 | Hippocampus |
| GSE210137 | Mus musculus | Apr 11 2023 | PRJNA864030 | Retina |
| GSE289259 | Allium mongolicum | Feb 08 2026 | PRJNA1222282 | Root |
| GSE310280 | Mus musculus | Jan 01 2026 | PRJNA1365693 | Stellate Ganglion |
| GSE283089 | Mus musculus | Jan 31 2025 | PRJNA1191850 | nal pigment epithelium (RPE) |
| GSE289318 | Mus musculus | Feb 19 2026 | PRJNA1222408 | Intestinal Mesenchyme |
| GSE233797 | Mus musculus | Aug 01 2023 | PRJNA977994 | Retina |
| GSE288895 | Homo sapiens | Feb 28 2026 | PRJNA1219841 | |
| GSE292273 | Homo sapiens | Jan 05 2026 | PRJNA1237812 | |
| GSE319496 | Homo sapiens | Feb 14 2026 | PRJNA1423615 | Whole blood |
| GSE317266 | Homo sapiens | Jan 22 2026 | PRJNA1406636 | Kidney |
| GSE297980 | Gallus gallus | Feb 09 2026 | PRJNA1267072 | spinal cord |
| GSE304359 | Mus musculus | Jan 01 2026 | PRJNA1300793 | DRG |
| GSE303912 | Homo sapiens | Feb 11 2026 | PRJNA1298218 | plasma-derived EV |
| GSE243065 | Homo sapiens;Mus musculus | Feb 24 2026 | PRJNA1016317 | embryo |

| | | | | |
|---|---|---|---|---|
| GSE313065 | Mus musculus | Jan 15 2026 | PRJNA1378204 | Peritoneal cavity |
| GSE246552 | Homo sapiens | Jan 01 2026 | PRJNA1033621 | |
| GSE269786 | Mus musculus | Jan 01 2026 | PRJNA1123691 | bone marrow |
| GSE308344 | Homo sapiens | Jan 19 2026 | PRJNA1330235 | Kidney |
| GSE301413 | Homo sapiens | Feb 26 2026 | PRJNA1284952 | |
| GSE135406 | Danio rerio;Mus musculus | May 01 2020 | PRJNA558770 | Retina |
| GSE302944 | Mus musculus | Jan 16 2026 | PRJNA1292872 | guinal white adipose tissue |
| GSE287872 | Homo sapiens | Jan 17 2026 | PRJNA1214838 | |
| GSE303488 | Homo sapiens | Feb 24 2026 | PRJNA1295509 | |
| GSE316471 | Homo sapiens | Jan 21 2026 | PRJNA1402966 | Bone marrow |
| GSE256188 | Homo sapiens | Feb 22 2026 | PRJNA1078384 | Colorectal cancer |
| GSE234724 | Homo sapiens | Feb 25 2026 | PRJNA982820 | |
| GSE192665 | Homo sapiens | Jun 21 2023 | PRJNA792579 | |
| GSE279121 | Aspergillus flavus | Feb 12 2026 | PRJNA1170915 | mycelium |
| GSE254752 | Mus musculus | Jan 31 2026 | PRJNA1071624 | ganglion |
| GSE309206 | Homo sapiens | Jan 06 2026 | PRJNA1334327 | |
| GSE303197 | Homo sapiens | Feb 05 2026 | PRJNA1294318 | peripheral blood |
| GSE310939 | Homo sapiens | Feb 24 2026 | PRJNA1367578 | |
| GSE293306 | Mus musculus | May 12 2026 | PRJNA1243982 | Retina |
| GSE163754 | Mus musculus | Dec 24 2020 | PRJNA687423 | retina |
| GSE312881 | Homo sapiens | Feb 15 2026 | PRJNA1377546 | iPSC-DSN |
| GSE239512 | Homo sapiens | Aug 31 2023 | PRJNA999619 | monocytes |
| GSE266034 | Homo sapiens | Jan 16 2026 | PRJNA1105372 | |
| GSE279643 | Homo sapiens | Feb 27 2026 | PRJNA1173754 | Bone marrow |
| GSE290933 | Mus musculus | Apr 17 2025 | PRJNA1231016 | retina |
| GSE293464 | Homo sapiens | Mar 22 2026 | PRJNA1244746 | retinal organoid |
| GSE314970 | Rattus norvegicus | Mar 05 2026 | | left atrium |
| GSE211349 | Homo sapiens | Aug 19 2022 | PRJNA869937 | |
| GSE244970 | Mus musculus | May 14 2024 | PRJNA1026540 | cerebellum |
| GSE278468 | Mus musculus | Jan 31 2026 | PRJNA1167319 | tches (30 mice) and blood (20 mice) |
| GSE162060 | Homo sapiens;Mus musculus | Jun 30 2021 | PRJNA680481 | |
| GSE296153 | Macaca mulatta | Feb 28 2026 | PRJNA1257933 | ACC |
| GSE254518 | Mus musculus | Jan 26 2026 | PRJNA1070699 | substantia nigra |
| GSE291289 | Mus musculus | Feb 19 2026 | PRJNA1232726 | |
| GSE312438 | Escherichia coli | Jan 21 2026 | PRJNA1373299 | |
| GSE142658 | Homo sapiens | Dec 26 2020 | PRJNA597881 | |
| GSE301265 | Homo sapiens | Feb 09 2026 | PRJNA1284371 | ntestinal neuroendocrine tumor |
| GSE189739 | Danio rerio | May 12 2023 | PRJNA784293 | |
| GSE317345 | Homo sapiens | Jan 28 2026 | PRJNA1406769 | tumor |
| GSE282349 | Homo sapiens | Feb 18 2026 | PRJNA1188171 | |
| GSE224068 | Mus musculus | Feb 21 2024 | PRJNA929506 | |
| GSE307121 | Acinetobacter baumannii | Jan 01 2026 | PRJNA1314615 | bacterial culture |
| GSE314501 | Mus musculus | Jan 23 2026 | PRJNA1391411 | Embryo |
| GSE290575 | Homo sapiens | Jan 23 2026 | PRJNA1228648 | |
| GSE201350 | Mus musculus | Dec 31 2022 | PRJNA830919 | |
| GSE289928 | Homo sapiens | Feb 23 2026 | PRJNA1225326 | Bone Marrow |
| GSE277564 | Danio rerio | Jan 01 2026 | PRJNA1162813 | pronephric duct |
| GSE286337 | Mus musculus | Jan 16 2026 | PRJNA1208941 | intestine |
| GSE292843 | Mus musculus | Feb 23 2026 | PRJNA1241719 | Kidney |
| GSE319228 | Homo sapiens | Feb 16 2026 | PRJNA1422639 | Leiomyoma |
| GSE308943 | Rattus norvegicus | Jan 28 2026 | PRJNA1333466 | islet |
| GSE193808 | Mus musculus | Dec 17 2022 | PRJNA798019 | RPE-sclera complex |
| GSE148263 | Homo sapiens | Sep 18 2020 | PRJNA623614 | |
| GSE306468 | Mus musculus | Feb 08 2026 | PRJNA1310867 | Spinal cord |
| GSE316191 | Mus musculus | Jan 16 2026 | PRJNA1401786 | splenic CD8+ T cells |
| GSE290220 | Homo sapiens | Jan 20 2026 | PRJNA1226596 | rectum |
| GSE240735 | Homo sapiens | Jan 09 2026 | PRJNA1005178 | Human embryonic kidney |
| GSE263945 | Mus musculus | Jan 08 2026 | PRJNA1100436 | |
| GSE314029 | Mus musculus | Feb 14 2026 | | Hypothalamus |
| GSE277231 | Mus musculus | Jan 01 2026 | PRJNA1161403 | |
| GSE175394 | Mus musculus | Aug 23 2021 | PRJNA732284 | Lens |
| GSE201678 | Homo sapiens | Apr 30 2022 | PRJNA832670 | |
| GSE291400 | Mus musculus | Feb 25 2026 | PRJNA1233295 | lung |

| | | | | |
|---|---|---|---|---|
| GSE290303 | Rattus norvegicus | Feb 28 2026 | PRJNA1227430 | re isolated from the hearts of newborn rats and cultured in vitro |
| GSE180814 | Mus musculus | Dec 03 2022 | PRJNA749619 | Retina |
| GSE272170 | Rattus norvegicus | Feb 05 2025 | PRJNA1135251 | Retina |
| GSE283723 | Mus musculus | Jan 01 2026 | PRJNA1195093 | pancreatic islets |
| GSE184386 | Homo sapiens | Dec 30 2021 | PRJNA764422 | |
| GSE275946 | Mus musculus | Jan 20 2026 | PRJNA1153931 | heart |
| GSE169519 | Mus musculus | May 04 2023 | PRJNA716964 | |
| GSE319141 | synthetic construct | Feb 11 2026 | PRJNA1422179 | |
| GSE286541 | Mus musculus | Jan 13 2026 | PRJNA1209885 | Brain |
| GSE269266 | Homo sapiens | Feb 10 2026 | PRJNA1120851 | hPSC-Islets |
| GSE315609 | Homo sapiens | Jan 09 2026 | PRJNA1398322 | PBMC |
| GSE298435 | Homo sapiens;Mus musculus | Feb 13 2026 | PRJNA1269436 | liver |
| GSE261232 | Gallus gallus | Jul 01 2024 | PRJNA1086001 | retina |
| GSE310309 | Homo sapiens | Feb 16 2026 | PRJNA1365738 | Endometrium |
| GSE298402 | Mus musculus | Jan 08 2026 | PRJNA1269089 | prostate |
| GSE311122 | Mus musculus | Jan 31 2026 | PRJNA1368708 | primary astrocyte |
| GSE205070 | Mus musculus | Aug 26 2022 | PRJNA843677 | BMDMs |
| GSE283569 | Mus musculus | Jan 23 2026 | PRJNA1194544 | liver |
| GSE282775 | Mus musculus | Dec 03 2024 | PRJNA1190394 | Retina |
| GSE318886 | Homo sapiens | Feb 12 2026 | PRJNA1420761 | |
| GSE318408 | Homo sapiens | Feb 15 2026 | PRJNA1418678 | |
| GSE237209 | Sus scrofa domesticus | Oct 01 2023 | PRJNA994173 | Retina |
| GSE244340 | Homo sapiens | Jun 24 2024 | PRJNA1022433 | cell line |
| GSE287517 | Mus musculus | Oct 29 2025 | PRJNA1213135 | E13.5 mouse embryos |
| GSE291437 | Homo sapiens | Jan 27 2026 | PRJNA1233596 | Human embryonic kidney |
| GSE308921 | Sus scrofa | Jan 01 2026 | PRJNA1333404 | Nucleus Pulposus |
| GSE294569 | Schmidtea mediterranea | Jan 06 2026 | | whole-body sample |
| GSE137128 | Mus musculus | Sep 01 2020 | PRJNA564599 | retina |
| GSE249755 | Danio rerio | Jun 21 2024 | PRJNA1050291 | eye |
| GSE295508 | Homo sapiens | Jan 14 2026 | PRJNA1255076 | PBMC |
| GSE300299 | Mus musculus | Feb 05 2026 | PRJNA1280064 | ankle |
| GSE160627 | Mus musculus | Jan 15 2021 | PRJNA673813 | |
| GSE315260 | Mus musculus | Jan 06 2026 | | |
| GSE318734 | Homo sapiens | Feb 23 2026 | PRJNA1419864 | pumlmonary epithelium |
| GSE300786 | Homo sapiens | Feb 12 2026 | | spinal cord |
| GSE269310 | Mus musculus | Jan 06 2026 | PRJNA1120926 | |
| GSE169170 | Homo sapiens | Jan 31 2026 | PRJNA715467 | serum |
| GSE285957 | Oryctolagus cuniculus | Dec 06 2025 | PRJNA1207048 | retina |
| GSE317803 | Mus musculus | Jan 30 2026 | PRJNA1415288 | Cerebral cortex |
| GSE299768 | Mus musculus | Feb 25 2026 | PRJNA1276469 | |
| GSE316691 | Histoplasma ohiense | Jan 26 2026 | PRJNA1403603 | |
| GSE302474 | Mus musculus | Feb 06 2026 | PRJNA1291072 | |
| GSE158585 | Plasmodium falciparum | Sep 26 2020 | PRJNA665719 | |
| GSE317192 | Homo sapiens | Jan 21 2026 | PRJNA1406248 | |
| GSE222386 | Homo sapiens | Jan 12 2023 | PRJNA921854 | retinal epithelium |
| GSE314236 | Caenorhabditis elegans | Feb 24 2026 | PRJNA1390510 | whole body |
| GSE310035 | Homo sapiens | Feb 25 2026 | PRJNA1365044 | |
| GSE228986 | Mus musculus | May 23 2023 | PRJNA952505 | Retina |
| GSE298709 | Homo sapiens | Feb 22 2026 | PRJNA1270988 | Blood |
| GSE287461 | Rattus norvegicus | Jan 25 2026 | PRJNA1213045 | Myocardial |
| GSE295103 | Mus musculus | Feb 19 2026 | PRJNA1253542 | Kidney |
| GSE317920 | Mus musculus | Jan 30 2026 | PRJNA1415825 | hippocampus microglia |
| GSE197937 | Gallus gallus | May 11 2022 | PRJNA812909 | |
| GSE301191 | Homo sapiens | Dec 04 2025 | PRJNA1284244 | |
| GSE314315 | Mus musculus | Jan 26 2026 | PRJNA1390704 | CT26 tumor tissues |
| GSE274945 | Homo sapiens | Feb 19 2026 | PRJNA1148599 | Melanoma |
| GSE230817 | Mus musculus | Jan 10 2024 | PRJNA962603 | |
| GSE310887 | Homo sapiens | Feb 20 2026 | PRJNA1367483 | |
| GSE310116 | Mus musculus | Feb 23 2026 | PRJNA1365156 | Lung |
| GSE160051 | Danio rerio | Jul 31 2021 | PRJNA671699 | |
| GSE319369 | Macaca fascicularis | Feb 16 2026 | PRJNA1423173 | Brain |
| GSE296290 | Bos taurus | Feb 25 2026 | PRJNA1258789 | Articular cartilage |
| GSE292902 | Mus musculus | Feb 26 2026 | PRJNA1241833 | testis |

| | | | | |
|---|---|---|---|---|
| GSE282075 | Arabidopsis thaliana | Feb 06 2026 | PRJNA1186953 | shoot |
| GSE319676 | Mus musculus | Feb 18 2026 | PRJNA1424947 | |
| GSE277425 | Mus musculus | Jan 13 2026 | PRJNA1162150 | spleen |
| GSE289869 | Homo sapiens | Feb 15 2026 | PRJNA1225223 | Pterygium |
| GSE172138 | Homo sapiens | Mar 03 2023 | PRJNA722233 | retinal organoids |
| GSE286514 | Homo sapiens | Jan 12 2026 | PRJNA1209838 | |
| GSE318308 | Homo sapiens | Feb 06 2026 | PRJNA1418576 | Ovarian tumor |
| GSE148168 | Homo sapiens | Oct 25 2021 | PRJNA623366 | |
| GSE241140 | Mus musculus | Feb 20 2026 | PRJNA1006635 | Cortex |
| GSE285843 | Homo sapiens | Jan 01 2026 | PRJNA1206194 | PBMC |
| GSE325620 | Danio rerio | Apr 02 2026 | PRJNA1441355 | retina |
| GSE181713 | Oryza sativa | Aug 12 2021 | PRJNA753085 | Panicle |
| GSE316127 | Homo sapiens | Jan 16 2026 | PRJNA1401584 | |
| GSE296350 | Mus musculus | Feb 10 2026 | PRJNA1258971 | |
| GSE305018 | Homo sapiens | Jan 14 2026 | PRJNA1303711 | cell line |
| GSE301704 | Mus musculus | Jan 07 2026 | | Mandible |
| GSE270442 | Danio rerio | Jul 01 2024 | PRJNA1126644 | larvae |
| GSE300765 | Homo sapiens | Jan 04 2026 | PRJNA1282076 | Brain |
| GSE222430 | Mus musculus | Jan 08 2026 | PRJNA922081 | hippocampus |
| GSE277287 | Mus musculus | Jan 13 2026 | PRJNA1161651 | |
| GSE309032 | erichia coli str. K-12 substr. MG1 | Jan 07 2026 | PRJNA1333829 | whole cells |
| GSE293132 | Mus musculus | Sep 10 2025 | PRJNA1242865 | retina |
| GSE223304 | Homo sapiens | Jan 18 2026 | PRJNA925550 | Laryngeal tissue |
| GSE309994 | Homo sapiens | Jan 26 2026 | PRJNA1364975 | |
| GSE305252 | Homo sapiens | Nov 04 2025 | PRJNA1304921 | umbilical vein |
| GSE316672 | Homo sapiens | Jan 22 2026 | PRJNA1403559 | lung |
| GSE287482 | Mus musculus | Jan 25 2025 | PRJNA1213080 | organoids |
| GSE315958 | Homo sapiens | Feb 25 2026 | PRJNA1400017 | |
| GSE316881 | Homo sapiens;Mus musculus | Feb 26 2026 | PRJNA1405471 | Brain |
| GSE189770 | Homo sapiens | Dec 01 2021 | PRJNA784443 | etinal pigment epithelium |
| GSE294933 | Mus musculus | Jan 06 2026 | PRJNA1252062 | Brain |
| GSE293026 | Mus musculus | Jan 14 2026 | PRJNA1242371 | cortex |
| GSE317052 | Mus musculus | Jan 30 2026 | PRJNA1405710 | heart |
| GSE307925 | Homo sapiens | Sep 19 2025 | PRJNA1327830 | |
| GSE135167 | Mus musculus | Feb 28 2020 | PRJNA557653 | |
| GSE295871 | Procambarus virginalis | Jan 29 2026 | PRJNA1256520 | Hemolymph |
| GSE223738 | Homo sapiens | Jan 30 2024 | PRJNA928081 | |
| GSE288330 | us hybridus;Amaranthus hypoch | Jan 29 2026 | PRJNA1216948 | Leaves |
| GSE212187 | Mus musculus | Nov 30 2022 | PRJNA874433 | bulk retina |
| GSE317730 | Homo sapiens | Feb 27 2026 | PRJNA1415119 | |
| GSE294181 | Homo sapiens | Feb 23 2026 | PRJNA1248516 | |
| GSE289433 | Homo sapiens | Jan 30 2026 | PRJNA1222899 | Liver |
| GSE166174 | Homo sapiens | Jul 26 2021 | PRJNA699543 | Retina |
| GSE318830 | Homo sapiens | Feb 13 2026 | PRJNA1420689 | lung |
| GSE316004 | anis lupus familiaris;Mus muscul | Jan 28 2026 | PRJNA1400327 | |
| GSE301338 | Mus musculus | Jan 01 2026 | PRJNA1284816 | peritoneal macrophage |
| GSE310853 | Homo sapiens | Jan 30 2026 | PRJNA1367174 | |
| GSE239605 | Homo sapiens | Aug 02 2023 | PRJNA1000495 | Abdominal Subcut. |
| GSE304638 | Homo sapiens | Jan 01 2026 | PRJNA1301964 | |
| GSE316995 | Mus musculus | Jan 25 2026 | PRJNA1405651 | |
| GSE304327 | Homo sapiens;synthetic construc | Jan 06 2026 | PRJNA1300393 | Mammary |
| GSE236905 | Gallus gallus | Dec 27 2023 | PRJNA993020 | Eye |
| GSE298483 | Homo sapiens | Jan 21 2026 | PRJNA1269528 | kidney tumor |
| GSE273836 | Mus musculus | Jan 01 2026 | PRJNA1143438 | Fully-grown GV oocyte |
| GSE271262 | Mus musculus | Dec 31 2024 | PRJNA1130615 | retina+choroid |
| GSE316279 | Homo sapiens | Jan 18 2026 | PRJNA1402154 | retina |
| GSE289679 | Homo sapiens | Feb 18 2026 | PRJNA1223938 | |
| GSE254445 | Homo sapiens | Feb 11 2026 | PRJNA1070592 | kidney |
| GSE290089 | apiens;Macaca fascicularis;Mus m | Jan 19 2026 | PRJNA1226179 | Ovary |
| GSE304504 | Macaca mulatta | Mar 30 2026 | PRJNA1301394 | Posterior eyecup |
| GSE317124 | Rattus norvegicus | Feb 08 2026 | | liver |
| GSE293150 | Caenorhabditis elegans | Feb 19 2026 | PRJNA1242887 | bulk |
| GSE263000 | Homo sapiens | Feb 15 2026 | PRJNA1095431 | Cell line |

| | | | | |
|---|---|---|---|---|
| GSE269391 | Homo sapiens | Feb 11 2026 | PRJNA1121367 | Blood |
| GSE315583 | Mus musculus | Jan 10 2026 | PRJNA1398284 | guinal white adipose tissue |
| GSE267561 | Homo sapiens | Jan 24 2026 | PRJNA1111855 | ICM |
| GSE313445 | Mus musculus | Jan 06 2026 | PRJNA1379527 | ascites |
| GSE294966 | Homo sapiens;Mus musculus | Feb 28 2026 | PRJNA1252116 | |
| GSE202017 | Homo sapiens | Jul 03 2022 | PRJNA834053 | |
| GSE281526 | Homo sapiens | Oct 14 2025 | | fovea centralis |
| GSE284026 | Mus musculus | Jan 23 2026 | PRJNA1196866 | |
| GSE316172 | Mus musculus | Jan 15 2026 | | brain |
| GSE261490 | Mus musculus | May 01 2025 | PRJNA1087306 | retina |
| GSE279422 | Homo sapiens | Jan 24 2026 | PRJNA1172682 | Skin |
| GSE309067 | Zea mays | Jan 26 2026 | PRJNA1333892 | Leaves |
| GSE311944 | Homo sapiens;Mus musculus | Jan 13 2026 | PRJNA1371974 | |
| GSE288513 | Mus musculus | Feb 13 2026 | PRJNA1217895 | NO |
| GSE315188 | Komagataella pastoris | Jan 06 2026 | PRJNA1395103 | |
| GSE227724 | Gallus gallus | Dec 20 2023 | PRJNA946718 | choroid |
| GSE313551 | Mus musculus | Feb 18 2026 | PRJNA1380306 | pancreas islet |
| GSE285606 | Mus musculus | Feb 19 2026 | PRJNA1204959 | |
| GSE301881 | Homo sapiens | Jan 24 2026 | PRJNA1287688 | human 4C embryo |
| GSE284378 | Arabidopsis thaliana | Feb 11 2026 | PRJNA1198909 | 10-day-old seedlings |
| GSE319833 | Caenorhabditis elegans | Feb 23 2026 | PRJNA1425278 | rmis - srf-3i expression domain |
| GSE229872 | Mus musculus | May 31 2023 | PRJNA956504 | retina |
| GSE305324 | Homo sapiens | Feb 13 2026 | PRJNA1305312 | |
| GSE312018 | Mus musculus | Jan 18 2026 | PRJNA1372058 | kidney |
| GSE223510 | Sus scrofa | Jan 19 2026 | PRJNA926619 | Dental epithelium |
| GSE229681 | Homo sapiens | Nov 03 2023 | PRJNA955280 | Retinal organoid |
| GSE306080 | Homo sapiens | Feb 25 2026 | PRJNA1309079 | |
| GSE242265 | Mus musculus | Feb 20 2026 | PRJNA1012733 | |
| GSE250146 | Mus musculus | Jan 01 2026 | PRJNA1052104 | Tumor |
| GSE284132 | Petromyzon marinus | Aug 18 2025 | PRJNA1197420 | retina |
| GSE308112 | Homo sapiens | Feb 26 2026 | PRJNA1328753 | lung |
| GSE319475 | Mus musculus | Feb 13 2026 | PRJNA1423509 | ated by FACS from medial habenula brain tissue |
| GSE254856 | Rattus norvegicus | Feb 06 2024 | PRJNA1072090 | Retina |
| GSE188698 | Homo sapiens | Jun 08 2022 | PRJNA779960 | |
| GSE296093 | Mus musculus | Feb 15 2026 | PRJNA1257771 | Spleen |
| GSE252763 | Homo sapiens | Jan 08 2026 | PRJNA1062406 | nasal epithelium |
| GSE208745 | Homo sapiens | Jan 21 2026 | PRJNA860936 | |
| GSE307263 | Homo sapiens | Feb 26 2026 | PRJNA1315084 | umbilical cord blood |
| GSE318414 | Homo sapiens | Feb 06 2026 | PRJNA1418692 | Skin |
| GSE237215 | emlineatus;Mustela putorius fur | Oct 01 2023 | PRJNA994164 | Retina |
| GSE283708 | Mus musculus | Feb 25 2026 | PRJNA1195074 | ochlear Sensory Epithelium |
| GSE295628 | Mus musculus | Jan 05 2026 | | Lung |
| GSE297954 | Homo sapiens | Jul 25 2025 | PRJNA1267027 | Retinal organoids |
| GSE305800 | Mus musculus | Jan 30 2026 | PRJNA1307634 | Mammary |
| GSE263689 | Mus musculus | Jan 23 2026 | PRJNA1098736 | primary lung tumor |
| GSE302754 | Mus musculus | Jan 12 2026 | | skin |
| GSE303735 | Homo sapiens | Jan 01 2026 | | PBMC |
| GSE291661 | Mus musculus | Mar 02 2026 | PRJNA1234617 | retina explant |
| GSE284698 | Rattus norvegicus | Jan 11 2026 | | Tumor |
| GSE240315 | Mus musculus | Aug 28 2023 | PRJNA1003142 | |
| GSE242941 | Homo sapiens | Jan 14 2026 | | BCC peri-tumor |
| GSE272311 | Rattus norvegicus | Jul 16 2024 | PRJNA1136409 | Retina |
| GSE304861 | Mus musculus | Jan 26 2026 | PRJNA1303081 | Spleen |
| GSE311748 | Homo sapiens | Jan 12 2026 | PRJNA1370872 | Kidney |
| GSE297877 | Homo sapiens | May 22 2025 | PRJNA1266668 | Human Tenon's capsule |
| GSE313803 | Mus musculus | Jan 01 2026 | PRJNA1381609 | cortex |
| GSE161310 | Mus musculus | Sep 30 2021 | PRJNA676207 | Retinal endothelial cells |
| GSE252677 | Homo sapiens | Jan 01 2026 | PRJNA1061784 | |
| GSE287891 | Drosophila melanogaster | Jan 23 2026 | PRJNA1214871 | ovaries |
| GSE199576 | Homo sapiens | Nov 17 2022 | PRJNA820745 | ived mesodermal cell aggregates |
| GSE306609 | Homo sapiens | Feb 26 2026 | PRJNA1311463 | Kidney PDX |
| GSE189590 | Mus musculus | Dec 22 2022 | PRJNA783798 | |
| GSE310729 | Homo sapiens | Jan 05 2026 | PRJNA1366996 | Peripheral blood |

| | | | | |
|---|---|---|---|---|
| GSE264312 | Danio rerio | Oct 21 2024 | PRJNA1101942 | eye |
| GSE312862 | Mus musculus | Mar 19 2026 | PRJNA1377522 | Retina |
| GSE315229 | Homo sapiens | Jan 04 2026 | | liver |
| GSE319556 | Mus musculus | Feb 20 2026 | PRJNA1423723 | leptomeninges |
| GSE286449 | Mus musculus | Jan 01 2026 | PRJNA1209388 | tongue |
| GSE291775 | Mus musculus | Sep 26 2025 | PRJNA1235223 | Retina |
| GSE314993 | Rattus norvegicus | Jan 06 2026 | PRJNA1394215 | heart |
| GSE279697 | Mus musculus | Jan 29 2026 | PRJNA1174106 | Epidermis of back skin |
| GSE213212 | Homo sapiens | Feb 23 2026 | PRJNA879757 | |
| GSE302640 | Homo sapiens | Jan 12 2026 | PRJNA1291633 | |
| GSE318537 | Homo sapiens | Feb 13 2026 | PRJNA1419191 | |
| GSE248537 | Mus musculus | Dec 26 2023 | PRJNA1044446 | nriched RGC mouse retina |
| GSE315063 | Mus musculus | Feb 18 2026 | PRJNA1394944 | cell line |
| GSE262581 | Homo sapiens | Jan 01 2026 | PRJNA1092592 | cell line |
| GSE272139 | Canis lupus familiaris | Jul 01 2025 | PRJNA1135184 | Neuroretina |
| GSE313975 | Fundulus heteroclitus | Jan 07 2026 | PRJNA1387461 | embryo |
| GSE318476 | Mus musculus | Feb 11 2026 | PRJNA1418808 | Glomeruli |
| GSE303002 | Mus musculus | Feb 09 2026 | PRJNA1293331 | testis |
| GSE261519 | Mus musculus | Feb 08 2026 | PRJNA1087370 | Spinal cord |
| GSE306502 | Mus musculus | Feb 11 2026 | PRJNA1310994 | BMDM-M0 |
| GSE319375 | Mus musculus | Feb 12 2026 | PRJNA1423170 | |
| GSE249375 | Homo sapiens | Jan 26 2024 | PRJNA1048906 | |
| GSE313299 | Mus musculus | Jan 05 2026 | PRJNA1378900 | lymph node |
| GSE295128 | Homo sapiens | Feb 13 2026 | PRJNA1253599 | |
| GSE316799 | Homo sapiens | Feb 23 2026 | PRJNA1404225 | Lung |
| GSE192903 | Homo sapiens | Jan 05 2022 | PRJNA793703 | |
| GSE169550 | Mus musculus | Jan 11 2022 | PRJNA716996 | |
| GSE236216 | Homo sapiens;Mus musculus | Jan 01 2026 | PRJNA989273 | bone marrow |
| GSE313786 | Mus musculus | Jan 31 2026 | PRJNA1381594 | CD71+ EB-derived erythroid cells, originally derived from ESC |
| GSE288814 | Citrullus lanatus | Feb 09 2026 | | leaves |
| GSE236221 | Mus musculus | Jul 15 2023 | PRJNA989276 | RPE/choroid |
| GSE269707 | Mus musculus | Jan 08 2026 | PRJNA1123274 | nouse sciatic nerve tissue |
| GSE287723 | Homo sapiens | Jan 06 2026 | PRJNA1214335 | colon |
| GSE222955 | Mus musculus | Feb 15 2023 | PRJNA924280 | retina |
| GSE313942 | Mus musculus | Jan 28 2026 | PRJNA1387409 | rived from fetal thymocytes |
| GSE131877 | Homo sapiens | May 28 2020 | PRJNA545219 | |
| GSE318441 | Homo sapiens | Feb 05 2026 | PRJNA1418735 | |
| GSE294461 | Mus musculus | Jan 22 2026 | PRJNA1249495 | 6/F10-BrM brain metastasis |
| GSE176472 | Mus musculus | Apr 06 2022 | PRJNA736401 | |
| GSE307729 | Homo sapiens | Jan 14 2026 | PRJNA1327222 | |
| GSE228276 | hlorocebus sabaeus;Homo sapie | Aug 30 2023 | PRJNA949182 | crovascular endothelial cells in cell culture |
| GSE298077 | Pseudomonas aeruginosa | Jan 12 2026 | PRJNA1268317 | |
| GSE201319 | Drosophila melanogaster | Feb 11 2026 | PRJNA830867 | testes |
| GSE288091 | Mus musculus | Jan 25 2026 | PRJNA1216177 | spleen |
| GSE317491 | Mus musculus | Jan 25 2026 | PRJNA1413915 | whole kidney |
| GSE306374 | Homo sapiens | Jan 26 2026 | PRJNA1310540 | terminal ileum, colon |
| GSE80687 | Arabidopsis thaliana | Jan 31 2026 | PRJNA319671 | seedlings |
| GSE315754 | Mus musculus | Jan 26 2026 | PRJNA1398895 | embryonic facial tissue |
| GSE117546 | Mus musculus | Jan 12 2026 | PRJNA482435 | ssue from pT3-vector plasmid injected mice |
| GSE315529 | Homo sapiens | Jan 31 2026 | PRJNA1398214 | |
| GSE303409 | Homo sapiens | Feb 25 2026 | PRJNA1295363 | |
| GSE315036 | Mus musculus | Feb 20 2026 | PRJNA1394901 | |
| GSE273231 | nearum;Schizosaccharomyces ja | Feb 24 2026 | PRJNA1140632 | |
| GSE279197 | Homo sapiens | Feb 25 2026 | PRJNA1171355 | |
| GSE315990 | Polypterus senegalus | Jan 15 2026 | PRJNA1400068 | pectoral fin |
| GSE314702 | Homo sapiens | Jan 30 2026 | | |
| GSE290014 | Homo sapiens | Feb 18 2026 | PRJNA1225736 | |
| GSE199571 | Homo sapiens | Nov 15 2022 | PRJNA820750 | rived mesodermal cell aggregates |
| GSE306311 | Gallus gallus | Jan 01 2026 | PRJNA1310447 | hypothalamus |
| GSE290559 | Homo sapiens | Sep 24 2025 | PRJNA1228620 | retinal endothelial cells |
| GSE315731 | Mus musculus | Jan 07 2026 | PRJNA1398864 | Liver |
| GSE311532 | Danio rerio | Jan 07 2026 | | whole body |
| GSE277735 | Mus musculus | Jan 04 2026 | PRJNA1163690 | liver |

| | | | | |
|---|---|---|---|---|
| GSE272235 | Mus musculus | Jan 01 2026 | PRJNA1135917 | Syngenic Tumor |
| GSE286166 | Mus musculus | Jan 01 2026 | PRJNA1208038 | Colon |
| GSE306473 | Homo sapiens | Jan 13 2026 | PRJNA1310946 | Lymph node |
| GSE315053 | Mus musculus | Jan 05 2026 | PRJNA1394933 | cutaneous pancreatic cancer |
| GSE312696 | Mus musculus | Feb 13 2026 | PRJNA1375724 | oVAT |
| GSE151947 | Gallus gallus | Jul 01 2021 | PRJNA637870 | |
| GSE316495 | Homo sapiens | Feb 27 2026 | PRJNA1403014 | |
| GSE269123 | Homo sapiens | Feb 13 2026 | PRJNA1120354 | thymus |
| GSE283333 | Mus musculus | Jan 31 2026 | PRJNA1193207 | Lung |
| GSE307658 | Mus musculus | Feb 21 2026 | PRJNA1322470 | Spleen |
| GSE284294 | Mus musculus | Feb 05 2026 | PRJNA1198356 | proximal colon |
| GSE315219 | Homo sapiens | Feb 09 2026 | PRJNA1395138 | |
| GSE186331 | Homo sapiens | Mar 08 2023 | PRJNA773409 | |
| GSE282352 | Homo sapiens | Feb 18 2026 | PRJNA1188173 | |
| GSE201984 | Homo sapiens | Jul 15 2022 | PRJNA833892 | |
| GSE307264 | Homo sapiens | Feb 26 2026 | PRJNA1315034 | iPSC-derived cells |
| GSE237212 | Ictidomys tridecemlineatus | Oct 01 2023 | PRJNA994176 | Retina |
| GSE314126 | Plasmodium falciparum | Jan 23 2026 | PRJNA1390134 | e grown in human red blood cells |
| GSE296905 | Synechocystis sp. PCC 6803 | Jan 26 2026 | PRJNA1261913 | total cell |
| GSE319472 | Homo sapiens | Feb 27 2026 | PRJNA1423506 | air liquid interface culture |
| GSE318371 | Homo sapiens | Feb 07 2026 | | blood |
| GSE201740 | Homo sapiens | Feb 08 2023 | PRJNA832801 | |
| GSE318346 | Homo sapiens | Feb 05 2026 | PRJNA1418616 | Nasal epithelium |
| GSE301879 | Homo sapiens | Jan 24 2026 | PRJNA1287692 | human 4C embryo |
| GSE151697 | Gallus gallus | Apr 19 2021 | PRJNA636978 | |
| GSE240312 | Mus musculus | Nov 02 2023 | PRJNA1003125 | Retina |
| GSE281680 | Bos taurus | Jan 08 2026 | PRJNA1185221 | |
| GSE318659 | Homo sapiens | Feb 11 2026 | | Cervical |
| GSE319445 | Homo sapiens | Feb 13 2026 | PRJNA1423290 | |
| GSE273377 | Homo sapiens | Jan 26 2026 | PRJNA1141508 | lung adenocarcinoma |
| GSE310470 | Homo sapiens | Jan 26 2026 | PRJNA1366401 | y human Donor lung fibroblasts |
| GSE156730 | Mus musculus | Jul 25 2023 | PRJNA658914 | retina |
| GSE273668 | Homo sapiens | May 06 2025 | PRJNA1142597 | |
| GSE311373 | Mus musculus | Feb 24 2026 | PRJNA1369365 | uterine tumor |
| GSE261054 | Homo sapiens | Feb 10 2026 | PRJNA1084905 | cell line |
| GSE319266 | Homo sapiens | Feb 13 2026 | PRJNA1422730 | |
| GSE293380 | Mus musculus | May 09 2025 | PRJNA1244319 | Retina |
| GSE319779 | Mus musculus;synthetic construc | Feb 17 2026 | PRJNA1425080 | marrow |
| GSE161417 | Homo sapiens | Mar 26 2023 | PRJNA678242 | |
| GSE295524 | Mus musculus | Feb 26 2026 | PRJNA1255119 | Lung |
| GSE197481 | Mus musculus | Nov 15 2022 | PRJNA810493 | |
| GSE264377 | Danio rerio | Mar 01 2026 | PRJNA1102288 | larvae |
| GSE290024 | Homo sapiens | Feb 24 2025 | PRJNA1225751 | Retinal organoid |
| GSE288096 | Mus musculus | Mar 01 2025 | PRJNA1216196 | Whole Retina |
| GSE317496 | Homo sapiens | Jan 31 2026 | PRJNA1414091 | |
| GSE222871 | Mus musculus | Jan 13 2026 | PRJNA923689 | Brain |
| GSE274391 | Xenopus laevis | Jan 01 2026 | PRJNA1146329 | animal cap |
| GSE315753 | Homo sapiens | Jan 07 2026 | PRJNA1398886 | stem cell-derived islet |
| GSE209662 | Homo sapiens | Jul 28 2022 | PRJNA862064 | Peripheral blood |
| GSE315764 | Homo sapiens | Jan 28 2026 | PRJNA1398925 | Blood cells |
| GSE288930 | Mus musculus | Feb 11 2025 | PRJNA1220049 | Retina |
| GSE266085 | Homo sapiens | Sep 02 2024 | PRJNA1105870 | |
| GSE319533 | Gallus gallus | Feb 18 2026 | PRJNA1423680 | embryo liver |
| GSE296844 | Homo sapiens | Jan 16 2026 | PRJNA1261803 | |
| GSE312830 | Homo sapiens | Feb 26 2026 | PRJNA1377490 | |
| GSE314067 | Homo sapiens | Feb 20 2026 | PRJNA1389921 | |
| GSE287152 | Mus musculus | Jan 31 2026 | PRJNA1211501 | |
| GSE318552 | Homo sapiens | Feb 09 2026 | PRJNA1419214 | Brain |
| GSE311205 | Homo sapiens | Jan 05 2026 | PRJNA1368805 | pulmonary blastoma lung lesion |
| GSE315519 | Mus musculus | Feb 08 2026 | | Spleen |
| GSE315997 | Homo sapiens | Feb 26 2026 | PRJNA1400085 | |
| GSE274285 | Mus musculus | Sep 01 2025 | PRJNA1145869 | |
| GSE273440 | Mus musculus | Feb 07 2026 | PRJNA1141927 | pineal gland |

| | | | | |
|---|---|---|---|---|
| GSE306267 | Homo sapiens | Feb 26 2026 | PRJNA1310367 | Ovary PDX |
| GSE302064 | Mus musculus | Feb 25 2026 | PRJNA1288437 | |
| GSE314144 | Mus musculus | Jan 01 2026 | PRJNA1390156 | Liver |
| GSE288813 | Citrullus lanatus | Feb 09 2026 | | leaves |
| GSE303005 | Escherichia coli | Feb 05 2026 | PRJNA1293343 | |
| GSE315125 | Homo sapiens | Jan 01 2026 | PRJNA1395025 | ovary |
| GSE298319 | Mus musculus | Feb 06 2026 | PRJNA1268951 | Bone Marrow |
| GSE310425 | Homo sapiens | Feb 23 2026 | PRJNA1365918 | |
| GSE299894 | Mus musculus | Feb 10 2026 | PRJNA1277270 | cortex and hippocampus |
| GSE295352 | Mus musculus | Jan 26 2026 | PRJNA1254258 | spleen |
| GSE319345 | Mus musculus | Feb 12 2026 | PRJNA1423130 | liver |
| GSE302731 | Monodelphis domestica | Jan 31 2026 | PRJNA1291816 | omphalopleure |
| GSE315883 | Homo sapiens | Jan 13 2026 | PRJNA1399499 | |
| GSE290665 | Mus musculus | Feb 23 2026 | PRJNA1229496 | skeletal muscle |
| GSE214928 | Homo sapiens | Oct 21 2022 | PRJNA887751 | |
| GSE287046 | Mus musculus | Jan 18 2026 | PRJNA1211047 | Ileum |
| GSE318446 | Homo sapiens | Feb 05 2026 | PRJNA1418740 | Placenta |
| GSE299132 | Mus musculus | Jan 28 2026 | PRJNA1272631 | hear |
| GSE213401 | Astyanax mexicanus | Feb 28 2026 | PRJNA880814 | Skeletal Muscle |
| GSE266191 | Homo sapiens | Sep 01 2025 | PRJNA1106186 | retinal organoid |
| GSE287724 | Homo sapiens | Jan 06 2026 | PRJNA1214338 | colon |
| GSE294186 | Homo sapiens | Jan 01 2026 | PRJNA1248528 | |
| GSE151477 | Rattus norvegicus | Sep 10 2020 | PRJNA635927 | |
| GSE235462 | Homo sapiens | Dec 04 2023 | PRJNA986106 | |
| GSE166173 | Homo sapiens | Jul 26 2021 | PRJNA699542 | Retina |
| GSE300114 | Mus musculus | Feb 06 2026 | PRJNA1279044 | ovasculature_Tg350_9 months _female |
| GSE291199 | Mus musculus | Feb 11 2026 | PRJNA1232195 | Breast tumor |
| GSE311802 | Mus musculus | Feb 26 2026 | PRJNA1370926 | neocortex |
| GSE307922 | Homo sapiens | Feb 26 2026 | PRJNA1327816 | |
| GSE271252 | Mus musculus | Nov 20 2024 | PRJNA1130532 | Retina |
| GSE102464 | Mus musculus | Jan 01 2020 | PRJNA397780 | |
| GSE298484 | Mus musculus | Jan 01 2026 | PRJNA1269532 | kidney |
| GSE263146 | Mus musculus | Feb 11 2026 | PRJNA1095976 | Mus musculus |
| GSE260580 | Homo sapiens | Jan 07 2026 | PRJNA1082270 | |
| GSE317062 | Rattus norvegicus | Feb 18 2026 | PRJNA1405731 | ileum |
| GSE296998 | Mus musculus | Jan 13 2026 | PRJNA1262341 | etastatic colorectal cancer |
| GSE284157 | Mus musculus | Jan 30 2026 | PRJNA1197474 | Heart |
| GSE310854 | Homo sapiens | Jan 30 2026 | PRJNA1367175 | |
| GSE306974 | Homo sapiens | Jan 20 2026 | | Small Intestine |
| GSE270204 | Homo sapiens | Jan 01 2026 | PRJNA1125407 | |
| GSE288300 | Mus musculus | Feb 16 2026 | PRJNA1216891 | |
| GSE236902 | Gallus gallus | Dec 27 2023 | PRJNA993024 | Eye |
| GSE289242 | Homo sapiens | Jan 28 2026 | PRJNA1221963 | |
| GSE159255 | Mus musculus | Oct 08 2023 | PRJNA668145 | |
| GSE184069 | Drosophila melanogaster | Dec 31 2021 | PRJNA763088 | Retina |
| GSE308054 | Homo sapiens | Sep 20 2025 | PRJNA1328664 | |
| GSE316120 | Polypterus senegalus | Jan 15 2026 | PRJNA1400634 | pectoral fin |
| GSE315968 | Mus musculus | Jan 13 2026 | PRJNA1400033 | |
| GSE144914 | Homo sapiens | Feb 08 2020 | PRJNA605325 | |
| GSE300991 | Mus musculus | Feb 24 2026 | PRJNA1282966 | breast tumor |
| GSE288223 | Homo sapiens | Feb 24 2026 | PRJNA1216591 | breast tumor |
| GSE193437 | Homo sapiens | Jun 14 2022 | PRJNA796230 | |
| GSE287485 | Mus musculus | Jan 25 2025 | PRJNA1213086 | organoids |
| GSE279158 | Drosophila melanogaster | Jan 27 2026 | PRJNA1170998 | Brain |
| GSE308063 | Homo sapiens | Jan 13 2026 | PRJNA1328679 | Brain |
| GSE313899 | Mus musculus | Jan 01 2026 | PRJNA1387222 | |
| GSE286386 | Mus musculus | Jan 01 2026 | PRJNA1209042 | |
| GSE310940 | Homo sapiens | Jan 14 2026 | PRJNA1367582 | Placenta |
| GSE160489 | Gallus gallus | Oct 31 2020 | PRJNA673388 | Thymus |
| GSE317176 | Macaca fascicularis | Jan 24 2026 | | hippocampus |
| GSE202238 | Danio rerio | May 07 2022 | PRJNA835105 | whole embryo |
| GSE254049 | Homo sapiens | Jan 19 2026 | PRJNA1068251 | |
| GSE155035 | Mus musculus | Aug 13 2023 | PRJNA648315 | |

| | | | | |
|---|---|---|---|---|
| GSE317469 | Homo sapiens | Jan 28 2026 | PRJNA1413175 | breast cancer |
| GSE199962 | Mus musculus | Aug 05 2022 | PRJNA822371 | Cortex |
| GSE316734 | Homo sapiens | Feb 19 2026 | PRJNA1404100 | |
| GSE138149 | Camellia sinensis | Dec 31 2020 | PRJNA574929 | root |
| GSE242537 | Homo sapiens | Dec 01 2023 | PRJNA1013821 | |
| GSE304899 | Mus musculus | Feb 19 2026 | | spleen |
| GSE289456 | Homo sapiens | Feb 10 2026 | PRJNA1222997 | bone |
| GSE306921 | Mus musculus | Feb 12 2026 | PRJNA1313926 | ovary |
| GSE317755 | Ambystoma mexicanum | Jan 28 2026 | | Olfactory Bulb |
| GSE304375 | Homo sapiens | Jan 12 2026 | PRJNA1301130 | |
| GSE292984 | Homo sapiens | Jan 08 2026 | PRJNA1242294 | |
| GSE203630 | Gallus gallus | May 19 2025 | PRJNA841645 | |
| GSE253587 | Homo sapiens;Rattus norvegicus | Feb 26 2026 | | Breast invasive carcinoma |
| GSE316061 | Leisingera sp. ANG-M7 | Jan 13 2026 | PRJNA1400543 | |
| GSE299198 | Arabidopsis thaliana | Jan 22 2026 | PRJNA1273061 | whole seedlings |
| GSE300141 | Mus musculus | Feb 12 2026 | PRJNA1279099 | heart |
| GSE297515 | Mus musculus | Jan 13 2026 | PRJNA1265126 | Muscle |
| GSE309309 | Homo sapiens | Feb 12 2026 | PRJNA1335126 | Peripheral blood |
| GSE281435 | Danio rerio | Jan 06 2026 | PRJNA1183855 | 28 hpf embryos |
| GSE308368 | Mus musculus | Feb 18 2026 | PRJNA1330294 | Cerebral tissue |
| GSE281648 | Homo sapiens | Sep 17 2025 | PRJNA1185156 | |
| GSE218039 | Homo sapiens | Jan 01 2023 | PRJNA901967 | Placenta |
| GSE151501 | Mus musculus | Jul 27 2021 | PRJNA636090 | Liver |
| GSE143622 | Mus musculus | Jan 04 2026 | PRJNA601130 | spleen and lymph node |
| GSE270345 | Canis lupus familiaris | Feb 19 2026 | PRJNA1126235 | mammary |
| GSE316142 | Danio rerio | Jan 12 2026 | PRJNA1401708 | spinal cord |
| GSE311787 | Homo sapiens | Mar 03 2026 | PRJNA1370914 | |
| GSE317123 | Rattus norvegicus | Feb 08 2026 | | liver |
| GSE301003 | Plasmodium vivax | Feb 12 2026 | PRJNA1283052 | Enriched blood sample |
| GSE223356 | Homo sapiens | Jan 20 2026 | PRJNA925864 | |
| GSE144785 | Homo sapiens | Aug 31 2022 | PRJNA604911 | |
| GSE179438 | Homo sapiens | Jul 08 2021 | PRJNA743802 | |
| GSE289217 | Homo sapiens | Feb 09 2026 | PRJNA1221918 | |
| GSE139276 | Rattus norvegicus | Dec 16 2021 | PRJNA579063 | |
| GSE312414 | Mus musculus | Jan 26 2026 | PRJNA1373272 | BM culture |
| GSE311943 | Homo sapiens | Feb 07 2026 | PRJNA1371946 | Lung tumor |
| GSE305555 | Homo sapiens | Jan 31 2026 | PRJNA1306494 | |
| GSE292101 | Homo sapiens | Jan 09 2026 | PRJNA1236960 | |
| GSE287869 | Homo sapiens | Jan 17 2026 | PRJNA1214835 | |
| GSE317676 | Oryctolagus cuniculus | Jan 31 2026 | PRJNA1414631 | peripheral blood |
| GSE301756 | Homo sapiens | Feb 08 2026 | PRJNA1287528 | |
| GSE228900 | Homo sapiens | Dec 12 2023 | PRJNA952113 | r blood-retinal barrier (iBRB) |
| GSE280480 | Homo sapiens | Jan 09 2026 | PRJNA1178681 | ALCL tumors |
| GSE279287 | Homo sapiens | Feb 10 2026 | PRJNA1171803 | |
| GSE311112 | Homo sapiens | Jan 27 2026 | PRJNA1368690 | peripheral blood |
| GSE203499 | Homo sapiens | Jun 23 2022 | PRJNA841015 | |
| GSE268200 | Mus musculus | Feb 09 2026 | PRJNA1115115 | liver |
| GSE273674 | Homo sapiens | May 06 2025 | PRJNA1142593 | |
| GSE245473 | Homo sapiens | Oct 01 2025 | PRJNA1028561 | endometrium |
| GSE310173 | Homo sapiens | Jan 20 2026 | PRJNA1365224 | lung cancer |
| GSE299453 | Drosophila melanogaster | Oct 22 2025 | PRJNA1274462 | thorax |
| GSE306053 | Mus musculus | Feb 11 2026 | PRJNA1309019 | |
| GSE180813 | Mus musculus | Dec 03 2022 | PRJNA749618 | Retina |
| GSE290304 | Homo sapiens | Nov 20 2025 | PRJNA1227441 | retinal organoids |
| GSE294418 | Mus musculus | Feb 06 2026 | PRJNA1249419 | sperm |
| GSE318881 | Homo sapiens | Feb 13 2026 | PRJNA1420767 | Peripheral blood |
| GSE275785 | Mus musculus | Feb 20 2026 | PRJNA1152977 | Tumor |
| GSE297978 | Mus musculus | Jan 01 2026 | PRJNA1267062 | cells |
| GSE315444 | Anser cygnoides | Feb 28 2026 | PRJNA1397162 | thymus |
| GSE314658 | Homo sapiens | Feb 11 2026 | PRJNA1392416 | Placenta |
| GSE310144 | Homo sapiens | Jan 20 2026 | PRJNA1365196 | lung cancer |
| GSE288381 | Mus musculus | Sep 18 2025 | PRJNA1217330 | retina |
| GSE316913 | Arabidopsis thaliana | Jan 26 2026 | PRJNA1405532 | leaf |

| | | | | |
|---|---|---|---|---|
| GSE309806 | Homo sapiens | Apr 06 2026 | PRJNA1363211 | cell line |
| GSE249171 | Mus musculus | Feb 18 2026 | PRJNA1047592 | ureter |
| GSE302505 | scherichia phage T4;Homo sapie | Feb 17 2026 | PRJNA1291096 | |
| GSE237073 | Mus musculus | Oct 01 2023 | PRJNA993672 | optic nerve head |
| GSE315326 | Homo sapiens | Jan 23 2026 | PRJNA1395637 | |
| GSE269749 | Homo sapiens | Jan 08 2026 | PRJNA1123612 | Bone |
| GSE294652 | Homo sapiens | Feb 12 2026 | | human CAR-T cells |
| GSE303158 | Homo sapiens | Feb 11 2026 | PRJNA1294256 | |
| GSE300910 | Mus musculus | Feb 09 2026 | PRJNA1282658 | Lung |
| GSE285905 | Homo sapiens | Jan 01 2026 | PRJNA1206919 | |
| GSE311997 | Mus musculus | Jan 30 2026 | PRJNA1372035 | testis |
| GSE284288 | Ovis aries | Jan 14 2026 | PRJNA1198348 | rumen |
| GSE314019 | Rattus norvegicus | Feb 25 2026 | PRJNA1389474 | Breast tumor |
| GSE268441 | Homo sapiens | Jan 16 2026 | PRJNA1117391 | PDA Cell Line |
| GSE304959 | Mus musculus | Sep 18 2025 | PRJNA1303595 | retina |
| GSE264309 | Danio rerio | Oct 21 2024 | PRJNA1101945 | eye |
| GSE247866 | Mus musculus | Jan 01 2026 | PRJNA1040839 | Submandibular gland |
| GSE287679 | Homo sapiens | Feb 09 2026 | PRJNA1214239 | embryonic kidney |
| GSE318279 | Mus musculus | Feb 05 2026 | PRJNA1418504 | |
| GSE317695 | Homo sapiens | Jan 31 2026 | PRJNA1414665 | |
| GSE268323 | Mus musculus | Oct 23 2024 | PRJNA1116214 | prachiasmatic nucleus (SCN) |
| GSE201402 | Mus musculus | May 02 2022 | PRJNA831452 | retina |
| GSE315550 | Homo sapiens | Jan 07 2026 | PRJNA1398251 | Lymph node |
| GSE318004 | feces metagenome | Feb 16 2026 | PRJNA1416305 | Mus musculus stool |
| GSE176528 | Mus musculus | Sep 12 2021 | PRJNA736614 | retina |
| GSE307673 | Mus musculus | Feb 25 2026 | PRJNA1322495 | iWAT |
| GSE313496 | Homo sapiens | Jan 16 2026 | PRJNA1379654 | whole blood |
| GSE275691 | Homo sapiens | Feb 20 2026 | PRJNA1152578 | |
| GSE283746 | Homo sapiens | Jan 15 2026 | PRJNA1195123 | Blood |
| GSE315411 | Homo sapiens | Jan 07 2026 | PRJNA1396685 | Lung |
| GSE246946 | Homo sapiens | Sep 30 2025 | PRJNA1035137 | |
| GSE134873 | Mus musculus | Mar 10 2020 | PRJNA556663 | Rod photoreceptors |
| GSE282045 | Mus musculus | Jan 30 2026 | PRJNA1186708 | tumor |
| GSE221897 | Homo sapiens | Jan 01 2026 | PRJNA916709 | |
| GSE216402 | Mus musculus | Feb 14 2024 | PRJNA893551 | Lymph nodes |
| GSE317927 | Mus musculus | Jan 30 2026 | PRJNA1415822 | hippocampus microglia |
| GSE301807 | psis thaliana;Selaginella moelle | Jan 01 2026 | PRJNA1287585 | Root |
| GSE285873 | Mus musculus | Jan 01 2026 | PRJNA1206688 | Colon tumor |
| GSE156533 | Mus musculus | Aug 21 2020 | PRJNA658272 | Whole retina |
| GSE288838 | Mus musculus | Feb 26 2026 | | Testes |
| GSE252078 | Danio rerio | Dec 25 2025 | PRJNA1057281 | brain |
| GSE315344 | Homo sapiens | Jan 27 2026 | PRJNA1395699 | J-Lat cell line |
| GSE314158 | Homo sapiens | Feb 18 2026 | PRJNA1390185 | skin |
| GSE276443 | Homo sapiens | Dec 24 2024 | PRJNA1157376 | |
| GSE261260 | Homo sapiens | Feb 25 2026 | PRJNA1086183 | T cell |
| GSE316782 | Homo sapiens | Jan 21 2026 | PRJNA1404196 | Tumor and lymph node |
| GSE215962 | Mus musculus | Feb 18 2026 | PRJNA891350 | noma cell line-derived tumor |
| GSE141327 | Homo sapiens | Feb 05 2020 | PRJNA593183 | retina |
| GSE226096 | Mus musculus | Feb 06 2026 | PRJNA938498 | |
| GSE282072 | Arabidopsis thaliana | Feb 06 2026 | PRJNA1186950 | root |
| GSE277422 | Mus musculus | Jan 13 2026 | PRJNA1162148 | FL-BMDC |
| GSE275837 | Mus musculus | Feb 28 2026 | PRJNA1153283 | lepa1-6 orthotopic tumor |
| GSE311064 | Mus musculus | Jan 01 2026 | PRJNA1368618 | Liver |
| GSE181095 | Mus musculus | Dec 02 2022 | PRJNA750706 | retina |
| GSE307339 | Mus musculus | Feb 24 2026 | | |
| GSE306125 | Mus musculus | Jan 14 2026 | PRJNA1309495 | Breast |
| GSE194211 | Mus musculus | Jan 01 2026 | PRJNA799741 | adrenal_glands |
| GSE311219 | Homo sapiens | Jan 20 2026 | PRJNA1368817 | liver |
| GSE253425 | Mus musculus | Jan 01 2026 | PRJNA1065779 | Brain |
| GSE153645 | Homo sapiens | Jul 10 2020 | PRJNA643537 | |
| GSE315505 | Mus musculus | Jan 06 2026 | PRJNA1398077 | lung |
| GSE237650 | Mus musculus | Dec 20 2023 | PRJNA996105 | retina |
| GSE261416 | Mus musculus | Feb 17 2026 | PRJNA1086908 | |

| | | | | |
|---|---|---|---|---|
| GSE306883 | Mus musculus | Jan 14 2026 | PRJNA1312852 | Heart |
| GSE277536 | Lyssavirus rabies | Feb 17 2026 | PRJNA1162744 | |
| GSE297480 | Homo sapiens | Jan 21 2026 | PRJNA1264983 | |
| GSE207928 | Mus musculus | Jul 11 2022 | PRJNA857704 | Retinal Tissue |
| GSE331109 | Danio rerio | May 20 2026 | PRJNA1466908 | eye |
| GSE161369 | Homo sapiens | Jan 16 2026 | PRJNA677965 | |
| GSE306112 | Homo sapiens | Jan 08 2026 | PRJNA1309142 | Prostate |
| GSE194226 | Mus musculus | Feb 22 2023 | PRJNA799766 | retina |
| GSE153672 | Mus musculus | Oct 23 2020 | PRJNA643590 | |
| GSE296383 | Homo sapiens | Jan 20 2026 | PRJNA1259030 | Breast |
| GSE315532 | Rattus norvegicus | Jan 07 2026 | PRJNA1398208 | testis |
| GSE289296 | Mus musculus | Feb 28 2026 | PRJNA1222364 | kidney |
| GSE160308 | Homo sapiens | Apr 19 2021 | PRJNA672929 | Retina Macula |
| GSE302473 | Mus musculus | Feb 06 2026 | PRJNA1291071 | |
| GSE314553 | Homo sapiens | Jan 14 2026 | PRJNA1391499 | DIV60 neurons |
| GSE160617 | Macaca fascicularis | Mar 31 2021 | PRJNA673804 | |
| GSE295538 | Mus musculus | May 13 2025 | PRJNA1255151 | retina |
| GSE315250 | Mus musculus | Jan 04 2026 | PRJNA1395479 | |
| GSE304737 | hila melanogaster;Drosophila si | Feb 09 2026 | PRJNA1302423 | Brain |
| GSE226462 | Mus musculus | Feb 28 2026 | PRJNA940388 | ower-limb skeletal muscle |
| GSE317317 | Homo sapiens | Jan 24 2026 | PRJNA1406731 | |
| GSE247317 | Mus musculus | Dec 22 2023 | PRJNA1037101 | |
| GSE152491 | Mus musculus | Nov 30 2021 | PRJNA639530 | urified rod photoreceptors |
| GSE301728 | Homo sapiens | Jan 20 2026 | | bone marrow |
| GSE192600 | Mus musculus | Apr 27 2022 | PRJNA792265 | |
| GSE289014 | Mus musculus | Sep 30 2025 | PRJNA1220615 | retina |
| GSE262252 | Homo sapiens | Jan 08 2026 | PRJNA1090906 | Breast |
| GSE285521 | Mus musculus | Jan 04 2026 | PRJNA1204296 | aortic arch |
| GSE300749 | Homo sapiens | Jan 01 2026 | PRJNA1282023 | Breast |
| GSE203527 | Euprymna berryi | Mar 20 2025 | PRJNA841223 | Suckers from arms |
| GSE218153 | Homo sapiens | Dec 28 2022 | PRJNA902418 | |
| GSE146887 | Homo sapiens | Apr 09 2020 | PRJNA612302 | |
| GSE305756 | Mus musculus | Feb 13 2026 | PRJNA1307562 | retinal |
| GSE261694 | Mus musculus | Feb 19 2026 | PRJNA1088435 | tumor |
| GSE281715 | Mus musculus | Feb 23 2026 | PRJNA1185278 | kin, blood, bone marrow |
| GSE313641 | Homo sapiens | Jan 26 2026 | | nasal epithelium |
| GSE276911 | Mus musculus | Jan 01 2026 | PRJNA1159702 | Bone Marrow |
| GSE254062 | synthetic construct | Jan 31 2026 | PRJNA1068483 | blood |
| GSE266579 | Mus musculus | Jan 01 2026 | PRJNA1107620 | anoids from mouse intestine |
| GSE315974 | Homo sapiens | Jan 15 2026 | PRJNA1400047 | differentiated |
| GSE308557 | Mus musculus | Jan 02 2026 | PRJNA1330881 | cortex |
| GSE299954 | Homo sapiens | Jan 20 2026 | PRJNA1277701 | |
| GSE209872 | Rattus norvegicus | Nov 25 2022 | | retina |
| GSE259254 | Homo sapiens | Feb 10 2025 | PRJNA1080538 | cell |
| GSE233822 | Mus musculus | May 16 2025 | PRJNA978051 | Retina |
| GSE267007 | Homo sapiens | Jan 12 2026 | PRJNA1109337 | |
| GSE247320 | Mus musculus | Dec 22 2023 | PRJNA1037095 | |
| GSE214210 | Homo sapiens | Apr 19 2023 | PRJNA884380 | |
| GSE305642 | Mus musculus | Jan 14 2026 | PRJNA1307398 | |
| GSE313762 | Mus musculus | Feb 18 2026 | PRJNA1381556 | |
| GSE312061 | Mus musculus | Jan 13 2026 | PRJNA1372128 | Liver |
| GSE239301 | Mus musculus | Jan 14 2026 | PRJNA998778 | Bone marrow |
| GSE119246 | Mus musculus | Aug 30 2021 | PRJNA488604 | retina |
| GSE299877 | Mus musculus | Jan 21 2026 | PRJNA1277210 | retina |
| GSE315857 | Homo sapiens | Jan 14 2026 | PRJNA1399448 | Eutopic endometrium |
| GSE310848 | Rattus norvegicus | Feb 24 2026 | PRJNA1367160 | Skeletal Muscle |
| GSE237293 | Mus musculus | Aug 15 2023 | PRJNA994529 | etinal pigment epithelium |
| GSE318492 | Mus musculus | Feb 05 2026 | PRJNA1419027 | Skin |
| GSE316028 | Homo sapiens | Jan 11 2026 | PRJNA1400489 | |
| GSE224684 | Mus musculus | Feb 06 2026 | PRJNA932037 | Heart |
| GSE313528 | Homo sapiens | Feb 11 2026 | PRJNA1380109 | hiPSC-derived |
| GSE300899 | Homo sapiens | Feb 09 2026 | PRJNA1282625 | |
| GSE299840 | Homo sapiens | Jan 14 2026 | PRJNA1277109 | |

| | | | | |
|---|---|---|---|---|
| GSE290686 | Mus musculus | Jan 15 2026 | PRJNA1229540 | Liver |
| GSE315860 | Mus musculus | Jan 08 2026 | PRJNA1399447 | testis |
| GSE313037 | Rattus norvegicus | Feb 19 2026 | PRJNA1378171 | EVs |
| GSE310393 | Homo sapiens | Jan 05 2026 | PRJNA1365874 | Peripheral blood |
| GSE271751 | Homo sapiens | May 28 2025 | PRJNA1133031 | Organoid |
| GSE317049 | Mus musculus | Feb 05 2026 | PRJNA1405704 | guinal white adipose tissue |
| GSE312549 | Homo sapiens | Feb 26 2026 | PRJNA1374147 | neuroblastoma |
| GSE163753 | Mus musculus | Dec 24 2020 | PRJNA687475 | retina, Muller glia |
| GSE318286 | Homo sapiens | Feb 05 2026 | PRJNA1418526 | diffuse midline glioma |
| GSE252860 | Mus musculus | Jan 01 2026 | PRJNA1062840 | retina |
| GSE164489 | Mus musculus | Jan 10 2021 | | retina |
| GSE201680 | Homo sapiens | Apr 30 2022 | PRJNA832667 | |
| GSE304037 | Mus musculus | Jun 11 2026 | PRJNA1298818 | Ear Fibroblasts |
| GSE279644 | Homo sapiens;Mus musculus | Feb 27 2026 | PRJNA1173753 | bone marrow |
| GSE202759 | Mus musculus | Jun 07 2023 | PRJNA837217 | |
| GSE238216 | Mus musculus | Jun 06 2024 | PRJNA998380 | Retina |
| GSE325371 | Solanum lycopersicum | Mar 20 2026 | | leaf |
| GSE313176 | Mus musculus | Dec 10 2025 | PRJNA1378747 | Retina |
| GSE305056 | Mus musculus | Jan 09 2026 | PRJNA1304377 | iWAT |
| GSE292402 | Homo sapiens | Jan 16 2026 | PRJNA1238320 | |
| GSE234723 | Drosophila melanogaster | Feb 24 2026 | PRJNA982810 | Ovary |
| GSE136399 | Homo sapiens | May 19 2020 | PRJNA562493 | |
| GSE246169 | Homo sapiens | Oct 30 2023 | PRJNA1031951 | retina |
| GSE193161 | Homo sapiens | Mar 04 2022 | PRJNA795163 | |
| GSE309236 | Mus musculus | Jan 12 2026 | PRJNA1334386 | Heart |
| GSE261832 | Homo sapiens | Jan 12 2026 | PRJNA1089182 | |
| GSE225960 | Arabidopsis thaliana | Feb 28 2026 | PRJNA938110 | leaf |
| GSE280209 | Rattus norvegicus | Jan 26 2026 | PRJNA1177365 | |
| GSE308748 | Mus musculus | Feb 25 2026 | PRJNA1332648 | Brain cortical plate |
| GSE317420 | Danio rerio | Jan 29 2026 | PRJNA1412571 | whole zebrafish |
| GSE285574 | Mus musculus | Jan 01 2026 | PRJNA1204912 | prostate |
| GSE274638 | Homo sapiens | Jan 14 2026 | PRJNA1147322 | oft tissue tumors in PDXs |
| GSE285335 | Homo sapiens | Jan 21 2026 | PRJNA1202919 | Peripheral blood |
| GSE316562 | Homo sapiens | Jan 23 2026 | PRJNA1403396 | liver |
| GSE289162 | Mus musculus | Feb 11 2026 | PRJNA1221832 | liver |
| GSE228486 | | Apr 10 2023 | | |
| GSE121820 | Homo sapiens | Jan 07 2020 | PRJNA498524 | Whole blood |
| GSE291841 | feces metagenome | Jan 28 2026 | | Feces |
| GSE254123 | Rattus norvegicus | May 31 2024 | PRJNA1068598 | neural retina |
| GSE290820 | Saccharomyces cerevisiae | Feb 14 2026 | PRJNA1230175 | |
| GSE289600 | Mus musculus | Feb 08 2026 | | bone marrow |
| GSE232902 | Danio rerio | Jun 30 2023 | PRJNA974400 | Retina |
| GSE153099 | Homo sapiens | Feb 17 2021 | PRJNA641428 | |
| GSE281654 | Mus musculus | Jan 13 2026 | PRJNA1185167 | Bone marrow |
| GSE263427 | Mus musculus | May 10 2024 | PRJNA1097520 | RPE/choroid |
| GSE308343 | Homo sapiens;Mus musculus | Jan 19 2026 | PRJNA1330237 | Kidney |
| GSE175750 | Homo sapiens | Sep 27 2021 | PRJNA733485 | |
| GSE304885 | Mus musculus | Feb 13 2026 | PRJNA1303169 | Small intestine |
| GSE317534 | Mus musculus | Jan 26 2026 | PRJNA1414431 | hippocampus tissue |
| GSE230361 | Bos taurus | Feb 27 2026 | PRJNA960816 | Whole blood |
| GSE252943 | Mus musculus | Feb 06 2026 | PRJNA1063288 | Kidney |
| GSE280160 | Homo sapiens | Jan 16 2026 | PRJNA1176580 | uman PDA cell line - HPAFII |
| GSE299822 | Gallus gallus | Jan 12 2026 | PRJNA1276570 | Embryonic spinal cord |
| GSE301669 | Homo sapiens | Feb 26 2026 | PRJNA1285951 | |
| GSE277197 | Mus musculus | Jan 21 2026 | PRJNA1161169 | Ileum |
| GSE142029 | Mus musculus | Mar 30 2020 | PRJNA595602 | retina |
| GSE272497 | Homo sapiens | Jan 12 2026 | PRJNA1136977 | epithelium |
| GSE315802 | Mus musculus | Jan 12 2026 | PRJNA1399337 | |
| GSE276694 | Homo sapiens | Mar 10 2025 | PRJNA1158713 | eye |
| GSE298852 | Homo sapiens;Mus musculus | Jan 21 2026 | PRJNA1271498 | Primary colorectal cancer |
| GSE304105 | Homo sapiens | Feb 18 2026 | PRJNA1298921 | |
| GSE291694 | Mus musculus | Jan 12 2026 | PRJNA1234669 | brain |
| GSE226250 | Arabidopsis thaliana | Feb 28 2026 | PRJNA939232 | Whole plant |

| | | | | |
|---|---|---|---|---|
| GSE182945 | Danio rerio | Aug 27 2022 | PRJNA758398 | |
| GSE308352 | Homo sapiens | Jan 15 2026 | PRJNA1330258 | cell |
| GSE276685 | Homo sapiens | Feb 18 2026 | PRJNA1158697 | MIBC |
| GSE318858 | Mus musculus | Feb 17 2026 | PRJNA1420723 | Heart |
| GSE309333 | Mus musculus | Jan 09 2026 | PRJNA1335190 | brain |
| GSE280613 | Mus musculus | Feb 18 2026 | PRJNA1179493 | |
| GSE314681 | Mus musculus | Jan 04 2026 | PRJNA1392443 | Heart |
| GSE308407 | Mus musculus | Jan 13 2026 | PRJNA1330349 | |
| GSE206622 | Mus musculus | Aug 20 2022 | PRJNA851595 | retina |
| GSE230425 | Mus musculus | Dec 18 2023 | PRJNA961117 | retina |
| GSE302289 | Mus musculus | Feb 17 2026 | PRJNA1289799 | guinal white adipose tissue |
| GSE168928 | Mus musculus | Mar 16 2021 | PRJNA714613 | |
| GSE292265 | Mus musculus | Feb 12 2026 | PRJNA1237778 | |
| GSE316211 | Mus musculus | Feb 23 2026 | PRJNA1401802 | Gingiva |
| GSE239410 | Danio rerio | Aug 31 2023 | PRJNA999234 | Retina |
| GSE243711 | Mus musculus | Jan 12 2026 | PRJNA1019678 | |
| GSE297765 | Mus musculus | Jan 07 2026 | PRJNA1266144 | epidermis |
| GSE319480 | Mus musculus | Feb 13 2026 | PRJNA1423518 | |
| GSE226778 | Homo sapiens | Oct 19 2023 | PRJNA941788 | |
| GSE281645 | Homo sapiens | Sep 17 2025 | PRJNA1185147 | |
| GSE317512 | Homo sapiens | Feb 11 2026 | PRJNA1414398 | primary osteosarcoma |
| GSE282283 | Homo sapiens;Mus musculus | Feb 19 2026 | PRJNA1188059 | eye |
| GSE312012 | Homo sapiens | Jan 31 2026 | PRJNA1372052 | |
| GSE309466 | Solanum tuberosum | Jan 01 2026 | PRJNA1336128 | tuber |
| GSE223439 | Mus musculus | Nov 08 2023 | PRJNA926323 | Retina |
| GSE275648 | Homo sapiens | Feb 11 2026 | PRJNA1152546 | Primary tumor |
| GSE315096 | Mus musculus | Jan 06 2026 | PRJNA1394989 | CD1 mouse fetal kidney |
| GSE277903 | Mus musculus | Jan 21 2026 | PRJNA1164729 | Epidermis |
| GSE215321 | Homo sapiens | Mar 31 2023 | PRJNA889765 | |
| GSE290083 | Arabidopsis thaliana | Jan 29 2026 | PRJNA1226175 | Seedlings |
| GSE214340 | Homo sapiens | Sep 25 2025 | PRJNA885025 | |
| GSE308759 | Mus musculus | Feb 13 2026 | PRJNA1332679 | kidney |
| GSE252846 | Mus musculus | Jan 01 2026 | PRJNA1062812 | Pancreas |
| GSE301511 | Mus musculus | Jan 01 2026 | PRJNA1285366 | P60 cortex |
| GSE226373 | Danio rerio | Mar 03 2023 | PRJNA940076 | Retina |
| GSE317406 | Mus musculus | Feb 11 2026 | PRJNA1412541 | RM-1 tumors |
| GSE312897 | Mus musculus | Jan 30 2026 | PRJNA1377565 | |
| GSE202257 | Mus musculus | May 07 2022 | PRJNA835142 | Superior Colliculus |
| GSE317119 | Oryctolagus cuniculus | Jan 24 2026 | PRJNA1406124 | iliac artery |
| GSE288997 | Danio rerio | Feb 06 2026 | PRJNA1220571 | Heart |
| GSE301039 | Homo sapiens | Feb 09 2026 | PRJNA1283111 | |
| GSE312664 | Homo sapiens | Jan 15 2026 | PRJNA1375676 | striatal organoids |
| GSE297113 | Homo sapiens | Feb 26 2026 | PRJNA1262741 | |
| GSE281033 | Mus musculus | Jun 23 2025 | PRJNA1181861 | Retina |
| GSE315930 | Dactylorhiza majalis | Jan 27 2026 | PRJNA1399969 | protocorm (seed) |
| GSE202474 | Homo sapiens | Sep 09 2022 | PRJNA836264 | Adult Retina |
| GSE315871 | Sus scrofa | Jan 12 2026 | PRJNA1399468 | Muscle |
| GSE295688 | Mus musculus | Oct 07 2025 | PRJNA1255593 | Retinal Explant |
| GSE288625 | Mus musculus | Jan 31 2026 | PRJNA1218719 | brain |
| GSE317225 | Mus musculus | Jan 27 2026 | PRJNA1406306 | bone marrow |
| GSE316039 | Homo sapiens | Jan 14 2026 | PRJNA1400508 | colorectum |
| GSE313539 | Mus musculus | Jan 31 2026 | PRJNA1380130 | |
| GSE130610 | Rattus norvegicus | May 01 2022 | PRJNA540770 | retina |
| GSE316244 | Mus musculus | Feb 24 2026 | PRJNA1401859 | Distal lung |
| GSE292230 | Mus musculus | Feb 24 2026 | PRJNA1237686 | gastrocnemius |
| GSE289644 | Bos taurus | Jan 08 2026 | PRJNA1223881 | nusculus longissimus dorsi |
| GSE310382 | Homo sapiens | Jan 05 2026 | PRJNA1365862 | Peripheral blood |
| GSE317547 | Arabidopsis thaliana | Jan 28 2026 | PRJNA1414443 | leaf |
| GSE244810 | Mus musculus | Aug 13 2024 | PRJNA1024937 | Eye |
| GSE302930 | Homo sapiens | Feb 16 2026 | PRJNA1292852 | h node biopsy; Muscle invasive; T3aN0Mx |
| GSE223705 | Mus musculus | Feb 16 2023 | PRJNA927319 | brain |
| GSE312070 | Mus musculus | Jan 13 2026 | PRJNA1372140 | |
| GSE303795 | Homo sapiens | Jan 27 2026 | PRJNA1297696 | cell line |

| | | | | |
|---|---|---|---|---|
| GSE285346 | Mus musculus | Jan 21 2026 | PRJNA1203201 | |
| GSE270716 | Homo sapiens | Jan 01 2026 | PRJNA1128071 | Skin |
| GSE283280 | Mus musculus | Feb 20 2026 | | E14_5_Gonad |
| GSE181325 | Mus musculus | Dec 20 2022 | PRJNA751573 | retina |
| GSE287912 | Homo sapiens | Feb 23 2026 | PRJNA1215268 | Spleen |
| GSE180778 | Danio rerio | Jul 01 2022 | PRJNA749559 | Eye |
| GSE311595 | Mus musculus | Jan 26 2026 | PRJNA1369980 | bone marrow |
| GSE277792 | Danio rerio | Jan 30 2025 | PRJNA1164198 | Retina |
| GSE268687 | Mus musculus | Feb 24 2026 | PRJNA1118268 | sal hippocampus hemisphere |
| GSE309245 | Mus musculus | Jan 22 2026 | PRJNA1334403 | Spleen |
| GSE294580 | Homo sapiens | Jan 14 2026 | PRJNA1250525 | Bone Marrow |
| GSE291911 | Mus musculus | Jan 16 2026 | PRJNA1235769 | Spleen |
| GSE289032 | Mus musculus | Feb 06 2026 | PRJNA1220674 | |
| GSE287488 | Mus musculus | Jan 25 2025 | PRJNA1213082 | organoids |
| GSE312319 | Homo sapiens | Jan 23 2026 | PRJNA1373026 | brain |
| GSE254726 | Mus musculus | Feb 15 2026 | PRJNA1071576 | |
| GSE274401 | Homo sapiens | Jan 20 2026 | PRJNA1146351 | Breast |
| GSE287197 | Mus musculus | Feb 17 2026 | PRJNA1211585 | Lung |
| GSE308213 | Mus musculus | Jan 13 2026 | PRJNA1329210 | Brain |
| GSE206529 | Homo sapiens | Aug 01 2022 | PRJNA851279 | ent derived Retinal organoids |
| GSE155038 | Mus musculus | Aug 13 2023 | PRJNA648317 | |
| GSE300758 | Homo sapiens | Jan 04 2026 | PRJNA1282052 | Brain |
| GSE293410 | Homo sapiens | Jan 12 2026 | PRJNA1244382 | aorta |
| GSE272538 | Homo sapiens | Jan 21 2026 | PRJNA1137308 | Breast |
| GSE211188 | Homo sapiens | Aug 14 2023 | PRJNA869450 | RPE |
| GSE222390 | Homo sapiens | Jan 12 2023 | PRJNA921858 | retinal epithelium |
| GSE308091 | Homo sapiens | Feb 17 2026 | PRJNA1328721 | Retina |
| GSE297491 | Mus musculus | Jan 27 2026 | PRJNA1265079 | |
| GSE288584 | Mus musculus | Jan 14 2026 | PRJNA1218633 | |
| GSE271383 | Mus musculus | Jan 18 2026 | PRJNA1131281 | Kidney glomeruli |
| GSE283409 | Homo sapiens | Feb 13 2026 | PRJNA1193593 | Endometrium |
| GSE186369 | Mus musculus | Oct 19 2022 | PRJNA773562 | Retina |
| GSE153101 | Homo sapiens | Feb 17 2021 | PRJNA641425 | |
| GSE298562 | Mus musculus | Feb 18 2026 | PRJNA1269940 | non-small cell lung cancer |
| GSE289287 | Homo sapiens | Feb 10 2026 | PRJNA1222348 | in vitro cell culture |
| GSE302462 | Arabidopsis thaliana | Jan 26 2026 | PRJNA1291041 | seedling |
| GSE286416 | Homo sapiens | Feb 14 2026 | PRJNA1209102 | |
| GSE171845 | Homo sapiens | Oct 01 2021 | PRJNA721187 | |
| GSE283121 | Lotus japonicus | Feb 05 2026 | PRJNA1192200 | root |
| GSE269306 | Mus musculus | Jan 06 2026 | PRJNA1120933 | |
| GSE172512 | Homo sapiens | Sep 21 2022 | PRJNA723590 | |
| GSE248722 | Mus musculus | Jan 22 2026 | PRJNA1045656 | Forebrain |
| GSE262967 | Homo sapiens | Jan 06 2026 | PRJNA1095010 | |
| GSE294060 | Homo sapiens;Mus musculus | Jan 28 2026 | PRJNA1247972 | |
| GSE300963 | Mus musculus | Jan 10 2026 | PRJNA1282747 | hippocampus |
| GSE311208 | Mus musculus | Feb 25 2026 | PRJNA1368813 | Bone marrow |
| GSE314708 | Mus musculus | Feb 13 2026 | PRJNA1392484 | Cortex |
| GSE314456 | Homo sapiens | Jan 12 2026 | PRJNA1391351 | ose stromal vascular fraction |
| GSE136563 | Mus musculus | Jan 08 2020 | PRJNA562697 | retina |
| GSE317901 | Homo sapiens | Jan 30 2026 | PRJNA1415770 | Oropharynx |
| GSE311156 | Mus musculus | Feb 23 2026 | PRJNA1368760 | striatum nucleus |
| GSE307569 | Homo sapiens | Feb 18 2026 | PRJNA1322298 | bone marrow |
| GSE314334 | Homo sapiens | Jan 09 2026 | PRJNA1390736 | |
| GSE290340 | Homo sapiens | Feb 24 2026 | PRJNA1227510 | |
| GSE211583 | Mus musculus | Dec 19 2023 | PRJNA870946 | |
| GSE303999 | Mus musculus | Feb 13 2026 | PRJNA1298607 | knee synovium |
| GSE318163 | Penicillium oxalicum | Jan 31 2026 | PRJNA1416830 | |
| GSE214912 | Homo sapiens | Mar 31 2023 | PRJNA887734 | |
| GSE287554 | Homo sapiens | Jan 01 2026 | PRJNA1213738 | liver |
| GSE316957 | Homo sapiens | Jan 21 2026 | PRJNA1405594 | Femoral head |
| GSE282054 | Homo sapiens | Jan 21 2026 | PRJNA1186722 | |
| GSE160047 | Danio rerio | Jul 31 2021 | PRJNA671696 | |
| GSE284803 | Mus musculus | Jan 01 2026 | PRJNA1200692 | iWAT |

| | | | | |
|---|---|---|---|---|
| GSE310100 | Mus musculus | Jan 31 2026 | PRJNA1365139 | mammary gland |
| GSE319657 | Escherichia coli K-12 | Feb 24 2026 | PRJNA1424921 | strain |
| GSE274953 | Danio rerio | Jan 01 2026 | PRJNA1148636 | Whole embryo |
| GSE310891 | Homo sapiens | Feb 20 2026 | PRJNA1367491 | |
| GSE325932 | Mus musculus | Apr 01 2026 | PRJNA1442636 | Retina |
| GSE245362 | Mus musculus | Sep 11 2024 | PRJNA1027889 | Ganglion cell layer |
| GSE315362 | Homo sapiens | Jan 06 2026 | PRJNA1396600 | PBMCs / B cells |
| GSE319135 | Mus musculus | Feb 16 2026 | PRJNA1422162 | |
| GSE318329 | Homo sapiens | Feb 11 2026 | PRJNA1418594 | Peripheral blood |
| GSE305384 | Homo sapiens | Feb 27 2026 | PRJNA1305419 | Blood |
| GSE307041 | Mus musculus | Feb 19 2026 | PRJNA1314157 | Thymus |
| GSE317936 | Homo sapiens | Jan 31 2026 | PRJNA1415851 | metastatic melanoma |
| GSE291748 | Escherichia coli | Jan 19 2026 | PRJNA1235181 | |
| GSE310723 | Homo sapiens | Jan 09 2026 | PRJNA1366985 | |
| GSE306603 | Mus musculus | Jan 01 2026 | PRJNA1311443 | kidney |
| GSE299203 | Mus musculus | Jan 01 2026 | PRJNA1273057 | |
| GSE254948 | Danio rerio | Jan 02 2025 | PRJNA1072679 | Retina |
| GSE309092 | Rattus | Dec 31 2025 | PRJNA1333942 | Retina |
| GSE316187 | Caenorhabditis elegans | Jan 12 2026 | | Whole organism |
| GSE302362 | Mus musculus | Jan 22 2026 | PRJNA1290252 | Lung |
| GSE239299 | Mus musculus | Jan 14 2026 | PRJNA998783 | Bone marrow |
| GSE245541 | Mus musculus | Jun 06 2024 | PRJNA1028972 | retina |
| GSE315576 | Homo sapiens | Jan 09 2026 | PRJNA1398281 | |
| GSE273771 | Mus musculus | Jan 01 2026 | PRJNA1143034 | whole ovary |
| GSE287422 | Homo sapiens | Jan 21 2026 | PRJNA1212732 | |
| GSE315069 | Mus musculus | Feb 18 2026 | PRJNA1394961 | cell line |
| GSE311986 | Homo sapiens | Jan 30 2026 | PRJNA1372012 | |
| GSE303334 | Homo sapiens | Jan 01 2026 | PRJNA1294858 | Chondrosarcoma |
| GSE309934 | Homo sapiens | Feb 23 2026 | PRJNA1364014 | |
| GSE144429 | Mus musculus | Mar 09 2020 | PRJNA603598 | tumor |
| GSE237700 | Schizosaccharomyces pombe | Jan 01 2026 | PRJNA996182 | |
| GSE311349 | Mus musculus | Feb 23 2026 | PRJNA1368995 | ovary |
| GSE294121 | Homo sapiens | Jan 20 2026 | | skin |
| GSE313593 | Canis lupus familiaris | Jan 08 2026 | PRJNA1380358 | |
| GSE275794 | Mus musculus | Jan 28 2026 | PRJNA1153003 | ileum |
| GSE252276 | Homo sapiens | Jan 01 2024 | PRJNA1059046 | |
| GSE226787 | Homo sapiens | Jan 01 2026 | PRJNA941795 | |
| GSE318663 | Homo sapiens | Feb 15 2026 | PRJNA1419736 | Tumor - Lung |
| GSE315628 | Homo sapiens;Mus musculus | Jan 06 2026 | PRJNA1398356 | bladder cancer |
| GSE252514 | Mus musculus | Jan 26 2024 | PRJNA1061067 | |
| GSE310328 | Ascaris suum | Jan 01 2026 | PRJNA1365768 | Whole organism |
| GSE307014 | Homo sapiens | Jan 22 2026 | PRJNA1314116 | |
| GSE309820 | Mus musculus | Jan 05 2026 | PRJNA1363765 | kidney |
| GSE284692 | Mus musculus | Feb 21 2026 | PRJNA1200052 | CD43- Splenic B cells |
| GSE128442 | Mus musculus | Feb 25 2022 | PRJNA527760 | retinal |
| GSE298423 | Mus musculus | Feb 28 2026 | PRJNA1269435 | |
| GSE279296 | Mus musculus | Jan 08 2026 | PRJNA1171817 | |
| GSE143937 | Homo sapiens | Jan 01 2021 | PRJNA602294 | bone marrow |
| GSE318654 | Mus musculus | Feb 16 2026 | PRJNA1419721 | se Lumbar Spinal Cord (L1-L5) |
| GSE287254 | Homo sapiens | Jan 01 2026 | PRJNA1211734 | SC-derived brain organoid |
| GSE233749 | Mus musculus | Sep 30 2023 | PRJNA977695 | retina |
| GSE241892 | Homo sapiens | Feb 18 2026 | PRJNA1010699 | |
| GSE283548 | Mus musculus | Jan 08 2026 | PRJNA1194509 | tumor |
| GSE282754 | Zea mays | Jan 29 2026 | PRJNA1190367 | leaf |
| GSE200233 | Mus musculus | Apr 05 2023 | PRJNA823714 | Retina |
| GSE211958 | Homo sapiens | Aug 28 2022 | PRJNA873054 | |
| GSE264046 | Homo sapiens | Jan 01 2026 | PRJNA1100597 | |
| GSE318429 | Homo sapiens | Feb 11 2026 | PRJNA1418718 | Liver |
| GSE277466 | Homo sapiens | Jan 14 2026 | PRJNA1162480 | Prostate |
| GSE294887 | Homo sapiens;Mus musculus | Feb 28 2026 | PRJNA1251980 | LNCaP_42D |
| GSE130392 | Mus musculus | Jul 02 2022 | PRJNA539976 | Brain |
| GSE212110 | Homo sapiens | Apr 23 2023 | PRJNA874026 | Pigmented epithelium; Retina |
| GSE313619 | acillus acidoterrestris;Alicycloba | Feb 14 2026 | PRJNA1380408 | |

| | | | | |
|---|---|---|---|---|
| GSE249906 | Mus musculus | Jan 14 2026 | PRJNA1051110 | bone marrow |
| GSE236389 | Mus musculus | Jul 10 2023 | PRJNA990744 | ocular tissue |
| GSE189579 | Mus musculus | Nov 25 2022 | PRJNA783645 | Kidney |
| GSE285051 | Homo sapiens | Feb 24 2026 | PRJNA1201119 | testis-like organoids |
| GSE237804 | Rattus norvegicus | Feb 28 2024 | PRJNA996639 | whole retina |
| GSE289206 | Homo sapiens | Mar 03 2025 | PRJNA1221904 | vascular |
| GSE287878 | us serotinus;Rhinolophus ferrum | Jan 01 2026 | | Trachea organoid |
| GSE281530 | Homo sapiens | Feb 09 2026 | PRJNA1184337 | |
| GSE278337 | Mus musculus | Feb 17 2026 | PRJNA1166676 | |
| GSE288778 | Mus musculus | Jan 21 2026 | PRJNA1219239 | Lung |
| GSE138719 | Mus musculus | Oct 09 2022 | PRJNA576897 | retina |
| GSE310933 | Mus musculus | Jan 09 2026 | PRJNA1367568 | erior dorsal cerebellar vermis |
| GSE274448 | Homo sapiens | Feb 15 2026 | PRJNA1146439 | |
| GSE243453 | Drosophila melanogaster | Feb 27 2026 | PRJNA1018515 | embryo |
| GSE281507 | Mus musculus | Jan 23 2026 | PRJNA1184158 | neuro, adBNST |
| GSE167208 | Homo sapiens | Feb 23 2021 | PRJNA703927 | |
| GSE245595 | Mus musculus | May 13 2024 | PRJNA1029042 | RPE-choroid eyecups |
| GSE311289 | Homo sapiens | Feb 11 2026 | PRJNA1368892 | Peripheral blood |
| GSE284996 | Homo sapiens | Jan 21 2026 | PRJNA1200768 | |
| GSE308745 | Homo sapiens | Feb 05 2026 | PRJNA1332649 | heral Blood Mononuclear Cells |
| GSE275136 | Mus musculus | Feb 24 2026 | PRJNA1149674 | cortex |
| GSE305211 | Homo sapiens | Jan 14 2026 | PRJNA1304766 | Thyroid |
| GSE280079 | Mus musculus | Feb 06 2026 | PRJNA1176149 | Tumor |
| GSE202763 | Homo sapiens | Jan 13 2026 | PRJNA837228 | |
| GSE292645 | Mus musculus | Jan 08 2026 | | oral palate |
| GSE102503 | Homo sapiens | Aug 10 2020 | PRJNA397896 | prostate epithelium |
| GSE235778 | Homo sapiens | Jun 01 2026 | PRJNA987610 | |
| GSE223658 | synthetic construct | Nov 08 2023 | PRJNA927050 | |
| GSE274635 | Homo sapiens | Jan 14 2026 | PRJNA1147320 | |
| GSE263987 | Homo sapiens | Oct 21 2024 | PRJNA1100507 | iPSC/SEAM_week0 |
| GSE304512 | Homo sapiens | Jan 28 2026 | PRJNA1301410 | Liver |
| GSE295911 | Homo sapiens | Jan 28 2026 | PRJNA1256768 | liver |
| GSE316070 | Homo sapiens | Jan 09 2026 | PRJNA1400555 | Cell line |
| GSE280638 | Mus musculus | Feb 23 2026 | PRJNA1179867 | ovary |
| GSE278223 | Mus musculus | Feb 25 2026 | PRJNA1166125 | spleen |
| GSE256096 | Homo sapiens | Feb 18 2026 | PRJNA1078243 | |
| GSE180147 | Homo sapiens | Nov 03 2022 | PRJNA746775 | |
| GSE192506 | Mus musculus | Aug 16 2022 | PRJNA791675 | Retina |
| GSE234447 | Mus musculus | Jan 03 2024 | PRJNA981457 | retina |
| GSE205085 | Danio rerio | Mar 15 2023 | PRJNA843695 | whole liver |
| GSE288411 | Gallus gallus | Jan 31 2026 | PRJNA1217399 | |
| GSE235344 | Homo sapiens | Jan 05 2025 | PRJNA985717 | |
| GSE317011 | Mus musculus | Jan 24 2026 | PRJNA1405661 | bone marrow |
| GSE296552 | Mus musculus | Feb 11 2026 | PRJNA1259958 | Liver |
| GSE142746 | Homo sapiens | Sep 30 2022 | PRJNA598192 | |
| GSE319825 | Rattus norvegicus | Feb 22 2026 | | heart |
| GSE288344 | Homo sapiens;Mus musculus | Jan 17 2026 | PRJNA1216978 | |
| GSE315481 | Homo sapiens | Jan 07 2026 | PRJNA1397222 | |
| GSE276699 | Homo sapiens | Mar 10 2025 | PRJNA1158718 | eye |
| GSE263660 | Homo sapiens | Jun 30 2025 | PRJNA1098720 | vitreous bodies |
| GSE154895 | Danio rerio | Feb 22 2021 | PRJNA647840 | eyes |
| GSE279008 | Mus musculus | Feb 20 2026 | PRJNA1169902 | heart ventricle |
| GSE279149 | Homo sapiens | Jan 29 2026 | PRJNA1170991 | |
| GSE316106 | Mus musculus | Jan 31 2026 | PRJNA1400607 | Left Quadricep muscle |
| GSE305039 | Homo sapiens | Jan 20 2026 | PRJNA1304274 | |
| GSE263043 | Homo sapiens;Mus musculus | Jan 06 2026 | PRJNA1095501 | lung |
| GSE309982 | Borreliella burgdorferi B31 | Jan 29 2026 | PRJNA1364871 | |
| GSE275163 | Danio rerio | Feb 26 2025 | | whole body cells |
| GSE174400 | Mus musculus | Jan 02 2022 | PRJNA729749 | |
| GSE285033 | Homo sapiens | Feb 08 2026 | PRJNA1201078 | bone narrow |
| GSE306682 | Mus musculus | Jan 13 2026 | PRJNA1311621 | GM-BMDC cultures |
| GSE279303 | Homo sapiens | Feb 10 2026 | PRJNA1171827 | |
| GSE241907 | Mus musculus | Feb 06 2026 | PRJNA1010735 | Heart |

| | | | | |
|---|---|---|---|---|
| GSE311096 | Homo sapiens | Feb 26 2026 | PRJNA1368650 | Kidney PDX |
| GSE180006 | Mus musculus | Sep 29 2021 | PRJNA746258 | retina |
| GSE296346 | Homo sapiens | Jan 01 2026 | PRJNA1258955 | |
| GSE163576 | Mus musculus | Dec 21 2021 | PRJNA686796 | |
| GSE226458 | Gallus gallus | Mar 11 2024 | PRJNA940372 | Retina |
| GSE313431 | Mus musculus | Jan 15 2026 | PRJNA1379500 | Bone marrow |
| GSE248932 | Ictidomys tridecemlineatus | Oct 23 2024 | PRJNA1046472 | Cortex |
| GSE289531 | Homo sapiens | Jan 12 2026 | PRJNA1223383 | tumor |
| GSE212914 | Homo sapiens | Sep 12 2022 | PRJNA878456 | |
| GSE270577 | Schizosaccharomyces pombe | Jan 27 2026 | PRJNA1127508 | |
| GSE293518 | Homo sapiens | Jan 28 2026 | PRJNA1244863 | |
| GSE199867 | Mus musculus | Apr 27 2022 | PRJNA821860 | Retina |
| GSE305350 | Mus musculus | Feb 10 2026 | PRJNA1305364 | implanted tumor |
| GSE303789 | Rattus norvegicus | Jan 27 2026 | PRJNA1297690 | Nucleus accumbens |
| GSE228867 | Homo sapiens | Feb 26 2026 | PRJNA951756 | |
| GSE231358 | Mus musculus | Sep 30 2023 | PRJNA964468 | retina |
| GSE308479 | Homo sapiens | Jan 01 2026 | PRJNA1330735 | Osteosarcoma |
| GSE305432 | Homo sapiens | Feb 13 2026 | PRJNA1305728 | |
| GSE318811 | Rattus norvegicus | Feb 18 2026 | PRJNA1420656 | left ventricular |
| GSE319683 | Mus musculus | Feb 18 2026 | PRJNA1424944 | Frontal cortex |
| GSE318826 | Homo sapiens | Feb 12 2026 | PRJNA1420688 | |
| GSE291819 | Mus musculus | Jan 16 2026 | PRJNA1235294 | cornea/limbus |
| GSE179568 | Homo sapiens | Oct 31 2021 | PRJNA744210 | RNV |
| GSE285110 | Mus musculus | Feb 10 2026 | PRJNA1201190 | bone marrow |
| GSE295867 | Procambarus virginalis | Jan 29 2026 | PRJNA1256517 | Hemolymph |
| GSE276808 | Mus musculus | Jan 22 2026 | PRJNA1159133 | lung |
| GSE203116 | Homo sapiens;Mus musculus | May 31 2023 | PRJNA838602 | Vitreous |
| GSE304464 | Mus musculus | Feb 19 2026 | PRJNA1301342 | Ileum |
| GSE174523 | Mus musculus | Nov 23 2022 | PRJNA730427 | Retinas and optic nerves |
| GSE242544 | Homo sapiens | Feb 11 2026 | PRJNA1013901 | Cervical adenocarcinoma |
| GSE291177 | Rattus norvegicus | Feb 25 2026 | PRJNA1232147 | Meibomian gland |
| GSE282557 | Homo sapiens | Jan 26 2026 | PRJNA1189131 | Heart |
| GSE285392 | Homo sapiens | Feb 11 2026 | PRJNA1203264 | cell line |
| GSE315892 | Mus musculus | Jan 09 2026 | PRJNA1399745 | Thymus |
| GSE286529 | Mus musculus | Jan 13 2025 | PRJNA1209861 | Retina |
| GSE287735 | Homo sapiens | Jan 08 2026 | PRJNA1214363 | |
| GSE315661 | Homo sapiens | Jan 11 2026 | PRJNA1398640 | |
| GSE256241 | Mus musculus | Jan 06 2026 | PRJNA1078766 | skeletal muscle |
| GSE305687 | Homo sapiens | Feb 11 2026 | PRJNA1307469 | |
| GSE298142 | Mus musculus | Jan 21 2026 | PRJNA1268419 | Bone marrow |
| GSE296976 | Mus musculus | Jan 13 2026 | PRJNA1262310 | liver |
| GSE290121 | Mus musculus | Jan 28 2026 | PRJNA1226235 | |
| GSE315349 | Rattus norvegicus | Jan 05 2026 | PRJNA1395867 | BAT EVs |
| GSE208053 | Homo sapiens | Aug 18 2022 | PRJNA858066 | Retinoblastoma |
| GSE298175 | Homo sapiens | Jan 22 2026 | PRJNA1268467 | |
| GSE307575 | Homo sapiens | Feb 15 2026 | PRJNA1322319 | skin |
| GSE314637 | Mus musculus | Jan 20 2026 | PRJNA1392352 | Livers |
| GSE225142 | Mus musculus | Aug 01 2024 | PRJNA934081 | retina |
| GSE307217 | Homo sapiens | Feb 17 2026 | PRJNA1314777 | B-Lymphocyte |
| GSE313963 | Homo sapiens;Mus musculus | Jan 06 2026 | PRJNA1387446 | Testis |
| GSE287060 | Homo sapiens | Jan 20 2026 | PRJNA1211074 | Plasma |
| GSE305843 | Macaca mulatta | Jan 01 2026 | PRJNA1308011 | PBMC |
| GSE307708 | Homo sapiens | Jan 05 2026 | PRJNA1327132 | connetive tissue |
| GSE290002 | Homo sapiens | Feb 20 2026 | PRJNA1225721 | |
| GSE270825 | Mus musculus | Feb 11 2026 | PRJNA1128581 | |
| GSE266905 | Oryza sativa | Jan 26 2026 | PRJNA1108699 | 14-day-old seedlings |
| GSE267797 | Homo sapiens | Jun 30 2025 | PRJNA1112889 | |
| GSE304090 | Mus musculus | Jan 01 2026 | PRJNA1298897 | Heart left ventricle |
| GSE198554 | Homo sapiens | Jun 26 2022 | PRJNA815960 | |
| GSE303428 | Rattus norvegicus | Jan 02 2026 | PRJNA1295390 | pancreatic islets |
| GSE315508 | Anthurium andraeanum | Jan 31 2026 | PRJNA1398173 | leaf |
| GSE315986 | Homo sapiens | Jan 12 2026 | PRJNA1400067 | |
| GSE156657 | Homo sapiens | Dec 31 2020 | PRJNA658590 | RB |

| | | | | |
|---|---|---|---|---|
| GSE289940 | Mus musculus | Jan 05 2026 | PRJNA1225604 | bone, bone marrow |
| GSE315017 | Laodelphax striatellus | Jan 03 2026 | PRJNA1394860 | midgut |
| GSE319768 | Homo sapiens | Feb 17 2026 | PRJNA1425072 | kidney tumor FFPE |
| GSE183117 | Mus musculus | Feb 01 2022 | PRJNA759217 | |
| GSE311223 | Homo sapiens | Jan 20 2026 | PRJNA1368833 | liver |
| GSE314723 | Bos taurus | Feb 23 2026 | PRJNA1392495 | |
| GSE261379 | Homo sapiens | Mar 12 2026 | PRJNA1086815 | |
| GSE271680 | Homo sapiens | Jan 07 2026 | PRJNA1132909 | |
| GSE301929 | Homo sapiens | Feb 06 2026 | PRJNA1287744 | |
| GSE177039 | Mus musculus | Jun 12 2021 | PRJNA736942 | whole retina |
| GSE303662 | Homo sapiens | Jan 28 2026 | | Brain |
| GSE286884 | Homo sapiens | Feb 18 2026 | PRJNA1210488 | breast cancer cell |
| GSE286115 | Homo sapiens | Jan 12 2026 | PRJNA1207954 | gnostic bone marrow sample |
| GSE254936 | Homo sapiens | Mar 04 2024 | PRJNA1072666 | Large intestine; Colon |
| GSE328096 | Mus musculus | Apr 15 2026 | PRJNA1453907 | retina |
| GSE318648 | Homo sapiens | Feb 28 2026 | | |
| GSE276895 | Homo sapiens | May 29 2025 | PRJNA1159360 | Retinoblastoma |
| GSE141660 | Homo sapiens | Dec 28 2022 | PRJNA594318 | stomach |
| GSE128141 | Homo sapiens | Jul 11 2020 | PRJNA526532 | |
| GSE183734 | Drosophila melanogaster | Mar 07 2022 | PRJNA761775 | retinas, optic lobes |
| GSE291677 | Mus musculus | Feb 26 2026 | PRJNA1234637 | retina explant |
| GSE302742 | Homo sapiens | Jan 29 2026 | PRJNA1292021 | |
| GSE264545 | Homo sapiens | Apr 15 2025 | PRJNA1103120 | |
| GSE300809 | Homo sapiens | Feb 12 2026 | | spinal cord |
| GSE290309 | Mus musculus | Feb 11 2026 | PRJNA1227463 | Ileum |
| GSE281855 | Mus musculus | Nov 15 2024 | PRJNA1186056 | Retina |
| GSE237203 | Mustela putorius furo | Oct 01 2023 | PRJNA994167 | Retina |
| GSE78181 | Blumeria hordei | Jan 31 2026 | PRJNA312918 | |
| GSE303714 | Mus musculus | Feb 17 2026 | PRJNA1297147 | |
| GSE294340 | Mus musculus | Feb 11 2026 | PRJNA1249058 | heart (ventricle) |
| GSE176353 | Rattus norvegicus | Jul 01 2021 | PRJNA735942 | retina |
| GSE256214 | Mus musculus | Nov 22 2024 | PRJNA1078424 | Retina |
| GSE292799 | Mus musculus | Jan 18 2026 | PRJNA1241277 | Ascites |
| GSE299614 | Homo sapiens | Feb 23 2026 | PRJNA1275683 | |
| GSE312960 | Mus musculus | Jan 13 2026 | PRJNA1377666 | spleen |
| GSE141657 | Homo sapiens | Dec 28 2022 | PRJNA594315 | stomach |
| GSE260982 | Mus musculus | Feb 14 2025 | PRJNA1084776 | Brain+Spinal Cord |
| GSE225642 | Homo sapiens | Mar 11 2024 | PRJNA937017 | etinal pigment epithelium |
| GSE301290 | Homo sapiens | Jan 22 2026 | PRJNA1284761 | euroblastoma Cancer Cells |
| GSE240575 | Danio rerio | Aug 24 2023 | PRJNA1004255 | Eye |
| GSE304790 | Danio rerio;Oncorhynchus mykis | Jan 08 2026 | PRJNA1302741 | (freshly fertilized egg/zygote) |
| GSE225003 | Homo sapiens | Feb 15 2023 | PRJNA933541 | Retina |
| GSE305093 | Mus musculus | Jan 05 2026 | | Pancreatic Islet |
| GSE318243 | Homo sapiens | Feb 05 2026 | | liver |
| GSE269667 | Mus musculus | Jun 21 2024 | PRJNA1123197 | retina |
| GSE263978 | Mus musculus | Apr 02 2025 | PRJNA1100496 | retina |
| GSE146186 | Mus musculus | May 19 2020 | PRJNA609681 | eye |
| GSE307403 | Homo sapiens | Feb 16 2026 | PRJNA1320683 | (dorsolateral prefrontal cortex) |
| GSE296800 | Homo sapiens | Feb 19 2026 | PRJNA1261067 | |
| GSE319577 | Homo sapiens | Feb 18 2026 | PRJNA1423778 | |
| GSE286177 | Homo sapiens | Jan 07 2026 | PRJNA1208069 | |
| GSE287674 | Mus musculus | Feb 25 2026 | PRJNA1214228 | Peritoneum and liver |
| GSE151956 | Mus musculus | Jun 08 2020 | PRJNA637887 | retina |
| GSE311241 | Mus musculus | Feb 12 2026 | PRJNA1368826 | adult mouse heart |
| GSE288298 | Mus musculus | Feb 16 2026 | PRJNA1216888 | |
| GSE298761 | Mus musculus | Jan 12 2026 | PRJNA1271064 | E13.5 fetal liver |
| GSE291529 | Homo sapiens | Feb 13 2026 | PRJNA1234098 | Liver |
| GSE287887 | Mus musculus | Jan 17 2026 | PRJNA1214862 | |
| GSE319215 | Homo sapiens | Feb 17 2026 | PRJNA1422638 | PBMCs |
| GSE153302 | Homo sapiens | Jun 25 2021 | PRJNA642134 | |
| GSE313815 | Mus musculus | Mar 02 2026 | PRJNA1381626 | |
| GSE168987 | Homo sapiens | Apr 05 2022 | PRJNA714867 | |
| GSE212468 | Homo sapiens | Nov 04 2022 | PRJNA875475 | |

| | | | | |
|---|---|---|---|---|
| GSE310894 | Anableps anableps | Apr 08 2026 | PRJNA1367496 | Dorsal retina |
| GSE318151 | Homo sapiens | Jan 30 2026 | PRJNA1416806 | |
| GSE186750 | Mus musculus | Sep 11 2023 | PRJNA775966 | |
| GSE249130 | Homo sapiens | Mar 01 2024 | PRJNA1047526 | |
| GSE309847 | Homo sapiens | Jan 31 2026 | PRJNA1363886 | |
| GSE315678 | Mus musculus | Jan 13 2026 | PRJNA1398775 | Liver |
| GSE283030 | Mus musculus | Jan 16 2026 | PRJNA1191470 | skull bone |
| GSE315367 | Homo sapiens | Jan 06 2026 | PRJNA1396606 | Bone marrow |
| GSE306747 | Homo sapiens | Jan 14 2026 | PRJNA1312029 | |
| GSE213887 | Mus musculus | May 22 2023 | PRJNA882852 | Retina |
| GSE291324 | Homo sapiens | Feb 06 2026 | PRJNA1232821 | |
| GSE315350 | Homo sapiens | Jan 05 2026 | PRJNA1395865 | Human skin (epidermis) |
| GSE236219 | Mus musculus | Jul 15 2023 | PRJNA989278 | RPE/choroid |
| GSE317095 | Mus musculus | Feb 16 2026 | PRJNA1405768 | |
| GSE220626 | Mus musculus | Sep 15 2023 | | retina |
| GSE286507 | Mus musculus | Jan 12 2026 | PRJNA1209825 | Renal cortex |
| GSE295127 | Homo sapiens | Feb 13 2026 | PRJNA1253601 | |
| GSE303512 | Mus musculus | Jan 15 2026 | PRJNA1295547 | E10.5 gonad |
| GSE316965 | Glycine max | Jan 20 2026 | PRJNA1405609 | leaves |
| GSE122970 | Homo sapiens | May 07 2020 | PRJNA507018 | retina |
| GSE282579 | Homo sapiens | Feb 23 2026 | PRJNA1189164 | Spleen |
| GSE310773 | Homo sapiens | Feb 11 2026 | PRJNA1367048 | Peripheral blood |
| GSE272777 | Homo sapiens | Feb 22 2026 | PRJNA1138714 | |
| GSE229730 | Mus musculus | Apr 18 2023 | PRJNA955717 | Retina |
| GSE318045 | Homo sapiens | Feb 16 2026 | PRJNA1416375 | rebral cortex - glioblastoma |
| GSE155288 | Homo sapiens | Aug 11 2021 | PRJNA649248 | retina |
| GSE315511 | Mus musculus | Apr 15 2026 | PRJNA1398191 | retina |
| GSE313859 | Homo sapiens | Jan 05 2026 | | Tumor |
| GSE283671 | Mus musculus | Jan 21 2026 | PRJNA1195021 | |
| GSE229218 | Mus musculus | Jun 06 2023 | PRJNA953420 | brain hemisection |
| GSE297494 | Mus musculus | Jan 23 2026 | PRJNA1265084 | l aortic perivascular adipose tissue |
| GSE316871 | Homo sapiens | Jan 20 2026 | PRJNA1405466 | Serum |
| GSE232373 | Mus musculus | Jun 07 2023 | PRJNA971879 | retina |
| GSE178842 | Danio rerio | Jan 25 2023 | PRJNA741229 | retina |
| GSE311047 | Tetraodon nigroviridis | Apr 08 2026 | PRJNA1368598 | Dorsal retina |
| GSE274399 | Homo sapiens | Aug 14 2024 | PRJNA1146349 | eye |
| GSE298078 | Pseudomonas aeruginosa | Jan 12 2026 | PRJNA1268316 | |
| GSE318651 | Mus musculus | Feb 10 2026 | | Bone-marrow derived |
| GSE311419 | Mus musculus | Jan 07 2026 | PRJNA1369432 | thyroid |
| GSE302526 | Mus musculus | Feb 18 2026 | PRJNA1291139 | liver |
| GSE228627 | Mus musculus | Dec 14 2023 | PRJNA950887 | Retina |
| GSE317951 | Homo sapiens | Jan 29 2026 | PRJNA1416084 | |
| GSE165520 | Rattus norvegicus | Jan 27 2021 | PRJNA694856 | extracted retinal tissue |
| GSE253547 | Mus musculus | Jan 18 2026 | PRJNA1066271 | Postnatal day P7 |
| GSE310167 | Homo sapiens | Jan 20 2026 | PRJNA1365230 | lung cancer |
| GSE218284 | Homo sapiens | Oct 11 2023 | PRJNA903087 | CD19 sorted cells from CNS |
| GSE233531 | Homo sapiens | Jan 01 2026 | PRJNA976619 | Liver |
| GSE311664 | Arabidopsis thaliana | Jan 29 2026 | PRJNA1370095 | of 3-week-old Arabidopsis plants |
| GSE290327 | Homo sapiens | Feb 24 2026 | | urothelial carcinoma of bladder |
| GSE275791 | Mus musculus | Jan 13 2026 | PRJNA1153009 | brain |
| GSE316096 | Homo sapiens | Jan 20 2026 | PRJNA1400587 | peripheral blood |
| GSE319318 | Mus musculus | Feb 18 2026 | PRJNA1422818 | reatic Ductal Adenocarcinoma |
| GSE166840 | Mus musculus | Feb 17 2021 | | Thymus |
| GSE309812 | Homo sapiens | Feb 23 2026 | PRJNA1363566 | |
| GSE277336 | Mus musculus | Feb 27 2026 | PRJNA1161785 | brain |
| GSE261216 | Mus musculus | Jan 20 2026 | PRJNA1085770 | Small intestine |
| GSE293912 | Mus musculus | Apr 08 2025 | PRJNA1247415 | Retina |
| GSE225344 | Mus musculus | May 03 2023 | PRJNA935311 | |
| GSE157271 | Mus musculus | Sep 02 2020 | PRJNA660735 | retina |
| GSE315332 | Bos taurus | Feb 23 2026 | PRJNA1395657 | |
| GSE291346 | Mus musculus | Jan 19 2026 | PRJNA1233192 | lung |
| GSE318666 | unclassified Streptomyces | Feb 13 2026 | PRJNA1419752 | |
| GSE212640 | Homo sapiens | Sep 05 2022 | PRJNA876300 | |

| | | | | |
|---|---|---|---|---|
| GSE302350 | Homo sapiens | Feb 17 2026 | PRJNA1290236 | SC-derived cerebral organoid |
| GSE319724 | cillus subtilis subsp. subtilis str. 1 | Feb 18 2026 | PRJNA1425009 | |
| GSE318538 | Homo sapiens | Feb 06 2026 | | |
| GSE230972 | Mus musculus | Jan 21 2026 | PRJNA962937 | Liver |
| GSE310073 | Rattus norvegicus | Feb 25 2026 | PRJNA1365087 | muscle |
| GSE311012 | Homo sapiens | Jan 14 2026 | PRJNA1368485 | |
| GSE314512 | Mus musculus | Jan 15 2026 | PRJNA1391437 | L929 cell line |
| GSE285080 | Homo sapiens | Feb 28 2026 | PRJNA1201154 | |
| GSE282645 | Gallus gallus | Feb 25 2026 | PRJNA1189531 | liver |
| GSE315211 | Mus musculus | Jan 14 2026 | PRJNA1395132 | Heart |
| GSE248745 | Mus musculus | Feb 06 2026 | PRJNA1045694 | ovary |
| GSE215191 | Mus musculus | Oct 13 2022 | PRJNA889069 | retina |
| GSE306606 | Homo sapiens | Feb 27 2026 | PRJNA1311464 | |
| GSE221053 | Homo sapiens | Aug 11 2023 | PRJNA912656 | |
| GSE311025 | ichia coli;Schizosaccharomyces p | Feb 16 2026 | PRJNA1368553 | strain:BW25113 |
| GSE310239 | Mus musculus | Jan 07 2026 | PRJNA1365517 | Liver |
| GSE298505 | Mus musculus | Feb 06 2026 | PRJNA1269570 | Hippocampus |
| GSE294030 | Mus musculus | Feb 25 2026 | PRJNA1247924 | ervical lymph nodes |
| GSE317681 | Homo sapiens | Jan 28 2026 | PRJNA1414632 | tumor |
| GSE245544 | Mus musculus | Jun 06 2024 | PRJNA1028973 | retina |
| GSE274186 | Mus | Feb 23 2026 | PRJNA1145376 | parotid gland |
| GSE311747 | Homo sapiens | Jan 12 2026 | PRJNA1370860 | KIDNEY |
| GSE282881 | Homo sapiens | Jan 04 2026 | PRJNA1190934 | Blood |
| GSE236112 | hatina fulica;Mizuhopecten yess | Jan 01 2026 | PRJNA989074 | heart |
| GSE319591 | Rattus norvegicus | Feb 18 2026 | PRJNA1423793 | whole brain hemisphere |
| GSE312103 | Mus musculus | Jan 25 2026 | PRJNA1372175 | spleen |
| GSE302874 | Mus musculus | Feb 27 2026 | PRJNA1292733 | liver |
| GSE270418 | Homo sapiens | Dec 30 2025 | PRJNA1126598 | |
| GSE267724 | Homo sapiens | Feb 13 2026 | PRJNA1112476 | |
| GSE297887 | Mus musculus | Jan 01 2026 | PRJNA1266676 | ppocampus;C57BL/6 male |
| GSE308009 | Mus musculus | Jan 20 2026 | PRJNA1328580 | Hindlimb knee joint |
| GSE254741 | Homo sapiens | Dec 06 2024 | PRJNA1071598 | |
| GSE225943 | Homo sapiens | Aug 18 2024 | PRJNA938058 | |
| GSE292109 | Mus musculus | Feb 25 2026 | PRJNA1237218 | |
| GSE138700 | Mus musculus | Nov 09 2021 | PRJNA576864 | spleen |
| GSE304776 | Mus musculus | Feb 13 2026 | PRJNA1302507 | Small Intestine |
| GSE279105 | Homo sapiens | Jan 01 2026 | PRJNA1170872 | Lung; Pleural effusion |
| GSE178179 | Homo sapiens;synthetic construc | Aug 31 2021 | PRJNA737515 | NA |
| GSE293322 | Homo sapiens | Feb 10 2026 | PRJNA1244020 | pancreas |
| GSE288756 | Mus musculus | Jan 07 2026 | PRJNA1219199 | bone marrow |
| GSE202018 | Homo sapiens | Jul 03 2022 | PRJNA834049 | |
| GSE223123 | Mus musculus | Feb 26 2026 | PRJNA925083 | Bone marrow |
| GSE132828 | Homo sapiens | Nov 25 2020 | PRJNA549242 | |
| GSE314790 | Homo sapiens | Jan 04 2026 | PRJNA1392606 | Cerebral Cortex |
| GSE316455 | Homo sapiens | Jan 21 2026 | PRJNA1402944 | Liver |
| GSE312134 | Mus musculus | Jan 10 2026 | PRJNA1372228 | |
| GSE317434 | Homo sapiens | Feb 09 2026 | PRJNA1412764 | |
| GSE301514 | Homo sapiens | Jan 29 2026 | | bone |
| GSE167573 | Homo sapiens | Jul 24 2021 | PRJNA704974 | Renal cell carcinoma |
| GSE163770 | Mus musculus | Jan 31 2021 | PRJNA687498 | Neural retina |
| GSE316337 | Homo sapiens | Jan 16 2026 | PRJNA1402332 | blood |
| GSE316069 | Homo sapiens | Jan 13 2026 | PRJNA1400547 | colon |
| GSE317275 | Homo sapiens | Jan 27 2026 | PRJNA1406651 | glioma |
| GSE147015 | Homo sapiens | Jul 15 2022 | PRJNA612787 | Retina |
| GSE300149 | Mus musculus | Jan 12 2026 | PRJNA1279127 | breast caner |
| GSE303090 | Homo sapiens | Jan 28 2026 | PRJNA1294139 | |
| GSE296701 | Homo sapiens | Jan 09 2026 | | |
| GSE280621 | Mus musculus | Jan 10 2026 | PRJNA1179503 | Liver |
| GSE298693 | Homo sapiens | Feb 15 2026 | PRJNA1270955 | |
| GSE278725 | Homo sapiens | Feb 17 2026 | PRJNA1168517 | |
| GSE291855 | Mus musculus | Feb 12 2026 | PRJNA1235636 | heart scar tissue |
| GSE317999 | Mus musculus | Jan 30 2026 | PRJNA1416292 | hippocampus microglia |
| GSE243714 | Mus musculus | Jan 12 2026 | PRJNA1019682 | |

| | | | | |
|---|---|---|---|---|
| GSE197500 | Mus musculus | Nov 15 2022 | PRJNA810714 | |
| GSE316223 | Danio rerio | Jan 17 2026 | PRJNA1401831 | Brain |
| GSE300303 | Mus musculus | Feb 05 2026 | PRJNA1280071 | |
| GSE231476 | Homo sapiens | Jan 29 2026 | PRJNA966120 | |
| GSE304891 | Homo sapiens | Jan 23 2026 | PRJNA1303183 | PBMC |
| GSE287942 | Homo sapiens | Jan 05 2026 | | blood |
| GSE245816 | Homo sapiens | Dec 11 2023 | PRJNA1030019 | Astrocytes |
| GSE267165 | Mus musculus | Feb 23 2026 | PRJNA1109993 | Lung |
| GSE310809 | Homo sapiens | Apr 30 2026 | PRJNA1367103 | |
| GSE266379 | Rattus norvegicus | May 10 2024 | PRJNA1106852 | Neuroretina |
| GSE262465 | Mus musculus | Jan 27 2026 | PRJNA1091932 | Spleen |
| GSE280616 | Homo sapiens | Feb 11 2026 | PRJNA1179497 | Cell Line |
| GSE309277 | Homo sapiens | Feb 13 2026 | PRJNA1335086 | |
| GSE305020 | Homo sapiens | Jan 20 2026 | PRJNA1303720 | |
| GSE281054 | Drosophila melanogaster | Jan 27 2026 | PRJNA1181894 | Outer photoreceptor cells |
| GSE230049 | Mus musculus | Feb 16 2025 | PRJNA957276 | Retina |
| GSE275307 | Mus musculus | Feb 07 2026 | PRJNA1150226 | pineal gland |
| GSE243100 | Mus musculus | Jan 01 2024 | PRJNA1016358 | Retina |
| GSE300520 | Homo sapiens | Feb 20 2026 | PRJNA1281100 | adipose tissue |
| GSE304041 | Homo sapiens | Jun 11 2026 | PRJNA1298814 | |
| GSE238260 | Mus musculus | Jan 06 2026 | PRJNA998475 | Ovary |
| GSE284729 | Danio rerio | Jun 02 2025 | PRJNA1200128 | retina |
| GSE317461 | Homo sapiens | Feb 06 2026 | PRJNA1413099 | Thyroid |
| GSE133492 | Mus musculus | Nov 19 2020 | PRJNA551612 | etinal ganglion cells (RGC) |
| GSE266072 | Homo sapiens | Sep 24 2024 | PRJNA1105633 | retina |
| GSE277797 | Homo sapiens | Jan 26 2026 | PRJNA1164211 | Nasal mucosa |
| GSE316355 | Histoplasma ohiense | Jan 26 2026 | PRJNA1402366 | |
| GSE277288 | Mus musculus | Jan 13 2026 | PRJNA1161656 | |
| GSE291914 | Mus musculus | Jan 16 2026 | PRJNA1235764 | Spleen |
| GSE165827 | Mus musculus | Dec 22 2021 | PRJNA697981 | retina |
| GSE220587 | Mus musculus | Dec 09 2022 | PRJNA910477 | Mouse Retina |
| GSE290299 | Pseudomonas aeruginosa PA14 | Feb 13 2026 | PRJNA1227193 | |
| GSE313428 | Mus musculus | Jan 15 2026 | PRJNA1379496 | Bone marrow |
| GSE317334 | Mus musculus | Jan 27 2026 | | brain |
| GSE316128 | Mus musculus | Jan 16 2026 | PRJNA1401583 | liver |
| GSE289676 | Homo sapiens | Feb 18 2026 | PRJNA1223922 | |
| GSE203138 | Mus musculus | May 19 2022 | PRJNA838637 | |
| GSE189416 | Danio rerio | Sep 07 2022 | PRJNA782975 | whole larvae |
| GSE304155 | Mus musculus | Feb 13 2026 | PRJNA1299265 | tumor |
| GSE324272 | Mus musculus | Mar 31 2026 | PRJNA1434345 | Embryonic retina |
| GSE317708 | Mus musculus | Feb 06 2026 | PRJNA1414946 | islet |
| GSE287917 | Mus musculus | Jan 23 2026 | PRJNA1215276 | Anterior hypothalamus |
| GSE301337 | Mus musculus | Jan 01 2026 | PRJNA1284817 | peritoneal macrophage |
| GSE304637 | Homo sapiens | Jan 01 2026 | PRJNA1301960 | embryonic stem cells |
| GSE297548 | Rattus norvegicus | Jan 11 2026 | PRJNA1265173 | mammary tumor |
| GSE316523 | Homo sapiens | Feb 18 2026 | PRJNA1403059 | Cell line |
| GSE202471 | Homo sapiens | Sep 09 2022 | PRJNA836269 | Adult Retina |
| GSE212188 | Mus musculus | Nov 30 2022 | PRJNA874436 | bulk retina |
| GSE152928 | Mus musculus | Feb 26 2021 | PRJNA641006 | retina |
| GSE207171 | Mus musculus | Jan 17 2024 | PRJNA854057 | Retina |
| GSE231376 | Mus musculus | May 05 2023 | PRJNA964700 | retinal |
| GSE216694 | Mus musculus | Jun 18 2024 | PRJNA894996 | mouse retina |
| GSE290861 | Saccharomyces cerevisiae | Jan 21 2026 | PRJNA1230277 | |
| GSE315687 | Mus musculus | Jan 08 2026 | PRJNA1398794 | Lung |
| GSE285109 | Mus musculus | Feb 10 2026 | PRJNA1201189 | bone marrow |
| GSE316241 | Mus musculus | Feb 24 2026 | PRJNA1401861 | Distal lung |
| GSE292235 | Mus musculus | Jan 01 2026 | PRJNA1237697 | forebrains |
| GSE224863 | Mus musculus | Apr 18 2023 | PRJNA932810 | |
| GSE305661 | Homo sapiens | Jan 14 2026 | PRJNA1307435 | |
| GSE263466 | Sus scrofa | Jan 25 2026 | PRJNA1097595 | dorsal root ganglion |
| GSE313741 | Homo sapiens | Feb 15 2026 | PRJNA1381513 | Xenographt tumor |
| GSE309129 | Homo sapiens | Feb 24 2026 | PRJNA1334190 | |
| GSE168129 | Mus musculus | Sep 10 2022 | PRJNA706189 | |

| | | | | |
|---|---|---|---|---|
| GSE289972 | Lumbricus rubellus | Jan 01 2026 | PRJNA1225672 | izzard, Nerve cord, Body wall, Seminal vesicles, Clitellum, Pharynx |
| GSE222389 | Homo sapiens | Jan 12 2023 | PRJNA921859 | retinal epithelium |
| GSE144059 | Caenorhabditis elegans | Jan 23 2020 | PRJNA602671 | |
| GSE226495 | Mus musculus | Sep 01 2023 | PRJNA940437 | brain |
| GSE298064 | Mus musculus | Jan 15 2026 | PRJNA1268223 | blood |
| GSE310758 | Hevea brasiliensis | Jan 05 2026 | PRJNA1367034 | leaf |
| GSE280947 | Mus musculus | Jan 20 2026 | PRJNA1181529 | Pancreas |
| GSE124252 | Homo sapiens | Jan 21 2021 | PRJNA511410 | retina |
| GSE270817 | Mus musculus | Jan 09 2026 | PRJNA1128568 | |
| GSE285958 | Homo sapiens | Jan 31 2026 | PRJNA1207055 | |
| GSE253667 | Helicoverpa armigera | Jan 14 2026 | PRJNA1066738 | wing disc |
| GSE319038 | Homo sapiens | Feb 18 2026 | PRJNA1422002 | conjuctiva |
| GSE315770 | Caenorhabditis elegans | Jan 07 2026 | | whole animal |
| GSE174843 | evere acute respiratory syndrom | Jan 01 2022 | PRJNA731890 | erived from Gibco episomal iPSC line A18945 |
| GSE264354 | llus;Homo sapiens;Pimephales | Feb 25 2026 | PRJNA1102016 | |
| GSE282446 | Rattus norvegicus | Nov 24 2024 | PRJNA1188594 | Optic Nerve Head |
| GSE104620 | Rattus norvegicus | Oct 01 2022 | PRJNA413308 | Retina |
| GSE240512 | Mus musculus | Feb 23 2026 | PRJNA1003949 | |
| GSE175822 | Mus musculus | Feb 13 2026 | PRJNA733896 | Calvaria |
| GSE191206 | Homo sapiens | Dec 25 2021 | PRJNA790451 | |
| GSE150794 | Mus musculus | May 18 2023 | PRJNA633594 | Retina |
| GSE241211 | Mus musculus | Jan 20 2026 | PRJNA1006731 | Cortex |
| GSE315983 | Caenorhabditis elegans | Jan 09 2026 | PRJNA1400057 | whole organism |
| GSE315653 | Gallus gallus;Homo sapiens | Jan 05 2026 | PRJNA1398405 | cell renal cell carcinoma_tumor |
| GSE317918 | Homo sapiens | Jan 31 2026 | PRJNA1415815 | n/a |
| GSE271791 | Mus musculus | Jul 09 2024 | PRJNA1133187 | PE/choroid/sclera complex |
| GSE294327 | Cucumis sativus | Jan 19 2026 | PRJNA1249032 | shoot apical meristem |
| GSE120338 | Homo sapiens | Mar 22 2021 | PRJNA492791 | y retinal pigment epithelial cells |
| GSE315131 | Mus musculus | Jan 31 2026 | PRJNA1395042 | subcutaneous tumor |
| GSE291145 | Homo sapiens | Mar 31 2026 | PRJNA1232088 | |
| GSE290646 | Homo sapiens | Jan 23 2026 | PRJNA1229476 | |
| GSE292583 | Homo sapiens | Jan 28 2026 | PRJNA1240256 | |
| GSE298612 | Mus musculus | Jan 22 2026 | PRJNA1270025 | incisor mesenchyme |
| GSE290671 | Macaca fascicularis | Jan 06 2026 | PRJNA1229513 | Blood |
| GSE197938 | Gallus gallus | May 11 2022 | PRJNA812904 | |
| GSE311467 | Mus musculus | Feb 20 2026 | PRJNA1369509 | Hippocampus |
| GSE300382 | Mus musculus | Jan 14 2026 | PRJNA1280175 | |
| GSE270934 | Mus musculus | Oct 16 2024 | PRJNA1129064 | Optic nerve head |
| GSE188280 | Homo sapiens | May 25 2022 | PRJNA778199 | |
| GSE201677 | Homo sapiens | Apr 30 2022 | PRJNA832665 | |
| GSE272221 | Homo sapiens | Jan 13 2026 | PRJNA1135884 | |
| GSE297697 | Homo sapiens | Feb 28 2026 | PRJNA1265723 | skin |
| GSE312871 | Mus musculus | Jan 08 2026 | PRJNA1377544 | spinal cord |
| GSE289910 | Homo sapiens | Feb 28 2026 | PRJNA1225282 | |
| GSE288782 | Homo sapiens | Feb 05 2026 | PRJNA1219396 | brain |
| GSE305930 | Mus musculus | Feb 18 2026 | PRJNA1308468 | brain |
| GSE223851 | Rattus norvegicus | Jan 01 2026 | PRJNA928672 | Cortical blood vessels |
| GSE307353 | Mus musculus | Feb 24 2026 | PRJNA1316976 | lung lavage |
| GSE147844 | Mus musculus | Apr 20 2020 | PRJNA622266 | Retina |
| GSE200621 | Mus musculus | Apr 16 2022 | PRJNA825553 | retina |
| GSE304795 | Homo sapiens | Jan 22 2026 | PRJNA1302761 | Primary T cell |
| GSE319738 | Escherichia coli | Feb 20 2026 | PRJNA1425035 | |
| GSE313827 | Homo sapiens | Feb 20 2026 | PRJNA1381646 | |
| GSE297853 | Macaca mulatta | Feb 13 2026 | PRJNA1266488 | Brain |
| GSE307431 | Homo sapiens | Jan 06 2026 | PRJNA1321660 | Skin |
| GSE203585 | Gallus gallus | May 19 2025 | PRJNA841502 | |
| GSE244980 | Homo sapiens | Feb 25 2026 | PRJNA1026555 | whole blood |
| GSE318246 | Sus scrofa | Feb 04 2026 | PRJNA1416961 | Retina |
| GSE303632 | omys tridecemlineatus;Mus mus | Jul 28 2025 | PRJNA1295948 | P5 retina |
| GSE178928 | Mus musculus | Jun 26 2021 | PRJNA741503 | |
| GSE261077 | Homo sapiens | Feb 28 2026 | PRJNA1085243 | Cell line |
| GSE317789 | Homo sapiens | Feb 06 2026 | PRJNA1415256 | human skin |
| GSE225125 | Homo sapiens | Jan 05 2026 | PRJNA934061 | |

| | | | | |
|---|---|---|---|---|
| GSE237206 | Callithrix jacchus | Oct 01 2023 | PRJNA994171 | Retina |
| GSE273354 | Homo sapiens | Jan 23 2026 | PRJNA1141477 | |
| GSE287007 | Mus musculus | Jan 24 2026 | PRJNA1210684 | Lymph nodes, Spleen |
| GSE164208 | Homo sapiens | Jan 09 2021 | PRJNA689555 | |
| GSE242796 | Gallus gallus | Sep 14 2023 | PRJNA1014752 | retina |
| GSE314132 | Homo sapiens | Feb 20 2026 | PRJNA1390140 | |
| GSE186206 | Homo sapiens | Oct 20 2021 | PRJNA772958 | |
| GSE308383 | Mus musculus | Dec 16 2025 | PRJNA1330319 | Myenteric layer of Ileum |
| GSE290459 | Mus musculus | Jan 15 2026 | PRJNA1228073 | Peritoneal cavity |
| GSE309480 | Homo sapiens | Feb 09 2026 | PRJNA1336138 | stem cell |
| GSE124324 | Mus musculus | Jan 01 2020 | PRJNA511608 | |
| GSE162184 | Homo sapiens | Jun 17 2021 | PRJNA680800 | |
| GSE309839 | Homo sapiens | Jan 31 2026 | PRJNA1363876 | adrenal mass |
| GSE304872 | Homo sapiens | Feb 19 2026 | PRJNA1303116 | |
| GSE128144 | Homo sapiens | Apr 22 2020 | PRJNA526531 | nal Pigmented Epithelial Cells |
| GSE233750 | Mus musculus | Sep 30 2023 | PRJNA977697 | retina |
| GSE306544 | Mus musculus | Jan 16 2026 | PRJNA1311095 | ovary P1 |
| GSE319333 | Homo sapiens | Feb 16 2026 | PRJNA1423034 | |
| GSE219084 | Mus musculus | Jan 11 2023 | PRJNA907000 | neuroretinal homogenate |
| GSE295191 | Mus musculus | Feb 12 2026 | | splenocytes |
| GSE309448 | Candida tropicalis | Feb 23 2026 | | |
| GSE227055 | Mus musculus | May 19 2023 | PRJNA942967 | retina |
| GSE304403 | agaria x ananassa subsp. ananas | Jan 31 2026 | PRJNA1301257 | Leaves |
| GSE300162 | Mus musculus | Feb 10 2026 | PRJNA1279156 | liver |
| GSE288341 | Homo sapiens | Jan 17 2026 | PRJNA1216966 | |
| GSE185475 | Homo sapiens | Sep 15 2025 | PRJNA769139 | |
| GSE310184 | Mus musculus | Jan 12 2026 | PRJNA1365245 | |
| GSE312241 | Homo sapiens | Jan 25 2026 | | |
| GSE315484 | Oncorhynchus mykiss | Jan 13 2026 | PRJNA1397221 | Visceral adipose tissue |
| GSE247776 | Mus musculus | May 23 2024 | PRJNA1040300 | before and after the ora serrata |
| GSE296202 | Homo sapiens | Jan 14 2026 | PRJNA1258202 | |
| GSE148387 | Homo sapiens | Aug 30 2020 | PRJNA624120 | Conjunctival melanoma |
| GSE262151 | Homo sapiens | Mar 25 2025 | PRJNA1090480 | RPE/choroid |
| GSE180142 | Mus musculus | Jul 15 2024 | PRJNA746767 | Retina |
| GSE291849 | onas macleodii;Thalassiosira pse | Feb 10 2026 | PRJNA1235624 | |
| GSE298390 | Homo sapiens | Jan 14 2026 | PRJNA1269061 | |
| GSE302290 | Mus musculus | Jan 05 2026 | PRJNA1289816 | of the embryonic small intestine (E15.5) |
| GSE260978 | Mus musculus | Feb 14 2025 | PRJNA1084767 | Optic Nerve |
| GSE286099 | Mus musculus | Jan 12 2026 | PRJNA1207923 | |
| GSE318685 | Mus musculus | Feb 27 2026 | PRJNA1419777 | |
| GSE300637 | Mus musculus | Feb 13 2026 | | Thymus |
| GSE152842 | Danio rerio | Nov 30 2020 | PRJNA640615 | retina |
| GSE313149 | Mus musculus | Jan 26 2026 | PRJNA1378702 | Lung |
| GSE315590 | Mus musculus | Jan 30 2026 | PRJNA1398302 | heart |
| GSE316449 | Homo sapiens | Jan 16 2026 | PRJNA1402931 | periodontal ligament |
| GSE217280 | Glycine max | Jan 31 2026 | PRJNA898330 | |
| GSE179104 | Danio rerio | Apr 05 2022 | PRJNA742218 | kidney |
| GSE307840 | Homo sapiens | Jan 23 2026 | PRJNA1327402 | liver |
| GSE193323 | Mus musculus | Jan 29 2026 | PRJNA795792 | |
| GSE254008 | Mus musculus | Dec 31 2025 | PRJNA1068106 | retina |
| GSE290585 | Mus musculus | Sep 26 2025 | PRJNA1228665 | Liver |
| GSE317100 | Mus musculus | Feb 16 2026 | PRJNA1405778 | |
| GSE271307 | Homo sapiens | Jan 14 2026 | PRJNA1130899 | Human breast |
| GSE226408 | Homo sapiens | Mar 15 2023 | PRJNA940148 | |
| GSE306816 | Homo sapiens | Feb 26 2026 | PRJNA1312449 | |
| GSE289561 | Mus musculus | Feb 14 2026 | PRJNA1223435 | liver |
| GSE285229 | Mus musculus | Feb 17 2026 | PRJNA1202078 | Colon |
| GSE312362 | Homo sapiens | Jan 18 2026 | PRJNA1373180 | |
| GSE253809 | Homo sapiens | Jan 29 2024 | PRJNA1067156 | Pigmented epithelium; Retina |
| GSE256187 | Homo sapiens | Feb 22 2026 | PRJNA1078385 | Colorectal cancer |
| GSE292884 | Mus musculus | Feb 23 2026 | PRJNA1241811 | Liver |
| GSE301603 | Mus musculus | Jan 19 2026 | PRJNA1285710 | stool |
| GSE180117 | Mus musculus | Nov 11 2021 | PRJNA746584 | |

| | | | | |
|---|---|---|---|---|
| GSE294803 | Mus musculus | Feb 18 2026 | PRJNA1251627 | Heart, ventricle |
| GSE308154 | Homo sapiens | Jan 04 2026 | PRJNA1329051 | |
| GSE307936 | Homo sapiens | Feb 14 2026 | PRJNA1327845 | |
| GSE317041 | Rattus norvegicus | Jan 27 2026 | PRJNA1405723 | Intestine |
| GSE288441 | Nematostella vectensis | Jan 12 2026 | PRJNA1217498 | ody wall with mesenteries |
| GSE285884 | Mus musculus | Jan 15 2026 | PRJNA1206706 | intestine |
| GSE244809 | Mus musculus | Aug 13 2024 | PRJNA1024938 | Eye |
| GSE297236 | Homo sapiens | Feb 10 2026 | PRJNA1263375 | |
| GSE221860 | Homo sapiens | Dec 29 2022 | PRJNA916510 | retina |
| GSE284621 | Mus musculus | Jan 05 2026 | PRJNA1199947 | Mesenteric Lymph Node |
| GSE312069 | Mus musculus | Jan 13 2026 | PRJNA1372142 | Cortex |
| GSE275072 | Mus musculus | Sep 15 2025 | PRJNA1149053 | Retina |
| GSE306960 | Mus musculus | Sep 04 2025 | PRJNA1314045 | retina |
| GSE308163 | Mus musculus | Jan 28 2026 | PRJNA1329126 | prostate |
| GSE316508 | Marchantia polymorpha | Jan 21 2026 | PRJNA1403027 | rchantia polymorpha Thallus |
| GSE280340 | Mus musculus | Jan 09 2026 | PRJNA1177878 | Liver |
| GSE311093 | Mus musculus | Feb 05 2026 | PRJNA1368664 | heart |
| GSE289256 | Homo sapiens | Feb 06 2026 | PRJNA1222271 | lung tumor |
| GSE312455 | Saccharomyces cerevisiae | Jan 29 2026 | PRJNA1373339 | |
| GSE301075 | Homo sapiens | Feb 18 2026 | PRJNA1283221 | primary pancreas tumor |
| GSE223320 | Mus musculus | Jan 20 2026 | PRJNA925802 | |
| GSE255514 | Mus musculus | Jan 26 2026 | PRJNA1075139 | |
| GSE308040 | Homo sapiens | Feb 26 2026 | PRJNA1328619 | B-cell lymphoma |
| GSE292140 | Homo sapiens | Jan 26 2026 | PRJNA1237270 | Primary Tumour |
| GSE274107 | Mus musculus | Feb 26 2025 | PRJNA1144968 | retina |
| GSE301720 | Homo sapiens | Feb 16 2026 | | Oropharynx |
| GSE181737 | Homo sapiens | Sep 14 2023 | PRJNA753130 | |
| GSE318900 | Escherichia coli DH5[alpha] | Feb 16 2026 | PRJNA1420791 | |
| GSE275604 | Mus musculus | Jan 01 2026 | PRJNA1152013 | Lung |
| GSE297477 | Homo sapiens | Jan 21 2026 | PRJNA1264970 | |
| GSE204806 | Homo sapiens | Sep 01 2022 | PRJNA842381 | |
| GSE189763 | Mus musculus | Feb 12 2026 | PRJNA784431 | |
| GSE325479 | Danio rerio;Mus musculus | Mar 31 2026 | PRJNA1440247 | retina |
| GSE285036 | Homo sapiens | Feb 11 2026 | PRJNA1201080 | Differentiating cells |
| GSE178652 | Homo sapiens | Jan 10 2026 | PRJNA739894 | |
| GSE178121 | Mus musculus | Aug 24 2021 | PRJNA737414 | retina |
| GSE319783 | Mus musculus | Feb 17 2026 | PRJNA1425088 | marrow |
| GSE274409 | Homo sapiens | Jan 20 2026 | PRJNA1146367 | Breast |
| GSE316112 | Homo sapiens | Jan 14 2026 | PRJNA1400622 | |
| GSE318080 | Homo sapiens | Feb 15 2026 | PRJNA1416428 | Tumor - Lung |
| GSE251731 | Rattus norvegicus | Nov 27 2024 | PRJNA1055140 | retina |
| GSE221952 | Mus musculus | Aug 08 2024 | PRJNA916923 | posterior of eye |
| GSE256389 | Mus musculus | Feb 16 2025 | PRJNA1079322 | Retina |
| GSE134745 | Homo sapiens | Jul 23 2020 | PRJNA556254 | |
| GSE247173 | Mus musculus | Jan 01 2025 | PRJNA1036568 | Eyes |
| GSE288573 | Mus musculus | Jan 03 2026 | | Eye |
| GSE309030 | Mus musculus | Jan 24 2026 | PRJNA1333816 | Bone Marrow |
| GSE311913 | ethanothermobacter marburgen | Jan 06 2026 | | |
| GSE300278 | Homo sapiens | Feb 25 2026 | PRJNA1280016 | skin |
| GSE223331 | Homo sapiens | Jan 12 2026 | PRJNA925825 | glioblastoma tumor |
| GSE317144 | Mus musculus | Feb 07 2026 | PRJNA1406168 | ovary |
| GSE127352 | Homo sapiens | Feb 28 2020 | PRJNA524714 | |
| GSE296352 | Mus musculus | Feb 10 2026 | PRJNA1258975 | |
| GSE312326 | Homo sapiens | Feb 27 2026 | PRJNA1373032 | Colorectal |
| GSE138247 | Mus musculus | Jan 08 2020 | PRJNA575217 | Retinal Microglia |
| GSE185512 | Homo sapiens | Oct 11 2021 | PRJNA769402 | |
| GSE163562 | Rattus norvegicus | Jul 28 2021 | PRJNA686776 | retina |
| GSE255505 | Homo sapiens | Feb 08 2026 | PRJNA1075125 | |
| GSE293013 | Homo sapiens | Jan 08 2026 | PRJNA1242347 | |
| GSE188607 | Mus musculus | Dec 02 2022 | PRJNA779630 | retina |
| GSE288178 | Homo sapiens;Mus musculus | Feb 22 2026 | PRJNA1216533 | adult mouse brain |
| GSE293982 | Homo sapiens | Nov 18 2025 | PRJNA1247529 | Retinal organoids |
| GSE297210 | Homo sapiens | Feb 17 2026 | PRJNA1263010 | Brain |

| | | | | |
|---|---|---|---|---|
| GSE223570 | Homo sapiens | Feb 16 2023 | PRJNA926900 | |
| GSE233596 | Homo sapiens | Jan 28 2026 | PRJNA976721 | blood |
| GSE270201 | Mus musculus | Dec 31 2024 | PRJNA1125395 | Choroid |
| GSE202154 | Mus musculus | Jun 28 2022 | PRJNA834807 | Retina |
| GSE211574 | Homo sapiens;Mus musculus | Jul 11 2023 | PRJNA870918 | embryonic stem cell |
| GSE282932 | Homo sapiens | Jan 31 2026 | PRJNA1191027 | PBMC |
| GSE306946 | Homo sapiens | Feb 13 2026 | | retina |
| GSE292045 | Homo sapiens | Feb 10 2026 | PRJNA1236374 | glioblastoma tumor |
| GSE290380 | Homo sapiens | Feb 23 2026 | PRJNA1227692 | |
| GSE190682 | Homo sapiens | Jul 01 2022 | PRJNA787955 | |
| GSE223225 | Homo sapiens | Jan 17 2026 | PRJNA925429 | |
| GSE317050 | Mus musculus | Jan 30 2026 | PRJNA1405709 | heart |
| GSE242411 | Mus musculus | Feb 20 2026 | PRJNA1013185 | |
| GSE142671 | Mus musculus | Jan 14 2020 | PRJNA597893 | Retina |
| GSE307866 | Homo sapiens | Feb 16 2026 | PRJNA1327429 | pons |
| GSE241946 | Mus musculus | Aug 03 2024 | PRJNA1010790 | containing retina, lens, and blood |
| GSE252879 | Mus musculus | Jan 08 2026 | PRJNA1062862 | heart |
| GSE163255 | Rattus norvegicus | Aug 26 2021 | PRJNA685424 | neuroretina |
| GSE317880 | Homo sapiens | Feb 12 2026 | PRJNA1415768 | |
| GSE304253 | Homo sapiens | Feb 27 2026 | PRJNA1299494 | Blood cells |
| GSE293607 | Mus musculus | Apr 02 2026 | PRJNA1245366 | relaxed ciliary muscle |
| GSE302395 | Mus musculus | Feb 23 2026 | PRJNA1290320 | retina |
| GSE308004 | Mus musculus | Jan 13 2026 | PRJNA1328499 | testis |
| GSE305550 | Homo sapiens | Sep 01 2025 | PRJNA1306489 | retinal organoid |
| GSE298295 | Mus musculus | Jan 21 2026 | PRJNA1268902 | thymus |
| GSE310927 | Homo sapiens | Feb 26 2026 | PRJNA1367546 | |
| GSE289547 | Mus musculus | Mar 11 2025 | PRJNA1223411 | retina |
| GSE222332 | Homo sapiens | Jan 02 2026 | PRJNA919029 | |
| GSE241782 | Mus musculus | Sep 06 2023 | PRJNA1009858 | UON |
| GSE315581 | Mus musculus | Jan 10 2026 | PRJNA1398285 | guinal white adipose tissue |
| GSE316458 | Mus musculus | Jan 16 2026 | PRJNA1402957 | Pancreas |
| GSE263002 | Homo sapiens | Feb 15 2026 | PRJNA1095435 | Cell line |
| GSE251997 | Homo sapiens | Dec 30 2025 | PRJNA1057047 | |
| GSE294288 | Mus musculus | Jan 01 2026 | PRJNA1248943 | |
| GSE284130 | Petromyzon marinus | Aug 18 2025 | PRJNA1197416 | retina |
| GSE300144 | Plasmodium vivax | Feb 12 2026 | PRJNA1279129 | Enriched blood sample |
| GSE318867 | Homo sapiens | Feb 15 2026 | PRJNA1420747 | SCLC |
| GSE306082 | Mus musculus | Aug 28 2025 | PRJNA1309089 | retina |
| GSE223512 | Homo sapiens | Sep 18 2024 | PRJNA926621 | breast |
| GSE229683 | Homo sapiens | Nov 03 2023 | PRJNA955277 | Retinal organoid |
| GSE179754 | Mus musculus | Jul 10 2021 | PRJNA744905 | retinal explant |
| GSE80571 | Arabidopsis thaliana | Jan 31 2026 | PRJNA319328 | seedlings |
| GSE283588 | Danio rerio | Jan 31 2025 | PRJNA1194578 | eye |
| GSE317032 | Mus musculus | Feb 25 2026 | PRJNA1405684 | Spleen |
| GSE288432 | Mus musculus | Jan 30 2026 | PRJNA1217466 | bone marrow |
| GSE311865 | Oryza sativa | Jan 26 2026 | PRJNA1371864 | 14-day-old seedlings |
| GSE316731 | Mus musculus | Jan 20 2026 | PRJNA1404098 | soleus muscles |
| GSE301883 | Homo sapiens | Jan 24 2026 | PRJNA1287687 | uman embryonic stem cell |
| GSE266898 | Oryza sativa | Jan 26 2026 | PRJNA1108694 | 14-day-old seedlings |
| GSE305311 | Drosophila melanogaster | Feb 16 2026 | | larva VNC MN1-Ib motoneurons |
| GSE281118 | Mus musculus | Jan 12 2026 | PRJNA1182318 | prostate |
| GSE306924 | Mus musculus | Jan 07 2026 | PRJNA1313932 | Substantia Nigra |
| GSE310804 | Mus musculus | Apr 30 2026 | PRJNA1367089 | RPE |
| GSE267168 | Homo sapiens | Feb 23 2026 | PRJNA1109990 | primary breast cancer |
| GSE318850 | Homo sapiens | Feb 13 2026 | PRJNA1420725 | MetLN |
| GSE248850 | Mus musculus | Feb 12 2026 | PRJNA1046108 | skin (face) |
| GSE278200 | Rattus norvegicus | Feb 12 2026 | PRJNA1166073 | tal Right Lung Homogenate |
| GSE290160 | Arabidopsis thaliana | Jan 29 2026 | PRJNA1226489 | Seedlings |
| GSE315308 | Arabidopsis thaliana | Feb 12 2026 | PRJNA1395612 | Whole seedling |
| GSE298134 | Mus musculus | Jan 21 2026 | PRJNA1268412 | Bone marrow |
| GSE315617 | Mus musculus | Jan 18 2026 | PRJNA1398334 | Spleen |
| GSE291663 | Mus musculus | Mar 02 2026 | PRJNA1234620 | retina explant |
| GSE294363 | Homo sapiens | Jan 06 2026 | | umbilical vein |

| | | | | |
|---|---|---|---|---|
| GSE314676 | Mus musculus | Feb 25 2026 | PRJNA1392436 | Brain |
| GSE302756 | Homo sapiens | Feb 25 2026 | PRJNA1292110 | kidney |
| GSE300493 | erichia coli str. K-12 substr. MG1 | Feb 28 2026 | PRJNA1281062 | |
| GSE304690 | Homo sapiens | Aug 11 2025 | PRJNA1302330 | cell line |
| GSE302249 | Mus musculus | Jan 26 2026 | PRJNA1289718 | |
| GSE310475 | Homo sapiens | Jan 26 2026 | PRJNA1366403 | lung |
| GSE224600 | Mus musculus | May 01 2023 | PRJNA931936 | Retina |
| GSE315142 | Mycolicibacterium smegmatis | Jan 01 2026 | PRJNA1395048 | cell |
| GSE313915 | Mus musculus | Feb 21 2026 | PRJNA1387237 | Mice prostate tissue |
| GSE288687 | Mus musculus | Jan 06 2026 | PRJNA1219070 | Lungs |
| GSE282516 | Mus musculus | Feb 06 2026 | PRJNA1189074 | e marrow from long bones and spine |
| GSE312298 | Mus musculus | Jan 01 2026 | PRJNA1372801 | Bone marrow |
| GSE287509 | Homo sapiens | Jan 16 2026 | PRJNA1213116 | organoid |
| GSE224328 | Homo sapiens | Feb 02 2024 | PRJNA930654 | |
| GSE291654 | Mus musculus | Aug 20 2025 | PRJNA1234598 | retina |
| GSE256200 | Cavia porcellus | Jun 26 2026 | PRJNA1078402 | Choroid |
| GSE155828 | Homo sapiens | Jan 08 2021 | PRJNA655721 | |
| GSE302003 | Homo sapiens | Jan 01 2026 | PRJNA1288268 | |
| GSE319477 | Hyalomma asiaticum | Feb 18 2026 | | Ticks as a whole |
| GSE311423 | Homo sapiens | Jan 01 2026 | PRJNA1369441 | Spleen |
| GSE276438 | Oryctolagus cuniculus | Sep 05 2024 | PRJNA1157360 | Retina |
| GSE288874 | Homo sapiens | Feb 05 2026 | PRJNA1219660 | |
| GSE262583 | Homo sapiens | Jan 01 2026 | PRJNA1092594 | cell line |
| GSE286329 | Mus musculus | Jan 20 2026 | PRJNA1208928 | Brain |
| GSE149549 | Mus musculus | Aug 24 2020 | PRJNA629051 | eyeball |
| GSE159921 | Homo sapiens | Jan 16 2026 | PRJNA670898 | |
| GSE302642 | Homo sapiens | Jan 13 2026 | PRJNA1291638 | |
| GSE310203 | Homo sapiens | Feb 09 2026 | PRJNA1365269 | |
| GSE270853 | Homo sapiens | Jan 16 2026 | PRJNA1128635 | A Cell Line injected in mice |
| GSE279695 | Homo sapiens | Jan 29 2026 | PRJNA1174100 | hiPSC-CM tissue |
| GSE298020 | Homo sapiens | Jan 29 2026 | PRJNA1267114 | |
| GSE291268 | Homo sapiens;Mus musculus | Jan 28 2026 | PRJNA1232693 | Left Ventricle |
| GSE290074 | Homo sapiens | Jan 19 2026 | PRJNA1226150 | Ovary |
| GSE155585 | Mus musculus | Jan 12 2026 | PRJNA650403 | inal center B cells from spleens |
| GSE288960 | Homo sapiens | Jan 29 2026 | PRJNA1220103 | cell line |
| GSE283463 | Mus musculus | Sep 01 2025 | PRJNA1194001 | Retina |
| GSE307417 | Mus musculus | Jan 22 2026 | PRJNA1321587 | Kidney |
| GSE290043 | Mus musculus | Jan 21 2026 | PRJNA1226090 | heart |
| GSE264310 | Danio rerio | Oct 21 2024 | PRJNA1101946 | head |
| GSE217746 | Homo sapiens | Feb 08 2023 | PRJNA900252 | tinal microvascular endothelial cell |
| GSE308799 | Mus musculus | Jan 09 2026 | PRJNA1332731 | Regenerating skin wound |
| GSE241255 | Mus musculus | Feb 20 2026 | PRJNA1007592 | |
| GSE299569 | Homo sapiens;Mus musculus | Feb 25 2026 | PRJNA1274998 | colon |
| GSE193498 | Mus musculus | Oct 09 2024 | PRJNA796364 | retina |
| GSE315549 | Homo sapiens | Jan 09 2026 | PRJNA1398244 | bone marrow |
| GSE245549 | Mus musculus | Jun 06 2024 | PRJNA1028963 | retina |
| GSE252675 | Homo sapiens | Jan 01 2026 | PRJNA1061786 | |
| GSE287893 | Mus musculus | Dec 03 2025 | PRJNA1214873 | bone marrow |
| GSE197384 | Mus musculus | Sep 23 2022 | PRJNA810059 | |
| GSE212524 | Homo sapiens;Mus musculus | Aug 18 2024 | PRJNA875892 | Retina |
| GSE195891 | Mus musculus | Jan 07 2024 | PRJNA802562 | retina |
| GSE310417 | Mus musculus | Feb 21 2026 | PRJNA1365913 | Renal pelvis |
| GSE290660 | Mus musculus | Jan 20 2026 | PRJNA1229498 | tumor tissue |
| GSE315886 | Homo sapiens | Jan 12 2026 | PRJNA1399508 | Heart |
| GSE298634 | Homo sapiens | Feb 26 2026 | PRJNA1270062 | |
| GSE311314 | Homo sapiens | Jan 06 2026 | PRJNA1368921 | astrocytes |
| GSE310375 | Mus musculus | Feb 28 2026 | PRJNA1365847 | gastrocnemius muscle |
| GSE288498 | Homo sapiens | Dec 03 2025 | PRJNA1217878 | Blood (Bone marrow) |
| GSE317098 | Homo sapiens | Feb 10 2026 | PRJNA1405775 | testis |
| GSE295435 | Homo sapiens | Feb 24 2026 | PRJNA1254674 | |
| GSE276145 | Homo sapiens | Jan 01 2026 | PRJNA1155180 | |
| GSE303991 | Danio rerio | Jan 01 2026 | PRJNA1298598 | bony fishes |
| GSE189386 | Candida albicans | Jan 12 2026 | PRJNA782880 | |

| | | | | |
|---|---|---|---|---|
| GSE303000 | Mus musculus | Feb 09 2026 | PRJNA1293335 | testis |
| GSE315120 | Rattus norvegicus | Jan 06 2026 | PRJNA1395022 | |
| GSE179188 | Homo sapiens | Jan 26 2023 | PRJNA742671 | |
| GSE290016 | Homo sapiens | Feb 28 2026 | PRJNA1225742 | |
| GSE234963 | Homo sapiens | Nov 29 2023 | PRJNA983820 | Retina |
| GSE184902 | Mus musculus | Mar 02 2022 | PRJNA767003 | retina |
| GSE294215 | Mus musculus | Feb 16 2026 | PRJNA1248565 | Skull |
| GSE266911 | s reinhardtii;Oryza sativa;Physco | Jan 26 2026 | PRJNA1108702 | 14-day-old seedlings |
| GSE300798 | Homo sapiens | Feb 12 2026 | | spinal cord |
| GSE287635 | Mus musculus | Jan 21 2026 | PRJNA1213855 | Brain |
| GSE315761 | Homo sapiens | Jan 12 2026 | PRJNA1398911 | |
| GSE282457 | Scyliorhinus canicula | Nov 25 2024 | | Retina |
| GSE294577 | Homo sapiens | Jan 07 2026 | PRJNA1250530 | |
| GSE329195 | Homo sapiens | Apr 29 2026 | PRJNA1458134 | corneal epithelial cell line |
| GSE306414 | Homo sapiens | Jan 13 2026 | PRJNA1310597 | Brain |
| GSE318560 | Homo sapiens | Feb 05 2026 | PRJNA1419230 | Hippocampus |
| GSE297499 | Homo sapiens | Jan 21 2026 | PRJNA1265093 | Cerebrospinal Fluid |
| GSE285949 | Mus musculus | Jan 07 2026 | PRJNA1207029 | |
| GSE309969 | Homo sapiens | Jan 09 2026 | PRJNA1364067 | |
| GSE275188 | Mus musculus | Feb 25 2026 | PRJNA1149757 | heart |
| GSE312193 | Homo sapiens | Feb 12 2026 | PRJNA1372642 | |
| GSE270197 | Mus musculus | Dec 31 2024 | PRJNA1125400 | Choroid |
| GSE310749 | Mus musculus | Jan 09 2026 | PRJNA1367031 | erior dorsal cerebellar vermis |
| GSE315249 | Mus musculus | Jan 04 2026 | PRJNA1395477 | |
| GSE272252 | Rattus norvegicus | Sep 04 2024 | PRJNA1135956 | Retina |
| GSE314055 | Mus musculus | Feb 20 2026 | PRJNA1389889 | lung |
| GSE315865 | Mus musculus | Jan 08 2026 | PRJNA1399451 | kidney |
| GSE287931 | Homo sapiens | Jan 29 2026 | PRJNA1215316 | bone marrow |
| GSE152939 | Homo sapiens | Feb 17 2021 | PRJNA641019 | |
| GSE262416 | Mus musculus | Jan 21 2026 | PRJNA1091684 | Testis |
| GSE309345 | Mus musculus | Feb 25 2026 | PRJNA1335210 | Brain cortical plate |
| GSE300370 | Mus musculus | Jan 12 2026 | PRJNA1280161 | Bone marrow |
| GSE224872 | Mus musculus | Feb 07 2026 | PRJNA932825 | |
| GSE315389 | Mus musculus | Jan 05 2026 | PRJNA1396646 | |
| GSE296538 | Homo sapiens | Jan 28 2026 | PRJNA1259928 | |
| GSE199858 | Mus musculus | May 02 2022 | PRJNA821837 | |
| GSE312053 | Homo sapiens | Jan 29 2026 | PRJNA1372126 | Lung |
| GSE259427 | Mus musculus | Feb 06 2026 | PRJNA1081463 | Mouse embryos |
| GSE288164 | Homo sapiens | Jan 28 2026 | PRJNA1216306 | Cartilage |
| GSE290847 | Homo sapiens | Jun 16 2025 | PRJNA1230231 | cell line |
| GSE314833 | Mus musculus | Feb 25 2026 | PRJNA1392910 | lippocampus microvessels |
| GSE302913 | Mus musculus | Jan 01 2026 | PRJNA1292810 | testis |
| GSE271990 | Homo sapiens | Aug 16 2025 | PRJNA1134251 | |
| GSE211077 | Mus musculus | Sep 15 2022 | PRJNA868778 | |
| GSE319394 | Siniperca chuatsi | Feb 16 2026 | PRJNA1423230 | |
| GSE314910 | Homo sapiens | Jan 05 2026 | PRJNA1393118 | Heart |
| GSE302830 | Homo sapiens | Jan 05 2026 | PRJNA1292235 | SKIN |
| GSE151707 | Gallus gallus | Apr 19 2021 | PRJNA637029 | |
| GSE313644 | Mus musculus | Feb 11 2026 | PRJNA1380452 | whole liver |
| GSE296651 | Mus musculus | May 26 2025 | PRJNA1260457 | Retina |
| GSE250019 | Mus musculus | Dec 13 2023 | PRJNA1051616 | retina |
| GSE316139 | Mus musculus | Feb 10 2026 | PRJNA1401706 | placenta |
| GSE315971 | vum;Triticum monococcum;Triti | Jan 08 2026 | PRJNA1400043 | Wheat grain |
| GSE253851 | Mus musculus | Jan 30 2026 | PRJNA1067575 | Spinal cord |
| GSE308552 | Saccharomyces cerevisiae | Jan 15 2026 | PRJNA1330865 | |
| GSE287825 | Homo sapiens | Feb 11 2026 | PRJNA1214812 | leukemia cells |
| GSE300988 | Homo sapiens | Feb 20 2026 | PRJNA1282955 | blood |
| GSE316426 | Homo sapiens | Feb 19 2026 | PRJNA1402887 | |
| GSE315946 | Danio rerio | Jan 13 2026 | PRJNA1399991 | Trunk tissue |
| GSE319280 | Arabidopsis thaliana | Feb 12 2026 | PRJNA1422742 | leaf |
| GSE303866 | Gallus gallus | Jan 04 2026 | PRJNA1298153 | posterior chicken end |
| GSE275316 | Homo sapiens | Nov 28 2024 | PRJNA1150243 | |
| GSE146671 | Homo | Oct 27 2020 | PRJNA611605 | creas ductal adenocarcinoma |

| | | | | |
|---|---|---|---|---|
| GSE304732 | Mus musculus | Sep 04 2025 | PRJNA1302420 | Retina |
| GSE310959 | Homo sapiens | Feb 20 2026 | PRJNA1367601 | |
| GSE225930 | Homo sapiens | Feb 22 2026 | PRJNA938059 | |
| GSE313673 | Mus musculus | Jan 14 2026 | PRJNA1380511 | Lung |
| GSE276923 | Mus musculus | Jan 15 2025 | PRJNA1159737 | retina |
| GSE307496 | Bos taurus | Feb 11 2026 | PRJNA1321880 | testis |
| GSE309504 | Bacteroides fragilis | Jan 15 2026 | PRJNA1336179 | |
| GSE288070 | Homo sapiens | Jan 24 2026 | PRJNA1215991 | Brain-derived |
| GSE138150 | Camellia sinensis | Dec 31 2020 | PRJNA574930 | root |
| GSE284679 | Mus musculus | Feb 11 2026 | PRJNA1200032 | RM-1 tumors |
| GSE309445 | Mus musculus | Oct 02 2025 | PRJNA1336094 | retina |
| GSE290812 | Mus musculus | Feb 10 2026 | PRJNA1230162 | |
| GSE317253 | Mus musculus | Feb 26 2026 | PRJNA1406626 | lung |
| GSE301373 | Homo sapiens | Oct 01 2025 | PRJNA1284870 | |
| GSE259327 | Homo sapiens | Jan 14 2026 | PRJNA1080652 | Plasma |
| GSE179260 | Homo sapiens | Aug 31 2021 | PRJNA743024 | |
| GSE313050 | Danio rerio | Feb 09 2026 | PRJNA1378201 | embryo |
| GSE203629 | Gallus gallus | May 19 2025 | PRJNA841647 | |
| GSE278734 | Homo sapiens | Sep 25 2025 | PRJNA1168533 | 2D culture |
| GSE267143 | Mus musculus | Jan 18 2026 | PRJNA1109845 | |
| GSE317019 | Mus musculus | Jan 27 2026 | PRJNA1405681 | Spleen |
| GSE163642 | Mus musculus | Jun 30 2022 | PRJNA686990 | |
| GSE247506 | Homo sapiens | Mar 07 2024 | PRJNA1039249 | arterial blood |
| GSE165784 | Homo sapiens | Jan 27 2022 | PRJNA697911 | Fibrous membrane |
| GSE280152 | Mus musculus | Dec 03 2024 | PRJNA1176557 | Bone marrow |
| GSE120794 | Homo sapiens | Jun 20 2020 | PRJNA494559 | |
| GSE174620 | Homo sapiens | Aug 26 2021 | PRJNA730800 | |
| GSE309233 | Homo sapiens | Feb 12 2026 | PRJNA1334379 | cell line |
| GSE317347 | Homo sapiens | Jan 22 2026 | PRJNA1406771 | Liver |
| GSE300206 | Plasmodium falciparum | Feb 16 2026 | PRJNA1279250 | |
| GSE295597 | Homo sapiens | Jan 01 2026 | PRJNA1255233 | |
| GSE304005 | Mus musculus | Jan 01 2026 | PRJNA1298740 | brain |
| GSE294294 | Mus musculus | Jan 01 2026 | PRJNA1248956 | Spleen |
| GSE242112 | Homo sapiens | Jan 14 2026 | PRJNA1011337 | |
| GSE164802 | Cavia porcellus | Jan 15 2021 | PRJNA692095 | retina |
| GSE296166 | Mus musculus | Feb 11 2026 | PRJNA1257955 | HEART |
| GSE318283 | Mus musculus | Feb 05 2026 | PRJNA1418512 | Bone Marrow |
| GSE314945 | Saccharomyces cerevisiae | Jan 08 2026 | PRJNA1398624 | |
| GSE317412 | Mus musculus | Feb 06 2026 | PRJNA1412552 | Heart |
| GSE312883 | Mus musculus | Jan 26 2026 | PRJNA1377560 | Hind limb muscle |
| GSE289711 | Homo sapiens | Feb 13 2026 | PRJNA1224239 | |
| GSE233681 | Mus musculus | Aug 01 2023 | PRJNA977590 | Retina |
| GSE297418 | Mus musculus | Jan 27 2026 | PRJNA1263809 | |
| GSE254750 | Axonopus compressus | Jan 31 2026 | PRJNA1071617 | Root |
| GSE227619 | Canis lupus familiaris | Jul 25 2024 | PRJNA945919 | macular RPE/choroid |
| GSE293304 | Homo sapiens | May 12 2026 | PRJNA1243983 | |
| GSE308507 | Mus musculus | Jan 21 2026 | PRJNA1330790 | Cecum |
| GSE287870 | Homo sapiens | Jan 17 2026 | PRJNA1214837 | |
| GSE270878 | Mus musculus | Jan 14 2026 | PRJNA1128680 | tumour |
| GSE315728 | Homo sapiens | Feb 24 2026 | PRJNA1398859 | Ileum |
| GSE272733 | Mus musculus | Apr 01 2026 | PRJNA1138553 | Lymph nodes |
| GSE319060 | Mus musculus | Feb 14 2026 | PRJNA1422032 | Whole lung lysate |
| GSE308785 | Homo sapiens | Feb 20 2026 | PRJNA1332723 | pancreatic islets |
| GSE297385 | Homo sapiens | Jan 13 2026 | PRJNA1263727 | |
| GSE310737 | Homo sapiens | Feb 26 2026 | PRJNA1367000 | |
| GSE210480 | Homo sapiens | Aug 02 2024 | PRJNA865758 | |
| GSE221042 | Homo sapiens | Jan 18 2023 | PRJNA912653 | Retina |
| GSE307676 | Mus musculus | Sep 11 2025 | PRJNA1322504 | Retina |
| GSE283602 | Homo sapiens | Feb 05 2026 | PRJNA1194601 | |
| GSE282101 | Mus musculus | Feb 21 2026 | PRJNA1187194 | pancreatic cancer tissue |
| GSE246802 | Mus musculus | Dec 31 2024 | PRJNA1034631 | retina |
| GSE316802 | Mus musculus | Jan 21 2026 | PRJNA1404236 | Paw |
| GSE271497 | Homo sapiens | Jan 01 2026 | PRJNA1131825 | |

| | | | | |
|---|---|---|---|---|
| GSE312390 | Homo sapiens | Feb 28 2026 | PRJNA1373223 | Lung; Bronchus |
| GSE316835 | Homo sapiens | Jan 27 2026 | PRJNA1404322 | Cartilage |
| GSE294509 | Mus musculus | Feb 18 2026 | PRJNA1250420 | lymphoma tumor tissues |
| GSE124568 | Homo sapiens | Mar 12 2020 | PRJNA512519 | |
| GSE314261 | Homo sapiens | Jan 13 2026 | PRJNA1390672 | heral blood mononuclear cells |
| GSE319735 | Mus musculus | Feb 17 2026 | PRJNA1425023 | mouse liver |
| GSE311992 | Arabidopsis thaliana | Jan 31 2026 | PRJNA1372039 | whole plant |
| GSE224575 | Mus musculus | Jan 29 2026 | PRJNA931903 | thyroid |
| GSE283448 | Homo sapiens | Jan 30 2026 | PRJNA1193674 | |
| GSE300836 | Mus musculus | Feb 26 2026 | PRJNA1282178 | spleen |
| GSE166851 | Homo sapiens | Jun 16 2021 | PRJNA702116 | Retina |
| GSE198893 | Mus musculus | Sep 06 2023 | PRJNA817253 | retina |
| GSE246598 | Mus musculus | Jun 12 2024 | PRJNA1033690 | Retina |
| GSE314342 | Homo sapiens | Feb 06 2026 | | blood |
| GSE313587 | Mus musculus | Apr 01 2026 | PRJNA1380354 | RPE/choroid |
| GSE286216 | Homo sapiens | Jan 01 2026 | PRJNA1208438 | Colorectal cacner |
| GSE318884 | Homo sapiens | Feb 16 2026 | PRJNA1420764 | prostate |
| GSE275780 | Mus musculus | Feb 20 2026 | PRJNA1152981 | spleen |
| GSE287768 | Mus musculus | Mar 26 2025 | PRJNA1214589 | RPE/choroid |
| GSE279282 | Homo sapiens | Feb 10 2026 | PRJNA1171808 | |
| GSE291360 | Rattus norvegicus | Jan 01 2026 | PRJNA1233234 | sciatic nerve |
| GSE129355 | Mus musculus | Mar 31 2025 | PRJNA531189 | lung |
| GSE182520 | Homo sapiens;blank sample | Aug 27 2021 | PRJNA756561 | |
| GSE290301 | Mus musculus | Feb 25 2026 | PRJNA1227198 | Round Spermatids |
| GSE319621 | Mus musculus | Feb 20 2026 | PRJNA1424879 | spleen |
| GSE295601 | Homo sapiens | Feb 05 2026 | PRJNA1255286 | or-infiltrating lymphocyte (TIL) |
| GSE310469 | Mus musculus | Jan 21 2026 | PRJNA1366384 | Fallopian tube (Oviduct) |
| GSE161732 | Homo sapiens | Feb 26 2021 | PRJNA679269 | |
| GSE308855 | Mus musculus | Jan 16 2026 | PRJNA1333212 | Lung |
| GSE316921 | Homo sapiens | Jan 30 2026 | PRJNA1405544 | Biliary tract tumor tissue |
| GSE306117 | Homo sapiens | Jan 01 2026 | PRJNA1309161 | Breast Tumor |
| GSE292814 | Mus musculus | Feb 24 2026 | PRJNA1241295 | uterine tumor |
| GSE261413 | Mus musculus | Feb 17 2026 | PRJNA1086904 | |
| GSE256408 | Mus musculus | Apr 22 2025 | PRJNA1079362 | |
| GSE308085 | Mus musculus | Feb 27 2026 | PRJNA1328719 | |
| GSE302314 | Homo sapiens | Jan 30 2026 | PRJNA1289850 | brain organoid |
| GSE314234 | Caenorhabditis elegans | Feb 24 2026 | PRJNA1390502 | whole body |
| GSE315255 | Mus musculus | Jan 26 2026 | PRJNA1395517 | Tumor |
| GSE283102 | Mus musculus | Jan 01 2026 | PRJNA1191867 | Spleen |
| GSE285955 | Mus musculus | Jan 06 2026 | PRJNA1207043 | extensor digitorum longus |
| GSE186662 | Homo sapiens | Feb 10 2022 | PRJNA774999 | colon |
| GSE286402 | Homo sapiens | Jan 30 2026 | PRJNA1209068 | |
| GSE158909 | Homo sapiens | Jun 23 2022 | PRJNA666930 | |
| GSE274388 | feces metagenome | Feb 05 2026 | PRJNA1146325 | fecal |
| GSE314556 | Homo sapiens | Feb 11 2026 | PRJNA1391507 | |
| GSE293842 | Homo sapiens | Jan 26 2026 | PRJNA1246578 | |
| GSE294209 | Mus musculus | Feb 09 2026 | PRJNA1248559 | Whole testis |
| GSE319035 | Mus musculus | Feb 16 2026 | PRJNA1422001 | Liver |
| GSE244561 | Arabidopsis thaliana | Feb 23 2026 | PRJNA1023576 | 14 days old seedlings |
| GSE135092 | Homo sapiens | Jan 22 2020 | PRJNA557438 | RPE, Macula |
| GSE312829 | Mus musculus | Jan 20 2026 | PRJNA1377479 | dney cortical collecting duct |
| GSE315262 | Homo sapiens | May 06 2026 | PRJNA1395516 | na; Retinal pigmented epithelium |
| GSE254587 | Mus musculus | Jun 12 2024 | PRJNA1071096 | Retina |
| GSE284092 | Mus musculus | Jan 30 2026 | PRJNA1196984 | Heart |
| GSE276718 | Homo sapiens | Mar 10 2025 | PRJNA1158748 | eye |
| GSE243848 | Homo sapiens | Jan 01 2026 | PRJNA1020158 | glioblastoma |
| GSE276207 | Mus musculus | Feb 17 2025 | PRJNA1155704 | Retina |
| GSE230090 | Aus musculus;synthetic construc | May 27 2023 | PRJNA957443 | retina |
| GSE315500 | Mus musculus | Jan 11 2026 | PRJNA1398073 | vaginal epithelial tissue |
| GSE220637 | Mus | Jan 15 2026 | PRJNA910562 | prefrontal cortex |
| GSE197093 | Homo sapiens | Feb 21 2025 | PRJNA808970 | |
| GSE209758 | Homo sapiens | Jul 26 2023 | PRJNA862360 | |
| GSE319409 | Mus musculus | Feb 12 2026 | PRJNA1423246 | brain |

| | | | | |
|---|---|---|---|---|
| GSE318615 | Siniperca chuatsi | Feb 11 2026 | PRJNA1419314 | |
| GSE217333 | Oryctolagus cuniculus | Nov 08 2022 | PRJNA898403 | |
| GSE273146 | Homo sapiens | Jan 02 2026 | PRJNA1140246 | liver |
| GSE271283 | Mus musculus | Jan 13 2026 | PRJNA1130854 | spleen |
| GSE317084 | Arabidopsis thaliana | Jan 26 2026 | PRJNA1405754 | whole seedlings |
| GSE229160 | Homo sapiens | May 01 2023 | PRJNA953085 | |
| GSE220155 | Homo sapiens | Jan 12 2023 | PRJNA908876 | etinal pigment epithelium |
| GSE302200 | Homo sapiens | Jan 28 2026 | PRJNA1289181 | |
| GSE298300 | Homo sapiens | Feb 13 2026 | PRJNA1268911 | tumor |
| GSE277427 | Mus musculus | Jan 09 2026 | PRJNA1162141 | |
| GSE318177 | Homo sapiens | Feb 21 2026 | PRJNA1416851 | whole blood |
| GSE296292 | Homo sapiens;synthetic construc | Jun 18 2026 | PRJNA1258872 | |
| GSE223593 | Mus musculus | Jun 14 2023 | PRJNA926937 | Retina |
| GSE246943 | Homo sapiens | Sep 30 2025 | PRJNA1035134 | |
| GSE311308 | Mus musculus | Feb 18 2026 | PRJNA1368915 | kidney |
| GSE311617 | Homo sapiens | Jan 26 2026 | PRJNA1370020 | |
| GSE318622 | Sus scrofa domesticus | Feb 08 2026 | PRJNA1419572 | Chronic venous thrombus |
| GSE197935 | Gallus gallus | May 11 2022 | PRJNA812910 | |
| GSE304493 | Homo sapiens | Feb 27 2026 | PRJNA1301382 | Blood cells |
| GSE319121 | Homo sapiens | Feb 10 2026 | PRJNA1422138 | |
| GSE169066 | Homo sapiens | Mar 18 2022 | PRJNA715097 | RPE |
| GSE317922 | Danio rerio | Jan 28 2026 | PRJNA1415820 | whole-larvae |
| GSE304355 | Mus musculus | Feb 16 2026 | PRJNA1300450 | Embryonic cortices E16.5 |
| GSE299490 | Mus musculus | Apr 11 2026 | PRJNA1274721 | |
| GSE279039 | Mus musculus | Jan 01 2026 | PRJNA1170220 | mammary |
| GSE306901 | Homo sapiens | Feb 25 2026 | PRJNA1312885 | clear cell carcinoma (CCC) |
| GSE235342 | Homo sapiens | Jan 05 2025 | PRJNA985722 | |
| GSE155154 | Homo sapiens | Aug 11 2021 | PRJNA648863 | |
| GSE318399 | Homo sapiens | Feb 07 2026 | PRJNA1418685 | cell |
| GSE308660 | Lagopus muta hyperborea | Dec 16 2025 | PRJNA1331821 | brain (hypothalamus) |
| GSE317986 | axillaris;Petunia integrifolia subs | Jan 30 2026 | PRJNA1416273 | upper leaves |
| GSE316714 | Homo sapiens | Feb 25 2026 | PRJNA1403821 | coronary artery |
| GSE227056 | Mus musculus | May 19 2023 | PRJNA942964 | retina |
| GSE300603 | Homo sapiens;Mus musculus | Jan 09 2026 | PRJNA1281525 | small intestine |
| GSE234476 | Danio rerio | Jan 22 2026 | PRJNA981502 | |
| GSE174547 | Mus musculus | Feb 17 2026 | PRJNA730519 | s residing in white adipose tissue (WAT) |
| GSE293054 | Mus musculus | Jan 01 2026 | PRJNA1242695 | lung |
| GSE281415 | Homo sapiens | Jan 22 2026 | PRJNA1183822 | ovary |
| GSE318842 | Penicillium oxalicum | Feb 12 2026 | PRJNA1420702 | strain |
| GSE282942 | Mus musculus | Jan 15 2026 | PRJNA1191051 | VAT |
| GSE289441 | Homo sapiens | Feb 12 2026 | PRJNA1222912 | |
| GSE303929 | Mus musculus | Feb 18 2026 | PRJNA1298246 | Ventricular heart |
| GSE198785 | Mus musculus | Apr 20 2022 | PRJNA816874 | retina |
| GSE317892 | Mus musculus | Jan 28 2026 | PRJNA1415761 | Brian, mPFC |
| GSE155040 | Mus musculus | Aug 13 2023 | PRJNA648325 | |
| GSE153817 | Mus musculus | Sep 29 2020 | PRJNA644202 | |
| GSE295920 | Mus musculus | Feb 25 2026 | PRJNA1256787 | |
| GSE319900 | Homo sapiens | Feb 23 2026 | PRJNA1425498 | human gingiva |
| GSE275418 | Mus musculus | Feb 06 2026 | PRJNA1150633 | colonic differentiated cell |
| GSE310935 | Homo sapiens | Jan 24 2026 | PRJNA1367576 | |
| GSE289562 | Danio rerio | Mar 26 2025 | PRJNA1223441 | Whole eye |
| GSE234237 | Homo sapiens | Jan 15 2026 | PRJNA980608 | |
| GSE297416 | Mus musculus | Jan 27 2026 | PRJNA1263807 | |
| GSE285735 | Canis lupus familiaris | Jun 01 2025 | PRJNA1205733 | Neurtoretina |
| GSE301741 | Homo sapiens | Feb 10 2026 | | Oral Cavity |
| GSE312361 | Mus musculus | Feb 23 2026 | PRJNA1373163 | teral motor cortex whole single cells |
| GSE317656 | Homo sapiens | Feb 12 2026 | PRJNA1414597 | |
| GSE317349 | Homo sapiens | Jan 22 2026 | PRJNA1406774 | Uveal melanoma |
| GSE160845 | Mus musculus | Mar 15 2021 | PRJNA674628 | |
| GSE178166 | Homo sapiens | Nov 21 2022 | PRJNA737493 | |
| GSE278306 | Rattus norvegicus | Sep 27 2024 | PRJNA1166306 | Retina |
| GSE297421 | Aedes albopictus | Feb 11 2026 | PRJNA1263820 | whole cell |
| GSE276978 | Mus musculus | Feb 11 2026 | PRJNA1159874 | TMJ |

| | | | | |
|---|---|---|---|---|
| GSE290099 | Homo sapiens | Feb 16 2026 | PRJNA1226207 | |
| GSE293990 | Saccharomyces cerevisiae | Feb 16 2026 | PRJNA1247544 | |
| GSE307093 | Rattus norvegicus | Jan 14 2026 | PRJNA1314251 | |
| GSE284622 | Mus musculus | Jan 05 2026 | PRJNA1199938 | Mesenteric Lymph Node |
| GSE309101 | Zea mays | Feb 25 2026 | PRJNA1333950 | 14DAS SAM |
| GSE180641 | Mus musculus | Nov 16 2022 | PRJNA749017 | etinal pigment epithelium |
| GSE289669 | Homo sapiens | Feb 26 2026 | PRJNA1223912 | |
| GSE301155 | Homo sapiens | Jan 08 2026 | PRJNA1284207 | |
| GSE311822 | Rattus norvegicus | Jan 11 2026 | PRJNA1370949 | hippocampus |
| GSE319685 | Mus musculus | Feb 23 2026 | PRJNA1424959 | Lung |
| GSE202146 | Homo sapiens | Mar 01 2025 | PRJNA834793 | Macula Choroid |
| GSE234341 | Homo sapiens | Jan 07 2026 | PRJNA981085 | |
| GSE316014 | Histoplasma ohiense | Jan 26 2026 | PRJNA1400458 | |
| GSE316985 | Homo sapiens | Feb 18 2026 | PRJNA1405641 | |
| GSE285674 | Mus musculus | Jan 12 2026 | PRJNA1205554 | spleen |
| GSE294191 | Homo sapiens | Jan 01 2026 | PRJNA1248522 | |
| GSE297557 | Homo sapiens | Jun 20 2025 | PRJNA1265179 | |
| GSE292057 | Homo sapiens | Aug 18 2025 | PRJNA1236401 | Retinal Organoid |
| GSE271038 | Mus musculus | Oct 31 2024 | PRJNA1129460 | retina+choroid |
| GSE307935 | Homo sapiens | Feb 14 2026 | PRJNA1327850 | |
| GSE310687 | Triticum aestivum | Feb 11 2026 | PRJNA1366938 | microspore cultures |
| GSE243241 | Pomacea canaliculata | Dec 31 2024 | PRJNA1017465 | retina |
| GSE169097 | Mus musculus | May 06 2025 | PRJNA715208 | |
| GSE293529 | Mus musculus | Feb 22 2026 | PRJNA1244883 | Kidney |
| GSE222534 | Mus musculus | Jan 01 2026 | PRJNA922481 | 2-cell embryo |
| GSE253868 | Mus musculus | Feb 01 2024 | PRJNA1067587 | Spinal cord |
| GSE316100 | Danio rerio | Jan 12 2026 | PRJNA1400599 | larvae |
| GSE185028 | Homo sapiens | Dec 21 2022 | PRJNA767404 | |
| GSE240957 | Homo sapiens | Oct 11 2023 | PRJNA1005937 | cell line |
| GSE309268 | Homo sapiens | Feb 18 2026 | PRJNA1335000 | neuroblastoma |
| GSE255520 | Mus musculus | Nov 04 2024 | PRJNA1075146 | retina |
| GSE301041 | Homo sapiens | Feb 09 2026 | PRJNA1283120 | |
| GSE167026 | Canis lupus familiaris | Feb 19 2021 | PRJNA702758 | |
| GSE274666 | Mus musculus | Jan 07 2025 | PRJNA1147614 | Retina |
| GSE298909 | Mus musculus | Feb 22 2026 | PRJNA1271777 | lung |
| GSE298587 | Mus musculus | Feb 13 2026 | PRJNA1269982 | |
| GSE199942 | Homo sapiens | Apr 04 2022 | PRJNA822305 | serum |
| GSE316655 | Homo sapiens | Jan 16 2026 | PRJNA1403544 | tumor |
| GSE317449 | Homo sapiens | Jan 31 2026 | PRJNA1412982 | Liver |
| GSE288049 | Homo sapiens | May 22 2025 | PRJNA1215937 | |
| GSE285993 | Homo sapiens | Jan 31 2026 | PRJNA1207445 | |
| GSE310793 | Homo sapiens | Feb 18 2026 | PRJNA1367083 | apillary Thyroid Carcinoma |
| GSE319955 | Drosophila melanogaster | Feb 23 2026 | PRJNA1425779 | Brain |
| GSE102205 | Homo sapiens | Jul 27 2020 | PRJNA397019 | |
| GSE284063 | Gallus gallus | Jan 14 2025 | PRJNA1196933 | optic cup |
| GSE255517 | Homo sapiens;Mus musculus | Jan 23 2026 | PRJNA1075135 | |
| GSE288234 | Rattus norvegicus | Jan 21 2026 | PRJNA1216604 | liver |
| GSE300986 | Homo sapiens | Feb 20 2026 | PRJNA1282960 | blood |
| GSE253653 | Homo sapiens | Jan 08 2026 | PRJNA1066715 | Bone |
| GSE301918 | Mus musculus | Jan 21 2026 | PRJNA1287743 | brainstem |
| GSE266903 | Physcomitrium patens | Jan 26 2026 | PRJNA1108701 | seedlings |
| GSE301596 | Homo sapiens | Jan 01 2026 | PRJNA1285712 | nor-free subventricular zone |
| GSE294207 | Drosophila melanogaster | Jan 07 2026 | PRJNA1248554 | Central brain |
| GSE299753 | mouse metagenome | Feb 11 2026 | PRJNA1276436 | Blood |
| GSE234971 | Homo sapiens | Nov 29 2023 | PRJNA983813 | Retina |
| GSE314070 | Mus musculus | Feb 11 2026 | PRJNA1389918 | small intestine |
| GSE313846 | Homo sapiens | Jan 16 2026 | PRJNA1381664 | notochord |
| GSE319246 | Mus musculus | Feb 16 2026 | PRJNA1422681 | brain |
| GSE307332 | Mus musculus | Jan 13 2026 | PRJNA1315255 | Intestinal epithelial cells |
| GSE296388 | Homo sapiens | Feb 05 2026 | PRJNA1259037 | |
| GSE220550 | Drosophila melanogaster | Dec 07 2023 | PRJNA910186 | retina |
| GSE230938 | Mus musculus | Jan 26 2026 | PRJNA962931 | heart |
| GSE313871 | Homo sapiens | Feb 11 2026 | PRJNA1381702 | |

| | | | | |
|---|---|---|---|---|
| GSE290033 | Mus musculus | Jan 07 2026 | PRJNA1225844 | bone marrow |
| GSE296864 | Mus musculus | Feb 19 2026 | PRJNA1261837 | |
| GSE313682 | Mus musculus | Jan 19 2026 | PRJNA1381101 | Lung |
| GSE242810 | Mus musculus | Sep 10 2023 | PRJNA1014780 | leptomeninges |
| GSE314558 | Homo sapiens | Feb 11 2026 | PRJNA1391501 | |
| GSE281197 | Mus musculus | Feb 17 2026 | PRJNA1182719 | liver |
| GSE289852 | Oryza sativa | Jan 26 2026 | PRJNA1225195 | 14-day-old seedlings |
| GSE273302 | Oryza sativa | Jan 19 2026 | PRJNA1141400 | stigma |
| GSE318451 | Mus musculus | Feb 07 2026 | PRJNA1418742 | Pancreas |
| GSE288833 | Mus musculus | May 15 2026 | PRJNA1219573 | retina |
| GSE264243 | Oryza sativa Japonica Group | Jan 24 2026 | PRJNA1101526 | Shoot |
| GSE224112 | Homo sapiens | Jan 30 2026 | PRJNA929923 | |
| GSE303747 | Homo sapiens | Feb 26 2026 | PRJNA1297603 | Breast |
| GSE176337 | Homo sapiens | Jun 08 2021 | PRJNA735779 | |
| GSE315650 | Parascaris univalens | Jan 10 2026 | PRJNA1398393 | 1-cell embryo |
| GSE286518 | Homo sapiens | Jan 20 2026 | PRJNA1209859 | CSF |
| GSE303770 | Homo sapiens | Jan 22 2026 | | |
| GSE208055 | Homo sapiens | Aug 17 2022 | PRJNA858063 | |
| GSE307573 | Mus musculus | Jan 19 2026 | PRJNA1322316 | Lung |
| GSE290127 | Mus musculus | Feb 20 2026 | PRJNA1226233 | |
| GSE319407 | Homo sapiens | Feb 25 2026 | | Teratoma |
| GSE319889 | Homo sapiens | Feb 23 2026 | PRJNA1425481 | brain |
| GSE313965 | Homo sapiens;Mus musculus | Jan 06 2026 | PRJNA1387444 | Testis |
| GSE295345 | Mus musculus | Jan 16 2026 | PRJNA1254236 | lung |
| GSE319365 | Mus musculus | Feb 20 2026 | PRJNA1423160 | Lung |
| GSE290704 | Homo sapiens | Feb 16 2026 | PRJNA1229598 | immune cells |
| GSE304796 | Homo sapiens | Jan 22 2026 | PRJNA1302758 | |
| GSE312845 | Mus musculus | Feb 18 2026 | PRJNA1377515 | cell line |
| GSE216060 | Danio rerio | Oct 25 2022 | PRJNA891853 | eye |
| GSE314012 | Homo sapiens | Jan 01 2026 | PRJNA1389466 | bone |
| GSE295566 | Mus musculus | Feb 20 2026 | PRJNA1255198 | lung |
| GSE249059 | Escherichia coli | Jan 28 2026 | PRJNA1047101 | |
| GSE280080 | Mus musculus | Feb 06 2026 | PRJNA1176142 | Tumor |
| GSE315711 | Homo sapiens | Feb 13 2026 | PRJNA1398826 | palatine tonsils |
| GSE279687 | Homo sapiens | Feb 25 2026 | PRJNA1174089 | |
| GSE160356 | Homo sapiens | Oct 27 2023 | PRJNA672989 | Endothelial Cells |
| GSE294252 | Mus musculus | Feb 05 2026 | | mammary tumor |
| GSE186311 | Homo sapiens | Jan 29 2026 | PRJNA773371 | |
| GSE317381 | Homo sapiens | Jan 30 2026 | PRJNA1412369 | dentate gyrus (DG) |
| GSE298278 | Homo sapiens | Jan 28 2026 | PRJNA1268873 | oids derived from primary human placentas |
| GSE294530 | Mus musculus | Feb 12 2026 | PRJNA1250457 | |
| GSE274199 | Homo sapiens | Feb 27 2026 | PRJNA1145391 | lung |
| GSE268328 | Mus musculus | Feb 23 2026 | PRJNA1116228 | Blood (sera) |
| GSE287751 | Mus musculus | Jan 01 2026 | PRJNA1214390 | bone-marrow |
| GSE303725 | Mus musculus | Feb 12 2026 | PRJNA1297160 | mPFC |
| GSE318351 | Mus musculus | Feb 04 2026 | PRJNA1418622 | retina |
| GSE315605 | Mus musculus | Jan 25 2026 | PRJNA1398324 | |
| GSE197959 | Homo sapiens | Mar 03 2023 | PRJNA812938 | |
| GSE200054 | Mus musculus | Apr 03 2025 | PRJNA822796 | retina |
| GSE304871 | Homo sapiens | Feb 19 2026 | PRJNA1303109 | |
| GSE305810 | Homo sapiens | Oct 15 2025 | PRJNA1307873 | |
| GSE235866 | Homo sapiens | Feb 29 2024 | PRJNA987727 | Retinal organoid |
| GSE185807 | Mus musculus | Jan 07 2022 | PRJNA770910 | amygdala |
| GSE314664 | Mus musculus | Jan 09 2026 | PRJNA1392425 | |
| GSE290610 | Homo sapiens | Jan 21 2026 | PRJNA1228980 | brain |
| GSE316596 | Rattus | Jan 22 2026 | PRJNA1403460 | atria |
| GSE316918 | Mus musculus | Jan 21 2026 | PRJNA1405535 | Paw |
| GSE241353 | Escherichia coli;Mus musculus | Sep 01 2023 | PRJNA1007735 | whole retina |
| GSE319307 | Bos taurus | Feb 16 2026 | | plasma |
| GSE290627 | Mus musculus | Feb 19 2026 | PRJNA1229439 | |
| GSE283718 | Homo sapiens | Jan 21 2026 | PRJNA1195091 | |
| GSE289489 | Homo sapiens | Feb 10 2026 | PRJNA1223303 | |
| GSE317295 | Sus scrofa | Feb 09 2026 | PRJNA1406695 | term placenta |

| | | | | |
|---|---|---|---|---|
| GSE286218 | Mus musculus | Jan 14 2026 | PRJNA1208435 | re-isolated satellite cells |
| GSE243907 | Danio rerio | Jan 07 2026 | PRJNA1020549 | brain |
| GSE237205 | Anolis sagrei | Oct 01 2023 | PRJNA994168 | Retina |
| GSE145330 | Danio rerio | May 19 2020 | PRJNA606767 | Retina |
| GSE255827 | Homo sapiens | Feb 18 2026 | PRJNA1076597 | |
| GSE302011 | Homo sapiens | Jan 27 2026 | PRJNA1288348 | breast cancer |
| GSE319465 | Homo sapiens | Feb 12 2026 | PRJNA1423328 | |
| GSE256212 | Mus musculus | Nov 22 2024 | PRJNA1078426 | Retina |
| GSE282266 | Homo sapiens | Jan 05 2026 | PRJNA1188017 | human lymphocytes |
| GSE293583 | Mus musculus | Jan 14 2026 | PRJNA1245301 | Pancreatic tumor |
| GSE176355 | Rattus norvegicus | Jul 01 2021 | PRJNA735937 | retina |
| GSE273435 | Homo sapiens | Nov 15 2024 | PRJNA1141914 | Blood |
| GSE138734 | Homo sapiens | Apr 15 2021 | PRJNA576920 | |
| GSE316149 | Homo sapiens | Jan 14 2026 | PRJNA1401725 | whole blood |
| GSE316456 | Mus musculus | Jun 17 2026 | PRJNA1402945 | Retina |
| GSE299921 | Homo sapiens | Jul 17 2025 | PRJNA1277328 | Retinal organoid |
| GSE279664 | Homo sapiens | Jan 31 2026 | PRJNA1174052 | |
| GSE294799 | Danio rerio | Feb 16 2026 | PRJNA1251618 | endoderm-enriched |
| GSE197420 | Mus musculus | Sep 23 2022 | PRJNA810190 | retina |
| GSE292377 | Mus musculus | Jan 22 2026 | PRJNA1238266 | Spleen |
| GSE289703 | Homo sapiens | Feb 19 2025 | PRJNA1224224 | |
| GSE317400 | Homo sapiens | Jan 30 2026 | PRJNA1412499 | rian serous adenocarcinoma |
| GSE304020 | Homo sapiens | Feb 27 2026 | PRJNA1298775 | Blood |
| GSE117959 | Homo sapiens;Mus musculus | Jan 24 2020 | PRJNA483791 | retina |
| GSE174167 | Homo sapiens | Dec 13 2021 | PRJNA728658 | |
| GSE312100 | Mus musculus | Jan 25 2026 | PRJNA1372182 | tumor |
| GSE261183 | Mus musculus | Mar 31 2025 | PRJNA1085718 | RPE/choroid complex |
| GSE226108 | Homo sapiens | Mar 14 2023 | PRJNA938514 | peripheral region of retina |
| GSE310929 | Homo sapiens | Feb 05 2026 | PRJNA1367552 | Whole blood |
| GSE297754 | Mus musculus | Feb 08 2026 | PRJNA1266133 | bone marrow stroma |
| GSE316220 | Mus musculus | Jan 16 2026 | PRJNA1401828 | Ovary |
| GSE288123 | Mycosarcoma maydis | Feb 18 2026 | PRJNA1216232 | |
| GSE315815 | Homo | Jan 08 2026 | PRJNA1399353 | |
| GSE314687 | Homo sapiens | Feb 24 2026 | PRJNA1392457 | |
| GSE287941 | Homo sapiens | Jan 01 2026 | PRJNA1215333 | |
| GSE297036 | Mus musculus | May 29 2026 | PRJNA1262423 | Neurosensory Retina |
| GSE152949 | Arabidopsis thaliana | Jun 23 2020 | PRJNA641038 | Whole plant |
| GSE274059 | Mus musculus | Feb 18 2026 | PRJNA1144845 | Mouse uterus |
| GSE222737 | Mus musculus | Jan 01 2026 | | |
| GSE297529 | Homo sapiens | Jan 22 2026 | PRJNA1265151 | PBMC |
| GSE310484 | Homo sapiens | Apr 01 2026 | PRJNA1366416 | |
| GSE301361 | Mus musculus | Feb 17 2026 | PRJNA1284852 | |
| GSE215233 | Arvicanthis niloticus | Dec 09 2022 | PRJNA889410 | Adipose |
| GSE267166 | Mus musculus | Feb 23 2026 | PRJNA1109992 | lymph node |
| GSE206624 | Mus musculus | Aug 20 2022 | PRJNA851597 | retina |
| GSE297990 | Mus musculus | Jan 21 2026 | PRJNA1267078 | liver |
| GSE287976 | Homo sapiens | Jan 07 2026 | PRJNA1215395 | peripheral blood |
| GSE331097 | Rattus norvegicus | May 19 2026 | PRJNA1466891 | Retina |
| GSE314843 | Mus musculus | Feb 23 2026 | PRJNA1392983 | skeletal muscle |
| GSE309460 | Mus musculus | Feb 16 2026 | PRJNA1336124 | |
| GSE318385 | Homo sapiens | Feb 15 2026 | PRJNA1418651 | |
| GSE287785 | Mus musculus | Oct 29 2025 | PRJNA1214699 | E13.5 mouse embryos |
| GSE274213 | Mus musculus | Aug 31 2025 | PRJNA1145412 | |
| GSE164904 | Mus musculus | Jul 21 2022 | PRJNA692431 | |
| GSE316708 | Homo sapiens | Jan 20 2026 | PRJNA1403813 | blood |
| GSE197503 | Mus musculus | Nov 15 2022 | PRJNA810711 | |
| GSE319486 | Homo sapiens;Mus musculus | Feb 16 2026 | PRJNA1423601 | |
| GSE292263 | Mus musculus | Feb 17 2026 | PRJNA1237779 | spleen |
| GSE316217 | Mus musculus | Jan 18 2026 | PRJNA1401818 | medial prefrontal cortex |
| GSE316356 | Mus musculus | Jan 14 2026 | | spleen |
| GSE234003 | Homo sapiens | Nov 29 2023 | PRJNA979192 | Retina |
| GSE283849 | Homo sapiens | Feb 15 2026 | PRJNA1195901 | |
| GSE266071 | Homo sapiens | Sep 24 2024 | PRJNA1105634 | retina |

| | | | | |
|---|---|---|---|---|
| GSE319949 | Mus musculus | Feb 18 2026 | PRJNA1425597 | Brain |
| GSE314093 | Homo sapiens | Jan 19 2026 | PRJNA1390081 | |
| GSE288055 | Mus musculus | Jan 01 2026 | PRJNA1215957 | hippocampal tissue |
| GSE317455 | Homo sapiens | Jan 31 2026 | PRJNA1412944 | primary prostate cancer |
| GSE252822 | Zea mays | Jan 07 2026 | PRJNA1062773 | Root |
| GSE299943 | Homo sapiens | Jan 27 2026 | | |
| GSE317628 | Homo sapiens | Jan 28 2026 | PRJNA1414568 | tumor |
| GSE143354 | Mus musculus | Jan 02 2026 | PRJNA600234 | Hippocampus |
| GSE308540 | Homo sapiens | Jan 22 2026 | PRJNA1330848 | |
| GSE312328 | Mus musculus | Jan 15 2026 | PRJNA1373029 | Spleen |
| GSE269813 | Mus musculus | Jun 28 2024 | PRJNA1124236 | |
| GSE304717 | Danio rerio | Jan 08 2026 | PRJNA1302399 | 1 hpf (~4-cell) |
| GSE188560 | Danio rerio | Nov 10 2022 | PRJNA779419 | |
| GSE317300 | Medicago sativa | Jan 27 2026 | PRJNA1406705 | flowers |
| GSE267027 | Homo sapiens | Feb 05 2026 | PRJNA1109373 | or-infiltrating lymphocyte (TIL) |
| GSE259274 | Homo sapiens | Jan 07 2026 | PRJNA1080565 | SKOV3 cells |
| GSE289003 | Homo sapiens | Jan 12 2026 | PRJNA1220590 | |
| GSE189263 | Homo sapiens | Feb 23 2022 | PRJNA782306 | |
| GSE319292 | Danio rerio | Feb 16 2026 | PRJNA1422787 | |
| GSE196376 | Mus musculus | Jan 22 2026 | PRJNA804542 | |
| GSE136800 | Bos taurus | Sep 01 2020 | PRJNA563691 | |
| GSE304042 | Homo sapiens | Jun 11 2026 | PRJNA1298815 | |
| GSE269907 | Escherichia coli | Feb 09 2026 | PRJNA1124095 | |
| GSE305891 | Mus musculus | Jan 21 2026 | PRJNA1308123 | lymph node |
| GSE254603 | Mus musculus | Jan 12 2026 | PRJNA1071122 | spleen |
| GSE317223 | Rattus norvegicus | Jan 26 2026 | PRJNA1406304 | serum |
| GSE278773 | Homo sapiens | Jan 12 2026 | PRJNA1168876 | |
| GSE315877 | Homo sapiens | Jan 09 2026 | PRJNA1399477 | Kidney |
| GSE192734 | Homo sapiens | Dec 01 2022 | PRJNA792993 | |
| GSE299857 | Homo sapiens | Jun 18 2026 | PRJNA1277163 | retinal organoids |
| GSE202472 | Homo sapiens | Sep 09 2022 | PRJNA836268 | Adult Retina |
| GSE300400 | Sus scrofa | Jan 08 2026 | PRJNA1280891 | endometrial |
| GSE297229 | Mus musculus | Jan 12 2026 | PRJNA1263207 | Lung |
| GSE314816 | Homo sapiens | Jan 06 2026 | PRJNA1392643 | Placenta |
| GSE298836 | Mus musculus | Feb 22 2026 | PRJNA1271472 | |
| GSE310384 | Homo sapiens | Jan 05 2026 | PRJNA1365861 | Peripheral blood |
| GSE263465 | Sus scrofa | Jan 25 2026 | PRJNA1097993 | dorsal root ganglion |
| GSE305662 | Mus musculus | Feb 18 2026 | PRJNA1307434 | liver |
| GSE308301 | Mus musculus | Jan 09 2026 | PRJNA1330160 | brain |
| GSE317069 | Homo sapiens;Mus musculus | Jan 26 2026 | PRJNA1405729 | Diaphysis, femur |
| GSE148187 | Homo sapiens | Sep 22 2020 | PRJNA623400 | |
| GSE303793 | Mus musculus | Sep 30 2025 | PRJNA1297691 | retina |
| GSE102458 | Rattus norvegicus | Jul 04 2020 | PRJNA397767 | Retinal ganglion cell |
| GSE317576 | Mus musculus | Jan 27 2026 | PRJNA1414502 | Lung |
| GSE253960 | Homo sapiens | Jan 26 2024 | PRJNA1068002 | na; Retinal pigmented epithelium |
| GSE181323 | Mus musculus | Dec 20 2022 | PRJNA751575 | retina |
| GSE110019 | Felis catus | Apr 07 2020 | PRJNA432566 | |
| GSE271982 | Mus musculus | Feb 24 2026 | PRJNA1134240 | Embryo |
| GSE312714 | Mus musculus | Apr 21 2026 | PRJNA1375756 | Retina |
| GSE267133 | Homo sapiens | Mar 26 2025 | PRJNA1109819 | |
| GSE277648 | Mus musculus | Jan 16 2026 | PRJNA1163280 | splenocytes |
| GSE312918 | Mus musculus | Jan 30 2026 | PRJNA1377595 | |
| GSE194158 | Danio rerio | Jan 24 2022 | PRJNA799333 | |
| GSE285853 | Homo sapiens | Jan 14 2026 | PRJNA1206222 | cell line |
| GSE316795 | Caenorhabditis elegans | Jan 21 2026 | PRJNA1404223 | rmaphroditic adult whole animals |
| GSE289395 | Homo sapiens | Jan 22 2026 | PRJNA1222848 | T cell |
| GSE314450 | Homo sapiens | Jan 09 2026 | PRJNA1391338 | |
| GSE136565 | Mus musculus | Jan 08 2020 | PRJNA562691 | retina |
| GSE220625 | Mus musculus | Sep 15 2023 | PRJNA910551 | choroid |
| GSE318318 | Mus musculus | Feb 06 2026 | PRJNA1418562 | islet |
| GSE233078 | Rattus norvegicus | Jan 12 2026 | PRJNA974862 | kidney |
| GSE224344 | Mus musculus | Feb 12 2026 | PRJNA930685 | Femur |
| GSE291445 | Homo sapiens | Oct 07 2025 | PRJNA1233770 | Pigmented epithelium;Retina |

| | | | | |
|---|---|---|---|---|
| GSE290346 | Mus musculus | Feb 24 2026 | PRJNA1227521 | |
| GSE269541 | Mus musculus | Feb 25 2026 | PRJNA1122301 | kidney |
| GSE310419 | Homo sapiens | Feb 23 2026 | PRJNA1365911 | |
| GSE175929 | Danio rerio | Aug 30 2021 | PRJNA734313 | |
| GSE303039 | Mus musculus | Feb 05 2026 | PRJNA1293395 | Bone Marrow |
| GSE166816 | Mus musculus | Sep 20 2022 | PRJNA702064 | Lung |
| GSE316951 | Cavia porcellus | Jan 27 2026 | PRJNA1405581 | cochlear |
| GSE253526 | Homo sapiens | Nov 12 2024 | PRJNA1066214 | |
| GSE311167 | Homo sapiens | Feb 25 2026 | PRJNA1368768 | |
| GSE184818 | Mus musculus | Jan 01 2022 | PRJNA766595 | whole retina |
| GSE310664 | Homo sapiens | Jan 14 2026 | PRJNA1366912 | organoids |
| GSE283672 | Homo sapiens | Jan 21 2026 | PRJNA1195019 | |
| GSE241239 | Mus musculus | Aug 20 2024 | PRJNA1006773 | retina |
| GSE166935 | Homo sapiens | Dec 01 2021 | PRJNA702442 | |
| GSE299505 | Mus musculus | Feb 16 2026 | PRJNA1274870 | |
| GSE316872 | Homo sapiens | Jan 24 2026 | PRJNA1405468 | Peripheral blood |
| GSE269337 | Mus musculus | Feb 13 2025 | PRJNA1121243 | retina |
| GSE144724 | Mus musculus | Sep 12 2023 | PRJNA604711 | |
| GSE302464 | Arabidopsis thaliana | Jan 19 2026 | PRJNA1291032 | seedling |
| GSE306378 | Homo sapiens | Jan 01 2026 | PRJNA1310548 | Blood |
| GSE161103 | Mus musculus | Apr 28 2021 | PRJNA675587 | |
| GSE320486 | Homo sapiens | Feb 26 2026 | | retinal pigment epithelium |
| GSE298564 | Homo sapiens | Jan 26 2026 | PRJNA1269945 | sal squamous cell carcinoma |
| GSE253367 | Acinetobacter baumannii AB507 | Jan 15 2026 | PRJNA1065420 | |
| GSE143870 | Homo sapiens | Jan 10 2026 | PRJNA601936 | |
| GSE180781 | Danio rerio;Homo sapiens | Jul 01 2022 | PRJNA749558 | Eye |
| GSE318724 | Mus musculus | Feb 11 2026 | PRJNA1419837 | Spleen |
| GSE106298 | Mus musculus | Oct 27 2020 | PRJNA416250 | etinal pigment epithelium |
| GSE280283 | Mus musculus | Jan 30 2026 | PRJNA1177523 | Trachea |
| GSE318046 | Homo sapiens | Feb 16 2026 | PRJNA1416380 | rebral cortex - glioblastoma |
| GSE235582 | Homo sapiens | Feb 15 2024 | PRJNA986510 | organoid |
| GSE316845 | Mus musculus | Jan 26 2026 | PRJNA1405125 | Heart |
| GSE195505 | Homo sapiens | Sep 23 2022 | PRJNA800825 | Cortical organoid |
| GSE318896 | Homo sapiens | Feb 12 2026 | | Cardiomyocytes |
| GSE318418 | Homo sapiens | Feb 09 2026 | PRJNA1418712 | hepatocellular carcinoma |
| GSE236005 | Petromyzon marinus | Mar 04 2024 | PRJNA988411 | Retina |
| GSE290324 | Mus musculus | Feb 09 2026 | PRJNA1227476 | Liver |
| GSE256073 | Mus musculus | Feb 18 2026 | PRJNA1077991 | |
| GSE317796 | Homo sapiens | Feb 06 2026 | PRJNA1415279 | vaginal |
| GSE300824 | Ceratotherium simum simum | Jan 14 2026 | PRJNA1282156 | In vivo cells |
| GSE265774 | Homo sapiens | Oct 14 2025 | | peripheral region of retina |
| GSE291427 | Homo sapiens | Jan 26 2026 | PRJNA1233428 | |
| GSE183206 | Mus musculus | Apr 13 2022 | PRJNA759613 | Retina |
| GSE150839 | Danio rerio | Jun 24 2020 | PRJNA633835 | retina |
| GSE319166 | [Clostridium] scindens | Feb 15 2026 | PRJNA1422238 | Strain |
| GSE309811 | Homo sapiens | Feb 26 2026 | PRJNA1363567 | |
| GSE210099 | Mus musculus | Aug 05 2022 | PRJNA863657 | PE-Choroid-slera complex |
| GSE319479 | Mus musculus | Feb 13 2026 | PRJNA1423516 | Tumor |
| GSE155826 | Homo sapiens | Jan 08 2021 | PRJNA655712 | |
| GSE140466 | Homo sapiens | Feb 05 2020 | PRJNA589853 | |
| GSE318652 | Mus musculus | Feb 10 2026 | PRJNA1419720 | Liver |
| GSE200048 | Mus musculus | Apr 03 2025 | PRJNA822804 | retina |
| GSE315306 | Oryctolagus cuniculus | Jan 07 2026 | PRJNA1395614 | |
| GSE201254 | Mus musculus | Jun 28 2022 | PRJNA830507 | retina |
| GSE317952 | Danio rerio | Jan 30 2026 | PRJNA1416083 | whole larvae |
| GSE186731 | Homo sapiens | Feb 10 2022 | PRJNA775808 | |
| GSE294110 | Homo sapiens | Jan 20 2026 | PRJNA1248404 | Skin |
| GSE273663 | Homo sapiens | May 06 2025 | PRJNA1142588 | |
| GSE286233 | Homo sapiens | Feb 20 2026 | PRJNA1208460 | cell line |
| GSE269514 | Mus musculus | Aug 08 2024 | PRJNA1122170 | Rod PR |
| GSE315225 | Homo sapiens | Jan 09 2026 | PRJNA1395367 | Heart tissue |
| GSE210492 | Mus musculus | Aug 10 2022 | PRJNA865781 | |
| GSE282671 | Homo sapiens | Feb 23 2026 | PRJNA1189580 | Spleen |

| | | | | |
|---|---|---|---|---|
| GSE298019 | Homo sapiens | Jan 29 2026 | PRJNA1267119 | |
| GSE140545 | Homo sapiens | Aug 01 2020 | PRJNA590151 | duced pluripotent stem cell |
| GSE215192 | Mus musculus | Oct 13 2022 | PRJNA889068 | retina |
| GSE232470 | Mus musculus | May 31 2023 | PRJNA972278 | retina |
| GSE186612 | Mus musculus | Dec 15 2021 | PRJNA774789 | central retinas |
| GSE314526 | Homo sapiens | Jan 01 2026 | PRJNA1391459 | BA44 |
| GSE212248 | Homo sapiens | Dec 31 2022 | PRJNA874710 | |
| GSE307106 | Homo sapiens | Feb 25 2026 | PRJNA1314286 | Bone marrow |
| GSE277221 | Homo sapiens | May 14 2025 | PRJNA1161384 | |
| GSE159918 | Homo sapiens | Jan 16 2026 | PRJNA670880 | |
| GSE245547 | Mus musculus | Jun 06 2024 | PRJNA1028975 | retina |
| GSE319710 | Homo sapiens | Feb 20 2026 | PRJNA1424988 | PBMCs |
| GSE225531 | Rattus norvegicus | Feb 15 2026 | PRJNA936125 | left ventricular |
| GSE316181 | Homo sapiens | Jan 15 2026 | PRJNA1401761 | |
| GSE286327 | Homo sapiens | Jan 10 2026 | PRJNA1208926 | |
| GSE319727 | cillus subtilis subsp. subtilis str. 1 | Feb 18 2026 | PRJNA1425010 | |
| GSE272074 | Homo sapiens;Mus musculus | Dec 01 2024 | | retina |
| GSE186458 | Homo sapiens | Mar 02 2022 | PRJNA773951 | Abdominal Subcut. |
| GSE306150 | Homo sapiens | Feb 05 2026 | PRJNA1309599 | |
| GSE310070 | Homo sapiens | Jan 26 2026 | PRJNA1365094 | plasma |
| GSE300907 | Homo sapiens | Feb 05 2026 | PRJNA1282649 | |
| GSE318259 | Homo sapiens | Feb 09 2026 | PRJNA1416989 | |
| GSE221578 | Homo sapiens | Feb 26 2026 | PRJNA915027 | |
| GSE319045 | Mus musculus | Feb 16 2026 | PRJNA1422016 | Left ventricle |
| GSE281280 | Homo sapiens | Nov 12 2024 | PRJNA1183307 | |
| GSE200443 | Mus musculus | Apr 08 2022 | PRJNA824511 | Retina |
| GSE169102 | Homo sapiens;Mus musculus | May 05 2025 | PRJNA715210 | |
| GSE277216 | Homo sapiens | May 14 2025 | PRJNA1161378 | |
| GSE287346 | Homo sapiens;Mus musculus | Feb 25 2026 | PRJNA1212172 | Bone Marrow |
| GSE156452 | Homo sapiens | Aug 19 2021 | PRJNA658007 | fetal eye |
| GSE229033 | Mus musculus | Dec 14 2023 | PRJNA952615 | Retina |
| GSE245561 | Homo sapiens | Nov 02 2023 | PRJNA1029000 | tinal fibrovascular membrane |
| GSE290709 | Homo sapiens | Feb 26 2026 | PRJNA1229608 | breast |
| GSE316836 | Homo sapiens | Jan 18 2026 | PRJNA1404336 | Enriched blood |
| GSE319736 | Mus musculus | Feb 18 2026 | PRJNA1425031 | Olfactory Epithelium |
| GSE216210 | Mus musculus | Feb 16 2026 | PRJNA892595 | lung |
| GSE186156 | Arabidopsis thaliana | Dec 12 2022 | PRJNA772750 | Seedling |
| GSE244262 | Homo sapiens | Jul 31 2025 | PRJNA1022020 | Retinal Organoids |
| GSE318757 | Mus musculus | Feb 27 2026 | PRJNA1419887 | Tumor-bearing lung |
| GSE191017 | Homo sapiens | Aug 22 2023 | PRJNA789332 | |
| GSE288948 | Mus musculus | Feb 06 2026 | PRJNA1220077 | Bone Marrow |
| GSE306623 | Danio rerio | Feb 24 2026 | PRJNA1311497 | testis |
| GSE287648 | Homo sapiens | Jan 12 2026 | PRJNA1214192 | derived bone marrow organoid |
| GSE318248 | Homo sapiens | Jan 31 2026 | PRJNA1416971 | human melanoma |
| GSE200452 | Homo sapiens | Apr 10 2022 | PRJNA824530 | |
| GSE302899 | Homo sapiens | Jan 20 2026 | PRJNA1292794 | |
| GSE314537 | Homo sapiens | Feb 11 2026 | PRJNA1391471 | |
| GSE201679 | Homo sapiens | Apr 30 2022 | PRJNA832669 | |
| GSE298999 | Homo sapiens | Jan 21 2026 | PRJNA1272089 | kidney |
| GSE277230 | Mus musculus | Jan 01 2026 | PRJNA1161400 | Corpus callosal axon |
| GSE302375 | Homo sapiens;synthetic construc | Feb 26 2026 | PRJNA1290274 | |
| GSE275697 | Homo sapiens | Feb 20 2026 | PRJNA1152583 | . |
| GSE142862 | Homo sapiens | Sep 14 2021 | PRJNA598699 | |
| GSE136360 | Mus musculus | Jul 26 2022 | PRJNA562293 | retina |
| GSE316190 | Mus musculus | Jan 14 2026 | PRJNA1401779 | splenic CD8+ T cells |
| GSE311755 | Mus musculus | Jan 10 2026 | PRJNA1370868 | Lung |
| GSE315556 | Mus musculus | Jan 21 2026 | PRJNA1398260 | skeletal muscle |
| GSE306228 | Anableps anableps | Nov 18 2025 | PRJNA1309719 | Whole retina |
| GSE311114 | Homo sapiens | Feb 25 2026 | PRJNA1368699 | |
| GSE220661 | Homo sapiens | Dec 14 2022 | PRJNA910936 | ll line-derived retinal organoid |
| GSE315608 | Mus musculus | Jan 09 2026 | PRJNA1398320 | Spleen |
| GSE224462 | Homo sapiens | Dec 31 2024 | PRJNA931287 | |
| GSE306737 | Mus musculus | Jan 13 2026 | PRJNA1311997 | sdLN |

| | | | | |
|---|---|---|---|---|
| GSE260530 | Mus musculus | Feb 18 2026 | PRJNA1081950 | |
| GSE248643 | Homo sapiens | May 29 2024 | PRJNA1045303 | retina |
| GSE212896 | Homo sapiens | May 22 2023 | PRJNA877840 | |
| GSE58785 | Homo sapiens | Jul 06 2021 | PRJNA253480 | retinoblastoma |
| GSE196935 | Mus musculus | Feb 20 2022 | PRJNA807898 | retina |
| GSE180815 | Mus musculus | Dec 02 2022 | PRJNA749616 | Retina |
| GSE212107 | Homo sapiens | Apr 23 2023 | PRJNA874035 | Pigmented epithelium; Retina |
| GSE310175 | Saccharomyces cerevisiae | Feb 10 2026 | PRJNA1366920 | |
| GSE311369 | Mus musculus | Dec 10 2025 | PRJNA1369363 | retina |
| GSE315475 | Mus musculus | Jan 03 2026 | PRJNA1397210 | Whole Lung |
| GSE299462 | Homo sapiens | May 20 2026 | PRJNA1274478 | |
| GSE316915 | Escherichia coli DH5[alpha] | Jan 23 2026 | PRJNA1405528 | |
| GSE287516 | Mus musculus | Oct 29 2025 | PRJNA1213133 | E13.5 mouse embryos |
| GSE287998 | Mus musculus | Jan 28 2026 | PRJNA1215434 | Liver |
| GSE274080 | Mus musculus | Jan 31 2026 | PRJNA1144915 | small intestine |
| GSE291436 | Homo sapiens | Jan 27 2026 | PRJNA1233595 | Human embryonic kidney |
| GSE315442 | Homo sapiens | Jan 15 2026 | PRJNA1397163 | Skin |
| GSE317298 | Mus musculus | Jan 22 2026 | PRJNA1406704 | tumor |
| GSE237208 | Peromyscus maniculatus | Oct 01 2023 | PRJNA994174 | Retina |
| GSE297292 | Mus musculus | Jan 01 2026 | PRJNA1263468 | skin |
| GSE216451 | Mus musculus | Feb 16 2026 | PRJNA893642 | lung |
| GSE290998 | Homo sapiens | Mar 11 2025 | PRJNA1231357 | Mammary |
| GSE244562 | Arabidopsis thaliana | Feb 19 2026 | PRJNA1023563 | 14 days old seedlings |
| GSE330159 | Danio rerio | May 12 2026 | PRJNA1462852 | whole embryo |
| GSE315261 | Homo sapiens | Jan 06 2026 | | |
| GSE291215 | Mus musculus | Jan 09 2026 | PRJNA1232258 | Lung |
| GSE300298 | Marchantia polymorpha | Jan 16 2026 | PRJNA1280062 | Thallus |
| GSE294715 | Mus musculus | Feb 23 2026 | PRJNA1251359 | tumor |
| GSE290274 | Homo sapiens | Feb 10 2026 | PRJNA1227119 | |
| GSE286354 | Homo sapiens | Jan 01 2026 | PRJNA1208962 | Blood |
| GSE249754 | Danio rerio | Jun 21 2024 | PRJNA1050286 | retina |
| GSE287148 | Mus musculus | Dec 23 2025 | PRJNA1211493 | organoids |
| GSE241700 | Mus musculus | Aug 29 2024 | PRJNA1009136 | Retina |
| GSE308920 | Mus musculus | Jan 08 2026 | PRJNA1333403 | liver |
| GSE310003 | Homo sapiens | Feb 05 2026 | PRJNA1364988 | |
| GSE285792 | Homo sapiens;Mus musculus | Jan 01 2026 | PRJNA1206093 | |
| GSE316854 | Mus musculus | Jan 22 2026 | PRJNA1405365 | brain |
| GSE314728 | Homo sapiens | Jan 30 2026 | PRJNA1392508 | KG1a cells |
| GSE308917 | Homo sapiens | Jan 01 2026 | PRJNA1333396 | Osteosarcoma |
| GSE153674 | Mus musculus | Oct 23 2020 | PRJNA643586 | |
| GSE287460 | Homo sapiens | Jan 31 2026 | PRJNA1213042 | corneal epithelium |
| GSE310034 | Mus musculus | Jan 26 2026 | PRJNA1365035 | Skeletal muscle |
| GSE298708 | Homo sapiens | Jan 14 2026 | PRJNA1270990 | Liver |
| GSE306114 | Homo sapiens | Jan 08 2026 | PRJNA1309149 | Prostate |
| GSE314237 | Caenorhabditis elegans | Feb 24 2026 | PRJNA1390503 | whole body |
| GSE222387 | Homo sapiens | Jan 12 2023 | PRJNA921857 | retinal epithelium |
| GSE283663 | Homo sapiens | Jan 21 2026 | PRJNA1195006 | |
| GSE261410 | Mus musculus | Feb 17 2026 | PRJNA1086897 | |
| GSE287302 | Mus musculus | Jan 21 2025 | PRJNA1212068 | organoids |
| GSE237892 | Homo sapiens | Dec 11 2023 | PRJNA996995 | |
| GSE310249 | Mus musculus | Jan 06 2026 | PRJNA1365633 | T cells |
| GSE270819 | Mus musculus | Jan 13 2026 | PRJNA1128570 | |
| GSE156257 | Homo sapiens | Aug 11 2021 | PRJNA657203 | |
| GSE315417 | Mus musculus | Jan 05 2026 | PRJNA1397123 | Whole Blood |
| GSE310408 | Homo sapiens | Jan 09 2026 | PRJNA1365902 | Small intestine |
| GSE298334 | Homo sapiens | Feb 21 2026 | PRJNA1268978 | OX9 recombinant protein |
| GSE116106 | Homo sapiens | May 07 2020 | PRJNA477287 | retina |
| GSE282721 | Homo sapiens | Feb 21 2026 | PRJNA1190064 | prostate cancer |
| GSE315375 | Homo sapiens | Jan 02 2026 | PRJNA1396615 | Thyroid tissue |
| GSE301801 | Mus musculus | Feb 15 2026 | | Kidney |
| GSE213104 | Mus musculus | Feb 06 2026 | PRJNA879195 | skin |
| GSE309855 | | Jan 31 2026 | | |
| GSE282209 | Mus musculus | Jan 13 2026 | PRJNA1187588 | pre-implantation embryos |

| | | | | |
|---|---|---|---|---|
| GSE293962 | Homo sapiens | Jan 20 2026 | PRJNA1247501 | Lung fibroblasts |
| GSE294636 | Acidithiobacillus ferrooxidans | Feb 11 2026 | PRJNA1250982 | biofilm |
| GSE283984 | Homo sapiens | Feb 18 2026 | PRJNA1196477 | peripheral blood |
| GSE277424 | Mus musculus | Jan 13 2026 | PRJNA1162152 | FL-BMDC |
| GSE227793 | Homo sapiens | Feb 16 2026 | PRJNA947086 | left cardiac tissue |
| GSE319677 | Mus musculus | Feb 18 2026 | PRJNA1424948 | |
| GSE316977 | Homo sapiens | Jan 22 2026 | PRJNA1405628 | Cell line |
| GSE173447 | Homo sapiens | Mar 03 2023 | PRJNA725539 | |
| GSE291454 | Mus musculus | Jan 14 2026 | PRJNA1233787 | |
| GSE190428 | Homo sapiens | Dec 10 2021 | PRJNA787110 | |
| GSE290648 | Ixodes scapularis | Feb 20 2026 | | salivary glands |
| GSE309691 | Homo sapiens | Jan 05 2026 | PRJNA1336746 | |
| GSE319368 | Mus musculus | Feb 12 2026 | PRJNA1423174 | dorsal brain |
| GSE312067 | Mus musculus | Dec 31 2025 | PRJNA1372129 | retina |
| GSE264286 | Homo sapiens | Jan 28 2026 | PRJNA1101869 | stage III/IV OCCC Tissue |
| GSE304458 | Mus musculus | Feb 10 2026 | PRJNA1301333 | mammary tumor |
| GSE300344 | Homo sapiens | Jan 01 2026 | PRJNA1280124 | prostate |
| GSE139605 | Mus musculus | Oct 29 2020 | PRJNA580470 | optic nerve head |
| GSE286288 | Mus musculus | Feb 17 2026 | PRJNA1208531 | |
| GSE289419 | Rattus norvegicus | Feb 12 2026 | PRJNA1222886 | Colon |
| GSE327855 | Homo sapiens | Apr 18 2026 | PRJNA1452879 | |
| GSE303946 | Mus musculus | Jan 27 2026 | PRJNA1298286 | cell line |
| GSE316531 | Homo sapiens | Jan 21 2026 | PRJNA1403272 | |
| GSE309346 | Homo sapiens | Feb 12 2026 | PRJNA1335212 | asis lesion from colorectal adenocarcinoma |
| GSE262415 | Mus musculus | Jan 21 2026 | PRJNA1091682 | Testis |
| GSE304612 | Homo sapiens | Feb 27 2026 | PRJNA1301922 | Blood cells |
| GSE269598 | Mus musculus | Jan 01 2025 | PRJNA1122599 | eye cup |
| GSE224871 | Mus musculus | Feb 07 2026 | PRJNA932823 | |
| GSE312050 | Rattus norvegicus | Feb 26 2026 | | Mammary microlesions |
| GSE151610 | Homo sapiens | Jun 03 2020 | PRJNA636612 | |
| GSE202540 | Mus musculus | May 12 2022 | PRJNA836532 | blastocyst |
| GSE284546 | Mus musculus | Jan 30 2026 | PRJNA1199505 | Heart |
| GSE308566 | Homo sapiens | Jan 29 2026 | PRJNA1331544 | |
| GSE315945 | Mus musculus | Jan 20 2026 | PRJNA1399992 | Heart |
| GSE289983 | Ixodes scapularis | Feb 19 2026 | PRJNA1225684 | salivary glands |
| GSE157806 | Mus musculus | Sep 12 2020 | PRJNA662876 | retina |
| GSE204808 | Homo sapiens | Sep 01 2022 | PRJNA842375 | |
| GSE274898 | Homo sapiens | Jan 31 2026 | PRJNA1148427 | Liver |
| GSE325477 | Mus musculus | Apr 02 2026 | PRJNA1440255 | retina |
| GSE288073 | Homo sapiens | Jan 21 2026 | PRJNA1215995 | |
| GSE283858 | Homo sapiens | Jan 06 2026 | PRJNA1195920 | |
| GSE316347 | Mus musculus | Jan 13 2026 | PRJNA1402351 | ngineered Colon Assembloids |
| GSE308233 | Mus musculus | Feb 24 2026 | PRJNA1329261 | |
| GSE301206 | Homo sapiens | Feb 12 2026 | PRJNA1284268 | ortic vascular smooth muscle cells |
| GSE303852 | Mus musculus | Feb 25 2026 | PRJNA1297769 | liver |
| GSE315972 | Homo sapiens | Jan 09 2026 | PRJNA1400041 | DLPFC_BA46 |
| GSE291588 | Mus musculus | Jan 12 2026 | PRJNA1234196 | Aorta |
| GSE227350 | Mus musculus | Jan 09 2024 | PRJNA944689 | |
| GSE318878 | Homo sapiens | Feb 09 2026 | PRJNA1420758 | peripheral blood |
| GSE247267 | Homo sapiens | Jul 08 2024 | PRJNA1037018 | Retina |
| GSE255626 | Homo sapiens | Jan 21 2026 | PRJNA1075795 | Brain tissue |
| GSE312994 | Homo sapiens | Jan 31 2026 | PRJNA1378075 | lung |
| GSE292272 | Homo sapiens | Feb 09 2026 | PRJNA1237814 | |
| GSE319188 | Mus musculus | Feb 16 2026 | PRJNA1422527 | |
| GSE162175 | Homo sapiens | Feb 15 2026 | PRJNA680792 | |
| GSE286931 | Homo sapiens | Jan 09 2026 | PRJNA1210566 | |
| GSE313731 | Homo sapiens | Feb 19 2026 | PRJNA1381500 | spleen |
| GSE309159 | Homo sapiens | Jan 30 2026 | PRJNA1334242 | In vitro derived islets |
| GSE281418 | Mus musculus | Feb 24 2026 | PRJNA1183828 | small intestine |
| GSE168908 | Macaca fascicularis | Aug 05 2021 | PRJNA714553 | retina |
| GSE311689 | human metagenome | Jan 30 2026 | PRJNA1370129 | nasopharyngeal swab |
| GSE285304 | Mus musculus | Jan 14 2026 | PRJNA1202196 | Embryo E11 |
| GSE301370 | Homo sapiens | Feb 17 2026 | PRJNA1284868 | |

| | | | | |
|---|---|---|---|---|
| GSE316553 | Homo sapiens | Mar 18 2026 | PRJNA1403376 | PL1 |
| GSE314946 | Saccharomyces cerevisiae | Jan 08 2026 | PRJNA1393400 | |
| GSE287680 | Homo sapiens | Feb 19 2026 | PRJNA1214240 | |
| GSE296165 | Mus musculus | Feb 11 2026 | PRJNA1257954 | heart |
| GSE318280 | Homo sapiens | Feb 11 2026 | PRJNA1418515 | gue squamous cell carcinoma |
| GSE234047 | Homo sapiens | Aug 07 2023 | PRJNA979238 | |
| GSE312880 | Mus musculus | Jan 08 2026 | PRJNA1377547 | C57Bl6 |
| GSE286183 | Mus musculus | Jan 08 2026 | PRJNA1208087 | Back skin |
| GSE139980 | Mus musculus | Dec 08 2020 | PRJNA587773 | retina |
| GSE293307 | Mus musculus | May 12 2026 | PRJNA1243981 | Retina |
| GSE288773 | Homo sapiens | Jan 14 2026 | PRJNA1219231 | Placenta |
| GSE299589 | Homo sapiens | Feb 18 2026 | PRJNA1275025 | In-vitro cell line |
| GSE316470 | Mus musculus | Jan 21 2026 | | |
| GSE278623 | Homo sapiens | Feb 05 2026 | PRJNA1168104 | |
| GSE313170 | Homo sapiens | Jan 21 2026 | PRJNA1378734 | Prostate |
| GSE316447 | Homo sapiens | Jan 16 2026 | PRJNA1402928 | periodontal ligament |
| GSE305067 | Homo sapiens | Feb 25 2026 | PRJNA1304383 | |
| GSE189738 | Danio rerio | May 12 2023 | PRJNA784297 | |
| GSE273286 | Homo sapiens | Feb 23 2026 | PRJNA1141364 | human embryo |
| GSE295594 | Mus musculus | Jan 05 2026 | PRJNA1255227 | nus endothelial cells (SECs) |
| GSE288026 | Homo sapiens | Jan 14 2026 | PRJNA1215735 | |
| GSE317426 | Homo sapiens | Jan 27 2026 | PRJNA1412573 | |
| GSE302851 | Homo sapiens | Jan 20 2026 | PRJNA1292280 | bone marrow |
| GSE316098 | Homo sapiens | Jan 12 2026 | PRJNA1400592 | Cervix |
| GSE287015 | Homo sapiens | Jan 16 2026 | PRJNA1210973 | plasma exosomes |
| GSE315141 | Burkholderia contaminans | Jan 12 2026 | PRJNA1395045 | microbe from plant root |
| GSE237214 | Danio rerio | Oct 01 2023 | PRJNA994180 | Retina |
| GSE160019 | Danio rerio | Jul 31 2021 | PRJNA671650 | |
| GSE307262 | Homo sapiens | Feb 26 2026 | PRJNA1315089 | umbilical cord blood |
| GSE276359 | Homo sapiens | Oct 03 2024 | PRJNA1156779 | |
| GSE319316 | Saccharomyces cerevisiae | Feb 16 2026 | PRJNA1422811 | SK1 |
| GSE201982 | Homo sapiens | Jul 15 2022 | PRJNA833894 | |
| GSE318377 | Mus musculus | Feb 09 2026 | PRJNA1418650 | retina |
| GSE322638 | Anableps anableps | Apr 08 2026 | PRJNA1430790 | Dorsal retina |
| GSE311420 | Mus musculus | Jan 16 2026 | PRJNA1369431 | |
| GSE304860 | Mus musculus | Jan 26 2026 | PRJNA1303082 | Liver |
| GSE172170 | Homo sapiens | Apr 16 2021 | PRJNA722287 | mary retinoblastoma tissue |
| GSE318340 | Homo sapiens | Feb 06 2026 | PRJNA1418599 | Prostate Cancer |
| GSE261052 | Homo sapiens | Feb 10 2026 | PRJNA1084908 | |
| GSE314675 | Homo sapiens | Feb 11 2026 | PRJNA1392428 | |
| GSE306556 | Sus scrofa | Feb 17 2026 | PRJNA1311112 | lung |
| GSE287022 | Homo sapiens | Jan 14 2026 | PRJNA1210993 | Non-Lesional |
| GSE297955 | Mus musculus | Aug 01 2025 | PRJNA1267025 | retina |
| GSE263688 | Mus musculus | Jan 23 2026 | PRJNA1098895 | primary lung tumor |
| GSE127934 | Homo sapiens | Mar 16 2021 | PRJNA525832 | colon tumor distal region |
| GSE319560 | Candida albicans | Feb 17 2026 | PRJNA1423732 | |
| GSE315228 | Homo sapiens | Feb 18 2026 | PRJNA1395377 | rectus femoris |
| GSE146191 | Mus musculus | Aug 06 2021 | PRJNA609690 | |
| GSE306608 | Homo sapiens | Feb 26 2026 | PRJNA1311465 | Kidney PDX |
| GSE299717 | Homo sapiens | Feb 26 2026 | PRJNA1276122 | iPSC-derived cells |
| GSE285928 | Oncorhynchus mykiss | Jan 06 2026 | PRJNA1206997 | liver tissues |
| GSE275394 | Mus musculus | Feb 08 2026 | PRJNA1150598 | retina |
| GSE252676 | Homo sapiens | Jan 01 2026 | PRJNA1061785 | |
| GSE208650 | Homo sapiens;Mus musculus | Oct 10 2022 | PRJNA860553 | |
| GSE292869 | Rattus norvegicus | Jan 01 2026 | PRJNA1241790 | midbrain |
| GSE307376 | Mus musculus | Feb 25 2026 | PRJNA1320549 | |
| GSE318501 | Homo sapiens | Feb 06 2026 | | |
| GSE165112 | Homo sapiens | Jan 20 2021 | PRJNA693278 | |
| GSE287101 | Homo sapiens | Mar 30 2025 | PRJNA1211122 | Retina |
| GSE245055 | Mus musculus | Feb 11 2026 | PRJNA1026869 | Lung |
| GSE291021 | Homo sapiens | Feb 22 2026 | PRJNA1231410 | Tear fluid |
| GSE307669 | Mus musculus | Feb 20 2026 | PRJNA1322494 | Heart |
| GSE303142 | Arabidopsis thaliana | Jan 29 2026 | PRJNA1294232 | Seedlings |

| | | | | |
|---|---|---|---|---|
| GSE315062 | Mus musculus | Feb 18 2026 | PRJNA1394945 | cell line |
| GSE232637 | Homo sapiens;Mus musculus | Aug 21 2025 | PRJNA973084 | Retinal Pigment Epithelial |
| GSE287429 | Homo sapiens | Jan 18 2026 | PRJNA1212740 | anoids from primary breast tumors |
| GSE329865 | Mus musculus | Jun 29 2026 | PRJNA1461971 | retina |
| GSE291509 | Homo sapiens;Mus musculus | Jan 14 2026 | PRJNA1234067 | |
| GSE213213 | Homo sapiens | Feb 23 2026 | PRJNA879759 | |
| GSE294009 | Homo sapiens | Feb 05 2026 | PRJNA1247589 | Skin |
| GSE286157 | Mus musculus | Jan 13 2026 | PRJNA1208026 | Bone marrow |
| GSE319557 | Macaca fascicularis | Feb 20 2026 | PRJNA1423731 | Brain |
| GSE295577 | Caulobacter vibrioides | Feb 25 2026 | PRJNA1255207 | |
| GSE236217 | Homo sapiens;Mus musculus | Jan 01 2026 | PRJNA989274 | cord blood |
| GSE290136 | Homo sapiens | Feb 19 2026 | PRJNA1226249 | alin-fixed paraffin embedded |
| GSE313787 | Mus musculus | Jan 31 2026 | PRJNA1381593 | Quaddel] primary adult APH treated spleen |
| GSE282215 | Mus musculus | Jun 25 2025 | PRJNA1187598 | retina |
| GSE316798 | Mus musculus | Jan 29 2026 | PRJNA1404226 | Brain tissue |
| GSE313298 | Mus musculus | Jan 05 2026 | PRJNA1378899 | lymph node |
| GSE311320 | Homo sapiens | Feb 11 2026 | PRJNA1368942 | EDTA Plasma |
| GSE299892 | Homo sapiens | Jan 21 2026 | PRJNA1277267 | |
| GSE318477 | Caenorhabditis elegans | Feb 05 2026 | PRJNA1418810 | Whole Animal |
| GSE313974 | Fundulus heteroclitus | Jan 07 2026 | PRJNA1387459 | embryo |
| GSE277881 | Mus musculus | Jan 30 2026 | PRJNA1164690 | Lung |
| GSE144168 | Mus musculus | Jan 28 2020 | PRJNA602922 | Retina |
| GSE281817 | Saccharomyces cerevisiae | Feb 12 2026 | PRJNA1185750 | |
| GSE302228 | Candida albicans | Jan 22 2026 | PRJNA1289683 | |
| GSE228436 | Homo sapiens | Jun 30 2024 | PRJNA949697 | differentiated iPSC cells |
| GSE184815 | Mus musculus | Jan 01 2022 | PRJNA766600 | whole retina |
| GSE282222 | Mus musculus | Dec 09 2025 | PRJNA1187608 | Retina |
| GSE128134 | Homo sapiens | Jul 11 2020 | PRJNA526530 | |
| GSE236220 | Mus musculus | Jul 15 2023 | PRJNA989279 | RPE/choroid |
| GSE319421 | Mus musculus | Feb 28 2026 | PRJNA1423264 | |
| GSE259286 | Homo sapiens | Feb 19 2026 | PRJNA1080588 | |
| GSE282463 | Homo sapiens | Feb 18 2026 | PRJNA1188629 | |
| GSE315528 | Homo sapiens | Jan 31 2026 | PRJNA1398200 | |
| GSE152474 | Mus musculus | Jun 16 2020 | PRJNA639510 | |
| GSE223498 | Mus musculus | Jan 31 2023 | PRJNA926592 | |
| GSE298714 | Homo sapiens | Jan 26 2026 | PRJNA1270998 | lung adenocarcinoma |
| GSE80686 | Arabidopsis thaliana | Jan 31 2026 | PRJNA319670 | seedlings |
| GSE302469 | Homo sapiens | Feb 25 2026 | PRJNA1291043 | Lung |
| GSE287601 | Homo sapiens | Oct 05 2025 | PRJNA1213800 | eye |
| GSE318201 | Canis lupus familiaris | Jan 31 2026 | PRJNA1416901 | ted bone primary osteosarcoma |
| GSE192816 | Homo sapiens | Oct 28 2022 | PRJNA793257 | |
| GSE306375 | Rattus norvegicus | Nov 17 2025 | PRJNA1310547 | Retina |
| GSE275494 | Homo sapiens | Dec 02 2024 | PRJNA1151087 | |
| GSE307476 | Homo sapiens | Feb 11 2026 | PRJNA1321789 | |
| GSE314061 | Danio rerio | Dec 22 2025 | PRJNA1389905 | Whole larvae |
| GSE300284 | Homo sapiens | Jun 24 2025 | | |
| GSE295515 | Drosophila melanogaster | Jan 08 2026 | | ovary |
| GSE266083 | Homo sapiens | Sep 02 2024 | PRJNA1105856 | |
| GSE315991 | Mus musculus | Jan 15 2026 | PRJNA1400074 | liver |
| GSE164644 | Homo sapiens | Jan 13 2021 | PRJNA691476 | |
| GSE319944 | Homo sapiens | Feb 18 2026 | PRJNA1425563 | uro-2a subcutaneous tumor |
| GSE274640 | Homo sapiens | Jan 14 2026 | PRJNA1147313 | |
| GSE279314 | Macaca mulatta | Feb 06 2026 | PRJNA1171842 | lymph node |
| GSE238259 | Mus musculus | Jan 06 2026 | PRJNA998476 | Ovary |
| GSE184570 | Anolis sagrei | Jun 21 2022 | PRJNA765051 | |
| GSE327286 | Mus musculus | Jul 02 2026 | PRJNA1450117 | retina |
| GSE278899 | Saccharomyces cerevisiae | Feb 16 2026 | PRJNA1169376 | |
| GSE198932 | Bos taurus | Mar 14 2025 | PRJNA817534 | muscle |
| GSE285771 | Mus musculus | Jan 07 2026 | | pancreas |
| GSE300997 | Mus musculus | Jan 01 2026 | PRJNA1282974 | P60 cortex |
| GSE296351 | Homo sapiens | Feb 10 2026 | PRJNA1258974 | |
| GSE312325 | Mus musculus | Jan 07 2026 | PRJNA1373027 | Substantia Nigra |
| GSE325621 | Danio rerio | Apr 02 2026 | PRJNA1441374 | retina |

| | | | | |
|---|---|---|---|---|
| GSE255506 | Mus musculus | Feb 05 2026 | PRJNA1075128 | |
| GSE305506 | Mus musculus | Feb 18 2026 | PRJNA1306116 | Tumor |
| GSE328375 | Homo sapiens | May 17 2026 | PRJNA1454870 | retinal organoid |
| GSE296366 | Homo sapiens | Feb 09 2026 | PRJNA1259000 | |
| GSE196106 | Homo sapiens;Mus musculus | Jul 13 2022 | PRJNA803274 | retina |
| GSE315959 | Homo sapiens | Feb 25 2026 | PRJNA1400010 | Prostate |
| GSE184547 | Mus musculus | May 06 2022 | PRJNA765016 | |
| GSE275174 | Mus musculus | Feb 24 2026 | PRJNA1149734 | cortex |
| GSE309995 | Homo sapiens | Jan 26 2026 | PRJNA1364977 | |
| GSE302496 | Mus musculus | Jan 05 2026 | PRJNA1291080 | male liver |
| GSE305253 | Homo sapiens | Feb 11 2026 | PRJNA1304928 | lesion |
| GSE301050 | Homo sapiens | Feb 24 2026 | PRJNA1283149 | |
| GSE303395 | Mus musculus | Feb 05 2026 | PRJNA1295195 | Lung |
| GSE297618 | Mus musculus | Feb 05 2026 | PRJNA1265272 | Pancreatic cancer |
| GSE210402 | Homo sapiens | Aug 03 2022 | PRJNA865654 | |
| GSE292046 | Homo sapiens | Feb 10 2026 | PRJNA1236382 | glioblastoma tumor |
| GSE166175 | Homo sapiens | Jul 26 2021 | PRJNA699531 | Retina |
| GSE214040 | Homo sapiens | Mar 13 2023 | PRJNA883672 | Retina (female) |
| GSE162376 | Mus musculus | Dec 01 2020 | PRJNA681624 | retina |
| GSE212186 | Mus musculus | Nov 30 2022 | PRJNA874434 | bulk retina |
| GSE309438 | Mus musculus | Feb 28 2026 | PRJNA1336082 | |
| GSE230606 | Homo sapiens | Jan 15 2026 | PRJNA961900 | PBMC |
| GSE296527 | Homo sapiens | Jan 04 2026 | PRJNA1259898 | |
| GSE295870 | Procambarus virginalis | Jan 29 2026 | PRJNA1256518 | Hemolymph |
| GSE254444 | Homo sapiens | Feb 11 2026 | PRJNA1070590 | kidney |
| GSE279237 | Homo sapiens | Feb 19 2025 | PRJNA1171451 | |
| GSE289678 | Homo sapiens | Feb 18 2026 | PRJNA1223928 | |
| GSE104593 | Mus musculus;Rattus norvegicus | Oct 01 2022 | PRJNA413143 | Retina |
| GSE158473 | Mus musculus | Jan 01 2026 | PRJNA665370 | ovary |
| GSE316767 | Rattus norvegicus | Feb 16 2026 | PRJNA1404170 | Nucleus pulposus |
| GSE298482 | Homo sapiens | Jan 26 2026 | PRJNA1269535 | kidney tumor |
| GSE296272 | Mus musculus | Jan 05 2026 | PRJNA1258765 | mall intestinal epithelium |
| GSE286294 | Mus musculus | Feb 17 2026 | PRJNA1208548 | |
| GSE312719 | Homo sapiens | Jan 27 2026 | PRJNA1375772 | |
| GSE318806 | Homo sapiens | Feb 11 2026 | PRJNA1420657 | PBMCs |
| GSE310852 | Homo sapiens | Jan 30 2026 | PRJNA1367173 | |
| GSE310913 | Homo sapiens | Jan 30 2026 | PRJNA1367523 | |
| GSE222331 | Homo sapiens | Jan 02 2026 | PRJNA919030 | |
| GSE311781 | Rattus norvegicus | Feb 05 2026 | PRJNA1370900 | ellar preduncle (ccp) and Inferior Olive |
| GSE317358 | Arabis alpina | Jan 31 2026 | PRJNA1406785 | meristem |
| GSE202016 | Homo sapiens | Jul 03 2022 | PRJNA834054 | |
| GSE278317 | Homo sapiens | Jan 31 2026 | PRJNA1166320 | |
| GSE297430 | Homo sapiens | Jan 13 2026 | PRJNA1263832 | |
| GSE288247 | Homo sapiens | Jan 27 2026 | PRJNA1216620 | |
| GSE315582 | Mus musculus | Jan 10 2026 | PRJNA1398283 | guinal white adipose tissue |
| GSE254267 | Homo sapiens | Jan 28 2026 | PRJNA1069108 | |
| GSE287858 | Mus musculus | Jan 23 2026 | PRJNA1214819 | islet beta cell |
| GSE263001 | Homo sapiens | Feb 15 2026 | PRJNA1095430 | Cell line |
| GSE285071 | Mus musculus | Jan 01 2026 | PRJNA1201145 | |
| GSE316339 | Mus musculus | Jan 18 2026 | PRJNA1402339 | Spleen |
| GSE291294 | Homo sapiens | Jan 05 2026 | PRJNA1232734 | Primary Human HSPC |
| GSE290088 | Mus musculus | Jan 19 2026 | PRJNA1226196 | Ovary |
| GSE266029 | Homo sapiens | Sep 25 2025 | PRJNA1105368 | 2D culture |
| GSE285046 | Homo sapiens | Feb 21 2026 | | None |
| GSE213937 | Mus musculus | Jul 01 2023 | PRJNA883230 | |
| GSE318783 | Mus musculus | Feb 18 2026 | PRJNA1420540 | |
| GSE309579 | Homo sapiens | Feb 16 2026 | PRJNA1336287 | brain tumor-derived stem cell line |
| GSE298789 | Homo sapiens | Feb 18 2026 | PRJNA1271109 | non-small cell lung cancer |
| GSE266254 | Mus musculus | Jan 08 2026 | PRJNA1106453 | colon |
| GSE305553 | Mus musculus | Feb 24 2026 | PRJNA1306501 | |
| GSE142881 | Mus musculus | Dec 29 2020 | PRJNA598726 | |
| GSE281679 | Homo sapiens | Jan 21 2026 | PRJNA1185214 | |
| GSE210543 | Homo sapiens | Jan 19 2023 | PRJNA866128 | |

| | | | | |
|---|---|---|---|---|
| GSE317031 | Mus musculus | Feb 25 2026 | PRJNA1405695 | Spleen |
| GSE297246 | Mus musculus | Feb 12 2026 | PRJNA1263390 | kidney |
| GSE301880 | Homo sapiens | Jan 24 2026 | PRJNA1287695 | human 4C embryo |
| GSE266415 | dulatus;Meriones unguiculatus;I | Sep 02 2025 | PRJNA1107131 | Retina |
| GSE316732 | Mus musculus | Jan 16 2026 | PRJNA1404104 | liver |
| GSE179760 | Homo sapiens | Feb 10 2025 | PRJNA744910 | paraffin embedded lymph node biopsy |
| GSE312253 | Mus musculus | Feb 24 2026 | PRJNA1372729 | Cochlea - Organ of Corti |
| GSE280618 | Homo sapiens | Feb 11 2026 | PRJNA1179496 | Cell Line |
| GSE311695 | Homo sapiens | Feb 20 2026 | PRJNA1370145 | organoid |
| GSE266377 | Mus musculus | Jul 30 2025 | PRJNA1106854 | Retina |
| GSE316050 | Mus musculus | Feb 17 2026 | PRJNA1400523 | AML cell line |
| GSE211515 | Mus musculus | Nov 29 2022 | PRJNA870585 | optic nerve head |
| GSE281433 | Homo sapiens | Jan 22 2026 | PRJNA1183854 | ovary |
| GSE318864 | Homo sapiens | Feb 13 2026 | PRJNA1420738 | |
| GSE160977 | Homo sapiens | Jan 16 2026 | PRJNA675041 | reatic Ductal Adenocarcinoma |
| GSE279537 | Penicillium expansum | Jan 01 2026 | PRJNA1173215 | |
| GSE255158 | ndida albicans;Candidozyma au | Jan 12 2026 | | |
| GSE312264 | Rattus norvegicus | Jan 13 2026 | PRJNA1372742 | Jejunum |
| GSE186589 | Homo sapiens;Mus musculus | Mar 13 2023 | PRJNA774649 | |
| GSE300458 | Homo sapiens | Feb 21 2026 | PRJNA1280993 | |
| GSE316578 | Crypthecodinium cohnii | Jan 20 2026 | PRJNA1403410 | |
| GSE313205 | Mus musculus | Jan 31 2026 | PRJNA1378789 | bone marrow |
| GSE192511 | Danio rerio | Dec 29 2021 | PRJNA791684 | embryos (only retina cells tagged) |
| GSE332621 | Microcebus murinus | May 31 2026 | PRJNA1468687 | Retina |
| GSE135133 | Homo sapiens | Jan 22 2020 | PRJNA557496 | Left |
| GSE160784 | Homo sapiens | Nov 17 2020 | PRJNA674413 | |
| GSE288888 | Homo sapiens | Feb 06 2026 | PRJNA1219679 | Hippocampus on autopsy |
| GSE171426 | Danio rerio | Oct 05 2021 | PRJNA719543 | |
| GSE311098 | Homo sapiens | Feb 26 2026 | PRJNA1368670 | Kidney PDX |
| GSE238240 | Danio rerio | Oct 02 2023 | PRJNA998413 | brain |
| GSE223634 | Homo sapiens | Jan 28 2023 | PRJNA927001 | |
| GSE196355 | Homo sapiens | Apr 07 2022 | PRJNA804510 | |
| GSE206013 | Mus musculus | May 31 2023 | PRJNA848800 | retina |
| GSE245784 | Homo sapiens | Jul 24 2024 | PRJNA1029963 | |
| GSE299957 | Homo sapiens;Mus musculus | Jan 14 2026 | PRJNA1277709 | |
| GSE300500 | Mus musculus | Jun 24 2025 | PRJNA1281061 | retina |
| GSE297154 | Rattus norvegicus | Jan 01 2026 | PRJNA1262899 | Kidney |
| GSE308554 | Mus musculus | Jan 13 2026 | PRJNA1330873 | spleen |
| GSE165830 | Mus musculus | Dec 22 2021 | PRJNA697990 | retina |
| GSE287823 | Rattus norvegicus | Jan 23 2026 | PRJNA1214783 | hippocampal |
| GSE247323 | Mus musculus | Jan 21 2026 | PRJNA1037105 | liver |
| GSE226456 | Mus musculus | Jul 26 2023 | PRJNA940371 | retina |
| GSE247314 | Mus musculus | Nov 08 2023 | PRJNA1037094 | Retina |
| GSE293360 | Homo sapiens | Jan 20 2026 | PRJNA1244253 | Lung |
| GSE189768 | Mus musculus | Feb 12 2026 | PRJNA784446 | |
| GSE254734 | Homo sapiens | Dec 06 2024 | PRJNA1071590 | |
| GSE292463 | Homo sapiens | Feb 19 2026 | PRJNA1238812 | Retinal Organoid |
| GSE308563 | Homo sapiens | May 22 2026 | PRJNA1330891 | cell line |
| GSE303860 | Homo sapiens | Jan 30 2026 | PRJNA1297771 | |
| GSE305037 | Homo sapiens | Feb 27 2026 | PRJNA1304277 | Blood cells |
| GSE146884 | Homo sapiens | Jul 08 2020 | PRJNA612295 | |
| GSE309502 | Arabidopsis thaliana | Feb 28 2026 | PRJNA1336183 | whole seedlings |
| GSE261697 | Homo sapiens | Jan 31 2026 | PRJNA1088437 | circulating blood |
| GSE183684 | Homo sapiens | Nov 16 2021 | PRJNA761679 | |
| GSE312469 | Mus musculus | Feb 12 2026 | PRJNA1373557 | cardiac tissue |
| GSE210745 | Homo sapiens | Apr 18 2023 | PRJNA867358 | melanoma cell lines |
| GSE279064 | Homo sapiens | Feb 17 2026 | PRJNA1170287 | |
| GSE176495 | Homo sapiens | Sep 27 2021 | PRJNA736434 | |
| GSE141800 | Homo sapiens | Apr 14 2021 | PRJNA594745 | |
| GSE315863 | Mus musculus | Jan 13 2026 | PRJNA1399453 | lung |
| GSE304308 | Mus musculus | Jan 25 2026 | PRJNA1300340 | dendrocyte isolated from brain |
| GSE290685 | Mus musculus | Feb 23 2026 | PRJNA1229542 | mouse spleen |
| GSE317728 | Mus musculus | Jan 29 2026 | PRJNA1415106 | orsal root ganglion (L4-L5) |

| | | | | |
|---|---|---|---|---|
| GSE234661 | Danio rerio | Jul 03 2023 | PRJNA982720 | Whole retina |
| GSE316534 | Homo sapiens | Jan 19 2026 | PRJNA1403283 | organoids |
| GSE289349 | Homo sapiens | Feb 11 2026 | PRJNA1222542 | Human meningioma |
| GSE247555 | Mus musculus | Feb 05 2026 | PRJNA1039709 | kidney |
| GSE316749 | Homo sapiens | Jan 17 2026 | PRJNA1404127 | lung |
| GSE297722 | Homo sapiens | Mar 05 2026 | PRJNA1265961 | ll line derived retinal spheroid |
| GSE158142 | Danio rerio | Sep 18 2020 | PRJNA664124 | brain |
| GSE263471 | Homo sapiens | Jun 30 2026 | PRJNA1097600 | retina |
| GSE190387 | Homo sapiens | Dec 10 2021 | PRJNA787111 | |
| GSE295869 | Procambarus virginalis | Jan 29 2026 | PRJNA1256519 | Hemolymph |
| GSE303298 | Rattus norvegicus | Jan 21 2026 | PRJNA1294785 | Brainstem |
| GSE307584 | Mus musculus | Jan 30 2026 | PRJNA1322343 | |
| GSE243992 | Mus musculus | Aug 16 2024 | | retina |
| GSE162647 | Mus musculus | Mar 01 2021 | PRJNA682538 | retina |
| GSE285354 | Toxoplasma gondii | Jan 01 2026 | PRJNA1203222 | |
| GSE310584 | Homo sapiens | Jan 16 2026 | PRJNA1366534 | PDA Cell Lines |
| GSE277911 | Homo sapiens | Jun 06 2025 | PRJNA1164753 | |
| GSE274446 | Homo sapiens | Feb 15 2026 | PRJNA1146437 | |
| GSE284998 | Homo sapiens | Jan 21 2026 | PRJNA1200761 | |
| GSE299935 | Danio rerio | Jan 31 2026 | PRJNA1277581 | retina |
| GSE294292 | Mus musculus | Jan 01 2026 | PRJNA1248951 | |
| GSE293457 | Homo sapiens | Mar 22 2026 | PRJNA1244739 | retinal organoid |
| GSE314974 | Danio rerio | Feb 16 2026 | PRJNA1393358 | ad with eyes, 3-5 heads combined |
| GSE263018 | Homo sapiens | Dec 15 2024 | PRJNA1095460 | Gastric Fundus (corpus) |
| GSE304003 | Homo sapiens | Jan 01 2026 | PRJNA1298742 | testis |
| GSE266030 | Homo sapiens | Sep 25 2025 | PRJNA1105370 | 2D culture |
| GSE319586 | Homo sapiens | Feb 28 2026 | PRJNA1423791 | |
| GSE316608 | Mus musculus | Jan 21 2026 | PRJNA1403462 | Ileum |
| GSE314943 | Mus musculus | Jan 30 2026 | PRJNA1393397 | lung |
| GSE243308 | Mus musculus | Jan 31 2026 | PRJNA1017789 | Dorsal root ganglia |
| GSE300555 | Mus musculus | Jan 21 2026 | PRJNA1281169 | hippocampus |
| GSE301749 | Mus musculus | Jan 28 2026 | PRJNA1287517 | spinal cord |
| GSE305055 | Mus musculus | Feb 25 2026 | PRJNA1304347 | e embryonic stem cells, mESC |
| GSE165865 | Homo sapiens | Feb 01 2021 | PRJNA698369 | |
| GSE301256 | Mus musculus | Feb 26 2026 | PRJNA1284341 | lung |
| GSE288776 | Mus musculus | Jan 21 2026 | PRJNA1219238 | Lung |
| GSE304886 | Mus musculus | Feb 13 2026 | PRJNA1303150 | Small intestine |
| GSE297740 | Homo sapiens | Jan 29 2026 | PRJNA1266080 | heral Blood Mononuclear Cell |
| GSE276864 | Mus musculus | Jan 15 2025 | PRJNA1159291 | retina |
| GSE326388 | Mus musculus | Apr 24 2026 | PRJNA1445194 | retina |
| GSE279006 | Mus musculus | Feb 20 2026 | PRJNA1169906 | heart ventricle |
| GSE310490 | Homo sapiens | Jan 01 2026 | PRJNA1366430 | |
| GSE171232 | Mus musculus | Apr 01 2022 | PRJNA718835 | retina |
| GSE316049 | Mus musculus | Jan 14 2026 | PRJNA1400520 | Spleen |
| GSE308415 | Mus musculus | Jan 22 2026 | PRJNA1330373 | |
| GSE300441 | Homo sapiens | Feb 17 2026 | | |
| GSE228485 | Mus musculus | Apr 10 2023 | PRJNA950035 | |
| GSE256098 | Mus musculus | Feb 18 2026 | PRJNA1078248 | |
| GSE162147 | Sus scrofa | Apr 20 2021 | | Adductores |
| GSE304120 | Homo sapiens | Feb 11 2026 | PRJNA1299296 | |
| GSE164910 | Homo sapiens | Jul 19 2022 | PRJNA692437 | FFPE |
| GSE297268 | Homo sapiens | Feb 15 2026 | PRJNA1263425 | Lung |
| GSE223775 | Mus musculus | Jan 04 2026 | PRJNA928353 | |
| GSE288100 | Mus musculus | Mar 01 2025 | PRJNA1216191 | Whole Retina |
| GSE192508 | Mus musculus | Aug 16 2022 | PRJNA791673 | Retina |
| GSE311848 | Homo sapiens | Jan 01 2026 | PRJNA1371624 | Cell line |
| GSE233793 | Mus musculus | Aug 01 2023 | PRJNA978001 | retina |
| GSE218539 | Homo sapiens | Jan 05 2026 | PRJNA904269 | donor heart |
| GSE283592 | Homo sapiens | Feb 17 2026 | PRJNA1194583 | Osteosarcoma |
| GSE269537 | Mus musculus | Feb 13 2026 | PRJNA1122220 | lung |
| GSE315158 | Oryza sativa | Jan 16 2026 | PRJNA1395069 | leaf |
| GSE287513 | Mus musculus | Jan 01 2026 | | Lung tumor |
| GSE148373 | Macaca fascicularis | May 19 2020 | PRJNA624096 | caque trabecular meshwork |

| | | | | |
|---|---|---|---|---|
| GSE315447 | Mus musculus | Feb 24 2026 | PRJNA1397169 | Cortex |
| GSE298406 | Mus musculus | Feb 05 2026 | PRJNA1269097 | etal Gut (stomach to anus) |
| GSE94982 | Mus musculus | Mar 10 2020 | PRJNA375080 | Brain |
| GSE297297 | Mus musculus | Jan 02 2026 | PRJNA1263470 | Heart |
| GSE249172 | Mus musculus | Feb 18 2026 | PRJNA1047591 | ureter |
| GSE141659 | Homo sapiens | Dec 28 2022 | PRJNA594319 | stomach |
| GSE276422 | Mus musculus | Jan 20 2026 | PRJNA1157330 | ain (Cortex, TBI insult area) |
| GSE310625 | Homo sapiens | Jan 05 2026 | PRJNA1366598 | Peripheral blood |
| GSE302019 | Mus musculus | Jan 06 2026 | | lung |
| GSE252019 | Catharanthus roseus | Feb 18 2026 | PRJNA1057131 | leaf |
| GSE212660 | Homo sapiens | Sep 07 2022 | PRJNA876579 | Retina |
| GSE240612 | Mus musculus | Jan 01 2026 | PRJNA1004308 | |
| GSE291366 | Mus musculus | Jan 14 2026 | PRJNA1233250 | |
| GSE307031 | Mus musculus | Sep 05 2025 | PRJNA1314146 | Retina |
| GSE248359 | Mus musculus | Jan 19 2026 | PRJNA1043588 | |
| GSE273884 | Mus musculus | Jan 14 2026 | PRJNA1143546 | Liver |
| GSE283238 | Mus musculus | Jan 21 2026 | PRJNA1192711 | Pancreas |
| GSE310170 | Homo sapiens | Jan 20 2026 | PRJNA1365229 | lung cancer |
| GSE253550 | Mus musculus | Jan 18 2026 | PRJNA1066274 | Postnatal day P7 |
| GSE161734 | Mus musculus | Feb 26 2021 | PRJNA679271 | |
| GSE141413 | Mus musculus | Jan 01 2022 | PRJNA593395 | |
| GSE313938 | Bos taurus | Feb 18 2026 | PRJNA1387392 | Peripheral Blood |
| GSE58780 | Homo sapiens | Jul 06 2021 | PRJNA253482 | retinoblastoma |
| GSE262803 | Mus musculus | Jan 05 2026 | PRJNA1093413 | |
| GSE290307 | Mus musculus | Feb 11 2026 | PRJNA1227452 | Liver |
| GSE286179 | Homo sapiens | Feb 07 2026 | PRJNA1208070 | |
| GSE306611 | Caenorhabditis elegans | Feb 26 2026 | PRJNA1311460 | Whole organism lysate |
| GSE285931 | | Jan 31 2026 | | |
| GSE214685 | Homo sapiens | Oct 07 2022 | PRJNA886684 | |
| GSE287889 | Homo sapiens | Jan 17 2026 | PRJNA1214861 | |
| GSE164608 | Homo sapiens | Jan 10 2026 | PRJNA691325 | |
| GSE308970 | Nothobranchius furzeri | Jan 04 2026 | | gut |
| GSE317696 | Homo sapiens | Jan 29 2026 | PRJNA1414663 | Gonad |
| GSE247696 | Mus musculus | Dec 01 2024 | PRJNA1040182 | retina |
| GSE199713 | Mus musculus | Apr 20 2022 | PRJNA821222 | Brain |
| GSE287118 | Homo sapiens | Jan 30 2026 | PRJNA1211145 | gastrocnemius muscle |
| GSE275692 | Mus musculus | Jan 04 2026 | | Glioblastoma |
| GSE286304 | Rhodotorula toruloides | Feb 07 2026 | PRJNA1208580 | |
| GSE196824 | Mus musculus | Jun 24 2022 | PRJNA807380 | |
| GSE290213 | Homo sapiens | Feb 25 2026 | PRJNA1226580 | Prostate |
| GSE283633 | Mus musculus | Jan 07 2026 | PRJNA1194952 | bone |
| GSE217516 | Homo sapiens | Feb 05 2026 | PRJNA899462 | ronchoalveolar lavage fluid |
| GSE315564 | Rattus norvegicus | Jan 10 2026 | PRJNA1398264 | brain tissues |
| GSE316833 | Mus musculus | Jan 21 2026 | PRJNA1404318 | Paw |
| GSE256144 | Homo sapiens | Feb 20 2026 | PRJNA1078325 | |
| GSE290571 | Mus musculus | Jan 15 2026 | PRJNA1228641 | peritoneal cavity |
| GSE204715 | Mus musculus | May 31 2022 | PRJNA842003 | retinal |
| GSE291333 | Homo sapiens | Feb 18 2026 | PRJNA1232855 | |
| GSE318613 | Macaca mulatta | Feb 06 2026 | PRJNA1419313 | blood |
| GSE296978 | Mus musculus | Jan 13 2026 | PRJNA1262312 | liver |
| GSE215961 | Mus musculus | Feb 18 2026 | PRJNA891351 | |
| GSE270908 | Mus musculus | Jan 12 2026 | PRJNA1128812 | spleen |
| GSE319110 | Homo sapiens | Feb 11 2026 | PRJNA1422111 | n cervical adenocarcinoma cell |
| GSE182211 | caca fascicularis;Macaca nemest | Sep 08 2021 | PRJNA755316 | |
| GSE282071 | rabidopsis thaliana;Brassica nap | Feb 06 2026 | PRJNA1186945 | root |
| GSE297088 | Dermacentor andersoni | Feb 20 2026 | PRJNA1262496 | salivary glands |
| GSE303505 | Mus musculus | Feb 13 2026 | PRJNA1295536 | Mammary fat pad |
| GSE292097 | Mus musculus | Aug 14 2025 | PRJNA1236858 | Retina |
| GSE306994 | Dehalococcoides mccartyi CBDB1 | Jan 28 2026 | PRJNA1314092 | |
| GSE314326 | Mus musculus | Apr 01 2026 | PRJNA1390720 | Retina |
| GSE284180 | Homo sapiens | Dec 13 2024 | PRJNA1197514 | Human fetal retinosphere |
| GSE307731 | Mus musculus | Jan 28 2026 | PRJNA1327228 | Lung |
| GSE286245 | Mus musculus | Jan 13 2025 | PRJNA1208478 | |

| | | | | |
|---|---|---|---|---|
| GSE318459 | Homo sapiens | Feb 05 2026 | PRJNA1418765 | |
| GSE287059 | Homo sapiens | Jan 15 2026 | PRJNA1211077 | cell culture |
| GSE246945 | Homo sapiens | Sep 30 2025 | PRJNA1035142 | |
| GSE154197 | Homo sapiens | Jul 11 2021 | PRJNA645398 | |
| GSE291466 | Mus musculus | Feb 10 2026 | PRJNA1233810 | |
| GSE307053 | Mus musculus | Feb 11 2026 | PRJNA1314182 | nfrarenal abdominal aorta |
| GSE317924 | Mus musculus | Jan 30 2026 | PRJNA1415828 | hippocampus microglia |
| GSE314473 | Homo sapiens | Jan 26 2026 | PRJNA1391374 | skin |
| GSE272674 | Mus musculus | Jan 07 2025 | PRJNA1137943 | Optic cup |
| GSE190667 | Mus musculus | May 06 2022 | PRJNA787937 | |
| GSE282724 | Mus musculus | Feb 21 2026 | PRJNA1190075 | Prostate |
| GSE255689 | Mus musculus | Feb 10 2026 | PRJNA1076168 | lung |
| GSE152172 | Mus musculus | Sep 14 2020 | PRJNA638545 | placenta |
| GSE319279 | Homo sapiens | Feb 20 2026 | PRJNA1422743 | icrovascular endothelial cell line |
| GSE249004 | Callithrix jacchus | Dec 26 2023 | PRJNA1046999 | Retina |
| GSE302470 | Mus musculus | Feb 06 2026 | PRJNA1291070 | |
| GSE122680 | Danio rerio | May 19 2020 | PRJNA506002 | whole retina |
| GSE293873 | Homo sapiens | Feb 11 2026 | PRJNA1246634 | |
| GSE315253 | Mus musculus | Jan 26 2026 | PRJNA1395515 | |
| GSE268420 | Mus musculus | Mar 19 2025 | PRJNA1117071 | retina |
| GSE273054 | Homo sapiens | Jan 16 2026 | PRJNA1139841 | human coronary artery |
| GSE326554 | Homo sapiens | Apr 05 2026 | PRJNA1445881 | Retinal organoid |
| GSE310753 | Mus musculus | Jan 09 2026 | PRJNA1367027 | erior dorsal cerebellar vermis |
| GSE199013 | Homo sapiens | Jul 11 2022 | PRJNA817894 | Right Eye |
| GSE291210 | Mus musculus | Feb 11 2026 | PRJNA1232208 | FFPE Embedded Tumor |
| GSE275834 | Homo sapiens | Jan 13 2026 | PRJNA1153133 | Human brain |
| GSE253344 | Homo sapiens | Jan 16 2025 | PRJNA1065372 | |
| GSE309944 | Mus musculus | Feb 25 2026 | PRJNA1364022 | forelimbs |
| GSE300782 | Homo sapiens | Feb 12 2026 | | spinal cord |
| GSE293844 | Homo sapiens | Jan 28 2026 | PRJNA1246574 | esophagus |
| GSE160141 | Danio rerio | Oct 27 2020 | PRJNA672094 | Retina |
| GSE306126 | Homo sapiens | Jan 14 2026 | PRJNA1309494 | |
| GSE272002 | Homo sapiens | Jan 20 2026 | PRJNA1134606 | |
| GSE249751 | Danio rerio | Jun 21 2024 | PRJNA1050282 | eye |
| GSE319751 | Spongilla lacustris | Feb 18 2026 | PRJNA1425054 | Whole body |
| GSE263145 | Mus musculus | Feb 11 2026 | PRJNA1095977 | Bone marrow |
| GSE293984 | Homo sapiens | Nov 18 2025 | PRJNA1247528 | Retinal organoid |
| GSE289362 | Homo sapiens | May 14 2025 | PRJNA1222786 | Lung |
| GSE298487 | Homo sapiens | Jan 14 2026 | PRJNA1269540 | |
| GSE305420 | Homo sapiens | Jan 31 2026 | PRJNA1305502 | |
| GSE310857 | Homo sapiens | Feb 20 2026 | PRJNA1367190 | cell line |
| GSE205797 | Mus musculus | Feb 10 2026 | PRJNA847618 | |
| GSE312203 | Neisseria gonorrhoeae | Jan 12 2026 | PRJNA1372663 | |
| GSE153786 | Homo sapiens | Aug 20 2020 | PRJNA658094 | Osteosarcoma cells |
| GSE162344 | Mus musculus | Mar 08 2021 | PRJNA681565 | |
| GSE171764 | Homo sapiens | Jan 31 2026 | PRJNA720832 | serum |
| GSE212183 | Mus musculus | Oct 19 2022 | PRJNA874429 | Retina |
| GSE316528 | Homo sapiens | Jan 20 2026 | PRJNA1403279 | Uterus; Cervix |
| GSE301614 | Homo sapiens | Jan 28 2026 | PRJNA1285798 | bladder cancer (SCC) |
| GSE220187 | Mus musculus | Dec 06 2022 | PRJNA909201 | Eye |
| GSE290386 | Mus musculus | Jan 06 2026 | PRJNA1227867 | Ear skin tissue |
| GSE271251 | Mus musculus | Nov 20 2024 | PRJNA1130529 | Eye |
| GSE232948 | Homo sapiens;Pan troglodytes | May 23 2023 | PRJNA974466 | |
| GSE305375 | Homo sapiens | Jan 31 2026 | PRJNA1305427 | |
| GSE317549 | Homo sapiens | Jan 31 2026 | PRJNA1414442 | Tumor |
| GSE219273 | Danio rerio | Dec 31 2024 | PRJNA907845 | whole larvae |
| GSE266472 | Mus musculus | Aug 01 2024 | PRJNA1107245 | suprachiasmatic nucleus |
| GSE199792 | Mus musculus | Jan 17 2023 | PRJNA821499 | retina |
| GSE196103 | Mus musculus | Jul 13 2022 | PRJNA803304 | retina |
| GSE296363 | Homo sapiens | Feb 06 2026 | PRJNA1258992 | oocyte |
| GSE278658 | Sus scrofa | May 28 2025 | PRJNA1168156 | e iPSC-derived retinal organoid |
| GSE286385 | Homo sapiens;Mus musculus | Jan 01 2026 | PRJNA1209043 | |
| GSE254728 | Mus musculus | Feb 15 2026 | PRJNA1071580 | |

| | | | | |
|---|---|---|---|---|
| GSE149798 | Drosophila melanogaster | Feb 12 2021 | PRJNA630174 | muscle |
| GSE134743 | Homo sapiens | Jul 23 2020 | PRJNA556256 | |
| GSE223628 | Mus musculus | Jan 31 2023 | PRJNA926996 | |
| GSE277283 | Mus musculus | Jan 13 2026 | PRJNA1161648 | FL-BMDC |
| GSE296129 | Mus musculus | Jan 02 2026 | PRJNA1257847 | fetal testis |
| GSE295961 | Homo sapiens | Jan 01 2026 | PRJNA1256863 | colonic organoid |
| GSE223337 | Homo sapiens | Jan 10 2026 | PRJNA925833 | |
| GSE301062 | Homo sapiens | Jan 08 2026 | PRJNA1283198 | |
| GSE142622 | Arabidopsis thaliana | Jun 21 2020 | PRJNA597766 | Seedling |
| GSE159256 | Mus musculus | Oct 08 2023 | PRJNA668146 | Retina |
| GSE306854 | Mus musculus | Jan 08 2026 | PRJNA1312530 | Brain |
| GSE294900 | Mus musculus | Jan 06 2026 | PRJNA1252011 | brain-hippocampus |
| GSE305503 | Homo sapiens | Mar 18 2026 | PRJNA1306117 | iPSC Retinal Organoid |
| GSE163564 | Rattus norvegicus | Jul 28 2021 | PRJNA686775 | retina |
| GSE308548 | Homo sapiens | Feb 19 2026 | PRJNA1330863 | |
| GSE296354 | Mus musculus | Feb 10 2026 | PRJNA1258973 | |
| GSE279473 | Homo sapiens | Feb 10 2026 | PRJNA1173091 | |
| GSE231328 | Homo sapiens | Jun 10 2024 | PRJNA963774 | |
| GSE227717 | Mus musculus | Jan 21 2026 | PRJNA946710 | |
| GSE284136 | Gallus gallus;Petromyzon marinu | Aug 18 2025 | PRJNA1197421 | retina |
| GSE300142 | Homo sapiens | Jan 05 2026 | PRJNA1279121 | tumour |
| GSE316062 | Homo sapiens | Jan 16 2026 | PRJNA1400544 | lenocarcinoma, Esophagus (Lower 1/3) |
| GSE296577 | Homo sapiens | Feb 12 2026 | PRJNA1260003 | Pre-treatment biopsy |
| GSE313200 | Homo sapiens | Feb 18 2026 | PRJNA1378785 | |
| GSE313237 | Homo sapiens | Feb 18 2026 | | |
| GSE301885 | Homo sapiens | Jan 24 2026 | PRJNA1287690 | uman embryonic stem cell |
| GSE296540 | Drosophila melanogaster | Jan 12 2026 | PRJNA1259938 | central brain dissection |
| GSE283937 | Mus musculus | Jan 08 2026 | PRJNA1196349 | Colon tumor |
| GSE316228 | Mus musculus | Feb 08 2026 | PRJNA1401825 | Murine lungs |
| GSE254669 | Mus musculus | Jul 05 2024 | PRJNA1071483 | retina |
| GSE284101 | Homo sapiens | Jan 12 2026 | PRJNA1197356 | |
| GSE300175 | Mus musculus | Sep 02 2025 | PRJNA1279178 | Retina |
| GSE316055 | Mus musculus | Jan 15 2026 | PRJNA1400537 | Hippocampus and cortex |
| GSE211038 | Mus musculus | Aug 18 2022 | PRJNA868726 | Retina |
| GSE274051 | Homo sapiens | Feb 18 2026 | PRJNA1144810 | EBV+ Burkitt Lymphoma |
| GSE203604 | Gallus gallus | May 19 2025 | PRJNA841578 | |
| GSE202746 | Homo sapiens | Jun 08 2023 | PRJNA837213 | neural retina |
| GSE188268 | Mus musculus | Nov 07 2021 | PRJNA778172 | bone marrow |
| GSE243314 | Homo sapiens | Dec 31 2024 | PRJNA1017811 | retina |
| GSE267230 | Mus musculus | Jan 12 2026 | PRJNA1110284 | Liver |
| GSE301037 | Homo sapiens | Feb 09 2026 | PRJNA1283117 | |
| GSE317117 | Homo sapiens | Feb 12 2026 | PRJNA1406038 | |
| GSE287386 | Homo sapiens | Jan 18 2026 | PRJNA1212542 | hepatocellular carcinoma |
| GSE215318 | Homo sapiens | Mar 31 2023 | PRJNA889755 | |
| GSE298293 | Mus musculus | Jan 21 2026 | PRJNA1268898 | thymus |
| GSE301755 | Homo sapiens;Rattus | Feb 08 2026 | PRJNA1287535 | |
| GSE186816 | Homo sapiens | Jun 22 2023 | PRJNA776175 | |
| GSE301762 | lbopictus;Culex pipiens pallens;C | Jan 19 2026 | PRJNA1287547 | midgut |
| GSE285716 | Homo sapiens | Jan 01 2026 | PRJNA1205700 | Kidney |
| GSE304262 | Homo sapiens | Feb 18 2026 | | kidney |
| GSE315587 | Mus musculus | Feb 06 2026 | PRJNA1398273 | cornea |
| GSE275359 | Mus musculus | Mar 31 2025 | PRJNA1150535 | Retina |
| GSE184275 | Mus musculus | Aug 22 2022 | PRJNA763875 | retina |
| GSE315098 | Homo sapiens | Jan 01 2026 | PRJNA1394978 | |
| GSE255561 | Homo sapiens | Feb 06 2026 | PRJNA1075440 | |
| GSE297435 | Homo sapiens | Jan 13 2026 | PRJNA1263845 | |
| GSE141799 | Homo sapiens | Apr 14 2021 | PRJNA594746 | |
| GSE292348 | Homo sapiens | Jan 31 2026 | PRJNA1238208 | |
| GSE304500 | Mus musculus | Feb 17 2026 | PRJNA1301391 | Brain |
| GSE289223 | Mus musculus | Mar 19 2025 | PRJNA1221932 | RETINA |
| GSE207271 | Mus musculus | Jul 01 2022 | PRJNA854612 | |
| GSE319230 | Mus musculus | Feb 24 2026 | PRJNA1422651 | Cerebellum,brainstem |
| GSE290710 | Homo sapiens | Feb 26 2026 | PRJNA1229609 | breast |

| | | | | |
|---|---|---|---|---|
| GSE314764 | Homo sapiens | Jan 09 2026 | PRJNA1392568 | |
| GSE318533 | Homo sapiens | Feb 05 2026 | PRJNA1419182 | Bronchial brushing |
| GSE211842 | Paracentrotus lividus | Sep 20 2022 | PRJNA872340 | |
| GSE231596 | Mus musculus | May 16 2025 | PRJNA966999 | Primary DRG culture |
| GSE319552 | Sus scrofa | Feb 16 2026 | PRJNA1423717 | Liver |
| GSE311988 | lycobacterium tuberculosis H37I | Feb 25 2026 | | mycobacterial culture |
| GSE282351 | Homo sapiens | Feb 18 2026 | PRJNA1188179 | |
| GSE315732 | Homo sapiens | Jan 30 2026 | PRJNA1398863 | |
| GSE208428 | Mus musculus | Jul 16 2025 | PRJNA859902 | retina |
| GSE277229 | Mus musculus | Jan 01 2026 | PRJNA1161401 | Forebrain |
| GSE318779 | Acinetobacter baumannii | Feb 12 2026 | PRJNA1420531 | |
| GSE296812 | sophila melanogaster;Homo sap | Feb 24 2026 | PRJNA1261086 | |
| GSE290045 | Homo sapiens | Jan 30 2026 | PRJNA1226088 | |
| GSE137318 | Mus musculus | Aug 19 2020 | PRJNA565157 | |
| GSE314031 | Homo sapiens | Feb 13 2026 | PRJNA1389628 | |
| GSE125524 | Pseudomonas aeruginosa PAO1 | Jan 05 2026 | PRJNA516666 | |
| GSE287104 | Mus musculus | Jan 15 2026 | PRJNA1211129 | bone marrow |
| GSE286318 | Homo sapiens | Feb 07 2026 | PRJNA1208665 | |
| GSE313807 | Mus musculus | Feb 18 2026 | PRJNA1381617 | otopic colorectal cancer tissue |
| GSE212522 | Homo sapiens | Aug 18 2024 | PRJNA875898 | retina |
| GSE294524 | Homo sapiens | Feb 18 2026 | PRJNA1250446 | brain organoid |
| GSE314753 | Mus musculus | Jan 22 2026 | PRJNA1392539 | incisor mesenchyme |
| GSE316818 | Homo sapiens | Jan 16 2026 | PRJNA1404284 | Hela |
| GSE311253 | Homo sapiens | Jan 19 2026 | PRJNA1368855 | nan colorectal cancer cell line |
| GSE163690 | Homo sapiens | Dec 31 2021 | PRJNA687184 | |
| GSE310473 | Homo sapiens | Jan 26 2026 | PRJNA1366400 | lung |
| GSE232726 | Mus musculus | Aug 21 2023 | PRJNA973645 | Retina |
| GSE313924 | Homo sapiens | Jan 30 2026 | PRJNA1387361 | |
| GSE311370 | Mus musculus | Dec 10 2025 | PRJNA1369362 | retina |
| GSE255195 | Homo sapiens | Feb 06 2026 | PRJNA1073924 | |
| GSE289182 | Citrullus lanatus | Feb 06 2026 | PRJNA1221859 | leaves |
| GSE307267 | Mus musculus | Feb 12 2026 | PRJNA1315038 | ovary |
| GSE261060 | Mus musculus | Jun 23 2024 | | retina |
| GSE308878 | Mus musculus | Feb 24 2026 | PRJNA1333311 | oviductal epithelium |
| GSE313082 | Homo sapiens | Feb 06 2026 | PRJNA1378226 | |
| GSE319313 | Mus musculus | Feb 18 2026 | PRJNA1422806 | reatic Ductal Adenocarcinoma |
| GSE144427 | Mus musculus | Mar 09 2020 | PRJNA603600 | Retina |
| GSE242781 | Homo sapiens | Aug 21 2024 | PRJNA1014549 | Pigmented epithelium; Retina |
| GSE237211 | Ovis aries | Oct 01 2023 | PRJNA994177 | Retina |
| GSE283571 | Mus musculus | Feb 18 2026 | PRJNA1194552 | renal cortex |
| GSE233770 | Homo sapiens | Jan 01 2026 | PRJNA977887 | Liver |
| GSE296097 | Homo sapiens | Jan 12 2026 | PRJNA1257790 | |
| GSE315626 | Homo sapiens | Jan 17 2026 | PRJNA1398350 | |
| GSE315767 | Homo sapiens | Jan 11 2026 | PRJNA1398929 | Colorectal cancer tissue |
| GSE209661 | Homo sapiens | Jul 28 2022 | PRJNA862070 | Peripheral blood |
| GSE288933 | Homo sapiens;Mus musculus | Jan 29 2026 | PRJNA1221005 | |
| GSE299258 | Homo sapiens | Jan 20 2026 | PRJNA1273172 | Skin |
| GSE280794 | Homo sapiens | Sep 30 2025 | PRJNA1180547 | |
| GSE318551 | Mus musculus | Feb 05 2026 | | gastrocnemius |
| GSE307639 | Mus musculus | Feb 05 2026 | PRJNA1322439 | Contra |
| GSE272501 | Mus musculus | Jan 08 2026 | PRJNA1136984 | |
| GSE314706 | Homo sapiens | Feb 25 2026 | PRJNA1392485 | Nasal tissue |
| GSE302626 | Homo sapiens | Jan 14 2026 | PRJNA1291601 | Plasma |
| GSE319252 | Mus musculus | Feb 11 2026 | PRJNA1422697 | Brain |
| GSE272536 | Homo sapiens | Jan 21 2026 | PRJNA1137310 | Breast |
| GSE233164 | Homo sapiens | May 23 2023 | PRJNA975158 | |
| GSE139904 | Mus musculus | Jan 27 2022 | PRJNA587503 | |
| GSE269620 | Homo sapiens | Jul 31 2025 | PRJNA1122781 | retinal organoid |
| GSE161414 | Homo sapiens | Mar 26 2023 | PRJNA678241 | |
| GSE303670 | Homo sapiens | Jan 16 2026 | PRJNA1297055 | kidney biopsy |
| GSE274392 | Xenopus laevis | Jan 01 2026 | PRJNA1146322 | animal cap |
| GSE184933 | Mus musculus | Dec 07 2021 | PRJNA767059 | Cerebral cortex |
| GSE310250 | Mus musculus | Jan 06 2026 | PRJNA1365632 | T cells |

| | | | | |
|---|---|---|---|---|
| GSE182164 | Capra hircus | Aug 19 2021 | PRJNA755202 | optic nerve |
| GSE311312 | Homo sapiens | Jan 06 2026 | PRJNA1368920 | astrocytes |
| GSE289846 | Homo sapiens | Feb 18 2026 | PRJNA1225182 | |
| GSE268991 | Homo sapiens | Feb 15 2026 | PRJNA1119639 | adjacent normal |
| GSE305695 | Mus musculus | Feb 05 2026 | PRJNA1307484 | |
| GSE296953 | Homo sapiens | Jan 05 2026 | PRJNA1261984 | breast cancer |
| GSE264257 | Mus musculus | Oct 16 2024 | PRJNA1101555 | retina |
| GSE299653 | Danio rerio | Feb 25 2026 | PRJNA1275838 | |
| GSE303753 | Arabidopsis thaliana | Feb 19 2026 | PRJNA1297610 | seedling |
| GSE310344 | Homo sapiens | Feb 24 2026 | PRJNA1365784 | Skeletal Muscle |
| GSE298478 | Homo sapiens | Jan 26 2026 | PRJNA1269533 | kidney tumor |
| GSE295433 | Homo sapiens | Feb 24 2026 | PRJNA1254673 | |
| GSE316282 | Mus musculus | Jan 31 2026 | | |
| GSE289682 | Homo sapiens | Feb 18 2026 | PRJNA1223935 | |
| GSE254830 | Homo sapiens | Feb 05 2024 | PRJNA1072034 | Retinal organoid |
| GSE312910 | Mus musculus | Jan 08 2026 | PRJNA1377593 | Thoracic spinal cord |
| GSE255851 | Mus musculus | Jun 05 2024 | PRJNA1076855 | retina |
| GSE269549 | Mus musculus | Jul 17 2024 | PRJNA1122502 | retina |
| GSE296797 | Mus musculus | Jan 01 2026 | PRJNA1261065 | |
| GSE230838 | Mus musculus | Jan 21 2026 | PRJNA962812 | Liver |
| GSE303006 | Mus musculus | Jan 18 2026 | | prefontal cortex(right) |
| GSE295351 | Mus musculus | Jan 26 2026 | PRJNA1254252 | pancreatic islet |
| GSE309688 | Homo sapiens | Jan 06 2026 | PRJNA1336745 | Blood |
| GSE261771 | Homo sapiens | Mar 12 2026 | PRJNA1088827 | |
| GSE307576 | Mus musculus | Jan 28 2026 | PRJNA1322313 | Lung |
| GSE290122 | Mus musculus | Jan 28 2026 | PRJNA1226234 | |
| GSE314156 | Mus musculus | Jan 05 2026 | PRJNA1390189 | Duodenum |
| GSE276152 | Homo sapiens | Jan 01 2026 | PRJNA1155189 | |
| GSE306275 | Homo sapiens | Feb 26 2026 | PRJNA1310402 | Kidney |
| GSE315655 | Homo sapiens | Jan 11 2026 | PRJNA1398617 | |
| GSE303775 | Mus musculus | Feb 18 2026 | PRJNA1297664 | Ventricular heart |
| GSE302569 | Homo sapiens | Feb 09 2026 | PRJNA1291199 | ileum |
| GSE311334 | Homo sapiens | Feb 11 2026 | PRJNA1368973 | prefrontal cortex BA9 |
| GSE292585 | Homo sapiens | Jan 16 2026 | PRJNA1240259 | |
| GSE315428 | Homo sapiens | Feb 11 2026 | PRJNA1397143 | |
| GSE303986 | Ictidomys tridecemlineatus | Aug 03 2025 | PRJNA1298588 | retina |
| GSE116426 | Mus musculus | Jan 24 2020 | PRJNA478452 | retina |
| GSE149100 | Homo sapiens | Apr 23 2020 | PRJNA627387 | |
| GSE173150 | Homo sapiens | Feb 07 2022 | PRJNA723889 | |
| GSE245137 | Mus musculus | Mar 03 2025 | PRJNA1026993 | Retina |
| GSE297914 | Homo sapiens | Jan 29 2026 | PRJNA1266723 | |
| GSE285391 | Homo sapiens | Feb 11 2026 | PRJNA1203268 | cell line |
| GSE271953 | Homo sapiens | Feb 25 2026 | PRJNA1134195 | |
| GSE279589 | Homo sapiens | Feb 27 2026 | PRJNA1173556 | bone marrow |
| GSE302723 | Mus musculus | Jan 27 2026 | PRJNA1291791 | Liver |
| GSE222940 | Mus musculus | Jan 15 2026 | PRJNA924267 | lung |
| GSE194176 | Mus musculus | Jan 24 2022 | PRJNA799358 | retina |
| GSE270932 | Mus musculus | Feb 26 2026 | PRJNA1129062 | lungs + nasal tissue |
| GSE317929 | Mus musculus | Jan 30 2026 | PRJNA1415835 | hippocampus microglia |
| GSE209764 | Mus musculus | Nov 08 2022 | PRJNA862370 | 13.5 eye peripheral retina |
| GSE256242 | Homo sapiens | Jan 12 2026 | PRJNA1078767 | cornea endothelium |
| GSE315662 | Homo sapiens | Feb 09 2026 | PRJNA1398639 | cell line |
| GSE304187 | Mus musculus | Jan 04 2026 | PRJNA1299327 | lung |
| GSE328966 | Mus musculus | Apr 28 2026 | PRJNA1457098 | Retina |
| GSE303742 | Homo sapiens | Jan 29 2026 | PRJNA1297583 | |
| GSE287736 | Homo sapiens;Mus musculus | Feb 19 2026 | PRJNA1214364 | liver |
| GSE221899 | Homo sapiens | Jan 01 2026 | PRJNA916680 | |
| GSE141601 | Mus musculus | Nov 04 2020 | PRJNA594036 | ırified cone photoreceptors |
| GSE264118 | Homo sapiens | Feb 23 2026 | PRJNA1101015 | |
| GSE306403 | Homo sapiens | Aug 27 2025 | PRJNA1310585 | Neuroblastoma |
| GSE302600 | Mus musculus | Feb 28 2026 | PRJNA1291471 | Liver |
| GSE319274 | Cephalotaxus harringtonia | Feb 25 2026 | | young inner needles |
| GSE294048 | Mus musculus | Feb 18 2026 | PRJNA1247960 | Middle phalanx |

| | | | | |
|---|---|---|---|---|
| GSE271683 | Homo sapiens | Jan 07 2026 | PRJNA1132920 | |
| GSE160306 | Homo sapiens | Apr 19 2021 | PRJNA672928 | Retina Macula |
| GSE299761 | Mus musculus | Feb 18 2026 | PRJNA1276454 | |
| GSE314042 | Mus musculus | Dec 16 2025 | | retina |
| GSE290001 | Homo sapiens | Feb 20 2026 | PRJNA1225714 | |
| GSE206970 | Mus musculus | Jan 03 2024 | PRJNA863295 | retina |
| GSE266906 | Oryza sativa | Jan 26 2026 | PRJNA1108705 | 14-day-old seedlings |
| GSE270826 | Mus musculus | Feb 11 2026 | PRJNA1128574 | |
| GSE315985 | Mus musculus | Jan 12 2026 | PRJNA1400055 | E13.5 fetal liver |
| GSE280785 | Mus musculus | Apr 22 2025 | PRJNA1180535 | retina |
| GSE243843 | Mus musculus | Feb 05 2024 | PRJNA1020144 | Liver |
| GSE299175 | Mus musculus | Feb 15 2026 | PRJNA1273026 | Ovary |
| GSE312790 | Mus musculus | Jan 07 2026 | PRJNA1376138 | Substantia Nigra |
| GSE318401 | Mus musculus | Feb 07 2026 | PRJNA1418687 | cell |
| GSE264571 | Homo sapiens;Mus musculus | Dec 01 2024 | PRJNA1103168 | retina |
| GSE146900 | Mus musculus | Mar 13 2020 | PRJNA612322 | retina |
| GSE140157 | Mus musculus | Mar 27 2021 | PRJNA588437 | Aorta |
| GSE189491 | Musca domestica | Nov 30 2021 | PRJNA783282 | whole body |
| GSE225641 | Homo sapiens | Mar 11 2024 | PRJNA937022 | etinal pigment epithelium |
| GSE275978 | Homo sapiens | Aug 30 2025 | PRJNA1154007 | skin |
| GSE319460 | Mus musculus | Feb 17 2026 | PRJNA1423320 | Intestine |
| GSE145148 | Homo sapiens | Oct 06 2020 | PRJNA606196 | |
| GSE169527 | Mus musculus | Aug 11 2021 | PRJNA716982 | retina at PN24 |
| GSE290140 | Mus musculus | Feb 18 2026 | PRJNA1226257 | ouse embryonic fibroblasts |
| GSE128142 | Homo sapiens | Apr 22 2020 | PRJNA526535 | nal Pigmented Epithelial Cells |
| GSE284392 | Homo sapiens | Feb 12 2026 | PRJNA1198931 | |
| GSE302023 | Mus musculus | Jan 28 2026 | PRJNA1288373 | Testis |
| GSE164044 | Mus musculus | Dec 31 2020 | PRJNA688704 | |
| GSE173378 | Mus musculus | Jul 09 2021 | PRJNA725271 | retina |
| GSE200051 | Mus musculus | Apr 03 2025 | PRJNA822802 | Retina |
| GSE306220 | Homo sapiens | Jan 30 2026 | PRJNA1309702 | embryonic stem cell |
| GSE318354 | Homo sapiens | Feb 06 2026 | | |
| GSE319148 | Homo sapiens | Feb 11 2026 | PRJNA1422205 | PDAC |
| GSE124168 | Xenopus laevis | Feb 10 2020 | PRJNA510932 | Retinal ganglion cell axon |
| GSE159822 | Danio rerio | Oct 22 2020 | PRJNA670550 | optic fissure retina |
| GSE291409 | Homo sapiens | Feb 25 2026 | PRJNA1233318 | choalveolar Lavage fluid cells |
| GSE244661 | Mus musculus | Nov 29 2023 | PRJNA1023978 | eye |
| GSE224617 | Oncorhynchus mykiss | Feb 07 2023 | PRJNA931955 | |
| GSE337180 | Mus musculus | Jul 02 2026 | PRJNA1484424 | Retina |
| GSE304687 | Mus musculus | Jan 26 2026 | PRJNA1302328 | Tumor |
| GSE164167 | Homo sapiens | Jun 16 2021 | PRJNA689320 | retina |
| GSE296803 | Homo sapiens | Feb 19 2026 | PRJNA1261068 | |
| GSE318768 | Homo sapiens | Feb 11 2026 | PRJNA1419912 | tissue |
| GSE318277 | Mus musculus | Feb 05 2026 | PRJNA1418517 | |
| GSE287884 | Homo sapiens | Jan 17 2026 | PRJNA1214856 | |
| GSE319709 | Homo sapiens | Feb 20 2026 | | liver |
| GSE287122 | Mus musculus | Feb 11 2026 | PRJNA1211148 | Heart |
| GSE290701 | Homo sapiens | Jan 21 2026 | PRJNA1229594 | |
| GSE315569 | Homo sapiens | Jan 06 2026 | PRJNA1398291 | |
| GSE303634 | Mus musculus | Jan 28 2026 | PRJNA1295966 | aortic tissues |
| GSE306334 | Mus musculus | Jan 23 2026 | PRJNA1310484 | |
| GSE284799 | Mus musculus | Dec 04 2025 | PRJNA1200687 | Retina |
| GSE286143 | Homo sapiens | Jan 14 2025 | PRJNA1207995 | Eye |
| GSE288940 | Homo sapiens | Feb 06 2026 | PRJNA1220069 | |
| GSE316150 | Mus musculus | Jan 16 2026 | PRJNA1401737 | Spleen |
| GSE289550 | Mus musculus | Jan 01 2026 | PRJNA1223409 | nbar Dorsal Root, Spinal Cord |
| GSE292124 | Mus musculus | Feb 12 2026 | PRJNA1237235 | |
| GSE315089 | Mus musculus | Jan 26 2026 | PRJNA1394987 | Lung |
| GSE308024 | Mus musculus | Feb 28 2026 | PRJNA1328602 | trigeminal ganglion |
| GSE296327 | Mus musculus | Feb 18 2026 | | kidney |
| GSE253838 | Mus musculus | Feb 22 2024 | PRJNA1067544 | |
| GSE244979 | Homo sapiens | Feb 25 2026 | PRJNA1026556 | blood cell |
| GSE314979 | Mus musculus | Feb 11 2026 | PRJNA1393427 | ervical Lymph Nodes (cLNs) |

| | | | | |
|---|---|---|---|---|
| GSE291280 | Homo sapiens;Mus musculus | Jan 28 2026 | PRJNA1232708 | Left Ventricle |
| GSE80327 | Gallus gallus | Dec 31 2020 | PRJNA318660 | Retina, RPE, choroid |
| GSE180705 | Homo sapiens | Mar 05 2022 | PRJNA749262 | |
| GSE110623 | Mus musculus | Feb 24 2022 | PRJNA434120 | |
| GSE274636 | Homo sapiens | Jan 14 2026 | PRJNA1147317 | |
| GSE278149 | Homo sapiens | Jan 19 2026 | PRJNA1165762 | Nasal mucosa |
| GSE249905 | Mus musculus | Jan 14 2026 | PRJNA1051118 | |
| GSE318797 | Mus musculus | Feb 09 2026 | PRJNA1420647 | Tumor |
| GSE165384 | Mus musculus | Jan 23 2026 | PRJNA694300 | Hippocampus |
| GSE242406 | Mus musculus | May 28 2025 | PRJNA1041428 | Retina |
| GSE238218 | Mus musculus | Jun 06 2024 | PRJNA998379 | |
| GSE202757 | Mus musculus | Jun 08 2023 | PRJNA837221 | |
| GSE304039 | Homo sapiens | Jun 11 2026 | PRJNA1298810 | |
| GSE287688 | Homo sapiens;Mus musculus | Feb 19 2026 | PRJNA1214249 | |
| GSE318288 | nterica subsp. enterica serovar T | Feb 05 2026 | PRJNA1418535 | |
| GSE275102 | Caenorhabditis elegans | Jan 01 2026 | PRJNA1149305 | whole worms |
| GSE153812 | Mus musculus | Jul 11 2020 | PRJNA644185 | Retina |
| GSE312364 | Mus musculus | Feb 18 2026 | PRJNA1373169 | or lateral motor cortex nucleus |
| GSE308982 | Sus scrofa | Apr 07 2026 | PRJNA1333530 | Retina |
| GSE310930 | Homo sapiens | Feb 24 2026 | PRJNA1367565 | |
| GSE231732 | Mus musculus | Feb 26 2026 | PRJNA967384 | |
| GSE179016 | Homo sapiens | Mar 01 2022 | PRJNA741923 | |
| GSE317025 | Homo sapiens | Jan 27 2026 | PRJNA1405676 | hindbrain |
| GSE227053 | Mus musculus | May 19 2023 | PRJNA942962 | retina |
| GSE265956 | Homo sapiens | Apr 03 2025 | PRJNA1105043 | |
| GSE316726 | Mus musculus | Jan 20 2026 | | Pituitary |
| GSE304367 | Mus musculus | Feb 18 2026 | PRJNA1301120 | |
| GSE310182 | Mus musculus | Nov 17 2025 | PRJNA1365243 | Retina |
| GSE281410 | Mus musculus | Mar 19 2025 | PRJNA1183818 | Eye |
| GSE311681 | Mus musculus | Feb 17 2026 | PRJNA1370110 | liver |
| GSE316044 | Bos taurus | Jan 14 2026 | PRJNA1400522 | |
| GSE289444 | Mus musculus | Jan 28 2026 | PRJNA1222928 | Liver |
| GSE144847 | Mus musculus | Mar 10 2021 | PRJNA605142 | |
| GSE303595 | Saccharomyces cerevisiae | Jan 21 2026 | PRJNA1295895 | Whole cells |
| GSE301650 | Mus musculus | Jan 01 2026 | PRJNA1285919 | Liver |
| GSE256095 | Mus musculus | Feb 18 2026 | PRJNA1078244 | |
| GSE192505 | Mus musculus | Aug 16 2022 | PRJNA791676 | Retina |
| GSE317012 | Homo sapiens | Feb 10 2026 | PRJNA1405660 | Kidney |
| GSE319950 | Mus musculus | Feb 23 2026 | PRJNA1425585 | |
| GSE169499 | Escherichia coli K-12 | Mar 25 2021 | PRJNA716815 | |
| GSE292624 | Homo sapiens | Feb 23 2026 | PRJNA1240324 | ated unguided cerebral organoid |
| GSE296138 | Mus musculus | Feb 13 2026 | PRJNA1257857 | kidney |
| GSE315789 | Homo sapiens | Jan 12 2026 | PRJNA1399175 | |
| GSE280018 | Mus musculus | Jan 01 2026 | PRJNA1175742 | brain |
| GSE296345 | Homo sapiens | Jan 27 2026 | PRJNA1258956 | umbilical vein |
| GSE157839 | Homo sapiens | Oct 15 2021 | PRJNA662948 | |
| GSE313432 | Mus musculus | Jan 15 2026 | PRJNA1379497 | Bone marrow |
| GSE317319 | Homo sapiens | Jan 29 2026 | PRJNA1406739 | hindbrain |
| GSE247319 | Mus musculus | Dec 22 2023 | PRJNA1037107 | |
| GSE319794 | Homo sapiens | Feb 17 2026 | PRJNA1425103 | |
| GSE189007 | Homo sapiens | Nov 17 2021 | PRJNA781140 | Whole blood |
| GSE202735 | Homo sapiens | Jun 08 2023 | PRJNA837192 | neural retina |
| GSE302482 | Mus musculus | Feb 09 2026 | PRJNA1291066 | Bone marrow proB cells |
| GSE263040 | Homo sapiens | Jan 06 2026 | PRJNA1095505 | |
| GSE306681 | Mus musculus | Jan 13 2026 | PRJNA1311622 | sdLNs |
| GSE274663 | Homo sapiens | Dec 27 2024 | PRJNA1147602 | primary tumor cells |
| GSE270221 | Drosophila melanogaster | Feb 06 2026 | PRJNA1125427 | Whole third-instar larvae |
| GSE295697 | Mus musculus | Feb 12 2026 | PRJNA1255616 | heart |
| GSE294805 | Mus musculus | Feb 18 2026 | PRJNA1251628 | Heart, atrium |
| GSE290688 | Danio rerio | Feb 27 2026 | PRJNA1229547 | Whole larvae |
| GSE146759 | Homo sapiens | Dec 01 2020 | PRJNA611825 | |
| GSE247725 | Gallus gallus | Feb 24 2025 | PRJNA1040220 | |
| GSE297230 | Mus musculus | Jan 12 2026 | PRJNA1263210 | Lung |

| | | | | |
|---|---|---|---|---|
| GSE312058 | Staphylococcus aureus | Feb 23 2026 | PRJNA1372123 | |
| GSE314481 | Passer montanus | Jan 31 2026 | PRJNA1391391 | caudal hippocampus |
| GSE305364 | Homo sapiens | Jan 26 2026 | PRJNA1305403 | Skin |
| GSE263163 | Homo sapiens | Feb 15 2026 | PRJNA1095996 | Cell line |
| GSE255364 | Nothobranchius furzeri | Aug 21 2024 | PRJNA1074745 | Neural Retina |
| GSE278517 | Mus musculus | Jan 31 2026 | PRJNA1167682 | Heart |
| GSE316744 | Mus musculus;Mus pahari | Jan 27 2026 | PRJNA1404109 | xtraembryonic mesoderm |
| GSE192550 | Octopus vulgaris | Nov 11 2022 | PRJNA791920 | Axial Nerve Cord |
| GSE307930 | Mus musculus | Sep 19 2025 | PRJNA1327831 | etinal pigment epithelium |
| GSE293033 | Mus musculus | Jan 14 2026 | PRJNA1242382 | cortex |
| GSE307907 | Mus musculus | Feb 16 2026 | PRJNA1327774 | Spleen |
| GSE288470 | Homo sapiens | Jan 30 2026 | PRJNA1217820 | plasma |
| GSE314838 | Mus musculus | Jan 01 2026 | PRJNA1392904 | aorta |
| GSE270574 | Schizosaccharomyces pombe | Jan 27 2026 | PRJNA1127512 | |
| GSE222988 | Homo sapiens | Jan 01 2026 | PRJNA924345 | exocrine pancreas |
| GSE304332 | Mus musculus | Feb 25 2026 | PRJNA1300402 | SC and CB |
| GSE299879 | Mus musculus | Jan 21 2026 | PRJNA1277214 | retina |
| GSE313511 | Homo sapiens | Jan 15 2026 | PRJNA1379693 | |
| GSE316011 | Mus musculus | Jan 31 2026 | PRJNA1400445 | cortex |
| GSE269305 | Mus musculus | Jan 06 2026 | PRJNA1120934 | |
| GSE309955 | Mus musculus | Feb 12 2026 | PRJNA1364040 | Adipose |
| GSE311076 | Mus musculus | Jan 28 2026 | PRJNA1368629 | Lung |
| GSE295002 | Homo sapiens | Feb 11 2026 | PRJNA1252421 | |
| GSE291201 | Homo sapiens | Jan 12 2026 | PRJNA1232191 | |
| GSE315275 | Homo sapiens | Jan 10 2026 | PRJNA1395553 | adipose tissue |
| GSE218818 | Mus musculus | Nov 30 2022 | PRJNA905686 | eye |
| GSE319740 | Escherichia coli | Feb 20 2026 | PRJNA1425034 | |
| GSE315999 | Polypterus senegalus | Jan 15 2026 | PRJNA1400083 | pectoral fin |
| GSE310017 | Mus musculus | Feb 12 2026 | PRJNA1365013 | liver |
| GSE292834 | Mus musculus | Jan 21 2026 | PRJNA1241693 | retinal vein |
| GSE319268 | human feces metagenome | Feb 16 2026 | PRJNA1422728 | Stool |
| GSE300957 | Danio rerio | Feb 06 2026 | PRJNA1282735 | whole animal |
| GSE283368 | Homo sapiens | Jan 28 2026 | PRJNA1193533 | Vastus lateralis |
| GSE302303 | Mus musculus | Jan 05 2026 | | cerebral cortex |
| GSE311723 | Caenorhabditis elegans | Jan 08 2026 | PRJNA1370839 | whole organism |
| GSE221037 | Homo sapiens | Jun 21 2023 | PRJNA912646 | |
| GSE276545 | Homo sapiens | Feb 18 2026 | PRJNA1157821 | cell line |
| GSE317187 | Homo sapiens | Jan 23 2026 | PRJNA1406234 | |
| GSE319015 | Mus musculus | Feb 11 2026 | PRJNA1421946 | Brain cortex |
| GSE302461 | Arabidopsis thaliana | Jan 26 2026 | PRJNA1291039 | seedling |
| GSE288098 | Mus musculus | Mar 01 2025 | PRJNA1216193 | Whole Retina |
| GSE317498 | Homo sapiens | Jan 28 2026 | PRJNA1414266 | |
| GSE317816 | Homo sapiens | Jan 28 2026 | PRJNA1415311 | Lung |
| GSE294177 | Rattus norvegicus | Feb 25 2026 | PRJNA1248518 | brain tissue |
| GSE316954 | Homo sapiens | Jan 27 2026 | PRJNA1405578 | heral blood mononuclear cells |
| GSE268270 | Homo sapiens | Jan 24 2026 | PRJNA1116059 | ICM |
| GSE318448 | Mus musculus | Feb 07 2026 | PRJNA1418745 | peritoneal lavage fluid |
| GSE319654 | Mus musculus | Feb 18 2026 | PRJNA1424922 | Tail tendon fascicles |
| GSE315361 | Mus musculus | Jan 26 2026 | PRJNA1396598 | |
| GSE317935 | Saccharomyces cerevisiae | Jan 28 2026 | | |
| GSE289390 | Homo sapiens | Jan 22 2026 | PRJNA1222835 | T cell |
| GSE290421 | Mus musculus | Feb 25 2026 | PRJNA1227980 | Lung |
| GSE310656 | Mus musculus | Feb 18 2026 | PRJNA1366830 | brain |
| GSE303276 | acillus toyonensis;Escherichia co | Feb 18 2026 | PRJNA1294728 | |
| GSE244337 | omo sapiens;unidentified plasm | Jun 24 2024 | PRJNA1022426 | plasmid |
| GSE310134 | Homo sapiens | Jan 01 2026 | PRJNA1365177 | Whole blood |
| GSE299414 | Homo sapiens | Jan 01 2026 | | embryonic kidneys |
| GSE292917 | Arabidopsis thaliana | Jan 04 2026 | PRJNA1241867 | leaf |
| GSE298608 | Mus musculus | Jan 22 2026 | PRJNA1270023 | incisor mesenchyme |
| GSE303514 | Homo sapiens | Feb 18 2026 | PRJNA1295552 | rimary human monocytes |
| GSE148300 | Homo sapiens | Jul 08 2020 | PRJNA623855 | |
| GSE157129 | Gallus gallus | Jul 02 2021 | PRJNA660186 | hal pigment epithelium (RPE) |
| GSE315217 | andida albicans;Candida tropical | Jan 28 2026 | PRJNA1395136 | cell |

| | | | | |
|---|---|---|---|---|
| GSE239941 | Mus musculus | Feb 28 2024 | PRJNA1001664 | retina and choroid |
| GSE310717 | Equus caballus | Feb 27 2026 | PRJNA1366981 | |
| GSE286440 | Homo sapiens | May 22 2025 | PRJNA1209364 | Human Umbilical Vein |
| GSE309937 | Arabidopsis thaliana | Jan 17 2026 | PRJNA1364015 | Roots |
| GSE315708 | Mus musculus | Feb 15 2026 | PRJNA1398827 | tumor s.c. |
| GSE179870 | Homo sapiens | Feb 09 2026 | PRJNA745442 | |
| GSE136421 | Homo sapiens | May 19 2020 | PRJNA562527 | |
| GSE199548 | Homo sapiens | Sep 20 2022 | PRJNA820706 | |
| GSE183320 | Homo sapiens | Feb 21 2022 | PRJNA759922 | CD31_choroid |
| GSE309900 | Homo sapiens | Feb 20 2026 | PRJNA1363969 | |
| GSE307103 | Homo sapiens | Jan 28 2026 | PRJNA1314270 | |
| GSE286168 | Mus musculus | Jan 01 2026 | PRJNA1208040 | Colon |
| GSE303300 | Homo sapiens | Jan 28 2026 | PRJNA1294793 | Tumor |
| GSE309091 | Homo sapiens | Jan 01 2026 | PRJNA1333941 | ıman osteosarcoma tissues |
| GSE290048 | Homo sapiens | Jan 20 2026 | PRJNA1226096 | |
| GSE274811 | Homo sapiens | Feb 19 2026 | PRJNA1148093 | Melanoma |
| GSE252361 | Mus musculus | Jan 09 2026 | PRJNA1060295 | |
| GSE291029 | Homo sapiens | Oct 05 2025 | PRJNA1231422 | Eye |
| GSE316184 | Histoplasma ohiense | Jan 26 2026 | PRJNA1401770 | |
| GSE245542 | Mus musculus | Jun 06 2024 | PRJNA1028971 | retina |
| GSE299562 | Homo sapiens | Feb 09 2026 | PRJNA1274985 | |
| GSE294368 | Drosophila melanogaster | Sep 04 2025 | PRJNA1249269 | Out photoreceptor |
| GSE132229 | Mus musculus | Jan 24 2020 | PRJNA546561 | retina |
| GSE319154 | Homo sapiens | Feb 15 2026 | PRJNA1422220 | |
| GSE278596 | Mus musculus | Jan 12 2026 | PRJNA1168059 | aorta |
| GSE300187 | Mus musculus | Feb 10 2026 | PRJNA1279188 | |
| GSE286236 | Homo sapiens | Jan 13 2025 | PRJNA1208464 | Control RGC soma |
| GSE319636 | Mus musculus | Feb 17 2026 | PRJNA1424912 | |
| GSE299441 | Mus musculus | Jan 09 2026 | PRJNA1274450 | Sciatic DRG neurons |
| GSE310161 | Homo sapiens | Jan 20 2026 | PRJNA1365213 | lung cancer |
| GSE294115 | Mus musculus | Feb 06 2026 | PRJNA1248408 | Aorta |
| GSE145601 | Mus musculus | Feb 21 2020 | PRJNA607723 | |
| GSE160011 | Mus musculus | Feb 16 2021 | PRJNA671602 | Retinal microglia |
| GSE288393 | Mus musculus | Feb 06 2026 | PRJNA1217350 | uodenum crypt epithelium |
| GSE300821 | Capra hircus | Feb 24 2026 | PRJNA1282151 | hair follicle stem cells from goat skin |
| GSE245456 | Mus musculus | Mar 14 2024 | PRJNA1028541 | Retina |
| GSE310449 | Fundulus heteroclitus | Apr 08 2026 | PRJNA1366356 | eye |
| GSE303069 | Homo sapiens | Jan 28 2026 | | brain |
| GSE319601 | Mus musculus | Feb 19 2026 | PRJNA1424708 | whole lung |
| GSE111782 | Homo sapiens | Mar 13 2021 | PRJNA438128 | |
| GSE260513 | Mus musculus | Feb 28 2026 | PRJNA1081911 | Striatum |
| GSE226784 | Homo sapiens | Jan 01 2026 | PRJNA942051 | |
| GSE317960 | Mus musculus | Feb 12 2026 | PRJNA1416225 | Liver |
| GSE252517 | Mus musculus | Jan 26 2024 | PRJNA1061061 | |
| GSE236762 | Mus musculus | Jan 09 2026 | PRJNA992391 | left ventricular tissue |
| GSE180158 | Homo sapiens;synthetic construc | Nov 12 2021 | PRJNA746784 | |
| GSE315827 | Mus musculus | Jan 12 2026 | PRJNA1399377 | gastruloid |
| GSE309618 | Mus musculus | Jan 01 2026 | PRJNA1336614 | Kidney |
| GSE317273 | Mus musculus | Jan 27 2026 | PRJNA1406653 | Thymus |
| GSE147013 | Homo sapiens | Jul 15 2022 | PRJNA612795 | Retina |
| GSE268936 | Homo sapiens | Jun 11 2025 | PRJNA1119479 | PBMC |
| GSE297766 | Mus musculus | Jan 07 2026 | PRJNA1266147 | epidermis |
| GSE313712 | Homo sapiens | Jan 22 2026 | PRJNA1381468 | |
| GSE275515 | Danio rerio | Jan 15 2026 | PRJNA1151600 | S sorted kdrl:mCherry+ cells |
| GSE292266 | Homo sapiens | Feb 12 2026 | PRJNA1237792 | cell line |
| GSE216834 | Homo sapiens | Oct 01 2025 | PRJNA895463 | |
| GSE285445 | Ovis aries | Jan 14 2026 | PRJNA1203644 | rumen |
| GSE290805 | Homo sapiens | Feb 11 2026 | PRJNA1230148 | Human meningioma |
| GSE178709 | Danio rerio | Jan 25 2023 | PRJNA740056 | dorsal retina |
| GSE162161 | Homo sapiens | Jun 20 2021 | PRJNA680778 | |
| GSE293552 | nerichia coli str. K-12 substr. W3 | Jan 06 2026 | PRJNA1245175 | |
| GSE325522 | ns;Macaca fascicularis;Mus mus | Mar 25 2026 | | Eye,Eye |
| GSE294688 | Mus musculus | Jan 27 2026 | PRJNA1251088 | Spleen |

| | | | | |
|---|---|---|---|---|
| GSE316225 | Mus musculus | Feb 16 2026 | PRJNA1401829 | Control Mouse skin |
| GSE305167 | Danio rerio | Feb 17 2026 | | Whole retinas |
| GSE278714 | Homo sapiens | Aug 14 2025 | PRJNA1168505 | |
| GSE42419 | Mus musculus | Feb 26 2026 | PRJNA181365 | heart |
| GSE316547 | Mus musculus | Jan 16 2026 | PRJNA1403298 | proximal small intestine |
| GSE316619 | Mus musculus | Jan 28 2026 | PRJNA1403481 | colon |
| GSE236689 | Homo sapiens | Jan 02 2024 | PRJNA992056 | |
| GSE312105 | Mus musculus | Jan 08 2026 | PRJNA1372181 | Colon tumor |
| GSE205491 | Mus musculus | Aug 04 2022 | PRJNA845993 | Retina |
| GSE278155 | Mus musculus | Feb 27 2026 | PRJNA1165768 | liver |
| GSE200191 | Mus musculus | Nov 11 2022 | PRJNA823648 | RPE |
| GSE249919 | Mus musculus | Dec 31 2024 | PRJNA1051126 | eyeball |
| GSE288278 | Mus musculus | Jan 21 2026 | PRJNA1216740 | m flash-frozen powderized whole pancreas |
| GSE315933 | Mus musculus | Jan 12 2026 | PRJNA1399980 | Liver |
| GSE281030 | Mus musculus | Jun 23 2025 | PRJNA1181862 | Retina |
| GSE275363 | Mus musculus | Jan 05 2026 | PRJNA1150544 | colon |
| GSE313153 | Homo sapiens | Feb 23 2026 | PRJNA1378721 | spleen |
| GSE315095 | Mus musculus | Jan 06 2026 | PRJNA1394991 | CD1 mouse fetal kidney |
| GSE279103 | Homo sapiens | Jan 12 2026 | PRJNA1170865 | |
| GSE223358 | Homo sapiens | Jan 20 2026 | PRJNA925862 | |
| GSE295580 | Caulobacter vibrioides | Feb 25 2026 | PRJNA1255210 | |
| GSE189551 | Mus musculus | Oct 16 2024 | PRJNA783539 | |
| GSE316331 | Mus musculus | Jan 31 2026 | PRJNA1402329 | |
| GSE269687 | Homo sapiens | Apr 02 2025 | PRJNA1123233 | DIV15 RPCs |
| GSE126438 | Homo sapiens | Feb 14 2026 | PRJNA521888 | lymphoblastoid cell line |
| GSE268184 | Homo sapiens | Jan 24 2026 | PRJNA1115061 | human 4C stage embryo |
| GSE312073 | Mus musculus | Jan 13 2026 | PRJNA1372146 | |
| GSE153978 | Homo sapiens | Jul 08 2020 | PRJNA644639 | |
| GSE263148 | Mus musculus | Feb 11 2026 | PRJNA1095978 | Ear skin |
| GSE296518 | Mus musculus | Feb 23 2026 | PRJNA1259807 | |
| GSE211702 | Mus musculus | Jan 06 2023 | PRJNA871778 | Retina |
| GSE307595 | Homo sapiens | Feb 26 2026 | PRJNA1322361 | lung |
| GSE222005 | Mus musculus | Oct 30 2023 | PRJNA917303 | retina |
| GSE274998 | Homo sapiens | Jan 15 2026 | PRJNA1148929 | |
| GSE296765 | Mus musculus | Jan 20 2026 | PRJNA1261023 | Liver |
| GSE284159 | Homo sapiens | Jan 12 2026 | PRJNA1197473 | |
| GSE316512 | Arabidopsis thaliana | Jan 20 2026 | PRJNA1403040 | otyls and the upper 5mm of root |
| GSE289112 | Scylla paramamosain | Jan 12 2026 | PRJNA1221561 | hemolymph |
| GSE305105 | Mus musculus | Feb 22 2026 | PRJNA1304440 | abdominal aorta |
| GSE288339 | Mus musculus | Feb 16 2026 | PRJNA1216961 | |
| GSE312726 | Homo sapiens | Jan 31 2026 | PRJNA1375784 | |
| GSE291197 | Homo sapiens | Jan 12 2026 | PRJNA1232184 | |
| GSE317226 | Homo sapiens | Feb 05 2026 | PRJNA1406305 | Kidney |
| GSE297733 | Homo sapiens | Jan 19 2026 | PRJNA1266069 | Oligodendroglioma |
| GSE300367 | Mus musculus | Feb 09 2026 | PRJNA1280158 | Heart |
| GSE274243 | Mus musculus | Feb 28 2026 | PRJNA1145448 | embryo E14.5 |
| GSE304164 | Mus musculus | Feb 25 2026 | PRJNA1299325 | |
| GSE316758 | Homo sapiens | Jan 21 2026 | PRJNA1404151 | hepatic |
| GSE312605 | Mus musculus | Jan 22 2026 | PRJNA1375517 | incisor mesenchyme |
| GSE286388 | Homo sapiens | Jan 07 2026 | PRJNA1209045 | |
| GSE317178 | Mus musculus | Jan 26 2026 | PRJNA1406224 | brain |
| GSE307481 | Homo sapiens | Jan 21 2026 | PRJNA1321817 | brain |
| GSE301058 | Escherichia coli MP1 | Jan 12 2026 | PRJNA1283177 | |
| GSE181550 | Homo sapiens | Nov 01 2021 | PRJNA752566 | retina |
| GSE288067 | Homo sapiens | Jan 30 2026 | PRJNA1215986 | pancreas |
| GSE309513 | Mus musculus | Jan 28 2026 | PRJNA1336198 | |
| GSE192458 | Mus musculus | Apr 20 2022 | PRJNA791604 | retina |
| GSE288050 | Homo sapiens | May 22 2025 | PRJNA1215938 | |
| GSE301570 | Homo sapiens | Jan 11 2026 | PRJNA1285463 | |
| GSE280004 | Homo sapiens | Feb 25 2026 | PRJNA1175725 | blood |
| GSE304070 | Mus musculus | Jan 19 2026 | PRJNA1298867 | Brain |
| GSE262277 | Homo sapiens | Oct 10 2024 | PRJNA1090949 | Retinal Pigment Epithelial |
| GSE290973 | Homo sapiens | Jan 07 2026 | PRJNA1231107 | steoarthritic knee cartilage |

| | | | | |
|---|---|---|---|---|
| GSE316353 | Mus musculus | Jan 20 2026 | PRJNA1402361 | eye |
| GSE292327 | Homo sapiens | Jan 12 2026 | PRJNA1237935 | meningioma |
| GSE311596 | Mus musculus | Jan 26 2026 | PRJNA1369979 | |
| GSE297627 | Rattus norvegicus | Jan 11 2026 | PRJNA1265495 | mammary tumor |
| GSE317332 | Homo sapiens | Jan 23 2026 | PRJNA1406761 | small intestine organoid |
| GSE324435 | Danio rerio | Mar 11 2026 | PRJNA1435103 | zebrafish whole brain |
| GSE287832 | Mus musculus | Jan 01 2026 | PRJNA1214791 | Skeletal Muscle |
| GSE308545 | Brassica rapa | Feb 18 2026 | PRJNA1330859 | leaf |
| GSE175895 | Mus musculus | Jul 16 2021 | PRJNA734268 | Retina |
| GSE304740 | Homo sapiens | Feb 27 2026 | PRJNA1302433 | Blood Cells |
| GSE292417 | Homo sapiens | Jan 14 2026 | PRJNA1238346 | cell line |
| GSE292899 | Homo sapiens | Feb 11 2026 | PRJNA1241830 | |
| GSE281037 | Mus musculus | Jan 02 2026 | PRJNA1181871 | Testis |
| GSE267725 | Homo sapiens | Feb 13 2026 | PRJNA1112475 | MCF 7.00 |
| GSE312102 | Mus musculus | Jan 25 2026 | PRJNA1372174 | spleen |
| GSE263039 | Homo sapiens | Jan 06 2026 | PRJNA1095503 | |
| GSE179431 | Mus musculus | Jul 07 2021 | PRJNA743679 | Lung |
| GSE289701 | Mus musculus | Aug 19 2025 | PRJNA1224216 | Retina |
| GSE316301 | Homo sapiens | Feb 06 2026 | PRJNA1402258 | PBMC |
| GSE312893 | Mus musculus | Jan 30 2026 | PRJNA1377569 | |
| GSE290087 | Macaca fascicularis | Jan 19 2026 | PRJNA1226180 | Ovary |
| GSE317435 | Homo sapiens | Jan 24 2026 | PRJNA1412757 | Breast, Mammary Gland |
| GSE288035 | Mus musculus | Jan 26 2025 | PRJNA1215752 | retina+choroid |
| GSE285561 | Homo sapiens | Jan 17 2025 | PRJNA1208989 | |
| GSE302842 | Mus musculus | Jan 27 2026 | PRJNA1292261 | Liver |
| GSE290916 | Staphylococcus aureus | Feb 25 2026 | PRJNA1230787 | |
| GSE279666 | Mus musculus | Feb 20 2026 | PRJNA1174054 | brain |
| GSE268183 | Homo sapiens | Jan 24 2026 | PRJNA1115062 | human 8C embryo |
| GSE304015 | Mus musculus | Feb 25 2026 | PRJNA1298762 | Subcutanous tumor |
| GSE235993 | Mus musculus | Jun 27 2024 | PRJNA988140 | Retina |
| GSE297120 | Homo sapiens | Feb 22 2026 | PRJNA1262826 | |
| GSE312348 | Mus musculus | Jan 20 2026 | PRJNA1373138 | |
| GSE287857 | Mus musculus | Jan 23 2026 | PRJNA1214817 | |
| GSE215325 | Homo sapiens | Mar 31 2023 | PRJNA889762 | |
| GSE309223 | Mus musculus | Sep 30 2025 | PRJNA1334369 | retina |
| GSE329104 | Homo sapiens | May 01 2026 | PRJNA1457992 | RPE |
| GSE288409 | Gallus gallus | Jan 31 2026 | PRJNA1217400 | |
| GSE184121 | Drosophila melanogaster | Dec 31 2021 | PRJNA763225 | |
| GSE296062 | Homo sapiens | Jan 30 2026 | PRJNA1257424 | |
| GSE164906 | Mus musculus | Jul 21 2022 | PRJNA692426 | |
| GSE278724 | Mus musculus | Jan 27 2026 | PRJNA1168522 | |
| GSE298692 | Homo sapiens | Feb 15 2026 | PRJNA1270960 | |
| GSE289177 | Cryptocercus punctulatus | Feb 10 2026 | PRJNA1221862 | all |
| GSE307292 | Homo sapiens | Feb 25 2026 | PRJNA1315155 | Tumor |
| GSE312774 | Danio rerio | Jan 05 2026 | PRJNA1375859 | Caudal Fin |
| GSE206626 | Mus musculus | Aug 20 2022 | PRJNA851588 | retina |
| GSE304654 | Mus musculus | Jan 15 2026 | PRJNA1302208 | Aorta |
| GSE317274 | Homo sapiens | Jan 27 2026 | PRJNA1406652 | |
| GSE280617 | Homo sapiens | Feb 11 2026 | PRJNA1179500 | Cell Line |
| GSE262464 | Mus musculus | Jan 27 2026 | PRJNA1091931 | spleen |
| GSE267164 | Mus musculus | Feb 23 2026 | PRJNA1109988 | Blood |
| GSE293728 | Mus musculus | Feb 22 2026 | PRJNA1246119 | |
| GSE289140 | Danio rerio | Feb 15 2025 | PRJNA1221797 | |
| GSE292534 | Mus musculus | Feb 12 2026 | PRJNA1239921 | |
| GSE223529 | Mus musculus | Feb 21 2024 | PRJNA926649 | |
| GSE222735 | Mus musculus | Jan 01 2026 | | |
| GSE297249 | Mus musculus | Feb 18 2026 | PRJNA1263389 | Bone Marrow |
| GSE196235 | Homo sapiens | Mar 09 2022 | PRJNA803994 | Retina |
| GSE304101 | Homo sapiens | Feb 18 2026 | PRJNA1298912 | |
| GSE260952 | Mus musculus | Aug 28 2024 | PRJNA1084724 | Retina |
| GSE300243 | Danio rerio | Feb 06 2026 | PRJNA1279962 | |
| GSE211611 | Homo sapiens | Aug 22 2022 | PRJNA871132 | |
| GSE231536 | Homo sapiens | Sep 11 2023 | PRJNA966249 | |

| | | | | |
|---|---|---|---|---|
| GSE284728 | Danio rerio | Jun 02 2025 | PRJNA1200121 | retina |
| GSE238261 | Mus musculus | Jan 06 2026 | PRJNA998473 | Ovary |
| GSE299785 | Bos taurus | Feb 06 2026 | PRJNA1276517 | |
| GSE293414 | Mus musculus | Jan 01 2026 | PRJNA1244387 | |
| GSE304040 | Mus musculus | Jun 11 2026 | PRJNA1298816 | |
| GSE316401 | Mus musculus | Jan 14 2026 | PRJNA1402753 | tumor |
| GSE289001 | Mus musculus | Jan 05 2026 | PRJNA1220584 | Kidney |
| GSE275306 | Mus musculus | Feb 07 2026 | PRJNA1150229 | Pineal tumor |
| GSE293376 | Homo sapiens;Mus musculus | Jun 30 2026 | PRJNA1244296 | |
| GSE282493 | Homo sapiens | Jan 01 2026 | PRJNA1189017 | |
| GSE212582 | Homo sapiens | Apr 20 2023 | PRJNA876116 | Retinal organoid |
| GSE304715 | Oncorhynchus mykiss | Jan 08 2026 | PRJNA1302393 | (freshly fertilized egg/zygote) |
| GSE313429 | Mus musculus | Jan 15 2026 | PRJNA1379498 | Bone marrow |
| GSE277289 | Mus musculus | Jan 13 2026 | PRJNA1161653 | |
| GSE298920 | Plasmodium falciparum | Jan 01 2026 | PRJNA1271790 | |
| GSE289754 | Mus | Feb 26 2026 | PRJNA1224451 | LIVER |
| GSE147328 | Mus musculus | Mar 22 2020 | PRJNA613823 | Retina |
| GSE263573 | Homo sapiens | Feb 28 2026 | PRJNA1098083 | man pancreatic cancer cells |
| GSE318809 | Bacteroides fragilis | Feb 09 2026 | PRJNA1420658 | |
| GSE274511 | Mus musculus | Jan 21 2026 | PRJNA1146901 | |
| GSE193002 | Mus musculus | Dec 30 2023 | PRJNA794114 | RPE-choroid complex |
| GSE289884 | Mus musculus | Feb 18 2026 | PRJNA1225244 | Hippocampus |
| GSE315842 | Mus musculus | Jan 12 2026 | PRJNA1399414 | Lung |
| GSE287598 | Homo sapiens | Jan 01 2026 | | Trachea organoid |
| GSE312209 | Homo sapiens | Feb 12 2026 | PRJNA1372671 | |
| GSE316515 | Mus musculus | Jan 19 2026 | PRJNA1403043 | eral Septum Brain Dissection |
| GSE286977 | Gallus gallus | Jan 30 2026 | | cecal tissue |
| GSE301454 | Mus musculus | Feb 27 2026 | PRJNA1285261 | muscle |
| GSE299691 | Mus musculus | Jul 01 2025 | PRJNA1276071 | retina |
| GSE148814 | Mus musculus | Mar 17 2021 | PRJNA625890 | |
| GSE319868 | Homo sapiens | Feb 18 2026 | PRJNA1425440 | ell-lline-Derived Xenograft |
| GSE315399 | Homo sapiens | Jan 07 2026 | PRJNA1396669 | ovary |
| GSE281614 | Mus musculus | Feb 09 2026 | PRJNA1184800 | PDAC |
| GSE276810 | Mus musculus | Jan 22 2026 | PRJNA1159143 | lung |
| GSE315686 | Mus musculus | Jan 09 2026 | PRJNA1398795 | Spleen and Lymph node |
| GSE274244 | Mus musculus | Feb 28 2026 | PRJNA1145447 | onic stage (E16.5:telencephalon) |
| GSE288143 | Mus musculus | Jan 21 2026 | PRJNA1216275 | lymph node |
| GSE317543 | Rattus norvegicus | Jan 31 2026 | PRJNA1414437 | |
| GSE278771 | Homo sapiens | Feb 18 2026 | PRJNA1168872 | |
| GSE253955 | Mus musculus | Jan 28 2026 | PRJNA1067991 | spleen |
| GSE267419 | Mus musculus | Sep 02 2024 | PRJNA1111474 | retina |
| GSE212189 | Mus musculus | Nov 30 2022 | PRJNA874435 | bulk retina |
| GSE271539 | Homo sapiens | Jan 01 2026 | PRJNA1132184 | |
| GSE287232 | Mus musculus | Jan 21 2025 | PRJNA1211662 | organoids |
| GSE304483 | Mus musculus | Feb 09 2026 | PRJNA1301371 | hippocampus |
| GSE315366 | Homo sapiens | Jan 06 2026 | PRJNA1396607 | Peripheral blood |
| GSE307045 | Mus musculus | Jan 08 2026 | PRJNA1314159 | inner ear cells |
| GSE317932 | Mus musculus | Jan 29 2026 | PRJNA1415838 | lung origine |
| GSE285866 | Stylosanthes guianensis | Jan 04 2026 | PRJNA1206474 | Root |
| GSE186751 | Homo sapiens | Jan 01 2022 | | |
| GSE303759 | Homo sapiens | Feb 13 2026 | PRJNA1297634 | |
| GSE144605 | Homo sapiens | May 26 2020 | PRJNA604188 | |
| GSE285695 | Homo sapiens | Jan 28 2026 | PRJNA1205665 | |
| GSE310104 | Streptomyces venezuelae | Feb 11 2026 | PRJNA1365143 | |
| GSE212469 | Homo sapiens | Nov 04 2022 | PRJNA875478 | |
| GSE310895 | Tetraodon nigroviridis | Apr 08 2026 | PRJNA1367495 | Dorsal retina |
| GSE274957 | Danio rerio | Jan 01 2026 | PRJNA1148645 | pronephric duct |
| GSE290344 | Homo sapiens | Feb 11 2026 | PRJNA1227516 | Retinal organoid |
| GSE311152 | Mus musculus | Feb 23 2026 | PRJNA1368720 | or lateral motor cortex nucleus |
| GSE313297 | Danio rerio | Jan 10 2026 | PRJNA1378886 | ed head tissue from 30 hpf embryos |
| GSE319106 | Staphylococcus aureus | Feb 11 2026 | PRJNA1422106 | |
| GSE220627 | Homo sapiens;Mus musculus | Sep 15 2023 | PRJNA910547 | retinal organoid |
| GSE317094 | Mus musculus | Feb 16 2026 | PRJNA1405765 | |

| | | | | |
|---|---|---|---|---|
| GSE274793 | Mus musculus | Aug 14 2024 | PRJNA1148067 | Retina |
| GSE313788 | Mus musculus | Jan 31 2026 | PRJNA1381597 | Quaddel] primary adult APH treated spleen |
| GSE236218 | Homo sapiens;Mus musculus | Jan 01 2026 | PRJNA989270 | bone marrow |
| GSE313858 | Homo sapiens | Jan 05 2026 | | Tumor |
| GSE163591 | Homo sapiens | Dec 22 2020 | PRJNA686886 | Retina |
| GSE283358 | Homo sapiens | Feb 17 2026 | PRJNA1193518 | cell line |
| GSE309952 | Mus musculus | Feb 25 2026 | PRJNA1364036 | Spleen |
| GSE269302 | Mus musculus | Jan 06 2026 | PRJNA1120929 | |
| GSE200423 | Ambystoma mexicanum | Apr 10 2022 | PRJNA824487 | Retina |
| GSE205123 | Mus musculus | Jun 03 2022 | PRJNA843960 | Retina |
| GSE318726 | Mus musculus | Feb 11 2026 | PRJNA1419843 | Liver cancer |
| GSE312839 | Homo sapiens | Apr 27 2026 | PRJNA1377498 | |
| GSE264349 | Mus musculus | Oct 10 2024 | PRJNA1102013 | Retina |
| GSE304597 | Sus scrofa | Feb 18 2026 | PRJNA1301868 | liver |
| GSE315272 | Caenorhabditis elegans | Jan 15 2026 | PRJNA1395555 | whole worm |
| GSE303365 | Homo sapiens | Jan 14 2026 | PRJNA1294894 | |
| GSE253365 | Mus musculus | Jan 01 2026 | PRJNA1065416 | Brain |
| GSE315245 | Mus musculus | Jan 05 2026 | PRJNA1395480 | |
| GSE309965 | Mus musculus | Feb 18 2026 | PRJNA1364059 | Liver |
| GSE213034 | Mus musculus | Sep 10 2022 | PRJNA878815 | |
| GSE311046 | Homo sapiens | Feb 06 2026 | PRJNA1368597 | phoid cells type 2 sorted from PBMCs |
| GSE303889 | Homo sapiens | Jan 08 2026 | PRJNA1298167 | |
| GSE303407 | Saccharomyces cerevisiae | Jan 21 2026 | PRJNA1295354 | total cells |
| GSE297495 | Mus musculus | Jan 23 2026 | PRJNA1265082 | eric perivascular adipose tissue |
| GSE306418 | Anableps anableps | Nov 18 2025 | PRJNA1310599 | Head |
| GSE283670 | Homo sapiens | Jan 21 2026 | PRJNA1195015 | |
| GSE310479 | Homo sapiens | Jan 20 2026 | PRJNA1366387 | y human Donor lung fibroblasts |
| GSE298345 | Mus musculus | Feb 05 2026 | PRJNA1269000 | Adipose tissue |
| GSE261542 | Mus musculus | Jan 21 2026 | PRJNA1087408 | Back skin_Dorsal |
| GSE263687 | Mus musculus | Jan 23 2026 | PRJNA1098896 | primary lung tumor |
| GSE105048 | Mus musculus | Oct 16 2020 | PRJNA414470 | |
| GSE316931 | Arabidopsis thaliana | Feb 05 2026 | | whole seedling |
| GSE218285 | Homo sapiens | Oct 10 2023 | PRJNA903084 | .9 sorted cells from buffy coat |
| GSE282032 | Mus musculus | Nov 30 2025 | PRJNA1186687 | retina |
| GSE123945 | Mus musculus | Dec 17 2021 | PRJNA510363 | Retina/coroid |
| GSE220672 | Arvicanthis niloticus | Aug 10 2023 | PRJNA910951 | Whole Retinal Vasculature |
| GSE132731 | Mus musculus | Mar 22 2020 | PRJNA548871 | |
| GSE168717 | Mus musculus | Mar 12 2021 | PRJNA713767 | heart |
| GSE303224 | Mus musculus | Jan 01 2026 | PRJNA1294365 | lung |
| GSE214298 | Mus musculus | Sep 30 2022 | PRJNA884839 | |
| GSE253213 | Homo sapiens | Jan 14 2026 | PRJNA1064573 | peripheral blood |
| GSE172457 | Mus musculus | Apr 29 2022 | PRJNA723339 | |
| GSE308681 | Mus musculus | Jan 19 2026 | PRJNA1332460 | Bone marrow |
| GSE289188 | Homo sapiens | Feb 11 2026 | PRJNA1221882 | |
| GSE288394 | Mus musculus | Feb 06 2026 | PRJNA1217349 | uodenum villi_epithelium |
| GSE273656 | Mus musculus | Jan 30 2026 | PRJNA1142572 | heart |
| GSE315451 | Homo sapiens | Jan 06 2026 | PRJNA1397176 | Bone marrow |
| GSE291425 | Homo sapiens | Feb 25 2026 | PRJNA1233424 | |
| GSE303571 | Mus musculus | Feb 23 2026 | PRJNA1295845 | heart |
| GSE316097 | Heliophilum fasciatum | Jan 14 2026 | PRJNA1400591 | Strain |
| GSE198112 | Mus musculus | Jul 27 2022 | PRJNA813635 | perior Colliculus (Deep layer) |
| GSE173129 | Homo sapiens | Apr 23 2021 | PRJNA723860 | |
| GSE291264 | Homo sapiens | Jan 28 2026 | PRJNA1232684 | |
| GSE268463 | Mus musculus | Dec 06 2024 | PRJNA1117415 | Retina |
| GSE253330 | Mus musculus | Jan 11 2026 | PRJNA1065314 | Tumor |
| GSE283458 | Homo sapiens | Feb 11 2026 | PRJNA1193988 | |
| GSE282126 | Homo sapiens | Jan 20 2026 | PRJNA1187244 | Breast ( Breast Cancer) |
| GSE304788 | Arabidopsis thaliana | Feb 06 2026 | PRJNA1302545 | Rosette leaves |
| GSE148759 | Mus musculus | Oct 27 2020 | PRJNA625656 | retina |
| GSE281959 | Mus musculus | Dec 01 2024 | | retina |
| GSE230944 | Mus musculus | Jan 26 2026 | PRJNA962932 | heart |
| GSE248980 | Homo sapiens | Nov 30 2023 | | retina |
| GSE261461 | Homo sapiens | Jan 27 2026 | PRJNA1086997 | tal cortex (Brodmann area 9), left hemisphere |

| | | | | |
|---|---|---|---|---|
| GSE316183 | Escherichia coli | Jan 15 2026 | PRJNA1401771 | |
| GSE245545 | Homo sapiens | Oct 22 2023 | PRJNA1028965 | |
| GSE156305 | Mus musculus | Oct 04 2022 | PRJNA657405 | |
| GSE224230 | Folsomia candida | Jan 31 2026 | PRJNA930366 | whole body |
| GSE316812 | Mus musculus | Jan 21 2026 | PRJNA1404277 | Peritoneal cavity |
| GSE315738 | Homo sapiens | Jan 10 2026 | PRJNA1398870 | iPSC-astrocytes |
| GSE200476 | Oryctolagus cuniculus | Apr 09 2022 | PRJNA824789 | etinal Pigment Epithelium |
| GSE297395 | Homo sapiens | Feb 20 2026 | PRJNA1263755 | |
| GSE319070 | Homo sapiens | Feb 14 2026 | PRJNA1422039 | prostate |
| GSE314386 | Homo sapiens | Feb 27 2026 | PRJNA1390802 | breast cancer derived xenograft |
| GSE270244 | Oryza sativa | Jan 01 2026 | PRJNA1125584 | Leaves from 3-week-old |
| GSE239642 | Homo sapiens | Jan 29 2026 | PRJNA1000470 | |
| GSE304402 | Mus musculus | Jan 01 2026 | PRJNA1301254 | |
| GSE180616 | Homo sapiens | Oct 20 2021 | PRJNA748889 | |
| GSE255301 | Arabidopsis thaliana | Feb 25 2026 | PRJNA1074403 | seedlings |
| GSE316721 | Homo sapiens | Feb 25 2026 | PRJNA1403834 | Skin |
| GSE300601 | Mus musculus | Jan 09 2026 | PRJNA1281532 | small intestine |
| GSE308662 | lexandromys fortis;Mus musculu | Feb 18 2026 | PRJNA1331834 | Blood |
| GSE178725 | Mus musculus | Jun 24 2021 | PRJNA740196 | ouse embryonic fibroblasts |
| GSE317984 | Mus musculus | Jan 30 2026 | PRJNA1416261 | Tumor tissue |
| GSE284642 | Mus musculus | Dec 19 2024 | PRJNA1200000 | retina |
| GSE243709 | Mus musculus | Jan 12 2026 | PRJNA1019672 | |
| GSE305629 | Homo sapiens | Feb 06 2026 | PRJNA1307385 | in FFPE blocks from MIBC patient |
| GSE288415 | Gallus gallus | Jan 31 2026 | PRJNA1217392 | |
| GSE160797 | Mus musculus | Jan 19 2021 | PRJNA674526 | Retina |
| GSE316074 | Homo sapiens | Jan 13 2026 | PRJNA1400560 | |
| GSE309603 | Mus musculus | Jan 21 2026 | PRJNA1336520 | Liver |
| GSE190834 | Mus musculus | May 01 2022 | PRJNA788542 | Retina |
| GSE290584 | Homo sapiens | Feb 26 2026 | PRJNA1228668 | |
| GSE317136 | Homo sapiens | Feb 23 2026 | PRJNA1406153 | |
| GSE294787 | Homo sapiens | Jan 10 2026 | PRJNA1251602 | |
| GSE319935 | Homo sapiens | Feb 18 2026 | PRJNA1425549 | acent uninvolved liver tissue |
| GSE309042 | Mus musculus | Jan 01 2026 | PRJNA1333846 | Aorta |
| GSE211658 | Homo sapiens;Mus musculus | Aug 22 2022 | PRJNA871260 | Retina |
| GSE180702 | Homo sapiens | Nov 03 2022 | PRJNA749260 | |
| GSE304998 | Rattus norvegicus | Jan 12 2026 | PRJNA1303666 | Tumor |
| GSE316157 | Ascaris suum | Jan 12 2026 | | embryo |
| GSE189737 | Danio rerio | May 12 2023 | PRJNA784298 | |
| GSE308023 | Mus musculus | Feb 28 2026 | PRJNA1328600 | trigeminal ganglion |
| GSE278304 | Mus musculus | Sep 29 2024 | PRJNA1166296 | retina |
| GSE198943 | Mus musculus | Feb 14 2023 | PRJNA817541 | retina |
| GSE194285 | Macaca mulatta | Jan 24 2022 | PRJNA800052 | Retina |
| GSE247663 | Homo sapiens | Nov 26 2023 | PRJNA1040128 | |
| GSE292885 | Mus musculus | Feb 23 2026 | PRJNA1241802 | Kidney |
| GSE279430 | Homo sapiens | Feb 12 2026 | PRJNA1172688 | PBMC |
| GSE293308 | Homo sapiens;Mus musculus | May 10 2026 | PRJNA1243980 | |
| GSE313460 | Homo sapiens | Feb 28 2026 | PRJNA1379587 | |
| GSE318963 | Homo sapiens | Feb 09 2026 | PRJNA1420894 | |
| GSE255363 | Nothobranchius furzeri | Aug 27 2024 | PRJNA1074746 | Neural Retina |
| GSE221861 | Homo sapiens | Dec 29 2022 | PRJNA916513 | retina |
| GSE297237 | Mus musculus | Jan 21 2026 | PRJNA1263376 | skeletal muscle |
| GSE222536 | Mus musculus | Jan 01 2026 | PRJNA922478 | 2-cell embryo |
| GSE285114 | Mus musculus | Jan 21 2026 | PRJNA1201199 | Knee joint |
| GSE307937 | Homo sapiens | Feb 14 2026 | PRJNA1327854 | |
| GSE269597 | Mus musculus | Jan 01 2025 | PRJNA1122597 | eye cup |
| GSE305401 | Homo sapiens | Jan 21 2026 | PRJNA1305462 | Stem cell |
| GSE158335 | Sus scrofa | Sep 23 2020 | PRJNA664933 | |
| GSE316021 | Homo sapiens | Jan 16 2026 | PRJNA1400483 | Ovary |
| GSE303949 | Kluyveromyces lactis | Jan 22 2026 | PRJNA1298293 | |
| GSE253949 | Mus musculus | Oct 09 2024 | PRJNA1067979 | PE-choroid-scleral complex |
| GSE267405 | Homo sapiens | May 17 2024 | PRJNA1111430 | retinal organoids |
| GSE162352 | Mus musculus | Feb 18 2021 | PRJNA681579 | fibroblasts |
| GSE286287 | Mus musculus | Feb 17 2026 | PRJNA1208529 | |

| | | | | |
|---|---|---|---|---|
| GSE308162 | Homo sapiens | Feb 09 2026 | PRJNA1329123 | plasma |
| GSE284620 | Mus musculus | Jan 05 2026 | PRJNA1199944 | Mesenteric Lymph Node |
| GSE219252 | Mus musculus | Jun 07 2023 | PRJNA907803 | RPE-choroid |
| GSE312336 | Homo sapiens | Feb 05 2026 | PRJNA1373113 | heral Blood Mononuclear Cells |
| GSE316135 | Mus musculus | Jan 16 2026 | PRJNA1401693 | Aorta |
| GSE102207 | Homo sapiens | Jul 27 2020 | PRJNA397006 | |
| GSE281561 | Homo sapiens | Jan 31 2026 | PRJNA1184725 | |
| GSE255515 | Mus musculus | Jan 26 2026 | PRJNA1075140 | |
| GSE304709 | Mus musculus | Jun 30 2026 | PRJNA1302379 | Retina |
| GSE318936 | Mus musculus | Feb 12 2026 | PRJNA1420858 | testis |
| GSE142449 | Homo sapiens | Feb 17 2020 | PRJNA597043 | |
| GSE295977 | Homo sapiens | Feb 05 2026 | PRJNA1257074 | heral Blood Mononuclear Cells |
| GSE301074 | Mus musculus | Jan 12 2026 | | ovary |
| GSE223321 | Mus musculus | Mar 30 2023 | PRJNA925805 | retina |
| GSE280500 | Candida albicans SC5314 | Jan 05 2026 | PRJNA1178708 | |
| GSE319957 | Macaca mulatta | Feb 23 2026 | PRJNA1425778 | |
| GSE289257 | Homo sapiens | Feb 06 2026 | PRJNA1222277 | |
| GSE246657 | Mus musculus | Feb 09 2026 | PRJNA1034073 | Brain |
| GSE313357 | Mus musculus | Jan 27 2026 | PRJNA1379312 | Hippocampus |
| GSE158543 | Homo sapiens | Dec 31 2023 | PRJNA665538 | |
| GSE178653 | Mus musculus | Jun 23 2021 | PRJNA739908 | retinal organoids |
| GSE302485 | Mus musculus | Feb 06 2026 | PRJNA1291079 | Epidermis |
| GSE311934 | Mus musculus | Feb 06 2026 | PRJNA1371954 | skin |
| GSE325478 | Mus musculus | Apr 02 2026 | PRJNA1440256 | retina |
| GSE319960 | Homo sapiens | Feb 24 2026 | PRJNA1425790 | Plasma |
| GSE296417 | Gallus gallus | Feb 13 2026 | PRJNA1259254 | Hematopoietic |
| GSE202050 | Drosophila melanogaster | Dec 31 2022 | PRJNA834090 | |
| GSE306875 | Klebsiella pneumoniae | Feb 09 2026 | PRJNA1312835 | |
| GSE218638 | Homo sapiens | Nov 27 2022 | PRJNA904736 | Pars plana |
| GSE281556 | Homo sapiens | Jan 06 2026 | PRJNA1184717 | |
| GSE274897 | Mus musculus | Jan 31 2026 | PRJNA1148424 | Liver |
| GSE318901 | Homo sapiens | Feb 16 2026 | PRJNA1420790 | whole blood |
| GSE285755 | Danio rerio | Feb 12 2026 | PRJNA1205765 | tumor in adrenal gland |
| GSE309775 | Homo sapiens;Mus musculus | Jan 15 2026 | PRJNA1363152 | blood |
| GSE164893 | Homo sapiens | Jan 10 2026 | PRJNA692392 | |
| GSE253666 | 3ombyx mandarina;Bombyx mor | Jan 14 2026 | PRJNA1066737 | wing disc |
| GSE291732 | Homo sapiens | Jan 23 2026 | PRJNA1235153 | Pancreas |
| GSE317483 | Homo sapiens | Jan 30 2026 | PRJNA1413410 | |
| GSE270816 | Mus musculus | Jan 13 2026 | PRJNA1128567 | FL-BMDC |
| GSE271684 | Homo sapiens | Jan 07 2026 | PRJNA1132919 | |
| GSE310246 | Mus musculus | Jan 23 2026 | PRJNA1365525 | Cochlea |
| GSE275809 | Drosophila melanogaster | Nov 30 2024 | PRJNA1152986 | |
| GSE315259 | Mus musculus | Jan 06 2026 | | |
| GSE208143 | Homo sapiens | Jul 18 2022 | PRJNA859173 | Tumor |
| GSE310759 | Mus musculus | Jan 01 2026 | PRJNA1367040 | |
| GSE141725 | Mus musculus | Dec 01 2020 | PRJNA594523 | retina |
| GSE318570 | Mus musculus | Feb 18 2026 | PRJNA1419241 | hippocampus |
| GSE332948 | Mus musculus | Jun 01 2026 | | brain |
| GSE315024 | Bos taurus | Jan 06 2026 | PRJNA1394894 | female |
| GSE291050 | Homo sapiens | Jan 13 2026 | PRJNA1231465 | Prostate |
| GSE227688 | Homo sapiens | Jan 28 2026 | PRJNA946650 | |
| GSE268657 | Mus musculus | Feb 07 2026 | PRJNA1118209 | of Langerhans, tdTomato+ cells |
| GSE289973 | Oryctolagus cuniculus | Jan 12 2026 | PRJNA1225669 | artery |
| GSE222388 | Homo sapiens | Jan 12 2023 | PRJNA921856 | retinal epithelium |
| GSE295253 | Homo sapiens;Rattus norvegicus | Jan 16 2026 | PRJNA1253942 | hNPCs |
| GSE222697 | Mus musculus | Jan 01 2026 | PRJNA923038 | Liver |
| GSE315982 | Mus musculus | Jan 09 2026 | PRJNA1400058 | testis |
| GSE335334 | Ambystoma mexicanum | Jun 16 2026 | PRJNA1477939 | Retina |
| GSE303133 | Arabidopsis thaliana | Jan 29 2026 | PRJNA1294226 | Seedlings |
| GSE313844 | Rhodotorula toruloides | Jan 20 2026 | PRJNA1381663 | |
| GSE328399 | Homo sapiens | Apr 24 2026 | PRJNA1454901 | heral blood samples of human |
| GSE289944 | Arabidopsis thaliana | Jan 29 2026 | PRJNA1225621 | Seedlings |
| GSE294567 | Homo sapiens | Jan 30 2026 | PRJNA1250507 | Tonsil |

| | | | | |
|---|---|---|---|---|
| GSE319526 | Homo sapiens | Feb 18 2026 | PRJNA1423676 | |
| GSE314072 | Homo sapiens | Jan 01 2026 | PRJNA1389917 | PBMC |
| GSE315771 | Mus musculus | Jan 07 2026 | PRJNA1398936 | liver |
| GSE273576 | Homo sapiens | Aug 12 2024 | | |
| GSE270821 | Mus musculus | Jan 13 2026 | PRJNA1128572 | |
| GSE292582 | Homo sapiens | Jan 28 2026 | PRJNA1240257 | |
| GSE316978 | Homo sapiens | Jan 22 2026 | PRJNA1405627 | |
| GSE299881 | Arabidopsis thaliana | Feb 28 2026 | PRJNA1277219 | whole seedlings |
| GSE297082 | Homo sapiens | Jan 01 2026 | PRJNA1262484 | or tissue of pancreatic cancer |
| GSE236566 | Homo sapiens | Jul 12 2023 | PRJNA991576 | Optic Nerve |
| GSE290647 | Ixodes scapularis | Feb 20 2026 | | salivary glands |
| GSE195438 | Mus musculus | Jan 29 2022 | PRJNA800666 | Macrophage |
| GSE315130 | Homo sapiens;Mus musculus | Jan 31 2026 | PRJNA1395035 | |
| GSE161516 | Arabidopsis thaliana | Mar 22 2021 | PRJNA678632 | Seedling |
| GSE228370 | Homo sapiens | Mar 31 2023 | PRJNA949575 | fetal eye |
| GSE327656 | Mus musculus | Apr 17 2026 | PRJNA1451773 | gastrocnemius muscle |
| GSE280852 | Homo sapiens | Feb 17 2026 | PRJNA1180922 | liver |
| GSE299672 | Homo sapiens | Feb 09 2026 | PRJNA1275959 | |
| GSE294326 | Cucumis sativus | Jan 19 2026 | PRJNA1249024 | shoot apical meristem |
| GSE291611 | ıs;Felis catus;Gallus gallus;Melop | Sep 02 2025 | PRJNA1234492 | Retina |
| GSE172101 | Mus musculus | Apr 15 2021 | PRJNA722013 | |
| GSE207802 | Homo sapiens | Jan 01 2023 | PRJNA857087 | fetal retina |
| GSE175937 | Homo sapiens | Nov 13 2023 | PRJNA734326 | |
| GSE271954 | Homo sapiens | Feb 25 2026 | PRJNA1134199 | |
| GSE307224 | Mus musculus | Feb 11 2026 | | |
| GSE216676 | Mus musculus | Apr 01 2025 | PRJNA894955 | Retina |
| GSE315896 | Mus musculus | Jan 13 2026 | PRJNA1399761 | spleen |
| GSE186471 | Homo sapiens | Mar 31 2022 | PRJNA773964 | |
| GSE267935 | Mus musculus | Jan 23 2025 | PRJNA1113669 | retina |
| GSE307365 | Mus musculus | Nov 15 2025 | PRJNA1316872 | Brain |
| GSE297389 | Homo sapiens | Jan 13 2026 | PRJNA1263737 | |
| GSE285582 | Mus musculus | Jan 21 2026 | PRJNA1204925 | m flash-frozen powderized whole pancreas |
| GSE281781 | Homo sapiens;Mus musculus | Feb 06 2026 | PRJNA1185699 | Bone |
| GSE318525 | Caenorhabditis elegans | Feb 10 2026 | PRJNA1419159 | Whole organism |
| GSE319739 | Escherichia coli | Feb 20 2026 | PRJNA1425039 | |
| GSE139947 | Homo sapiens | Oct 01 2021 | PRJNA587715 | |
| GSE316839 | Mus musculus | Feb 18 2026 | PRJNA1404823 | pancreatic islets |
| GSE294344 | Homo sapiens | Feb 17 2026 | PRJNA1249068 | Lung |
| GSE137705 | Mus musculus | Jul 14 2021 | PRJNA566356 | Retina |
| GSE319178 | Mus musculus | Feb 15 2026 | PRJNA1422509 | breast |
| GSE318364 | Drosophila melanogaster | Feb 27 2026 | PRJNA1418638 | Ovary |
| GSE317297 | Mus musculus | Feb 07 2026 | PRJNA1406703 | ovary |
| GSE237207 | Didelphis marsupialis | Oct 01 2023 | PRJNA994170 | Retina |
| GSE311351 | Mus musculus | Jan 20 2026 | PRJNA1369185 | o Adipose tissue macrophages |
| GSE316594 | Sus scrofa | Jan 30 2026 | PRJNA1403464 | cine preimplantation embryo |
| GSE208760 | Mus musculus | May 01 2023 | PRJNA860959 | |
| GSE212108 | Homo sapiens | Apr 23 2023 | PRJNA874034 | Pigmented epithelium; Retina |
| GSE240465 | Mus musculus | Oct 12 2023 | PRJNA1003849 | Retina |
| GSE289832 | Mus musculus | Feb 17 2026 | | |
| GSE313932 | Homo sapiens | Feb 24 2026 | PRJNA1387386 | |
| GSE283550 | Mus musculus | Jan 08 2026 | PRJNA1194505 | macrophages |
| GSE288853 | Mus musculus | Feb 09 2026 | PRJNA1219632 | Retina |
| GSE315318 | Mesocricetus auratus | Jan 15 2026 | PRJNA1395632 | |
| GSE272303 | Mus musculus | Feb 11 2026 | PRJNA1136138 | Coronal brain section |
| GSE302024 | Rattus norvegicus | Feb 11 2026 | PRJNA1288371 | brain |
| GSE231494 | Sus scrofa | Jan 29 2026 | PRJNA966156 | Lung tissue |
| GSE296927 | Homo sapiens | Feb 09 2026 | PRJNA1261943 | |
| GSE318353 | Homo sapiens | Feb 06 2026 | | |
| GSE303054 | Mus musculus | Oct 16 2025 | PRJNA1293418 | optic nerve |
| GSE318420 | Homo sapiens | Feb 04 2026 | PRJNA1418700 | patocellular Carcinoma (HCC) |
| GSE313923 | Mus musculus | Feb 06 2026 | PRJNA1387362 | P3 digit |
| GSE290603 | Homo sapiens | Feb 26 2026 | PRJNA1228960 | |
| GSE314677 | Rattus norvegicus | Jan 20 2026 | PRJNA1392439 | kidney |

| | | | | |
|---|---|---|---|---|
| GSE294362 | Mus musculus | Jan 07 2026 | PRJNA1249087 | Spleen |
| GSE240316 | | Aug 28 2023 | | |
| GSE270946 | Homo sapiens | Jan 23 2026 | PRJNA1129086 | |
| GSE203481 | Mus | Dec 05 2022 | PRJNA840925 | inferior-cone |
| GSE309829 | Homo sapiens | Jan 26 2026 | | Heart Organoid |
| GSE318342 | Homo sapiens | Feb 06 2026 | PRJNA1418615 | |
| GSE261732 | human feces metagenome | Dec 05 2024 | PRJNA1088508 | Stool |
| GSE263879 | Mus musculus | Jan 22 2026 | PRJNA1099666 | Liver |
| GSE198362 | Homo sapiens | Nov 16 2022 | PRJNA814699 | |
| GSE292293 | Mus musculus | Jan 22 2026 | PRJNA1237837 | |
| GSE155829 | Homo sapiens | Jan 08 2021 | PRJNA655716 | |
| GSE286076 | Homo sapiens | Feb 01 2025 | | Eye; Pigmented epithelium; Retina |
| GSE302002 | Homo sapiens | Jan 01 2026 | PRJNA1288267 | |
| GSE315621 | Enterobacteria phage T6 | Feb 05 2026 | PRJNA1398346 | |
| GSE289185 | Homo sapiens | Feb 07 2026 | PRJNA1221879 | Tumor tissue |
| GSE302760 | Mus musculus | Feb 06 2026 | PRJNA1292114 | |
| GSE319314 | Homo sapiens | Feb 16 2026 | PRJNA1422813 | |
| GSE261578 | Homo sapiens | Mar 12 2026 | PRJNA1087708 | |
| GSE303063 | Mus musculus | Feb 09 2026 | PRJNA1293428 | testis |
| GSE304976 | Homo sapiens | Jan 25 2026 | | |
| GSE272206 | Mus musculus | Dec 31 2024 | PRJNA1135691 | |
| GSE232448 | Homo sapiens | Sep 25 2025 | PRJNA972001 | lorsal forebrain organoids |
| GSE277219 | Homo sapiens | May 14 2025 | PRJNA1161382 | |
| GSE310202 | Homo sapiens;Mus musculus | Jan 29 2026 | PRJNA1365264 | Skin |
| GSE298743 | Homo sapiens | Jun 06 2025 | PRJNA1271036 | null |
| GSE160138 | Danio rerio | Oct 27 2020 | PRJNA672100 | Retina |
| GSE268613 | Mus musculus | Feb 11 2026 | PRJNA1117902 | liver |
| GSE291014 | Homo sapiens | Jan 20 2026 | PRJNA1231395 | Colon |
| GSE243837 | Mus musculus | Jan 12 2026 | PRJNA1020138 | |
| GSE306440 | Homo sapiens | Feb 11 2026 | PRJNA1310636 | |
| GSE230949 | Mus musculus | Jan 26 2026 | PRJNA962923 | heart |
| GSE287103 | Mus musculus | Jan 15 2026 | PRJNA1211130 | bone marrow |
| GSE294523 | Homo sapiens | Jan 14 2026 | PRJNA1250443 | |
| GSE317392 | Mus musculus | Jan 31 2026 | | Colon |
| GSE314754 | Mus musculus | Jun 18 2026 | PRJNA1392547 | Retina |
| GSE315548 | Homo sapiens | Feb 08 2026 | PRJNA1398246 | cell line |
| GSE287892 | Homo sapiens | Jan 01 2026 | PRJNA1214870 | gastric cancer |
| GSE316491 | Homo sapiens | Jan 14 2026 | PRJNA1402998 | |
| GSE246491 | Homo sapiens | Jan 15 2026 | PRJNA1033336 | Blood |
| GSE315735 | Homo sapiens | Jan 30 2026 | PRJNA1398869 | |
| GSE199575 | Homo sapiens | Nov 17 2022 | PRJNA820749 | ived mesodermal cell aggregates |
| GSE174488 | Drosophila melanogaster | Dec 31 2021 | PRJNA730035 | |
| GSE312861 | Mus musculus | Jan 08 2026 | PRJNA1377527 | brain |
| GSE184817 | Mus musculus | Jan 01 2022 | PRJNA766601 | photoreceptor cell |
| GSE311168 | Mus musculus | Jan 05 2026 | PRJNA1368769 | small intestine epithelium |
| GSE222956 | Mus musculus | Feb 15 2023 | PRJNA924281 | retina |
| GSE318320 | Pinus pinaster | Feb 07 2026 | PRJNA1418560 | Stem |
| GSE271945 | Canis lupus familiaris | Oct 01 2024 | PRJNA1134167 | Neuroretina |
| GSE298635 | Homo sapiens | Feb 26 2026 | PRJNA1270065 | |
| GSE319341 | Mus musculus | Feb 12 2026 | PRJNA1423050 | cortex |
| GSE253529 | Mus musculus | Nov 12 2024 | PRJNA1066215 | |
| GSE156756 | Mus musculus | Mar 29 2021 | PRJNA658977 | Retina |
| GSE310416 | Homo sapiens | Feb 23 2026 | PRJNA1365917 | |
| GSE318442 | Homo sapiens | Feb 11 2026 | PRJNA1418734 | |
| GSE313941 | Mus musculus | Jan 27 2026 | | Spleen |
| GSE291162 | Mus musculus | Jan 07 2026 | PRJNA1232126 | Brain |
| GSE299101 | Homo sapiens | Feb 18 2026 | PRJNA1272585 | pithelial Adenocarcinoma |
| GSE310421 | Homo sapiens | Feb 23 2026 | PRJNA1365914 | |
| GSE287075 | Homo sapiens;Mus musculus | Jan 20 2026 | PRJNA1211098 | |
| GSE303001 | Mus musculus | Feb 09 2026 | PRJNA1293336 | testis |
| GSE253001 | Mus musculus | Jan 19 2026 | PRJNA1063626 | aortic tissues |
| GSE207824 | Homo sapiens | Feb 01 2023 | PRJNA857290 | retinal organoid |
| GSE283276 | Mus musculus | Dec 02 2024 | PRJNA1193014 | retina |

| | | | | |
|---|---|---|---|---|
| GSE276339 | Saposhnikovia divaricata | Nov 30 2024 | PRJNA1156701 | leaves |
| GSE303990 | Mus musculus | Jan 13 2026 | PRJNA1298599 | Retina |
| GSE317586 | Homo sapiens | Feb 18 2026 | PRJNA1414503 | |
| GSE247586 | Homo sapiens;Mus musculus | Jun 26 2024 | PRJNA1039768 | retina enriched sample |
| GSE256263 | Mus musculus | Sep 29 2024 | PRJNA1078801 | brain |
| GSE318608 | Mus musculus | Feb 10 2026 | PRJNA1419304 | |
| GSE313785 | Mus musculus | Jan 31 2026 | PRJNA1381596 | 71+ EB-derived erythroid cells, originally derived from ESC |
| GSE291328 | Homo sapiens | Jan 01 2026 | PRJNA1232831 | CSF |
| GSE280843 | Homo sapiens | Apr 23 2025 | PRJNA1180903 | Gastric |
| GSE313855 | Oryza sativa | Jan 19 2026 | PRJNA1381683 | leaf |
| GSE305975 | Mus musculus | Jan 21 2026 | PRJNA1308574 | lymph node |
| GSE287449 | Homo sapiens | Jan 19 2026 | PRJNA1212783 | |
| GSE315993 | Ambystoma mexicanum | Jan 15 2026 | PRJNA1400073 | forelimb |
| GSE318234 | Mus musculus | Feb 07 2026 | PRJNA1416951 | pineal |
| GSE232135 | Homo sapiens | May 15 2023 | PRJNA971113 | |
| GSE266910 | Physcomitrium patens | Jan 26 2026 | PRJNA1108707 | seedlings |
| GSE319500 | Homo sapiens | Feb 17 2026 | PRJNA1423630 | |
| GSE148063 | Mus musculus | Feb 08 2021 | PRJNA622978 | Ocular area |
| GSE309968 | Mus musculus | Feb 20 2026 | PRJNA1364065 | hippocampus |
| GSE288092 | Mus musculus | Jan 14 2026 | PRJNA1216182 | Lung |
| GSE224022 | Homo sapiens | Feb 03 2023 | PRJNA929438 | Normal_retina |
| GSE192814 | Homo sapiens | Oct 28 2022 | PRJNA793258 | |
| GSE160310 | Homo sapiens | Apr 19 2021 | PRJNA672925 | Retina Macula |
| GSE285948 | Mus musculus | Jan 04 2026 | PRJNA1207027 | heart tissue |
| GSE213244 | Mus musculus | Dec 07 2022 | PRJNA880244 | cerebellum |
| GSE207930 | Mus musculus | Jul 11 2022 | PRJNA857697 | Retinal Tissue |
| GSE287161 | Mus musculus | Feb 16 2026 | PRJNA1211524 | Mediobasal Hypothalamus |
| GSE267255 | Homo sapiens | Feb 23 2026 | PRJNA1110755 | Spleen |
| GSE293106 | Homo sapiens | Jan 28 2026 | PRJNA1242824 | Breast |
| GSE182483 | Mus musculus | Aug 22 2021 | PRJNA756372 | retina |
| GSE134744 | Homo sapiens | Jul 23 2020 | PRJNA556257 | |
| GSE146411 | Gallus gallus | Oct 31 2020 | PRJNA610527 | |
| GSE305251 | Homo sapiens | Nov 04 2025 | PRJNA1304923 | umbilical vein |
| GSE126178 | Homo sapiens | Feb 14 2026 | PRJNA521250 | lymphoblastoid cell line |
| GSE197165 | Homo sapiens | Sep 07 2022 | PRJNA809304 | |
| GSE293664 | Homo sapiens | Feb 19 2026 | PRJNA1245462 | |
| GSE158718 | Mus musculus | Sep 01 2021 | PRJNA666343 | retina |
| GSE196104 | Mus musculus | Jul 13 2022 | PRJNA803305 | retina |
| GSE243413 | Mus musculus | Feb 15 2024 | PRJNA1018454 | retina |
| GSE288738 | Homo sapiens | Feb 09 2026 | PRJNA1219172 | |
| GSE316124 | Mus musculus | Jan 16 2026 | PRJNA1401577 | forelimb |
| GSE255504 | Homo sapiens | Feb 08 2026 | PRJNA1075126 | |
| GSE163563 | Rattus norvegicus | Jul 28 2021 | PRJNA686777 | retina |
| GSE301707 | Mus musculus | Jan 07 2026 | | head |
| GSE296353 | Homo sapiens | Feb 10 2026 | PRJNA1258970 | |
| GSE221797 | Mus musculus | Jan 14 2026 | PRJNA916226 | Medial prefrontal cortex |
| GSE285980 | Mus musculus | Jan 07 2025 | PRJNA1207114 | retina |
| GSE277284 | Mus musculus | Jan 13 2026 | PRJNA1161657 | |
| GSE316646 | Mus musculus | Jan 20 2026 | | Liver |
| GSE317145 | Mus musculus | Feb 16 2026 | PRJNA1406167 | |
| GSE312445 | Homo sapiens | Jan 15 2026 | PRJNA1373325 | c and GABAergic neurons derived from iPS cells |
| GSE236097 | Homo sapiens | Feb 07 2025 | PRJNA988887 | Organoid |
| GSE319696 | Mus musculus | Feb 17 2026 | PRJNA1424970 | |
| GSE318804 | Mus musculus | Feb 11 2026 | PRJNA1420648 | BMDM |
| GSE202155 | Mus musculus | Jun 28 2022 | PRJNA834803 | retina |
| GSE292073 | Glycine max | Feb 11 2026 | PRJNA1236645 | seed |
| GSE270200 | Mus musculus | Dec 31 2024 | PRJNA1125403 | Choroid |
| GSE304324 | Homo sapiens | Jan 30 2026 | PRJNA1300376 | |
| GSE312204 | Homo sapiens | Feb 10 2026 | PRJNA1372664 | normal colon |
| GSE283289 | Homo sapiens | Jan 31 2026 | PRJNA1193049 | Colon |
| GSE288304 | Homo sapiens | Feb 09 2026 | PRJNA1216886 | early embryo |
| GSE308611 | Homo sapiens | Jan 28 2026 | PRJNA1331709 | Lung |
| GSE316765 | Oryza sativa Japonica Group | Jan 25 2026 | PRJNA1404168 | |

| | | | | |
|---|---|---|---|---|
| GSE293983 | Homo sapiens | Jan 03 2026 | PRJNA1247521 | Eye |
| GSE254446 | Homo sapiens | Feb 11 2026 | PRJNA1070586 | kidney |
| GSE306783 | Mus musculus | Jan 09 2026 | PRJNA1312105 | |
| GSE277390 | Mus musculus | Sep 25 2024 | PRJNA1162090 | Retina |
| GSE316752 | Homo sapiens | Jan 16 2026 | PRJNA1404144 | |
| GSE298839 | Mus musculus | Feb 22 2026 | PRJNA1271471 | |
| GSE315878 | Homo sapiens | Jan 13 2026 | PRJNA1399478 | |
| GSE300881 | Drosophila melanogaster | Jan 20 2026 | PRJNA1282577 | Ring Gland |
| GSE301613 | Homo sapiens | Jun 01 2026 | PRJNA1285789 | Human Umbilical Vein |
| GSE317733 | Homo sapiens | Jan 27 2026 | PRJNA1415118 | Prostate |
| GSE305410 | Mus musculus | Jan 14 2026 | PRJNA1305485 | Parietal bone |
| GSE297544 | Rattus norvegicus | Jan 11 2026 | | mammary tumor |
| GSE140689 | Mus musculus | Sep 30 2020 | PRJNA590512 | |
| GSE298294 | Mus musculus | Jan 21 2026 | PRJNA1268903 | thymus |
| GSE310926 | Homo sapiens | Feb 24 2026 | PRJNA1367545 | |
| GSE318972 | Mus musculus | Feb 12 2026 | PRJNA1420900 | Pancreatic islet |
| GSE259219 | Homo sapiens | Mar 01 2024 | | Retina |
| GSE305551 | Drosophila melanogaster | Aug 25 2025 | PRJNA1306499 | eye |
| GSE251752 | Homo sapiens | Aug 28 2024 | PRJNA1055163 | |
| GSE296306 | Homo sapiens | Jun 18 2026 | PRJNA1258898 | |
| GSE315939 | Homo sapiens | Jan 12 2026 | PRJNA1399985 | |
| GSE202741 | Mus musculus | Feb 15 2023 | PRJNA837193 | retina |
| GSE289213 | Homo sapiens | Feb 08 2026 | PRJNA1221911 | |
| GSE117956 | Homo sapiens | Feb 01 2026 | PRJNA483793 | retina |
| GSE317110 | Rattus norvegicus | Jan 26 2026 | PRJNA1406029 | Intestinal contents |
| GSE283813 | Homo sapiens | Feb 15 2026 | PRJNA1195609 | |
| GSE309064 | Mus musculus | Jan 28 2026 | PRJNA1333886 | prostate |
| GSE297678 | Rattus norvegicus | Jan 21 2026 | PRJNA1265680 | |
| GSE308278 | Mus musculus | Jan 14 2026 | PRJNA1330046 | |
| GSE319924 | Homo sapiens | Feb 18 2026 | PRJNA1425531 | Stroma |
| GSE304507 | Canis lupus familiaris | Jan 08 2026 | PRJNA1301401 | |
| GSE289224 | Homo sapiens | Feb 23 2026 | PRJNA1221931 | Spleen |
| GSE267562 | Homo sapiens | Jan 24 2026 | PRJNA1111854 | ICM |
| GSE168193 | Homo sapiens | Mar 04 2021 | PRJNA706436 | |
| GSE315580 | Homo sapiens | Jan 19 2026 | PRJNA1398288 | |
| GSE222333 | Homo sapiens | Jan 02 2026 | PRJNA919026 | |
| GSE297432 | Homo sapiens | Jan 13 2026 | PRJNA1263833 | |
| GSE289618 | Mus musculus | Jan 12 2026 | PRJNA1223834 | liver |
| GSE316218 | Mus musculus | Jan 31 2026 | PRJNA1401819 | Liver |
| GSE307973 | Homo sapiens | Feb 13 2026 | PRJNA1327902 | Stem cells |
| GSE317004 | Mus musculus | Feb 10 2026 | PRJNA1405665 | heart |
| GSE316707 | Mus musculus | Feb 12 2026 | | liver |
| GSE303583 | Mus musculus | Feb 25 2026 | PRJNA1295872 | Liver |
| GSE299483 | Homo sapiens | Aug 31 2025 | PRJNA1274714 | |
| GSE301646 | Mus musculus | Feb 14 2026 | PRJNA1285916 | Pancreas |
| GSE223513 | Homo sapiens | Sep 18 2024 | PRJNA926620 | breast |
| GSE229682 | Homo sapiens | Nov 03 2023 | PRJNA955281 | Retinal organoid |
| GSE284131 | Petromyzon marinus | Aug 18 2025 | PRJNA1197415 | retina |
| GSE301359 | Homo sapiens | Feb 17 2026 | PRJNA1284842 | |
| GSE306925 | Mus musculus | Jan 07 2026 | PRJNA1313934 | Hippocampus |
| GSE222227 | Mus musculus | Jan 01 2026 | PRJNA918635 | |
| GSE267169 | Mus musculus | Feb 23 2026 | PRJNA1109989 | mammary gland |
| GSE297039 | Homo sapiens | Feb 11 2026 | PRJNA1262428 | Peripheral blood |
| GSE315494 | Homo sapiens | Jan 28 2026 | | PBMC |
| GSE222738 | Mus musculus | Jan 01 2026 | | |
| GSE265940 | Homo sapiens | Jun 04 2024 | PRJNA1105010 | |
| GSE301882 | Homo sapiens | Jan 24 2026 | PRJNA1287694 | human 8C embryo |
| GSE235366 | Homo sapiens | Aug 08 2023 | PRJNA985752 | |
| GSE311864 | Oryza sativa | Jan 26 2026 | PRJNA1371868 | 14-day-old seedlings |
| GSE223246 | Mus musculus;Rattus norvegicus | Feb 11 2026 | PRJNA925476 | |
| GSE268327 | Mus musculus | Feb 23 2026 | PRJNA1116221 | Brain (prefrontal cortex) |
| GSE158799 | Mus musculus | Feb 22 2021 | PRJNA666571 | Retina |
| GSE318000 | Homo sapiens | Feb 25 2026 | PRJNA1416306 | cervical cancer |

| | | | | |
|---|---|---|---|---|
| GSE267938 | Mus musculus | Jan 23 2025 | PRJNA1113667 | retina |
| GSE300923 | Mus musculus | Jan 19 2026 | PRJNA1282683 | intestinal tissue |
| GSE294020 | Homo sapiens | Jan 21 2026 | PRJNA1247880 | |
| GSE172230 | Mus musculus | Oct 11 2022 | PRJNA722495 | |
| GSE275695 | Homo sapiens | Feb 20 2026 | PRJNA1152585 | |
| GSE275866 | Sus scrofa | Jan 06 2026 | PRJNA1153395 | luced pluripotent stem cells |
| GSE318762 | Mus musculus | Feb 11 2026 | PRJNA1419895 | heart |
| GSE295041 | Homo sapiens | Jan 15 2026 | PRJNA1252700 | mary ovarian cancer tissues |
| GSE283161 | Homo sapiens | Jan 14 2026 | PRJNA1192258 | breast |
| GSE279688 | Homo sapiens | Feb 25 2026 | PRJNA1174084 | |
| GSE281293 | Homo sapiens | Feb 24 2026 | PRJNA1183339 | |
| GSE252422 | Homo sapiens;Mus musculus | Jan 01 2026 | PRJNA1060622 | cervical tissue |
| GSE285901 | Homo sapiens | Jan 05 2026 | PRJNA1206761 | |
| GSE318755 | Mus musculus | Jun 23 2026 | PRJNA1419876 | Choroid |
| GSE291008 | Homo sapiens | Jun 03 2025 | PRJNA1231369 | |
| GSE287128 | Zea mays | Jan 10 2026 | PRJNA1211164 | primary and 2 seminal roots per seedling) |
| GSE221209 | Mus musculus | Apr 30 2023 | PRJNA913195 | Retina |
| GSE188957 | Mus musculus | Jan 01 2024 | PRJNA780895 | Liver |
| GSE287437 | Cavia porcellus | Jan 31 2026 | PRJNA1212750 | primary visual cortex |
| GSE284960 | Mus musculus | Feb 21 2026 | PRJNA1200728 | CD43- Splenic B cells |
| GSE316834 | Homo sapiens | Jan 27 2026 | PRJNA1404323 | Cartilage |
| GSE256143 | Homo sapiens | Feb 20 2026 | PRJNA1078318 | |
| GSE277326 | Homo sapiens | Jan 15 2025 | PRJNA1161713 | macula |
| GSE249175 | Mus musculus | Feb 18 2026 | PRJNA1047595 | ureter |
| GSE265994 | Mus musculus | May 21 2024 | PRJNA1105098 | Optic Nerve Head |
| GSE199162 | Gallus gallus | Jun 30 2022 | PRJNA818671 | |
| GSE312969 | Ovis aries | Feb 12 2026 | PRJNA1377679 | pituitary |
| GSE282985 | Homo sapiens | Jan 14 2026 | PRJNA1191415 | |
| GSE289808 | llus pleuropneumoniae serovar | Feb 28 2026 | PRJNA1224859 | Bacteria |
| GSE111794 | Homo sapiens | Mar 13 2021 | PRJNA438150 | |
| GSE318885 | Homo sapiens | Feb 16 2026 | PRJNA1420763 | prostate |
| GSE286217 | Mus musculus | Jan 15 2026 | PRJNA1208436 | bone-marrow |
| GSE300837 | Homo sapiens | Feb 25 2026 | PRJNA1282181 | blood |
| GSE291434 | Mus musculus | Jan 19 2026 | PRJNA1233585 | spleen and bone marrow |
| GSE318114 | Homo sapiens | Feb 20 2026 | PRJNA1416689 | Skin |
| GSE148374 | Macaca mulatta | May 19 2020 | PRJNA624095 | caque trabecular meshwork |
| GSE292954 | Bos taurus | Jan 26 2026 | PRJNA1242229 | Endometrium |
| GSE287523 | Mus musculus | Jan 22 2026 | PRJNA1213143 | Lung |
| GSE303557 | Homo sapiens | Jan 24 2026 | PRJNA1295814 | uman embryonic stem cell |
| GSE161733 | Homo sapiens | Feb 26 2021 | PRJNA679272 | |
| GSE244374 | omo sapiens;unidentified plasm | Jun 24 2024 | PRJNA1022486 | plasmid |
| GSE189822 | Homo sapiens | Oct 11 2023 | PRJNA784597 | .9 sorted cells from buffy coat |
| GSE306735 | Mus musculus | Jan 13 2026 | PRJNA1311996 | GM-BMDC cultures |
| GSE287241 | Mus musculus | Jan 15 2026 | PRJNA1211677 | bone marrow |
| GSE319142 | Homo sapiens | Feb 16 2026 | PRJNA1422172 | Stool |
| GSE302477 | Mus musculus | Feb 06 2026 | PRJNA1291060 | Epidermis |
| GSE311057 | Danio rerio | Jan 31 2026 | PRJNA1368608 | retina |
| GSE303374 | Mus musculus | Jan 01 2026 | PRJNA1294960 | |
| GSE173233 | Mus musculus | Aug 10 2021 | PRJNA724735 | Retina |
| GSE306674 | Penaeus vannamei | Feb 18 2026 | PRJNA1311611 | Hepatopancreas |
| GSE317191 | Homo sapiens | Jan 24 2026 | PRJNA1406245 | Placental DNA |
| GSE228269 | Homo sapiens | Aug 30 2023 | PRJNA949174 | crovascular endothelial cells in cell culture |
| GSE290241 | Drosophila melanogaster | Feb 18 2026 | PRJNA1226640 | adult ovary |
| GSE261412 | Mus musculus | Feb 17 2026 | PRJNA1086895 | |
| GSE299009 | Homo sapiens | Feb 25 2026 | PRJNA1272108 | g alveolar basal epithelium |
| GSE222385 | Homo sapiens | Jan 12 2023 | PRJNA921855 | retinal epithelium |
| GSE310529 | Homo sapiens | Feb 25 2026 | PRJNA1366474 | stomach |
| GSE309787 | Mus musculus | Feb 12 2026 | PRJNA1363175 | |
| GSE310036 | Homo sapiens | Jan 30 2026 | PRJNA1365034 | |
| GSE217573 | Mus musculus | Nov 13 2022 | PRJNA899563 | |
| GSE292822 | Mus musculus | Jan 07 2026 | PRJNA1241306 | Frontal cortex |
| GSE249756 | Danio rerio | Jun 21 2024 | PRJNA1050288 | eye |
| GSE314202 | Mus musculus | Feb 28 2026 | PRJNA1390244 | Mammary Tumour |

| | | | | |
|---|---|---|---|---|
| GSE173079 | Mus musculus | Nov 25 2024 | Murine retinal endothelial cells | |
| GSE283656 | Homo sapiens | Jan 14 2026 | PRJNA1194992 | Breast Tissue |
| GSE161645 | acchus;Macaca fascicularis;Mus | Oct 19 2021 | PRJNA679024 | Retina |
| GSE191077 | Mus musculus | Dec 21 2021 | PRJNA789577 | etinal pigment epithelium |
| GSE299243 | Homo sapiens | Jan 24 2026 | PRJNA1273157 | SC-derived gastric organoids |
| GSE189954 | Mus musculus | Dec 21 2021 | PRJNA785265 | kidney |
| GSE295985 | Homo sapiens | Feb 26 2026 | PRJNA1257084 | l Colorectal Cancer organoids |
| GSE293843 | Homo sapiens | Jan 26 2026 | PRJNA1246577 | |
| GSE311060 | odopseudomonas palustris CGA( | Jan 01 2026 | PRJNA1368614 | |
| GSE301917 | Bos taurus;Homo sapiens | Feb 06 2026 | PRJNA1287725 | milk |
| GSE277267 | Homo sapiens | Jan 12 2026 | PRJNA1161621 | |
| GSE277139 | Capra hircus | Feb 11 2026 | PRJNA1160859 | TMJ |
| GSE287576 | Homo sapiens | Jan 01 2026 | PRJNA1213770 | brachialradialis muscle |
| GSE268251 | Mus musculus | Jan 21 2026 | PRJNA1115794 | Heart |
| GSE224648 | Mus musculus | Jun 08 2023 | PRJNA932003 | eye organoid |
| GSE277426 | Mus musculus | Jan 13 2026 | PRJNA1162149 | |
| GSE287069 | Mycobacterium tuberculosis | Jan 31 2026 | PRJNA1211083 | Bacterial Cell lysate |
| GSE220154 | Homo sapiens | Feb 27 2023 | PRJNA908874 | retinal organoid |
| GSE302201 | Mus musculus | Jan 01 2026 | PRJNA1289196 | Tumor |
| GSE271282 | Mus musculus | Jan 13 2026 | PRJNA1130855 | |
| GSE310640 | Homo sapiens | Jan 28 2026 | PRJNA1366637 | Skeletal muscle |
| GSE318614 | Siniperca chuatsi | Feb 11 2026 | PRJNA1419312 | spleen |
| GSE298463 | Homo sapiens | Jan 26 2026 | PRJNA1269495 | kidney tumor |
| GSE276158 | Mus musculus | Jan 13 2026 | PRJNA1155202 | lung |
| GSE220636 | Mus | Jan 20 2026 | PRJNA910657 | prefrontal cortex |
| GSE190657 | Mus musculus | Dec 31 2023 | PRJNA787928 | |
| GSE303748 | Homo sapiens | Feb 26 2026 | PRJNA1297602 | Breast |
| GSE302554 | synthetic construct | Jan 28 2026 | PRJNA1291184 | |
| GSE319120 | Danio rerio | Feb 10 2026 | PRJNA1422136 | Whole Embryo |
| GSE315668 | Homo sapiens | Jan 13 2026 | PRJNA1398655 | PBMC |
| GSE314474 | Petromyzon marinus | Feb 25 2026 | PRJNA1391375 | Embryos |
| GSE268504 | Mus musculus | Apr 23 2025 | PRJNA1117473 | organ Of Corti |
| GSE249642 | Homo sapiens | Dec 09 2024 | PRJNA1049872 | |
| GSE286242 | Drosophila melanogaster | Jan 08 2026 | PRJNA1208461 | whole prepupae |
| GSE180875 | Mus musculus;Sus scrofa | Nov 11 2021 | PRJNA749804 | retina |
| GSE246942 | Homo sapiens | Sep 30 2025 | PRJNA1035135 | |
| GSE311309 | Mus musculus | Feb 18 2026 | PRJNA1368913 | kidney |
| GSE201547 | Mus musculus | Feb 26 2024 | PRJNA832213 | |
| GSE314609 | Mus musculus | Feb 06 2026 | PRJNA1392210 | Hippocampus |
| GSE268266 | Homo sapiens | Feb 23 2026 | PRJNA1116061 | |
| GSE259426 | Mus musculus | Feb 06 2026 | PRJNA1081464 | Mouse embryos |
| GSE206100 | Mus musculus | Jun 24 2022 | PRJNA849198 | Retina |
| GSE227524 | Tupaia belangeri | Nov 27 2024 | PRJNA945453 | Retina |
| GSE155411 | Mus musculus | Nov 18 2020 | PRJNA649688 | whole retina |
| GSE278002 | Homo sapiens | Jan 05 2026 | PRJNA1165206 | |
| GSE159359 | Homo sapiens | Sep 05 2021 | PRJNA668653 | Retina periphery |
| GSE296539 | Homo sapiens | Jan 28 2026 | PRJNA1259940 | |
| GSE300371 | Mus musculus | Jan 12 2026 | PRJNA1280159 | Bone Marrow |
| GSE316251 | Homo sapiens | Jan 25 2026 | PRJNA1401883 | plasma cell-free DNA |
| GSE113686 | Rattus norvegicus | Mar 31 2020 | PRJNA453611 | Retina |
| GSE308158 | Homo sapiens | Feb 11 2026 | PRJNA1329048 | |
| GSE312730 | Homo sapiens | Jan 30 2026 | PRJNA1375789 | derived engineered heart tissue |
| GSE271037 | Mus musculus | Jan 13 2026 | PRJNA1129461 | spleen |
| GSE262108 | Mus musculus | Jan 27 2026 | | Bone marrow |
| GSE315864 | Mus musculus | Jan 08 2026 | PRJNA1399452 | testis |
| GSE308447 | Mus musculus | Feb 27 2026 | PRJNA1330608 | Kidney |
| GSE222746 | Mus musculus | Jan 01 2026 | PRJNA923599 | |
| GSE316533 | Homo sapiens | Jan 21 2026 | PRJNA1403275 | |
| GSE303973 | Mus musculus | Jan 07 2026 | PRJNA1298334 | heart |
| GSE269923 | Mus musculus | Jun 14 2024 | PRJNA1124124 | Retina |
| GSE154447 | Mus musculus | Jul 01 2024 | PRJNA646306 | |
| GSE312707 | Danio rerio | Jan 01 2026 | PRJNA1375755 | Gill |
| GSE285353 | Toxoplasma gondii | Jan 01 2026 | PRJNA1203215 | |

| | | | | |
|---|---|---|---|---|
| GSE317207 | Homo sapiens | Feb 12 2026 | | liver |
| GSE281632 | Mus musculus | Feb 11 2026 | PRJNA1184847 | |
| GSE305646 | Mus musculus | Feb 23 2026 | PRJNA1307404 | heart |
| GSE289666 | Homo sapiens | Feb 26 2026 | PRJNA1223914 | |
| GSE307583 | Mus musculus | Jan 30 2026 | PRJNA1322337 | |
| GSE149388 | Mus musculus | Nov 29 2025 | PRJNA628601 | retina |
| GSE302912 | Mus musculus | Jan 01 2026 | PRJNA1292809 | testis |
| GSE303780 | Mus musculus | Feb 18 2026 | PRJNA1297674 | Ventricular heart |
| GSE285431 | Mus musculus | Feb 25 2026 | PRJNA1203625 | |
| GSE289379 | Homo sapiens | Jan 21 2026 | PRJNA1222822 | sputum |
| GSE300989 | Homo sapiens | Feb 20 2026 | PRJNA1282958 | Blood |
| GSE292453 | Homo sapiens | Feb 25 2026 | | Bone Marrow |
| GSE316427 | Homo sapiens | Feb 19 2026 | PRJNA1402879 | |
| GSE287824 | Rattus norvegicus | Jan 20 2026 | PRJNA1214770 | hippocampal |
| GSE308553 | Homo sapiens | Jan 14 2026 | PRJNA1330871 | orbital tissue |
| GSE317324 | Mus musculus | Jan 23 2026 | PRJNA1406741 | liver |
| GSE316345 | Mus musculus | Feb 15 2026 | PRJNA1402347 | |
| GSE308231 | Homo sapiens | Jan 14 2026 | | stomach |
| GSE245783 | Homo sapiens | Jul 24 2024 | PRJNA1029964 | |
| GSE279615 | Homo sapiens | Feb 25 2026 | | human embryonic kidney |
| GSE206014 | Mus musculus | May 31 2023 | PRJNA848799 | retina |
| GSE312171 | Mus musculus | Jan 12 2026 | PRJNA1372593 | Colon |
| GSE239573 | Mus musculus | Mar 12 2024 | PRJNA999717 | retina |
| GSE316372 | Mus musculus | Feb 05 2026 | PRJNA1402385 | |
| GSE313672 | Mus musculus | Jan 14 2026 | PRJNA1380506 | Lung |
| GSE310958 | Homo sapiens | Feb 20 2026 | PRJNA1367603 | |
| GSE277943 | Mus musculus | Jan 05 2026 | PRJNA1164783 | testis |
| GSE306878 | Mus musculus | Jan 21 2026 | PRJNA1312842 | mouse uterus |
| GSE262534 | Drosophila melanogaster | Jan 27 2026 | PRJNA1092153 | |
| GSE309267 | Homo sapiens | Feb 18 2026 | PRJNA1334992 | primary neuroblastoma |
| GSE270617 | Mus musculus | Jan 12 2026 | PRJNA1127574 | spleen |
| GSE289010 | Mus musculus | Feb 28 2026 | PRJNA1220602 | |
| GSE308564 | Mus musculus | Feb 20 2026 | PRJNA1330890 | Pancreatic beta cells |
| GSE296239 | Homo sapiens | Jan 12 2026 | PRJNA1258261 | breast carcinoma |
| GSE123758 | Mus musculus | Dec 11 2021 | PRJNA509760 | |
| GSE310188 | Homo sapiens | Jan 01 2026 | PRJNA1365252 | PBMC |
| GSE245197 | Mus musculus | Dec 31 2023 | PRJNA1027340 | P7, RETINA |
| GSE317252 | Mus musculus | Feb 26 2026 | PRJNA1406628 | lung |
| GSE215220 | Homo sapiens | Jan 11 2026 | PRJNA889380 | brain |
| GSE292247 | Homo sapiens | Jan 01 2026 | PRJNA1237741 | |
| GSE314867 | Mus musculus | Jun 18 2026 | PRJNA1393018 | Retina |
| GSE290813 | Staphylococcus aureus | Feb 28 2026 | PRJNA1230166 | |
| GSE290824 | Mus musculus | Feb 27 2026 | PRJNA1230181 | Retina |
| GSE242518 | Akkermansia muciniphila | Jan 01 2026 | | |
| GSE197510 | Mus musculus | Nov 15 2022 | PRJNA810722 | |
| GSE312996 | Pseudomonas aeruginosa | Jan 25 2026 | PRJNA1378078 | |
| GSE281650 | Homo sapiens | Sep 17 2025 | PRJNA1185160 | |
| GSE284350 | Homo sapiens | Apr 02 2026 | PRJNA1198864 | |
| GSE189619 | Mustela putorius furo | Dec 31 2021 | PRJNA783897 | Lung tissues |
| GSE300159 | Homo sapiens;Mus musculus | Feb 20 2026 | PRJNA1279150 | |
| GSE267142 | Mus musculus | Jan 18 2026 | PRJNA1109846 | |
| GSE279529 | Mus musculus | Jan 21 2026 | PRJNA1173196 | Lung |
| GSE317424 | Homo sapiens | Jan 31 2026 | PRJNA1412587 | primary prostate cancer |
| GSE289238 | Homo sapiens | Feb 10 2026 | PRJNA1221953 | Colon |
| GSE304004 | Mus musculus | Jan 01 2026 | PRJNA1298739 | connective tissue |
| GSE180210 | Homo sapiens | Jul 16 2021 | PRJNA747028 | |
| GSE316327 | Homo sapiens | Jan 20 2026 | PRJNA1402322 | mammary gland |
| GSE300207 | Mus musculus | Feb 14 2026 | PRJNA1279251 | brain |
| GSE290096 | Micropterus salmoides | Feb 20 2026 | PRJNA1226189 | gonad |
| GSE282859 | Homo sapiens | Dec 23 2024 | PRJNA1190897 | etinal pigment epithelium |
| GSE118128 | Octopus bimaculoides | Jan 31 2021 | PRJNA484556 | Salivary_Gland-Anterior |
| GSE129964 | Homo sapiens | May 13 2020 | PRJNA533299 | |
| GSE287871 | Homo sapiens | Jan 17 2026 | PRJNA1214836 | |

| | | | | |
|---|---|---|---|---|
| GSE143312 | Danio rerio | May 06 2020 | PRJNA599983 | |
| GSE305052 | piens;Mus musculus;Persicaria | Jan 16 2026 | PRJNA1304375 | |
| GSE293305 | Homo sapiens | May 12 2026 | PRJNA1243984 | |
| GSE288771 | Homo sapiens | Jan 16 2026 | PRJNA1219225 | |
| GSE317371 | Homo sapiens | Jan 27 2026 | PRJNA1411834 | |
| GSE312671 | Homo sapiens | Jan 21 2026 | PRJNA1375687 | |
| GSE303194 | Mus musculus | Jan 21 2026 | PRJNA1294315 | condylar cartilage |
| GSE308019 | Homo sapiens | Jan 14 2026 | PRJNA1328599 | agus, squamous cell carcinoma |
| GSE270117 | Homo sapiens | Jan 16 2026 | PRJNA1125021 | PDA Cell Lines |
| GSE223379 | Homo sapiens | Jan 30 2023 | PRJNA925899 | Retinal organoid |
| GSE312882 | Mus musculus | Jan 23 2026 | PRJNA1377553 | |
| GSE305730 | Homo sapiens | Feb 11 2026 | PRJNA1307511 | |
| GSE155550 | Mus musculus | Aug 03 2020 | PRJNA650285 | retina |
| GSE165691 | Mus musculus | Jan 29 2021 | PRJNA695467 | |
| GSE309567 | Drosophila melanogaster | Feb 09 2026 | PRJNA1336255 | ptic lobes from adult brain |
| GSE153818 | Mus musculus | Sep 29 2020 | PRJNA644203 | |
| GSE136929 | Homo sapiens | Sep 05 2020 | PRJNA564075 | |
| GSE273807 | Pantherophis guttatus | Mar 03 2025 | PRJNA1143394 | Dorsal skin |
| GSE314492 | Caenorhabditis elegans | Jan 05 2026 | PRJNA1391406 | whole animal |
| GSE241192 | Rattus norvegicus | Aug 31 2024 | PRJNA1006718 | eye |
| GSE289357 | Homo sapiens | Jan 27 2026 | PRJNA1222773 | |
| GSE199840 | Homo sapiens | Apr 28 2022 | PRJNA821677 | Retina explant |
| GSE210526 | Gallus gallus | Jul 26 2024 | PRJNA866108 | RPE |
| GSE300368 | Plasmodium falciparum | Feb 16 2026 | PRJNA1280155 | |
| GSE317054 | Mus musculus | Jan 30 2026 | PRJNA1405708 | heart |
| GSE307923 | Homo sapiens | Mar 10 2026 | PRJNA1327834 | induced retinal organoid |
| GSE308141 | Bos taurus;Homo sapiens | Jan 06 2026 | PRJNA1329012 | Endometrium |
| GSE297541 | Mus musculus | Jan 01 2026 | PRJNA1265164 | |
| GSE146255 | Homo sapiens | Feb 16 2021 | PRJNA609898 | |
| GSE291809 | Mus musculus | Feb 21 2026 | PRJNA1235272 | Whole gonad |
| GSE203653 | Homo sapiens;Mus musculus | Feb 14 2023 | PRJNA841679 | |
| GSE288301 | Mus musculus | Feb 16 2026 | PRJNA1216893 | |
| GSE211570 | Homo sapiens;Mus musculus | Jul 11 2023 | PRJNA870924 | embryonic stem cell |
| GSE318801 | Mus musculus | Feb 11 2026 | PRJNA1420643 | Bone-marrow derived |
| GSE310855 | Homo sapiens | Jan 30 2026 | PRJNA1367176 | |
| GSE296999 | Mus musculus | Jan 13 2026 | PRJNA1262340 | colon |
| GSE317063 | Rattus norvegicus | Feb 18 2026 | PRJNA1405721 | ileum |
| GSE263147 | Mus musculus | Feb 11 2026 | PRJNA1095979 | Mus musculus |
| GSE309408 | is;Columba livia;Gallus gallus;Ta | Nov 11 2025 | PRJNA1336006 | Eye |
| GSE297214 | Homo sapiens | Jan 12 2026 | PRJNA1263019 | |
| GSE316760 | Homo sapiens | Feb 18 2026 | PRJNA1404153 | Cutaneous melanoma |
| GSE234435 | Danio rerio | Nov 10 2023 | PRJNA981359 | Whole retina |
| GSE300990 | Mus musculus | Feb 16 2026 | PRJNA1282965 | |
| GSE285776 | Mus musculus | Jan 12 2026 | PRJNA1206069 | spleen |
| GSE235577 | Homo sapiens | Feb 15 2024 | PRJNA986496 | organoid |
| GSE296356 | Mus musculus | Feb 10 2026 | PRJNA1258980 | |
| GSE310976 | Homo sapiens | Feb 20 2026 | PRJNA1367648 | |
| GSE223335 | Homo sapiens | Jan 10 2026 | PRJNA925834 | |
| GSE215132 | Mus musculus | Nov 10 2022 | PRJNA888974 | dorsal root ganglion |
| GSE279313 | Macaca mulatta | Feb 06 2026 | PRJNA1171838 | lymph node |
| GSE297628 | Rattus norvegicus | Jan 11 2026 | PRJNA1265468 | mammary tumor |
| GSE301548 | Drosophila melanogaster | Jan 19 2026 | PRJNA1285423 | Brain |
| GSE297300 | Mus musculus | Feb 14 2026 | PRJNA1263481 | Ovary |
| GSE301057 | Homo sapiens | Feb 25 2026 | PRJNA1283169 | n vitro differentiated cells |
| GSE278138 | Homo sapiens | Jan 26 2026 | PRJNA1165763 | SNSCC tumor |
| GSE198093 | Homo sapiens | Mar 14 2022 | PRJNA813591 | |
| GSE313416 | Homo sapiens | Mar 11 2026 | PRJNA1379470 | |
| GSE300529 | Homo sapiens | Feb 06 2026 | PRJNA1281113 | |
| GSE317615 | Homo sapiens | Jan 31 2026 | PRJNA1414554 | |
| GSE287484 | Mus musculus | Jan 25 2025 | PRJNA1213079 | ESCs |
| GSE312315 | Homo sapiens | Jan 16 2026 | PRJNA1373015 | PDA Cell Lines |
| GSE316548 | Homo sapiens | Feb 09 2026 | PRJNA1403297 | |
| GSE285600 | Brassica rapa subsp. chinensis | Jan 01 2026 | PRJNA1204947 | Leaf |

| | | | | |
|---|---|---|---|---|
| GSE306920 | Mus musculus | Feb 12 2026 | PRJNA1313917 | ovary |
| GSE310800 | Mus musculus | Feb 25 2026 | PRJNA1367090 | embryonic limb bud tissue |
| GSE284103 | Homo sapiens | Jan 12 2026 | PRJNA1197359 | |
| GSE304898 | Mus musculus | Feb 19 2026 | | spleen |
| GSE279265 | Homo sapiens | Feb 25 2026 | PRJNA1171760 | |
| GSE317036 | Mus musculus | Jan 22 2026 | PRJNA1405696 | Brown adipose tissue |
| GSE283935 | Mus musculus | Jan 08 2026 | PRJNA1196354 | Colon tumor |
| GSE294687 | Homo sapiens | Dec 03 2025 | PRJNA1251089 | |
| GSE297241 | Homo sapiens | Feb 11 2026 | PRJNA1263383 | |
| GSE316735 | Homo sapiens | Jan 20 2026 | PRJNA1404102 | Breast Tissue |
| GSE266412 | Homo sapiens | Sep 30 2025 | PRJNA1107120 | |
| GSE138148 | Camellia sinensis | Dec 31 2020 | PRJNA574928 | root |
| GSE246702 | Mus musculus | Jan 01 2025 | PRJNA1034145 | Eyes |
| GSE317990 | Mus musculus | Feb 27 2026 | | thymus |
| GSE312501 | Homo sapiens | Jan 12 2026 | PRJNA1374071 | |
| GSE287290 | Mus musculus | Jan 12 2026 | PRJNA1212041 | pancreas |
| GSE197509 | Mus musculus | Nov 15 2022 | PRJNA810720 | retina |
| GSE281649 | Homo sapiens | Sep 17 2025 | PRJNA1185157 | |
| GSE292269 | Mus musculus | Jan 06 2026 | PRJNA1237794 | Retina |
| GSE285155 | Mus musculus | Jan 26 2026 | PRJNA1201316 | |
| GSE311856 | Homo sapiens | Jan 26 2026 | PRJNA1371766 | Adrenal mass |
| GSE318863 | Homo sapiens | Feb 11 2026 | PRJNA1420732 | Umbilical cord blood |
| GSE159168 | Rattus norvegicus | Mar 24 2021 | PRJNA668010 | retina |
| GSE180157 | Homo sapiens | Nov 12 2021 | PRJNA746785 | |
| GSE317763 | Mus musculus | May 06 2026 | PRJNA1415193 | RPE/choroid |
| GSE299486 | Homo sapiens | Feb 11 2026 | PRJNA1274715 | |
| GSE223357 | Homo sapiens | Jan 20 2026 | PRJNA925872 | |
| GSE126437 | Homo sapiens | Feb 14 2026 | PRJNA521892 | lymphoblastoid cell line |
| GSE291293 | Homo sapiens | Jan 05 2026 | PRJNA1232731 | Primary Human HSPC |
| GSE270344 | Canis lupus familiaris | Feb 19 2026 | PRJNA1126236 | mammary |
| GSE318940 | Mus musculus | Feb 16 2026 | PRJNA1420855 | Bone Marrow |
| GSE281517 | Homo sapiens | Jan 05 2026 | PRJNA1184173 | |
| GSE240085 | Pomacea canaliculata | Dec 31 2024 | PRJNA1002264 | eye stalk |
| GSE285714 | Mus musculus | Feb 16 2026 | PRJNA1205695 | inguinal lymph nodes |
| GSE273782 | Mus musculus | Jan 13 2026 | PRJNA1143051 | Brain |
| GSE299592 | Homo sapiens | Jan 09 2026 | PRJNA1275032 | Cell line |
| GSE211119 | Mus musculus | Aug 15 2022 | PRJNA869031 | retina |
| GSE226102 | Arabidopsis thaliana | Feb 28 2026 | PRJNA938505 | Siliques |
| GSE298291 | Mus musculus | Jan 21 2026 | PRJNA1268897 | thymus |
| GSE307691 | Homo sapiens | Jan 05 2026 | PRJNA1322531 | connective tissue |
| GSE239775 | Homo sapiens | Feb 23 2026 | PRJNA1000950 | Whole blood |
| GSE310923 | Mus musculus | Jan 15 2026 | PRJNA1367541 | PBMC |
| GSE302391 | Homo sapiens | Sep 17 2025 | PRJNA1290313 | |
| GSE318977 | Mus musculus | Feb 15 2026 | PRJNA1420914 | Peritoneal cavity |
| GSE211406 | Mus musculus | May 01 2026 | PRJNA870115 | |
| GSE305554 | Mus musculus | Feb 24 2026 | PRJNA1306497 | |
| GSE247115 | Homo sapiens | Apr 23 2025 | PRJNA1036239 | Gastric |
| GSE301035 | Homo sapiens | Feb 09 2026 | PRJNA1283113 | |
| GSE318784 | Mus musculus | Feb 18 2026 | PRJNA1420539 | |
| GSE300736 | Mus musculus | Feb 11 2026 | PRJNA1281997 | tumor |
| GSE279659 | Mus musculus | Jan 21 2026 | PRJNA1173857 | een, then expanded and transduced |
| GSE263031 | Homo sapiens | Feb 15 2026 | PRJNA1095479 | Cell line |
| GSE314751 | Clostridium perfringens | Jan 20 2026 | PRJNA1392546 | |
| GSE183165 | Mus musculus | Sep 03 2021 | PRJNA759424 | |
| GSE225024 | Mus musculus | Jan 31 2026 | PRJNA934588 | |
| GSE291026 | Homo sapiens | Jan 14 2026 | PRJNA1231424 | hematopoietic |
| GSE318506 | Homo sapiens | Feb 27 2026 | PRJNA1419128 | Pancreatic Islet |
| GSE287106 | Penicillium oxalicum | Jan 31 2026 | PRJNA1211132 | Mycelia |
| GSE310552 | Mus musculus | Feb 25 2026 | | |
| GSE212520 | Mus musculus | Aug 18 2024 | PRJNA875896 | Retina |
| GSE297682 | Homo sapiens | Jan 14 2026 | PRJNA1265681 | Breast |
| GSE286167 | Mus musculus | Jan 01 2026 | PRJNA1208039 | Colon |
| GSE315707 | Hormonema carpetanum | Jan 13 2026 | PRJNA1398825 | mycelium |

| | | | | |
|---|---|---|---|---|
| GSE314995 | Homo sapiens | Jan 31 2026 | | PBMC |
| GSE287653 | Mus musculus | Jan 21 2026 | PRJNA1214198 | Small intestine |
| GSE290712 | Caenorhabditis elegans | Feb 27 2026 | PRJNA1229610 | worm |
| GSE213214 | Homo sapiens | Feb 23 2026 | PRJNA879741 | |
| GSE308689 | Mus musculus | Jan 28 2026 | PRJNA1332540 | lung lysate |
| GSE287770 | Homo sapiens;Mus musculus | Feb 16 2026 | PRJNA1214674 | NA |
| GSE292296 | Mus musculus | Jan 22 2026 | PRJNA1237841 | Spleen |
| GSE307507 | Canis lupus familiaris | Jan 27 2026 | PRJNA1321892 | Saliva |
| GSE297796 | Mus musculus | Jan 06 2026 | PRJNA1266193 | |
| GSE240446 | Homo sapiens | Aug 09 2024 | PRJNA1003811 | peripheral blood |
| GSE282512 | Homo sapiens | Jan 01 2026 | PRJNA1189068 | Blood plasma |
| GSE281845 | Homo sapiens | Sep 17 2025 | PRJNA1185802 | |
| GSE126796 | Rattus norvegicus | Jan 30 2022 | PRJNA523340 | liver |
| GSE255831 | Homo sapiens | Feb 18 2026 | PRJNA1076642 | |
| GSE315146 | Homo sapiens | Jan 12 2026 | PRJNA1395057 | |
| GSE237213 | Tupaia belangeri chinensis | Oct 01 2023 | PRJNA994175 | Retina |
| GSE242783 | Homo sapiens | Jun 30 2025 | PRJNA1014717 | |
| GSE310159 | Drosophila melanogaster | Jan 07 2026 | PRJNA1365221 | larval guts |
| GSE298665 | Mus musculus | Dec 17 2025 | PRJNA1270835 | etinal pigment epithelium (RPE) |
| GSE208743 | Homo sapiens | Jan 26 2026 | PRJNA860941 | |
| GSE299479 | Rattus norvegicus | Feb 26 2026 | PRJNA1274705 | |
| GSE315459 | Mus musculus | Feb 19 2026 | PRJNA1397184 | istal part of small intestine |
| GSE201985 | Mus musculus | May 02 2025 | PRJNA833931 | brain microglial cell |
| GSE133247 | Homo sapiens | Mar 01 2020 | PRJNA550448 | |
| GSE273669 | Homo sapiens | Feb 17 2026 | PRJNA1142590 | s:Melanoma-Full Thickness-A375 |
| GSE322475 | Mus musculus | Feb 28 2026 | PRJNA1429411 | retina explant |
| GSE319326 | Homo sapiens | Feb 13 2026 | PRJNA1422838 | |
| GSE301394 | Mus musculus | Feb 18 2026 | PRJNA1284922 | liver |
| GSE290319 | Homo sapiens | Jan 08 2026 | PRJNA1227478 | |
| GSE318425 | Mus musculus | Feb 28 2026 | | subcutaneous tumor |
| GSE273414 | Mus musculus | Jan 19 2026 | PRJNA1141570 | |
| GSE279785 | Mus musculus | Feb 26 2026 | PRJNA1174684 | embryo |
| GSE318347 | Homo sapiens | Feb 05 2026 | PRJNA1418609 | Nasal epithelium |
| GSE303672 | Homo sapiens | Jan 27 2026 | PRJNA1297058 | |
| GSE264669 | Homo sapiens | Jan 01 2026 | PRJNA1103775 | Ascites Fluid Cells |
| GSE274390 | Xenopus laevis | Jan 01 2026 | PRJNA1146328 | animal cap |
| GSE288097 | Homo sapiens | Mar 01 2025 | PRJNA1216190 | Retina |
| GSE209654 | Mus musculus | Apr 14 2023 | PRJNA862060 | Retina |
| GSE301939 | Mus musculus | Feb 24 2026 | PRJNA1287755 | |
| GSE318719 | Mus musculus | Feb 11 2026 | PRJNA1419831 | kidney |
| GSE316394 | Mus musculus | Feb 15 2026 | PRJNA1402440 | 5 Medial Ganglionic Eminence |
| GSE161416 | Homo sapiens | Mar 26 2023 | PRJNA678246 | |
| GSE311551 | Mus musculus | Mar 19 2026 | PRJNA1369852 | retina |
| GSE228270 | Chlorocebus sabaeus | Aug 30 2023 | PRJNA949176 | retina/choroid |
| GSE293381 | Mus musculus | May 09 2025 | PRJNA1244320 | Retina |
| GSE143396 | Mus musculus | Jan 10 2020 | PRJNA600281 | etinal pigment epithelium |
| GSE308582 | Homo sapiens | Feb 18 2026 | PRJNA1331621 | PBMC |
| GSE315518 | Homo sapiens | Jan 08 2026 | PRJNA1398199 | Liver |
| GSE315996 | Homo sapiens | Feb 26 2026 | PRJNA1400082 | |
| GSE285789 | Homo sapiens | Jan 08 2026 | PRJNA1206090 | Peripheral blood |
| GSE310018 | Mus musculus | Feb 12 2026 | PRJNA1364991 | Liver |
| GSE240018 | Mus musculus | Aug 05 2023 | PRJNA1001927 | Retina |
| GSE312831 | Mus musculus | Jan 16 2026 | PRJNA1377483 | lung |
| GSE302146 | Homo sapiens | Jan 09 2026 | PRJNA1289067 | |
| GSE304911 | Rattus norvegicus | Jan 12 2026 | | Mammary gland |
| GSE290012 | Danio rerio | Feb 19 2026 | | head |
| GSE236333 | Rattus norvegicus | Jul 05 2023 | PRJNA990646 | Retina |
| GSE315765 | Mus musculus | Jan 07 2026 | | |
| GSE271977 | Sus scrofa | Jan 08 2026 | PRJNA1134234 | |
| GSE295353 | Mus musculus | Jan 26 2026 | PRJNA1254251 | pancreatic islet |
| GSE334500 | Homo sapiens | Jun 09 2026 | PRJNA1475359 | C-derived cardiac organoid |
| GSE244627 | Mus musculus | Jul 03 2024 | PRJNA1023918 | PBMC |
| GSE310424 | Homo sapiens | Feb 23 2026 | PRJNA1365924 | |

| | | | | |
|---|---|---|---|---|
| GSE287070 | Carya illinoinensis;Pinus armandi | Jan 05 2026 | PRJNA1211086 | Leaf |
| GSE318470 | Rattus norvegicus | Feb 18 2026 | PRJNA1418795 | Heart |
| GSE315124 | Homo sapiens | Jan 01 2026 | PRJNA1395017 | ovary |
| GSE316280 | Mus musculus | Jan 18 2026 | PRJNA1402153 | retina |
| GSE225630 | Mus musculus | Feb 20 2026 | PRJNA936844 | lymph node |
| GSE288812 | Citrullus lanatus | Feb 09 2026 | | leaves |
| GSE310659 | Mus musculus | Dec 01 2025 | PRJNA1366829 | eye cup |
| GSE317583 | Homo sapiens | Jan 31 2026 | PRJNA1414505 | |
| GSE294332 | Mus musculus | Feb 20 2026 | PRJNA1249036 | dorsal hippocampus |
| GSE315646 | Homo sapiens | Jan 06 2026 | PRJNA1398384 | Nasopharynx |
| GSE299651 | Homo sapiens | Jan 12 2026 | PRJNA1275837 | |
| GSE318325 | Homo sapiens | Feb 07 2026 | PRJNA1418554 | |
| GSE183746 | Drosophila melanogaster | Dec 31 2021 | PRJNA761801 | Retina |
| GSE327886 | Mus musculus | Apr 17 2026 | PRJNA1452944 | brain |
| GSE232746 | Mus musculus | Sep 30 2024 | PRJNA973685 | retina |
| GSE195895 | Mus musculus | Jul 20 2022 | PRJNA802567 | Retina |
| GSE310413 | Homo sapiens | Feb 23 2026 | PRJNA1365910 | |
| GSE313944 | Mus musculus | Jan 01 2026 | PRJNA1387406 | |
| GSE213362 | Mus musculus | Jan 31 2023 | PRJNA880599 | retina |
| GSE224379 | Gallus gallus | Feb 07 2023 | PRJNA930741 | Fissure margin cells |
| GSE319344 | Danio rerio | Feb 16 2026 | PRJNA1423049 | meninges |
| GSE291478 | Danio rerio | Feb 11 2026 | PRJNA1233852 | Fin |
| GSE197463 | Mus musculus | Feb 28 2022 | PRJNA810437 | gastrocnemius |
| GSE244923 | Mus musculus | Dec 30 2023 | PRJNA1026146 | P5, RETINAL |
| GSE301729 | Homo sapiens | Jan 20 2026 | | bone marrow |
| GSE238008 | Mus musculus | Oct 30 2023 | PRJNA997367 | |
| GSE192601 | Mus musculus | Apr 27 2022 | PRJNA792269 | retina |
| GSE315942 | Homo sapiens | Jan 16 2026 | PRJNA1399996 | |
| GSE284069 | Mus musculus | Jan 21 2026 | PRJNA1196943 | |
| GSE224791 | Homo sapiens | Feb 11 2026 | PRJNA932715 | |
| GSE305993 | Homo sapiens | Feb 18 2026 | PRJNA1308854 | Kidney |
| GSE231768 | Mus musculus | Feb 17 2026 | PRJNA967657 | Gastrocnemius |
| GSE259255 | Mus musculus | Feb 10 2025 | PRJNA1080537 | retina at P17 |
| GSE166565 | Homo sapiens | Feb 11 2021 | PRJNA701280 | |
| GSE316422 | Mus musculus | Jan 29 2026 | PRJNA1402793 | heart |
| GSE299955 | Mus musculus | Jan 20 2026 | PRJNA1277708 | telencephalon |
| GSE268388 | Homo sapiens | Jan 16 2026 | PRJNA1117001 | )A Cell Line injected in mice |
| GSE308556 | Mus musculus | Jan 02 2026 | PRJNA1330875 | |
| GSE165832 | Mus musculus | Dec 22 2021 | PRJNA697987 | retina |
| GSE317443 | Homo sapiens | Jan 28 2026 | PRJNA1412926 | blood |
| GSE290960 | Mus musculus | Feb 11 2026 | PRJNA1231087 | Distal Colon |
| GSE202212 | Danio rerio | May 10 2023 | PRJNA835051 | adult retina |
| GSE237292 | Mus musculus | Aug 15 2023 | PRJNA994534 | etinal pigment epithelium |
| GSE163459 | Homo sapiens;Mus musculus | May 02 2022 | PRJNA686183 | |
| GSE318493 | Homo sapiens | Feb 09 2026 | PRJNA1419031 | Peripheral blood |
| GSE310849 | Rattus norvegicus | Feb 24 2026 | PRJNA1367169 | Skeletal Muscle |
| GSE239300 | Mus musculus | Jan 14 2026 | PRJNA998782 | Bone marrow |
| GSE299876 | Homo sapiens | Jan 21 2026 | PRJNA1277212 | e-etinal pigment epithelium |
| GSE296014 | Mus musculus | Feb 19 2026 | PRJNA1257322 | Fibroblasts |
| GSE284628 | Homo sapiens | Sep 16 2025 | PRJNA1199965 | |
| GSE307099 | Homo sapiens | Jan 20 2026 | PRJNA1314260 | |
| GSE308317 | Mus musculus | Jan 04 2026 | PRJNA1330185 | lung |
| GSE104588 | Rattus norvegicus | Oct 01 2021 | PRJNA413139 | Retina |
| GSE184160 | Mus musculus | Sep 22 2021 | PRJNA763483 | |
| GSE317048 | Mus musculus | Jan 30 2026 | PRJNA1405706 | left ventricle heart tissue |
| GSE315692 | Mus musculus | Jan 06 2026 | PRJNA1398805 | Small intestine |
| GSE273495 | Mycosarcoma maydis | Feb 18 2026 | PRJNA1142014 | |
| GSE242057 | Homo sapiens | Dec 31 2024 | PRJNA1011271 | retina |
| GSE310392 | Mus musculus | Jan 01 2026 | PRJNA1365897 | Liver |
| GSE315861 | Homo sapiens | Jan 30 2026 | PRJNA1399454 | whole blood |
| GSE290687 | Mus musculus | Feb 27 2026 | PRJNA1229543 | retina |
| GSE305116 | Homo sapiens | Feb 17 2026 | PRJNA1304461 | d matched normal gastric mucosa tissues |
| GSE313036 | Rattus norvegicus | Feb 19 2026 | PRJNA1378167 | breast cancer cell |

| | | | | |
|---|---|---|---|---|
| GSE300898 | Homo sapiens | Feb 09 2026 | PRJNA1282626 | |
| GSE145390 | Mus musculus | Mar 15 2022 | PRJNA607064 | retina |
| GSE210747 | Mus musculus | Apr 18 2023 | PRJNA867356 | Cortex (P56) |
| GSE294484 | Saccharomyces cerevisiae | Feb 11 2026 | PRJNA1250287 | yeast cells |
| GSE317235 | Homo sapiens | Jan 26 2026 | PRJNA1406324 | glioma |
| GSE225499 | Drosophila melanogaster | May 05 2023 | PRJNA935805 | |
| GSE288774 | Homo sapiens | Jan 14 2026 | PRJNA1219232 | Placenta |
| GSE309200 | Mus musculus | Jan 20 2026 | PRJNA1334314 | sagittal suture |
| GSE316477 | Homo sapiens | Jan 15 2026 | PRJNA1402981 | |
| GSE270670 | Mycobacterium tuberculosis | Jan 15 2026 | PRJNA1127660 | cription of sheared Mtb genomic DNA |
| GSE297103 | Mus musculus | Jan 09 2026 | PRJNA1262524 | Bone |
| GSE202758 | Mus musculus | Jun 08 2023 | PRJNA837223 | eye organoid |
| GSE317416 | Homo sapiens | Jan 31 2026 | PRJNA1412564 | primary prostate cancer |
| GSE114896 | Rattus norvegicus | Feb 02 2021 | PRJNA472991 | Retina |
| GSE238217 | Mus musculus | Jun 06 2024 | PRJNA998385 | Retina |
| GSE252861 | Mus musculus | May 16 2024 | PRJNA1062844 | E14 retina |
| GSE304036 | Homo sapiens | Jan 22 2026 | PRJNA1298820 | Epidermis |
| GSE161497 | Gallus gallus | Nov 16 2020 | PRJNA678523 | Retina |
| GSE316315 | Mus musculus | Jan 20 2026 | PRJNA1402281 | Brain |
| GSE313615 | Homo sapiens | Feb 11 2026 | PRJNA1380401 | |
| GSE281741 | Homo sapiens | Feb 19 2026 | PRJNA1185628 | cultured neural cells |
| GSE263532 | Sus scrofa | Jan 25 2026 | PRJNA1097995 | dorsal root ganglion |
| GSE308256 | Mus musculus | Jan 12 2026 | PRJNA1329310 | |
| GSE274639 | Homo sapiens | Jan 14 2026 | PRJNA1147323 | soft tissue tumors in PDXs |
| GSE290093 | Mus musculus | Feb 20 2026 | PRJNA1226200 | Whole heart tissue slices |
| GSE269694 | Homo sapiens | Jun 11 2025 | | |
| GSE298956 | Mus musculus | Feb 25 2026 | | Midbrain |
| GSE218107 | Danio rerio | Aug 21 2023 | PRJNA902356 | Retina |
| GSE302856 | Mus musculus | Feb 18 2026 | PRJNA1292470 | |
| GSE225961 | Arabidopsis thaliana | Feb 28 2026 | PRJNA938109 | leaf |
| GSE274444 | Homo sapiens | Feb 15 2026 | PRJNA1146435 | |
| GSE312643 | Oryctolagus cuniculus | Jan 01 2026 | PRJNA1375646 | facial nerve |
| GSE308375 | Homo sapiens | Jan 29 2026 | PRJNA1330305 | |
| GSE281655 | Mus musculus | Jan 13 2026 | PRJNA1185174 | bone marrow |
| GSE289601 | Mus musculus | Feb 08 2026 | | bone marrow |
| GSE234154 | Mus musculus | Sep 08 2023 | PRJNA980211 | Liver |
| GSE276851 | Homo sapiens | Jan 15 2025 | PRJNA1159282 | eye |
| GSE206150 | Mus musculus | Jan 04 2023 | PRJNA849417 | retina |
| GSE162145 | Sus scrofa | Apr 20 2021 | | Adductores |
| GSE293576 | Homo sapiens | Feb 28 2026 | PRJNA1245284 | |
| GSE298875 | Mus musculus | Jan 07 2026 | PRJNA1271537 | |
| GSE275264 | Homo sapiens | Feb 20 2026 | PRJNA1150139 | Lung airway epithelium |
| GSE291840 | Mus musculus | Jan 28 2026 | PRJNA1235612 | Peyer's patches |
| GSE219045 | ;Macaca fascicularis;Mus muscu | Dec 12 2022 | PRJNA906635 | Cornea |
| GSE228487 | Mus musculus | Apr 10 2023 | PRJNA950032 | |
| GSE284128 | Gallus gallus | Aug 18 2025 | PRJNA1197409 | retina |
| GSE281891 | Rattus norvegicus | May 31 2025 | | retina |
| GSE285303 | Homo sapiens;Mus musculus | Jan 14 2026 | PRJNA1202190 | Cervical cancer |
| GSE277196 | Mus musculus | Jan 21 2026 | PRJNA1161170 | Ileum |
| GSE309441 | Aspergillus fumigatus | Feb 26 2026 | | mycelium |
| GSE317535 | Homo sapiens | Feb 27 2026 | PRJNA1414429 | |
| GSE319829 | Mus musculus | Feb 18 2026 | PRJNA1425165 | pooled LN and spleen |
| GSE175751 | Homo sapiens | Sep 27 2021 | PRJNA733486 | |
| GSE304884 | Mus musculus | Feb 13 2026 | PRJNA1303170 | Small intestine |
| GSE276866 | Mus musculus | Jan 21 2026 | PRJNA1159293 | Proximal Small Intestine |
| GSE254688 | Phyllostachys violascens | Jan 31 2026 | PRJNA1071523 | culms |
| GSE316925 | Mus musculus | Jan 28 2026 | PRJNA1405545 | Trigeminal ganglion |
| GSE265755 | Schizosaccharomyces pombe | Feb 25 2026 | PRJNA1104156 | |
| GSE298433 | Mus musculus | Jun 19 2026 | PRJNA1269437 | Rod PR |
| GSE319147 | Tabanus nipponicus | Feb 15 2026 | PRJNA1422206 | salivary gland |
| GSE191104 | Mus musculus | May 12 2022 | PRJNA789767 | etinal pigment epithelium |
| GSE318644 | Gallus gallus | Feb 10 2026 | PRJNA1419705 | Optic fissure margin |
| GSE299618 | Homo sapiens | Feb 16 2026 | PRJNA1275676 | |

| | | | | |
|---|---|---|---|---|
| GSE270968 | Mus musculus | Feb 20 2026 | PRJNA1129125 | |
| GSE319170 | Mus musculus | Feb 14 2026 | PRJNA1422245 | |
| GSE253718 | Homo sapiens | Jan 01 2026 | PRJNA1066819 | cancer tumor tissues |
| GSE141139 | Homo sapiens | Jun 11 2020 | PRJNA592189 | |
| GSE298679 | Mus musculus | Feb 17 2026 | PRJNA1270941 | |
| GSE314659 | Homo sapiens | Feb 11 2026 | PRJNA1392419 | |
| GSE315445 | Danio rerio | Jan 02 2026 | PRJNA1397167 | whole embryo |
| GSE148371 | Homo sapiens | May 19 2020 | PRJNA624097 | uman trabecular meshwork |
| GSE311646 | Homo sapiens | Jan 06 2026 | PRJNA1370067 | FP, NoNVP2, Rep1 (EnChAMP-seq) |
| GSE183572 | Mus musculus | Sep 08 2021 | PRJNA761340 | inal Pigment Epithelium cells |
| GSE313583 | Mus musculus | Feb 19 2026 | PRJNA1380343 | Epicardium |
| GSE201356 | Homo sapiens | Dec 10 2022 | PRJNA830973 | Retinal organoid |
| GSE197847 | Homo sapiens | May 10 2023 | PRJNA812505 | retinal organoid |
| GSE235294 | Homo sapiens | Jan 05 2025 | PRJNA985660 | |
| GSE285904 | Homo sapiens | Jan 27 2026 | PRJNA1206920 | |
| GSE277562 | Danio rerio | Jan 01 2026 | PRJNA1162812 | pronephric duct |
| GSE161622 | Mus musculus | Jan 20 2021 | PRJNA678960 | |
| GSE298245 | Mus musculus | Jan 13 2026 | PRJNA1268563 | embryo |
| GSE275690 | Homo sapiens | Feb 20 2026 | PRJNA1152577 | |
| GSE289597 | Mus musculus | Feb 08 2026 | | bone marrow |
| GSE290739 | Mus musculus | Feb 28 2026 | PRJNA1229677 | |
| GSE317694 | Plasmodium falciparum | Feb 19 2026 | PRJNA1414658 | |
| GSE316806 | Homo sapiens | Jan 22 2026 | PRJNA1404256 | liver |
| GSE221559 | Mus musculus | Dec 14 2023 | PRJNA914782 | Liver |
| GSE318278 | Mus musculus | Feb 05 2026 | PRJNA1418533 | |
| GSE291247 | Homo sapiens | Feb 10 2026 | PRJNA1232656 | normal cervical tissue |
| GSE315233 | Homo sapiens | Feb 11 2026 | PRJNA1395382 | Engineered in vitro model |
| GSE174491 | Drosophila melanogaster | Dec 31 2021 | PRJNA730039 | |
| GSE307881 | Mus musculus | Jan 29 2026 | PRJNA1327674 | E18.5 ovary |
| GSE282726 | Homo sapiens;Mus musculus | Feb 21 2026 | PRJNA1190070 | prostate cancer |
| GSE288839 | Mus musculus | Feb 26 2026 | | Testes |
| GSE315372 | Mus musculus | Jan 04 2026 | PRJNA1396608 | bone marrow |
| GSE304497 | Homo sapiens | Jan 14 2026 | PRJNA1301387 | t-mortem fresh frozen brain |
| GSE285872 | Mus musculus | Jan 01 2026 | PRJNA1206687 | Colon |
| GSE302551 | Ovis aries | Jan 28 2026 | PRJNA1291175 | Intervertebral disc |
| GSE287739 | Homo sapiens | Jan 08 2026 | PRJNA1214360 | |
| GSE134872 | Mus musculus | Mar 10 2020 | PRJNA556668 | Rod photoreceptors |
| GSE300867 | Homo sapiens | Feb 26 2026 | PRJNA1282482 | |
| GSE315410 | Mus musculus | Jan 21 2026 | PRJNA1396684 | Liver |
| GSE295667 | mmotragus lervia;Ovis canadens | Feb 13 2026 | PRJNA1255556 | Lung |
| GSE196950 | Mus musculus | Feb 20 2022 | PRJNA808036 | brain |
| GSE277423 | Mus musculus | Jan 13 2026 | PRJNA1162147 | |
| GSE302204 | Homo sapiens | Feb 17 2026 | PRJNA1289201 | |
| GSE315138 | Homo sapiens | Jan 31 2026 | PRJNA1395034 | Duodenal |
| GSE282073 | Arabidopsis thaliana | Feb 06 2026 | PRJNA1186952 | root |
| GSE225333 | Mus musculus | Feb 20 2026 | PRJNA935287 | Cortex |
| GSE261261 | Homo sapiens | Feb 24 2026 | PRJNA1086182 | T cell |
| GSE220633 | Mus | Jan 20 2026 | PRJNA910564 | prefrontal cortex |
| GSE276442 | Homo sapiens | Dec 24 2024 | PRJNA1157377 | |
| GSE141639 | Homo sapiens | Jan 24 2020 | PRJNA594287 | |
| GSE294631 | Xenopus laevis | Feb 18 2026 | PRJNA1250967 | dorsal trunk |
| GSE302079 | Mus musculus | Feb 25 2026 | PRJNA1288480 | |
| GSE318611 | Drosophila melanogaster | Feb 06 2026 | PRJNA1419303 | brain |
| GSE302327 | Homo sapiens | Jan 14 2026 | PRJNA1290081 | Intestine |
| GSE290273 | Homo sapiens | Feb 10 2026 | PRJNA1227120 | |
| GSE281596 | Mus musculus | Feb 11 2026 | PRJNA1184770 | Sp5C, Brain |
| GSE262977 | Homo sapiens | Sep 30 2024 | PRJNA1095371 | retina |
| GSE310995 | Mus musculus | Feb 06 2026 | PRJNA1367910 | fecal |
| GSE277500 | Homo sapiens | Sep 15 2025 | PRJNA1162536 | retina |
| GSE291570 | Homo sapiens | Feb 11 2026 | PRJNA1234169 | Differentiating cells |
| GSE316853 | Rattus norvegicus | Feb 28 2026 | PRJNA1405362 | Liver |
| GSE310004 | Homo sapiens | Feb 24 2026 | PRJNA1364989 | |
| GSE299759 | Homo sapiens | Jan 04 2026 | PRJNA1276455 | |

| | | | | |
|---|---|---|---|---|
| GSE266909 | Chlamydomonas reinhardtii | Jan 26 2026 | PRJNA1108708 | cells |
| GSE310279 | Mus musculus | Jan 01 2026 | PRJNA1365692 | scapular brown adipose tissue |
| GSE315266 | Homo sapiens | Jan 07 2026 | PRJNA1395527 | blood |
| GSE126838 | Mus musculus | Feb 01 2022 | PRJNA523435 | retina |
| GSE294712 | Mus musculus | Feb 18 2026 | PRJNA1251356 | kidney |
| GSE313688 | Homo sapiens | Jan 18 2026 | | Endothelial cells (HUVEC) |
| GSE315251 | Homo sapiens | Jan 30 2026 | PRJNA1395503 | |
| GSE294725 | Drosophila melanogaster | Feb 25 2026 | PRJNA1251508 | brain and eye disc |
| GSE216520 | Mus musculus | Jan 23 2024 | PRJNA894148 | Retina |
| GSE319006 | Homo sapiens | Feb 16 2026 | PRJNA1421165 | Ovarian |
| GSE311052 | Homo sapiens | Feb 24 2026 | PRJNA1368603 | heart |
| GSE153673 | Mus musculus | Oct 23 2020 | PRJNA643589 | |
| GSE318067 | Homo sapiens | Feb 06 2026 | PRJNA1416417 | |
| GSE296382 | Homo sapiens | Jan 20 2026 | PRJNA1259027 | Breast |
| GSE308910 | Homo sapiens | Jan 13 2026 | PRJNA1333374 | Brain |
| GSE306113 | Homo sapiens | Jan 08 2026 | PRJNA1309143 | Prostate |
| GSE161368 | Homo sapiens | Jan 16 2026 | PRJNA677968 | |
| GSE207929 | Mus musculus | Jul 11 2022 | PRJNA857703 | Retinal Tissue |
| GSE197192 | Mus musculus | Feb 25 2022 | PRJNA809347 | |
| GSE314543 | Homo sapiens | Feb 11 2026 | PRJNA1391483 | |
| GSE161104 | Mus musculus | Apr 28 2021 | PRJNA675588 | |
| GSE316686 | Mus musculus | Jan 20 2026 | PRJNA1403584 | |
| GSE298563 | Homo sapiens | Jan 26 2026 | PRJNA1269941 | sal squamous cell carcinoma |
| GSE209659 | Homo sapiens | Jul 28 2022 | PRJNA862069 | Peripheral blood |
| GSE206959 | Homo sapiens | Feb 15 2023 | PRJNA853053 | |
| GSE315240 | Homo sapiens | Feb 18 2026 | PRJNA1395394 | 2D cell culture |
| GSE288585 | Mus musculus | Jan 14 2026 | PRJNA1218635 | |
| GSE317185 | Mus musculus | Feb 16 2026 | PRJNA1406232 | |
| GSE310740 | Mus musculus | Jan 06 2026 | PRJNA1367011 | Lung |
| GSE297490 | Mus musculus | Feb 09 2026 | PRJNA1265075 | Colon |
| GSE222391 | Homo sapiens | Jan 12 2023 | PRJNA921852 | retinal epithelium |
| GSE287476 | Mus musculus | Feb 25 2026 | PRJNA1213066 | |
| GSE211189 | Homo sapiens | Aug 14 2023 | PRJNA869447 | |
| GSE314709 | Homo sapiens | Jan 30 2026 | PRJNA1392477 | lung |
| GSE235585 | Homo sapiens | Feb 15 2024 | PRJNA986502 | organoid |
| GSE300962 | Mus musculus | Jan 10 2026 | PRJNA1282748 | hippocampus |
| GSE311209 | Homo sapiens | Jan 05 2026 | PRJNA1368780 | pulmonary blastoma lung lesion |
| GSE262966 | Mus musculus | Jan 06 2026 | PRJNA1095009 | lung |
| GSE294703 | Homo sapiens | Feb 07 2026 | PRJNA1251343 | Male |
| GSE269307 | Mus musculus | Jan 06 2026 | PRJNA1120932 | |
| GSE315768 | Homo sapiens | Jan 11 2026 | | |
| GSE300791 | Homo sapiens | Feb 12 2026 | | spinal cord |
| GSE292915 | Arabidopsis thaliana | Jan 04 2026 | PRJNA1241865 | leaf |
| GSE303998 | Mus musculus | Feb 18 2026 | PRJNA1298600 | BAT |
| GSE273631 | Pantherophis guttatus | Mar 03 2025 | PRJNA1142529 | Brain and spinal cord |
| GSE307715 | Schistosoma mansoni | Jan 28 2026 | | whole worm |
| GSE292084 | Mus musculus | Feb 12 2026 | PRJNA1236831 | |
| GSE283761 | Ipomoea batatas | Jan 12 2026 | PRJNA1195365 | leaf |
| GSE315129 | Homo sapiens | Jan 31 2026 | PRJNA1395037 | |
| GSE269546 | Mus musculus | Feb 22 2026 | PRJNA1122490 | |
| GSE290341 | Mus musculus | Jan 21 2026 | PRJNA1227517 | Craniofacial |
| GSE307568 | Homo sapiens | Feb 14 2026 | PRJNA1322299 | Organoid |
| GSE312591 | Homo sapiens | Jan 01 2026 | PRJNA1374262 | Isolated skin fibroblasts |
| GSE315354 | Rattus norvegicus | Jan 04 2026 | PRJNA1396587 | nucleus pulposus |
| GSE285854 | Homo sapiens | Jan 01 2026 | PRJNA1206225 | tibial tumor |
| GSE295123 | Homo sapiens | Feb 27 2026 | PRJNA1253593 | |
| GSE236702 | Homo sapiens | Feb 16 2024 | | iPSC |
| GSE224109 | Mus musculus | Mar 15 2023 | PRJNA929917 | |
| GSE317937 | Homo sapiens | Jan 31 2026 | PRJNA1415858 | metastatic melanoma |
| GSE287728 | Homo sapiens | Jan 06 2026 | PRJNA1214346 | colon |
| GSE305385 | Mus musculus | Jan 05 2026 | PRJNA1305435 | liver |
| GSE221116 | Mus musculus | Dec 30 2022 | PRJNA912849 | |
| GSE165224 | Homo sapiens | Jan 21 2024 | PRJNA693698 | |

| | | | | |
|---|---|---|---|---|
| GSE145103 | Mus musculus | Feb 12 2020 | PRJNA606042 | |
| GSE315363 | Mus musculus | Jan 26 2026 | PRJNA1396597 | |
| GSE307722 | Gallus gallus | Jan 01 2026 | PRJNA1327142 | Liver |
| GSE310890 | Homo sapiens | Feb 20 2026 | PRJNA1367488 | |
| GSE311602 | Homo sapiens | Feb 12 2026 | PRJNA1369978 | Bone marrow |
| GSE305869 | Homo sapiens | Jan 27 2026 | PRJNA1308074 | e-derived xenografts FFPE tissue |
| GSE310101 | Streptomyces venezuelae | Feb 11 2026 | PRJNA1365131 | |
| GSE245540 | Mus musculus | Jun 06 2024 | PRJNA1028970 | retina |
| GSE115819 | Homo sapiens | Feb 23 2026 | PRJNA476132 | |
| GSE239298 | Mus musculus | Jan 14 2026 | PRJNA998781 | Bone marrow |
| GSE193491 | Homo sapiens | May 13 2022 | PRJNA796355 | blood |
| GSE316499 | Homo sapiens | Jan 19 2026 | PRJNA1403009 | skin |
| GSE244243 | Mus musculus | Oct 01 2024 | PRJNA1021989 | Retina |
| GSE252363 | Mus musculus | Jan 09 2026 | PRJNA1060289 | |
| GSE289586 | Mus musculus | Feb 10 2026 | PRJNA1223476 | Blood |
| GSE331464 | Homo sapiens | May 22 2026 | PRJNA1468192 | |
| GSE264319 | Homo sapiens | Feb 09 2026 | PRJNA1101965 | |
| GSE303302 | Mus musculus | Sep 30 2025 | PRJNA1294788 | bulk retina |
| GSE310722 | Homo sapiens | Jan 09 2026 | PRJNA1366984 | |
| GSE309902 | Homo sapiens | Feb 20 2026 | PRJNA1363961 | |
| GSE167931 | Homo sapiens | Mar 23 2022 | PRJNA705603 | |
| GSE317876 | Homo sapiens | Feb 28 2026 | PRJNA1415747 | |
| GSE268179 | Danio rerio | May 27 2024 | PRJNA1115053 | Eye |
| GSE179872 | Homo sapiens | Feb 09 2026 | PRJNA745441 | |
| GSE314009 | taurus;Equus caballus;Homo sap | Jan 14 2026 | PRJNA1389012 | cell line |
| GSE315215 | Homo sapiens | Jan 07 2026 | PRJNA1395135 | |
| GSE294761 | Danio rerio | Feb 16 2026 | PRJNA1251568 | endoderm-enriched |
| GSE229034 | Mus musculus | Dec 14 2023 | PRJNA952614 | Retina |
| GSE286320 | Homo sapiens | Feb 07 2026 | PRJNA1208670 | |
| GSE273770 | Mus musculus | Jan 01 2026 | PRJNA1143033 | Fully-grown GV oocyte |
| GSE311135 | Coccidioides posadasii | Feb 26 2026 | PRJNA1368715 | |
| GSE292784 | Mus musculus | Nov 27 2025 | PRJNA1241248 | Retina |
| GSE315629 | Homo sapiens | Jan 23 2026 | PRJNA1398355 | |
| GSE306716 | Mus musculus | Feb 22 2026 | PRJNA1311965 | mammary gland tumor |
| GSE226786 | Homo sapiens | Jan 01 2026 | PRJNA942049 | |
| GSE296587 | Mus musculus | Jan 15 2026 | PRJNA1260021 | Embryonic stem cells |
| GSE275795 | Mus musculus | Jan 28 2026 | PRJNA1153002 | small intestinal muscle |
| GSE319603 | Mus musculus | Feb 20 2026 | PRJNA1424711 | gastrocnemius muscle |
| GSE294120 | Mus musculus | Jan 20 2026 | | skin |
| GSE300823 | Sus scrofa | Jan 15 2026 | PRJNA1282154 | |
| GSE315454 | Mus musculus | Feb 19 2026 | PRJNA1397175 | istal part of small intestine |
| GSE291420 | Mus musculus | Feb 06 2026 | PRJNA1233337 | heart |
| GSE178930 | Mus musculus | Jun 26 2021 | PRJNA741510 | nal cord; intervertebral disc |
| GSE144428 | Mus musculus | Mar 09 2020 | PRJNA603599 | Retina EGFP- cells |
| GSE303574 | Homo sapiens | Jan 07 2026 | PRJNA1295857 | helium from umbilical cord vein |
| GSE316934 | Mus musculus | Jan 23 2026 | PRJNA1405552 | mal white adipose tissue (eWAT) |
| GSE291899 | allimastix lanati (nom. inval.);Pse | Jan 06 2026 | PRJNA1235741 | |
| GSE295614 | Xenopus laevis | Feb 05 2026 | PRJNA1255475 | f a Nieuwkoop and Faber stage 9 Xenopus laevis embryo (animal caps) |
| GSE113472 | Homo sapiens | Apr 10 2020 | PRJNA451514 | |
| GSE236035 | Drosophila melanogaster | Feb 24 2026 | PRJNA988475 | Ovary |
| GSE318428 | Homo sapiens | Feb 10 2026 | PRJNA1418716 | iPSC-derived RPE |
| GSE287028 | Homo sapiens | Jan 30 2026 | PRJNA1211012 | |
| GSE302240 | Rattus norvegicus | Jan 08 2026 | PRJNA1289696 | marrow |
| GSE283549 | Mus musculus | Jan 08 2026 | PRJNA1194508 | Bone marrow |
| GSE154684 | Homo sapiens | Oct 27 2021 | PRJNA647189 | |
| GSE279297 | Mus musculus | Jan 08 2026 | PRJNA1171815 | |
| GSE284693 | Mus musculus | Jan 05 2026 | PRJNA1200053 | primary islets |
| GSE293921 | Mus musculus | Jan 22 2026 | PRJNA1247434 | |
| GSE307022 | Homo sapiens | Jan 01 2026 | | |
| GSE319156 | Mus musculus | Feb 23 2026 | PRJNA1422219 | skin |
| GSE303932 | Homo sapiens | Sep 26 2025 | PRJNA1298254 | cell line |
| GSE291866 | Mus musculus | Jan 31 2026 | PRJNA1235686 | Liver |
| GSE142526 | Homo sapiens | Jan 10 2020 | PRJNA597325 | |

| | | | | |
|---|---|---|---|---|
| GSE279015 | Mus musculus | Feb 20 2026 | PRJNA1169913 | heart ventricle |
| GSE240483 | Mus musculus | Feb 13 2026 | | Heart |
| GSE301366 | Mus musculus | Feb 17 2026 | PRJNA1284857 | |
| GSE231472 | o sapiens;Mycobacterium tuberc | Feb 17 2026 | PRJNA966113 | Biopsy |
| GSE289627 | Mus musculus | Feb 08 2026 | PRJNA1223847 | bone marrow |
| GSE182944 | Danio rerio | Aug 27 2022 | PRJNA758397 | |
| GSE214255 | Homo sapiens | Jan 15 2026 | PRJNA884645 | |
| GSE142244 | Gallus gallus | Mar 01 2020 | PRJNA596379 | retina |
| GSE289338 | Homo sapiens | Feb 06 2026 | | Kidney tissue |
| GSE274214 | Mus musculus | Jul 06 2025 | PRJNA1145411 | whole retina |
| GSE263128 | Mus musculus | Sep 30 2024 | PRJNA1095897 | retina |
| GSE317513 | Mus musculus | Jan 30 2026 | PRJNA1414400 | liver |
| GSE242982 | Mus musculus | Dec 03 2024 | PRJNA1015665 | Retina |
| GSE164092 | omo sapiens;Oryctolagus cunicul | Feb 04 2021 | PRJNA688888 | |
| GSE243710 | Mus musculus | Jan 12 2026 | PRJNA1019670 | |
| GSE292264 | Mus musculus | Feb 17 2026 | PRJNA1237781 | spleen |
| GSE206623 | Mus musculus | Aug 20 2022 | PRJNA851590 | retina |
| GSE137910 | Homo sapiens | Sep 03 2022 | PRJNA573847 | |
| GSE292988 | Homo sapiens | Jan 08 2026 | PRJNA1242295 | |
| GSE307297 | Homo sapiens | Feb 06 2026 | PRJNA1315167 | Human Lung |
| GSE293444 | Homo sapiens | Feb 16 2026 | PRJNA1244634 | cell line |
| GSE285564 | Homo sapiens;Mus musculus | Dec 12 2025 | PRJNA1204365 | Retina |
| GSE294281 | Mus musculus | Jan 01 2026 | PRJNA1248934 | |
| GSE302847 | Homo sapiens | Feb 20 2026 | PRJNA1292265 | peripheral blood |
| GSE304010 | Mus musculus | Feb 09 2026 | PRJNA1298755 | CNS |
| GSE268186 | Homo sapiens | Jan 24 2026 | PRJNA1115059 | velopment-arrest 8C stage embryo |
| GSE270134 | Mus musculus | Jun 19 2026 | PRJNA1125309 | Rod PR |
| GSE316333 | Homo sapiens | Jan 14 2026 | PRJNA1402331 | |
| GSE317352 | Mus musculus | Feb 14 2026 | PRJNA1406780 | |
| GSE113917 | Macaca fascicularis | Oct 20 2020 | PRJNA454464 | Retina |
| GSE277902 | Mus musculus | Jan 21 2026 | PRJNA1164732 | Epidermis |
| GSE309226 | Homo sapiens | Jan 27 2026 | PRJNA1334367 | |
| GSE315097 | Homo sapiens | Jan 01 2026 | PRJNA1394977 | |
| GSE316451 | Homo sapiens;synthetic construc | Jan 18 2026 | PRJNA1402941 | |
| GSE284505 | Mus musculus | Dec 22 2024 | | Retina |
| GSE291972 | Mus musculus | Dec 02 2025 | PRJNA1236275 | retina |
| GSE281032 | Mus musculus | Jun 23 2025 | PRJNA1181863 | Retina |
| GSE139996 | Mus musculus | Jan 31 2020 | PRJNA587943 | etinal pigment epithelium |
| GSE303180 | Mus musculus | Dec 04 2025 | | Mouse retina |
| GSE274462 | Homo sapiens | Jan 15 2026 | PRJNA1146461 | |
| GSE317118 | Mus musculus | Jan 21 2026 | PRJNA1406122 | heart |
| GSE317407 | Caenorhabditis elegans | Jan 27 2026 | PRJNA1412540 | whole worm |
| GSE192881 | Homo sapiens | Oct 28 2022 | PRJNA793673 | |
| GSE203555 | Gallus gallus | May 19 2025 | PRJNA841372 | |
| GSE314950 | Mus musculus | Jan 12 2026 | PRJNA1393387 | brainstem tissue |
| GSE244811 | Mus musculus | Aug 13 2024 | PRJNA1024939 | Eye |
| GSE267661 | Homo sapiens | May 19 2024 | PRJNA1112365 | umbilical cord |
| GSE310383 | Homo sapiens | Jan 05 2026 | PRJNA1365859 | Peripheral blood |
| GSE247546 | Mus musculus | Jun 26 2024 | PRJNA1039519 | retina enriched sample |
| GSE281611 | Homo sapiens;Mus musculus | Feb 09 2026 | PRJNA1184797 | Bone |
| GSE188639 | ichia coli;Homo sapiens;Mus mu | Nov 11 2021 | PRJNA779733 | |
| GSE288459 | Homo sapiens | Jan 30 2026 | PRJNA1217799 | Bronchoalveolar lavage |
| GSE317224 | Homo sapiens | Feb 11 2026 | PRJNA1406303 | Rabdoid Sarcoma |
| GSE297053 | Homo sapiens | Feb 26 2026 | PRJNA1262448 | |
| GSE289127 | Sparus aurata | Feb 07 2026 | PRJNA1221596 | Proximal intestine |
| GSE181324 | Mus musculus | Dec 20 2022 | PRJNA751576 | retina |
| GSE315847 | Cryptococcus neoformans | Jan 12 2026 | PRJNA1399418 | Yeast culture |
| GSE296278 | Mus musculus | Mar 10 2026 | PRJNA1258771 | retina |
| GSE316510 | Mus musculus | Jan 21 2026 | PRJNA1403026 | Aorta |
| GSE279040 | Mus musculus | Jan 01 2026 | PRJNA1170221 | mammary |
| GSE304633 | Toxoplasma | Jan 01 2026 | PRJNA1301952 | Toxoplasma gondii |
| GSE159107 | Gallus gallus | Jan 04 2021 | PRJNA667705 | Retina |
| GSE138672 | Mus musculus | Nov 25 2020 | PRJNA576715 | retina |

| | | | | |
|---|---|---|---|---|
| GSE285347 | Homo sapiens | Jan 21 2026 | PRJNA1203208 | Peripheral blood |
| GSE306978 | Mus musculus | Jan 14 2026 | PRJNA1314061 | |
| GSE281339 | Homo sapiens | Feb 27 2026 | PRJNA1183405 | pheral venous blood basophils |
| GSE296005 | Mus musculus | Sep 08 2025 | PRJNA1257309 | |
| GSE312071 | Mus musculus | Jan 13 2026 | PRJNA1372141 | Cortex |
| GSE267656 | Homo sapiens | May 01 2025 | PRJNA1112343 | |
| GSE306294 | Homo sapiens | Jan 30 2026 | PRJNA1310424 | |
| GSE299944 | Plasmodium falciparum | Feb 16 2026 | PRJNA1277590 | |
| GSE316433 | Mus musculus | Jan 20 2026 | PRJNA1402878 | heral blood mononuclear cells |
| GSE291910 | Mus musculus | Jan 16 2026 | PRJNA1235762 | Spleen |
| GSE313133 | Mus musculus | Feb 13 2026 | PRJNA1378694 | |
| GSE269814 | Mus musculus | Feb 18 2026 | PRJNA1123740 | ureter |
| GSE262999 | Homo sapiens | Feb 15 2026 | PRJNA1095432 | Cell line |
| GSE297458 | Mus musculus | Jan 22 2026 | PRJNA1263882 | Calvaria |
| GSE223338 | Homo sapiens | Jan 10 2026 | PRJNA925838 | tumor |
| GSE292325 | Homo sapiens | Jan 12 2026 | PRJNA1237938 | meningioma |
| GSE263576 | Apis mellifera | Jan 21 2026 | PRJNA1098086 | |
| GSE180779 | Homo sapiens | Jul 01 2022 | PRJNA749560 | |
| GSE288065 | Homo sapiens | Jan 22 2026 | PRJNA1215984 | rhabdomyosarcoma |
| GSE293411 | Homo sapiens | Jan 12 2026 | PRJNA1244383 | aorta |
| GSE317465 | Homo sapiens | Jan 30 2026 | PRJNA1413098 | |
| GSE146897 | Mus musculus | Aug 25 2020 | PRJNA612318 | |
| GSE155039 | Mus musculus | Aug 13 2023 | PRJNA648319 | |
| GSE159771 | Mus musculus | Jun 30 2021 | PRJNA670470 | retina |
| GSE312165 | Rattus norvegicus | Jan 14 2026 | PRJNA1372505 | Dorsal root ganglion |
| GSE232864 | Mus musculus | Feb 25 2026 | PRJNA974248 | Mouse brain |
| GSE276936 | Mus musculus | Jan 13 2026 | PRJNA1159777 | brain |
| GSE283879 | Mus musculus | Jan 05 2026 | PRJNA1195955 | testis |
| GSE149797 | Homo sapiens | Feb 12 2021 | PRJNA630173 | cortical brain organoids |
| GSE308212 | Mus musculus | Jan 13 2026 | PRJNA1329221 | |
| GSE143406 | Homo sapiens | May 07 2020 | PRJNA600307 | |
| GSE282496 | Homo sapiens | Feb 12 2026 | PRJNA1189025 | Cord blood |
| GSE254727 | Mus musculus | Feb 15 2026 | PRJNA1071585 | |
| GSE296673 | Danio rerio | May 01 2026 | PRJNA1260495 | eye |
| GSE318596 | Homo sapiens | Mar 23 2026 | PRJNA1419285 | |
| GSE274400 | Homo sapiens | Jan 20 2026 | PRJNA1146350 | Breast |
| GSE287489 | Mus musculus | Jan 25 2025 | PRJNA1213075 | Organoids |
| GSE319295 | Homo sapiens | Feb 18 2026 | PRJNA1422786 | |
| GSE215959 | Mus musculus | Feb 18 2026 | PRJNA891349 | anoma cell line-derived tumor |
| GSE287734 | Mus musculus | Jan 24 2025 | PRJNA1214356 | Retina |
| GSE304185 | Homo sapiens | Jan 04 2026 | PRJNA1299348 | |
| GSE290408 | Homo sapiens | Feb 09 2026 | PRJNA1227924 | lung |
| GSE268013 | Homo sapiens | Feb 26 2026 | PRJNA1114041 | |
| GSE315893 | Mus musculus | Jan 12 2026 | PRJNA1399744 | Lung |
| GSE314601 | Homo sapiens | Jan 21 2026 | PRJNA1392191 | ad and Neck Cancer cell line |
| GSE212470 | Homo sapiens | Nov 04 2022 | PRJNA875473 | |
| GSE310402 | Danio rerio | Jul 01 2026 | PRJNA1365889 | whole embryo |
| GSE285393 | Homo sapiens | Jun 01 2025 | PRJNA1203270 | Retinal organoid |
| GSE315102 | Mus musculus | Feb 24 2026 | PRJNA1395006 | Lymph node |
| GSE305842 | Mus musculus | Jan 30 2026 | PRJNA1308006 | Mammary |
| GSE245135 | Mus musculus | Jan 08 2026 | PRJNA1026990 | Heart |
| GSE193975 | Sus scrofa domesticus | Jan 21 2022 | PRJNA798674 | Area Postrema |
| GSE208730 | Mus musculus | Feb 16 2026 | PRJNA860829 | Back skin epidermis |
| GSE149102 | Oryctolagus cuniculus | Jul 21 2020 | PRJNA627392 | retina |
| GSE314636 | Homo sapiens | Jan 01 2026 | PRJNA1392354 | |
| GSE270907 | Mus musculus | Jan 12 2026 | PRJNA1128811 | spleen |
| GSE276150 | Homo sapiens | Jan 01 2026 | PRJNA1155181 | |
| GSE298174 | Homo sapiens | Jan 22 2026 | PRJNA1268462 | |
| GSE316291 | Bos taurus | Jan 19 2026 | PRJNA1402247 | Mammary fat pad |
| GSE296977 | Mus musculus | Jan 13 2026 | PRJNA1262309 | liver |
| GSE272353 | Mus musculus | Jan 23 2025 | PRJNA1136503 | Retina |
| GSE315348 | Rattus norvegicus | Jan 05 2026 | PRJNA1395863 | Septum |
| GSE290120 | Mus musculus | Jan 28 2026 | PRJNA1226230 | |

| | | | | |
|---|---|---|---|---|
| GSE313841 | Mus musculus | Jan 22 2026 | PRJNA1381660 | Pancreatic |
| GSE335331 | Pleurodeles waltl | Jun 16 2026 | PRJNA1477926 | Retina |
| GSE230908 | Macaca mulatta | Feb 14 2026 | | Bone marrow |
| GSE282442 | Homo sapiens | Feb 28 2026 | PRJNA1188590 | Colon |
| GSE319241 | Homo sapiens | Feb 11 2026 | PRJNA1422661 | |
| GSE315987 | Mus musculus | Jan 13 2026 | PRJNA1400065 | heart |
| GSE166919 | Rattus norvegicus | Feb 14 2022 | PRJNA702293 | retinal tissue |
| GSE223837 | Homo sapiens | Jan 26 2026 | PRJNA928488 | |
| GSE315774 | Homo sapiens | Feb 25 2026 | PRJNA1398938 | |
| GSE270824 | Mus musculus | Feb 11 2026 | PRJNA1128580 | |
| GSE311068 | Homo sapiens | Feb 18 2026 | PRJNA1368624 | |
| GSE294200 | Mus musculus | Feb 13 2026 | PRJNA1248541 | nixed sample oVAT/Spleen |
| GSE302157 | Mus musculus | Jan 29 2026 | PRJNA1289094 | Retina |
| GSE190263 | Homo sapiens | Dec 28 2022 | PRJNA786562 | |
| GSE225702 | nchius furzeri;Nothobranchius o | Jan 28 2026 | PRJNA937196 | brain |
| GSE184086 | Caenorhabditis elegans | Dec 31 2022 | PRJNA763157 | Whole animal |
| GSE313685 | Mus musculus | Feb 17 2026 | PRJNA1381337 | heart |
| GSE291228 | Mus musculus | Jan 12 2026 | PRJNA1232607 | cochlea |
| GSE294728 | Saccharomyces cerevisiae | Feb 18 2026 | PRJNA1251513 | yeast cell |
| GSE286885 | Homo sapiens | Feb 18 2026 | PRJNA1210487 | |
| GSE312186 | Mus musculus | Jan 12 2026 | | Embryonic stem cell |
| GSE148077 | Homo sapiens | Jun 22 2020 | PRJNA623011 | |
| GSE271681 | Homo sapiens | Jan 07 2026 | PRJNA1132918 | |
| GSE317486 | Mus musculus | Feb 28 2026 | PRJNA1413414 | Brain tumor |
| GSE310243 | Homo sapiens | Feb 15 2026 | PRJNA1365523 | |
| GSE294237 | Mus musculus | Feb 06 2026 | PRJNA1248857 | Aorta tissue |
| GSE307302 | Mus musculus | Jun 03 2026 | PRJNA1315172 | Retina |
| GSE316869 | mouse gut metagenome | Jan 21 2026 | PRJNA1405456 | |
| GSE161365 | Homo sapiens | Jan 16 2026 | PRJNA677961 | |
| GSE160179 | Danio rerio | Jul 31 2021 | PRJNA672149 | |
| GSE205370 | Homo sapiens | Oct 05 2022 | PRJNA844894 | human retina explant |
| GSE315021 | Oryza sativa | Jan 16 2026 | | leaf |
| GSE295608 | Mus musculus | Apr 24 2025 | PRJNA1255295 | stem cell-derived neuroectoderm organoid |
| GSE297943 | Rattus norvegicus | May 01 2026 | PRJNA1267001 | retina |
| GSE314890 | Rattus norvegicus | Jan 01 2026 | PRJNA1393091 | urethral tissue |
| GSE128140 | Homo sapiens | Jul 11 2020 | PRJNA526536 | |
| GSE284390 | Homo sapiens | Feb 12 2026 | PRJNA1198925 | |
| GSE307521 | Mus musculus | Jul 01 2026 | PRJNA1321908 | choroidia |
| GSE304876 | Mus musculus | Jan 13 2026 | PRJNA1303134 | Cortex |
| GSE200053 | Mus musculus | Apr 03 2025 | PRJNA822801 | RPE/choroid/sclera |
| GSE318649 | Mus musculus | Feb 07 2026 | PRJNA1419708 | |
| GSE260983 | Mus musculus | Feb 14 2025 | PRJNA1084778 | Brain |
| GSE314136 | Mus musculus | Jan 07 2026 | PRJNA1390159 | Large intestine |
| GSE312961 | Homo sapiens | Jan 04 2026 | PRJNA1377665 | cell culture |
| GSE294341 | Mus musculus | Feb 12 2026 | PRJNA1249072 | mesenteric adipose tissue |
| GSE176352 | Rattus norvegicus | Jul 01 2021 | PRJNA735941 | retina |
| GSE310335 | Homo sapiens | Feb 25 2026 | PRJNA1365773 | lung |
| GSE256215 | Mus musculus;synthetic construc | Nov 22 2024 | PRJNA1078421 | Retina |
| GSE199317 | Mus musculus | Feb 03 2023 | PRJNA819392 | |
| GSE237202 | Bos taurus | Oct 01 2023 | PRJNA994166 | Retina |
| GSE281854 | Mus musculus | Nov 15 2024 | PRJNA1186057 | Liver |
| GSE136120 | Homo sapiens | Aug 20 2022 | PRJNA561341 | |
| GSE145069 | Homo sapiens | Feb 11 2020 | PRJNA605854 | |
| GSE164152 | Rattus norvegicus | Apr 27 2021 | PRJNA689298 | Retinal tissue |
| GSE186321 | Homo sapiens | Jan 05 2026 | PRJNA773393 | |
| GSE299229 | Mus musculus | Jan 21 2026 | PRJNA1273111 | bone marrow cells |
| GSE299736 | Mus musculus | Jan 10 2026 | PRJNA1276341 | kidney |
| GSE305092 | Mus musculus | Jan 05 2026 | | Pancreatic Islet |
| GSE191260 | Mus musculus | Nov 11 2022 | PRJNA790816 | Retina |
| GSE315074 | Homo sapiens | Jan 26 2026 | PRJNA1394957 | |
| GSE297857 | Mus musculus | Feb 27 2026 | PRJNA1266596 | liver |
| GSE303154 | Macaca mulatta | Feb 25 2026 | PRJNA1294242 | granuloma |
| GSE176513 | Homo sapiens | Mar 09 2022 | PRJNA736472 | |

| | | | | |
|---|---|---|---|---|
| GSE268630 | Homo sapiens | May 30 2024 | PRJNA1117936 | peripheral region of retina |
| GSE248517 | Mus musculus | Mar 03 2025 | PRJNA1044416 | 6-week old mouse eye |
| GSE313814 | Homo sapiens | Feb 25 2026 | PRJNA1381625 | kidney |
| GSE282899 | Drosophila melanogaster | Jan 21 2026 | PRJNA1190966 | larval brain |
| GSE288299 | Mus musculus | Feb 16 2026 | PRJNA1216885 | |
| GSE282108 | Mus musculus | Jan 17 2026 | PRJNA1187209 | substantia nigra |
| GSE315721 | Saccharomyces cerevisiae | Jan 14 2026 | PRJNA1398847 | |
| GSE319069 | Podarcis muralis | Feb 18 2026 | PRJNA1422040 | Embryo |
| GSE336951 | Mus musculus | Jun 29 2026 | PRJNA1483627 | Dorsal |
| GSE286176 | Homo sapiens | Jan 07 2026 | PRJNA1208062 | |
| GSE296801 | Homo sapiens | Feb 19 2026 | PRJNA1261064 | |
| GSE146187 | Sus scrofa | May 19 2020 | PRJNA609680 | corneoscleral wedges |
| GSE317379 | Homo sapiens | Jan 30 2026 | PRJNA1412079 | dlPFC |
| GSE310932 | Homo sapiens | Feb 24 2026 | PRJNA1367566 | |
| GSE302380 | Homo sapiens | Sep 17 2025 | PRJNA1290284 | |
| GSE305545 | Homo sapiens | Jan 19 2026 | PRJNA1306488 | |
| GSE281531 | Homo sapiens | Feb 09 2026 | PRJNA1184334 | |
| GSE299583 | Homo sapiens | Jan 12 2026 | PRJNA1275018 | HUDEP1 |
| GSE170893 | Homo sapiens | Mar 30 2021 | | |
| GSE331198 | Mus musculus | May 21 2026 | PRJNA1467467 | Retina |
| GSE308980 | Escherichia coli | Jan 07 2026 | PRJNA1333525 | bacterial cell |
| GSE286496 | Mus musculus | Jan 01 2026 | | kidney |
| GSE281253 | Homo sapiens | Apr 24 2025 | PRJNA1182812 | Osteosarcoma |
| GSE285050 | Homo sapiens | Feb 24 2026 | PRJNA1201120 | undifferentiated iPSCs |
| GSE309070 | Homo sapiens | Jan 18 2026 | PRJNA1333896 | cell lines |
| GSE319589 | Mus musculus | Feb 19 2026 | PRJNA1423794 | Brain |
| GSE317133 | Homo sapiens | Jan 27 2026 | PRJNA1406138 | liver |
| GSE151273 | Danio rerio | Jan 06 2021 | PRJNA635369 | Head |
| GSE274634 | Homo sapiens | Jan 14 2026 | PRJNA1147318 | |
| GSE254513 | Homo sapiens | Nov 16 2025 | PRJNA1070689 | Stomach (fundus) |
| GSE316630 | Caenorhabditis elegans | Jan 15 2026 | | whole organism |
| GSE202762 | Homo sapiens | Jan 13 2026 | PRJNA837230 | |
| GSE223659 | Mus musculus;synthetic construc | Nov 08 2023 | PRJNA927048 | Retina |
| GSE214642 | Danio rerio | Nov 23 2022 | PRJNA886442 | head |
| GSE185718 | Mus musculus | Oct 01 2022 | PRJNA770620 | hypothalamus |
| GSE280078 | Mus musculus | Feb 06 2026 | PRJNA1176148 | Tumor |
| GSE234718 | Drosophila melanogaster | Feb 24 2026 | PRJNA982813 | Ovary |
| GSE251771 | Homo sapiens | Feb 18 2026 | PRJNA1055444 | |
| GSE284997 | Homo sapiens | Jan 21 2026 | PRJNA1200769 | |
| GSE158759 | Mus musculus | Jan 13 2022 | PRJNA666431 | |
| GSE315594 | Drosophila melanogaster | Jan 05 2026 | PRJNA1398313 | gut |
| GSE316152 | Homo sapiens | Jan 15 2026 | PRJNA1401734 | Stomach |
| GSE275655 | Mus musculus | Feb 11 2026 | PRJNA1152507 | Retina |
| GSE243452 | Drosophila melanogaster | Feb 27 2026 | PRJNA1018521 | embryo |
| GSE167157 | Homo sapiens | Jan 16 2026 | PRJNA703472 | |
| GSE151350 | Mus musculus | Sep 05 2021 | PRJNA635599 | Retina |
| GSE308667 | Mus musculus | Jan 13 2026 | PRJNA1331851 | testis |
| GSE162148 | sapiens;Macaca mulatta;Mus mu | Apr 20 2021 | PRJNA680759 | ominal subcutaneous adipose |
| GSE316713 | Homo sapiens | Feb 25 2026 | PRJNA1403812 | coronary artery |
| GSE289313 | Homo sapiens | Feb 28 2026 | PRJNA1222395 | |
| GSE192507 | Mus musculus | Aug 16 2022 | PRJNA791674 | Retina |
| GSE174215 | Homo sapiens | Sep 01 2022 | PRJNA728798 | SC-derived retinal organoid |
| GSE144845 | Mus musculus | Mar 10 2021 | PRJNA605145 | |
| GSE256097 | Mus musculus | Feb 18 2026 | PRJNA1078249 | |
| GSE317772 | Mus musculus | Jan 30 2026 | PRJNA1415209 | |
| GSE305451 | Homo sapiens | Jan 13 2026 | PRJNA1306003 | Brain |
| GSE155631 | Homo sapiens;Mus musculus | Feb 11 2026 | PRJNA650522 | |
| GSE307794 | Thermus thermophilus HB27 | Feb 25 2026 | PRJNA1327337 | |
| GSE292005 | Homo sapiens | Jan 17 2026 | PRJNA1236319 | |
| GSE284125 | Mus musculus | Apr 28 2025 | PRJNA1197399 | etinal pigment epithelium |
| GSE325741 | Homo sapiens | Mar 25 2026 | | Eye; Pigmented epithelium; Retina |
| GSE279009 | Mus musculus | Feb 20 2026 | PRJNA1169905 | heart ventricle |
| GSE329009 | Aquarana catesbeiana | Apr 30 2026 | PRJNA1457374 | Liver |

| | | | | |
|---|---|---|---|---|
| GSE316046 | Mus musculus | Jan 14 2026 | PRJNA1400515 | |
| GSE294865 | Homo sapiens | Jan 21 2026 | PRJNA1251885 | jection subcutaneously immune-deficient mice. |
| GSE168093 | Mus musculus | Jun 18 2021 | PRJNA706014 | retina |
| GSE209908 | Mus musculus | Jul 29 2022 | PRJNA862918 | RPE/choroid |
| GSE223530 | Mus musculus | Feb 21 2024 | PRJNA926655 | Retina |
| GSE301665 | Homo sapiens | Feb 26 2026 | PRJNA1285945 | |
| GSE326387 | Mus musculus | Apr 24 2026 | PRJNA1445195 | retina |
| GSE313224 | Mus musculus | Jan 13 2026 | PRJNA1378936 | FL-BMDC |
| GSE153933 | Salmo salar | Jul 08 2020 | PRJNA644578 | Lens |
| GSE288138 | Homo sapiens | Jan 22 2026 | PRJNA1216264 | brain |
| GSE319824 | Homo sapiens | Feb 24 2026 | PRJNA1425158 | Bone marrow |
| GSE278068 | Homo sapiens;Mus musculus | Jan 10 2026 | PRJNA1165323 | |
| GSE247166 | Mus musculus | Feb 18 2026 | PRJNA1036557 | |
| GSE210414 | Mus musculus | Aug 08 2023 | PRJNA865691 | RPE |
| GSE309012 | Homo sapiens | Jan 01 2026 | PRJNA1333718 | |
| GSE303383 | Mus musculus | Feb 27 2026 | PRJNA1294980 | Liver |
| GSE319965 | Mus musculus | Feb 20 2026 | PRJNA1425796 | colonic tumor |
| GSE317479 | Mus musculus | Jan 29 2026 | PRJNA1413379 | Whole heart |
| GSE263042 | Mus musculus | Jan 06 2026 | PRJNA1095508 | lung |
| GSE304059 | Homo sapiens | Feb 28 2026 | PRJNA1298846 | Kidney, tumor tissue |
| GSE312179 | Homo sapiens | Feb 12 2026 | PRJNA1372607 | |
| GSE302480 | Mus musculus | Feb 06 2026 | PRJNA1291063 | Epidermis |
| GSE317604 | Rattus norvegicus | Jan 29 2026 | PRJNA1414533 | adrenal gland |
| GSE304224 | Homo sapiens | Jan 29 2026 | PRJNA1299432 | |
| GSE296370 | Homo sapiens | Feb 09 2026 | PRJNA1259001 | Tonsil |
| GSE230051 | Homo sapiens | Jan 09 2026 | PRJNA957288 | fetal liver organoids from iPSCs |
| GSE319289 | Mus musculus | Feb 12 2026 | PRJNA1422780 | Skeletal Muscle, Plantaris |
| GSE214175 | Homo sapiens | Mar 13 2023 | PRJNA884311 | Retina (female) |
| GSE239731 | Mus musculus | Jan 15 2024 | PRJNA1000634 | |
| GSE316130 | Mus musculus | Jan 12 2026 | | brain |
| GSE180518 | Danio rerio | Oct 11 2021 | PRJNA748446 | whole retina |
| GSE309258 | Mus musculus | Sep 30 2025 | PRJNA1334826 | retina |
| GSE313430 | Mus musculus | Jan 15 2026 | PRJNA1379503 | Bone marrow |
| GSE285767 | Homo sapiens | Jan 31 2026 | PRJNA1205798 | cell line |
| GSE299958 | Mus musculus | Jan 14 2026 | PRJNA1277720 | spleen |
| GSE272963 | Plasmodium falciparum | Feb 25 2026 | PRJNA1139717 | Enriched blood sample |
| GSE296347 | Mus musculus | Feb 09 2026 | PRJNA1258962 | Embryo |
| GSE311097 | Mus musculus | Jan 21 2026 | PRJNA1368667 | brain |
| GSE314597 | Mus musculus | Feb 18 2026 | PRJNA1392189 | Tendon tissue |
| GSE241906 | Caenorhabditis elegans | Jul 16 2024 | PRJNA1010732 | whole worm |
| GSE315294 | Homo sapiens | Jan 01 2026 | PRJNA1395596 | cell |
| GSE297567 | Mus musculus | Feb 17 2026 | PRJNA1265200 | small intestine |
| GSE316013 | Mus musculus | Jan 14 2026 | PRJNA1400460 | |
| GSE306964 | Mus musculus | Sep 04 2025 | PRJNA1314055 | retina |
| GSE301630 | Canis lupus familiaris | Jan 12 2026 | PRJNA1285841 | Blood, PBMC |
| GSE292578 | Mus musculus | Jan 10 2026 | PRJNA1240242 | racture Callus, Periosteum |
| GSE203642 | Homo sapiens;Mus musculus | Feb 14 2023 | PRJNA841670 | |
| GSE307094 | Homo sapiens | Feb 10 2026 | PRJNA1314248 | Brain |
| GSE316771 | Homo sapiens | Jan 21 2026 | PRJNA1404173 | breast |
| GSE270576 | Schizosaccharomyces pombe | Jan 27 2026 | PRJNA1127509 | |
| GSE293519 | Mus musculus | Jan 21 2026 | PRJNA1244867 | Stomach |
| GSE333449 | Homo sapiens | May 28 2026 | PRJNA1471117 | |
| GSE312572 | Mus musculus | Feb 17 2026 | PRJNA1374201 | Hippocampus |
| GSE174515 | Drosophila melanogaster | Dec 31 2021 | PRJNA730340 | |
| GSE295851 | Mus musculus | Jan 13 2026 | PRJNA1256485 | |
| GSE299397 | Homo sapiens | Feb 11 2026 | PRJNA1274359 | |
| GSE62020 | Mus musculus | Sep 25 2020 | PRJNA262966 | retina |
| GSE255679 | Mus musculus | Feb 13 2026 | PRJNA1076149 | Hippocampus |
| GSE231882 | Mus musculus | Dec 18 2023 | PRJNA970059 | Neuroretina |
| GSE174522 | Mus musculus | Nov 23 2022 | PRJNA730426 | Retinas and optic nerves |
| GSE250379 | Danio rerio | Oct 23 2024 | PRJNA1053781 | retina |
| GSE285111 | Mus musculus | Feb 10 2026 | PRJNA1201182 | bone marrow |
| GSE246746 | Homo sapiens | Jan 01 2026 | PRJNA1034548 | cell line |

| | | | | |
|---|---|---|---|---|
| GSE285673 | Mus musculus | Jan 12 2026 | PRJNA1205557 | spleen |
| GSE318827 | Mus musculus | Feb 12 2026 | PRJNA1420681 | |
| GSE295695 | Mus musculus | Feb 12 2026 | PRJNA1255614 | heart |

Table S2: List of queries for benchmarking

| Query number | GEOBERT Query | GEO query | QueryType |
|---|---|---|---|
| q1 | Are there any proteomics or plasma-based datasets? | proteomics OR plasma-based | Rare |
| q2 | Are there datasets that study specific genetic variants in retinal disease? | retinal disease AND genetic variants | 2 clauses |
| q3 | Control-1 vs Disease-1 comparisons in human blood samples with single-cell RNA-seq ? | human blood samples AND single-cell RNA-seq with control vs disease | 2+ clauses |
| q4 | Dataset having filtered_feature_bc_matrix.h5? | filtered_bc_matrix.h5 | DataType |
| q5 | Dual Programmed Death Receptor-1 and Vascular Endothelial Growth Factor Receptor-2 info containg GSE ? | Dual Programmed Death Receptor-1 AND Vascular Endothelial Growth Factor Receptor-2 | Rare |
| q6 | find me human retina datasets with transcription factor chipseq ? | human retina AND transcription factor chipseq | 2 clauses |
| q7 | Which GSEs use 10x Genomics,Äìbased single-cell sequencing, and which use bulk RNA-seq? | 10x Genomics,Äìbased single-cell sequencing AND bulk RNA-seq | 2+ clauses |
| q8 | find retina scRNA-seq datasets that provide a gene-by-cell count matrix (e.g., matrix.mtx, filtered_feature_bc_matrix.h5) | retina AND scRNA-seq with gene-by-cell count matrix | DataType |
| q9 | give me retina datasets where the samples were collected within 24h of decease? | retina AND 24h of decease | 2 clauses |
| q10 | How many transcription factor chipsed data have? | transcription factor chipseq | 2 clauses |

Table S3: Retrived GSEIDS by GEO, GEO-BERT and Gemini and the corresponding groundtruth with respect to the query

| GSE ID | Ground Truth | GEO Decision | GEO-BERT Decision | Gemini Decision | Query Number | Comments |
|---|---|---|---|---|---|---|
| GSE153672 | FALSE | | FALSE | | q1 | |
| GSE153674 | FALSE | | FALSE | | q1 | |
| GSE161369 | FALSE | | FALSE | | q1 | |
| GSE168129 | FALSE | TRUE | | | q1 | |
| GSE183117 | FALSE | TRUE | | | q1 | |
| GSE186458 | FALSE | | FALSE | | q1 | |
| GSE190615 | FALSE | TRUE | | | q1 | proteome profiling was done but no data here |
| GSE196106 | FALSE | | FALSE | | q1 | |
| GSE199867 | FALSE | TRUE | | | q1 | abstract mentions proteomics, bt no proteomics data |
| GSE210745 | FALSE | | FALSE | | q1 | |
| GSE234454 | FALSE | TRUE | | | q1 | proteomics was done but not data |
| GSE236627 | FALSE | TRUE | | | q1 | mentions about doing maass/spec, but not data |
| GSE237203 | FALSE | | FALSE | | q1 | |
| GSE237205 | FALSE | | FALSE | | q1 | |
| GSE237206 | FALSE | | FALSE | | q1 | |
| GSE237209 | FALSE | | FALSE | | q1 | |
| GSE237211 | FALSE | | FALSE | | q1 | |
| GSE237212 | FALSE | | FALSE | | q1 | |
| GSE253809 | FALSE | TRUE | | | q1 | proteomics without data |
| GSE255194 | FALSE | TRUE | | | q1 | proteomics without data |
| GSE255195 | FALSE | TRUE | | | q1 | mention protein but not proteomics or data |
| GSE255561 | FALSE | TRUE | | | q1 | mention protein but not proteomics or data |
| GSE261379 | TRUE | TRUE | | | q1 | Did proteomics but no data |
| GSE261578 | FALSE | TRUE | FALSE | | q1 | made pacbio library but not proteomics data |
| GSE263576 | FALSE | | FALSE | | q1 | |
| GSE272353 | FALSE | TRUE | | | q1 | Mention proteomics but no data |
| GSE283858 | FALSE | TRUE | | | q1 | |
| GSE285561 | TRUE | | FALSE | | q1 | Phospho-seq |
| GSE287770 | FALSE | TRUE | FALSE | | q1 | Mentions proteomics but no data |
| GSE289257 | FALSE | TRUE | | | q1 | |
| GSE289585 | FALSE | TRUE | | | q1 | Mentions proteomics but no data |
| GSE289920 | FALSE | TRUE | | | q1 | |
| GSE289973 | FALSE | TRUE | | | q1 | |
| GSE290324 | FALSE | TRUE | | | q1 | |
| GSE290325 | FALSE | TRUE | | | q1 | Abstract has the word proteomics |
| GSE295433 | FALSE | TRUE | | | q1 | |
| GSE295435 | FALSE | TRUE | | | q1 | |
| GSE296927 | FALSE | TRUE | | | q1 | |
| GSE297733 | FALSE | TRUE | | | q1 | |
| GSE298286 | FALSE | TRUE | | | q1 | |
| GSE299477 | FALSE | TRUE | | | q1 | |
| GSE299736 | FALSE | TRUE | | | q1 | |
| GSE301720 | FALSE | TRUE | | | q1 | |
| GSE304020 | FALSE | TRUE | | | q1 | PROSeq |
| GSE304185 | FALSE | TRUE | | | q1 | |
| GSE304253 | FALSE | TRUE | | | q1 | SLAM-seq |
| GSE304493 | FALSE | TRUE | | | q1 | RNASeq |
| GSE304612 | FALSE | TRUE | | | q1 | ChipSeq of transcription factor protein |
| GSE304740 | FALSE | TRUE | | | q1 | CRISPRscreen |
| GSE305037 | FALSE | TRUE | | | q1 | PerturbSeq |
| GSE305384 | FALSE | TRUE | | | q1 | SuperSeries |
| GSE306473 | FALSE | | FALSE | | q1 | |
| GSE307099 | FALSE | TRUE | | | q1 | Used external proteomics data |
| GSE307920 | FALSE | TRUE | | | q1 | ChIPSeq of Tf protein and probabaly used external proteomics data |
| GSE307922 | FALSE | TRUE | | | q1 | RNASeq and probabaly used external proteomics data |
| GSE308162 | TRUE | TRUE | TRUE | | q1 | plasma sample |
| GSE308745 | FALSE | | FALSE | | q1 | |
| GSE308921 | FALSE | TRUE | | | q1 | analysis included proteomics data |
| GSE309277 | FALSE | TRUE | | | q1 | scRNASeq; did proteomics |
| GSE309691 | FALSE | TRUE | | | q1 | has the word proteostasis in abstract |
| GSE310070 | FALSE | | FALSE | | q1 | microRNA |
| GSE310929 | FALSE | | FALSE | | q1 | |
| GSE311312 | FALSE | TRUE | | | q1 | Cut&Tag; probably sed external proteomics |
| GSE311314 | FALSE | TRUE | | | q1 | RNASeq; probably sed external proteomics |
| GSE311320 | TRUE | TRUE | TRUE | | q1 | Plasma samples; did proteomics |
| GSE312336 | TRUE | | FALSE | | q1 | surface protein expression (ADT) |
| GSE313668 | FALSE | TRUE | | | q1 | journal name contains proteomics |
| GSE313682 | FALSE | TRUE | | | q1 | External proteomics analysis |
| GSE314132 | FALSE | TRUE | | | q1 | proteomics done; no data |
| GSE314950 | FALSE | TRUE | | | q1 | RNASeq; mentions the word proteomics in abstract |
| GSE315605 | FALSE | TRUE | | | q1 | External proteomics analysis |
| GSE315877 | FALSE | TRUE | FALSE | | q1 | External proteomics analysis |
| GSE315959 | FALSE | | FALSE | | q1 | |
| GSE315996 | FALSE | TRUE | | | q1 | ChIPSeq; external proteimcs data |
| GSE315997 | FALSE | TRUE | | | q1 | ChIPSeq; external proteimcs data |
| GSE316009 | FALSE | TRUE | | | q1 | ChIPSeq; external proteimcs data |

| | | | | | | |
|---|---|---|---|---|---|---|
| GSE316495 | FALSE | TRUE | | | q1 | ChIPSeq of TF protein |
| GSE316811 | TRUE | TRUE | | | q1 | milti-omics including sptail proteomics |
| GSE316823 | FALSE | TRUE | | | q1 | RNASeq; probably external proteomics data was used |
| GSE316872 | FALSE | | FALSE | | q1 | |
| GSE317789 | FALSE | TRUE | | | q1 | external proteomics data used |
| GSE318067 | TRUE | TRUE | TRUE | | q1 | |
| GSE318248 | TRUE | | FALSE | | q1 | protein array by protein profiling |
| GSE318608 | FALSE | TRUE | | | q1 | ribosome profiling |
| GSE319154 | FALSE | TRUE | | | q1 | external proteomics used |
| GSE319215 | FALSE | TRUE | | | q1 | external proteomics data for plasma cell used |
| GSE319465 | FALSE | TRUE | | | q1 | RNASeq; mention proteomics |
| GSE319710 | FALSE | TRUE | | | q1 | external proteomics data for plasma cell used |
| GSE319779 | FALSE | TRUE | | | q1 | CRISPR; might have used external proteomics |
| GSE319783 | FALSE | TRUE | | | q1 | RNASeq; might have used external proteomics |
| GSE325522 | FALSE | | FALSE | | q1 | proteomics done but not present |
| GSE325523 | FALSE | | FALSE | | q1 | |
| GSE117956 | FALSE | | FALSE | | q2 | |
| GSE165322 | TRUE | | TRUE | | q2 | |
| GSE172170 | TRUE | | | TRUE | q2 | |
| GSE179754 | TRUE | | | TRUE | q2 | |
| GSE180641 | TRUE | TRUE | | | q2 | |
| GSE183684 | TRUE | | TRUE | | q2 | |
| GSE188280 | FALSE | | FALSE | | q2 | |
| GSE193161 | TRUE | | TRUE | | q2 | risk genes were identified |
| GSE196235 | TRUE | | TRUE | | q2 | eye disease |
| GSE201677 | TRUE | TRUE | TRUE | | q2 | |
| GSE201678 | TRUE | TRUE | TRUE | | q2 | |
| GSE201679 | TRUE | TRUE | TRUE | | q2 | |
| GSE201680 | TRUE | TRUE | TRUE | | q2 | |
| GSE202471 | TRUE | TRUE | TRUE | | q2 | |
| GSE202472 | TRUE | TRUE | TRUE | | q2 | |
| GSE202473 | TRUE | TRUE | TRUE | | q2 | |
| GSE206529 | TRUE | | TRUE | | q2 | |
| GSE226108 | TRUE | | TRUE | | q2 | |
| GSE231536 | TRUE | | TRUE | | q2 | |
| GSE236097 | TRUE | TRUE | TRUE | | q2 | |
| GSE243413 | FALSE | | TRUE | | q2 | |
| GSE247157 | TRUE | | TRUE | | q2 | |
| GSE266191 | TRUE | | TRUE | | q2 | |
| GSE266191 | TRUE | | TRUE | | q2 | |
| GSE272353 | TRUE | | TRUE | | q2 | |
| GSE275494 | TRUE | TRUE | | | q2 | |
| GSE276851 | TRUE | | TRUE | | q2 | |
| GSE276864 | TRUE | | TRUE | | q2 | |
| GSE276923 | TRUE | | TRUE | | q2 | |
| GSE277326 | TRUE | | TRUE | | q2 | |
| GSE277361 | TRUE | | TRUE | | q2 | |
| GSE281526 | TRUE | | TRUE | | q2 | |
| GSE289703 | TRUE | TRUE | TRUE | | q2 | |
| GSE293982 | TRUE | TRUE | | | q2 | |
| GSE293984 | TRUE | TRUE | TRUE | | q2 | |
| GSE303302 | TRUE | TRUE | | | q2 | |
| GSE303793 | TRUE | TRUE | | | q2 | |
| GSE309408 | FALSE | | TRUE | | q2 | |
| GSE312253 | TRUE | TRUE | | | q2 | |
| GSE312445 | FALSE | | TRUE | | q2 | |
| GSE235177 | FALSE | | FALSE | | q3 | |
| GSE239775 | FALSE | | TRUE | | q3 | not a scRNASeq |
| GSE240218 | FALSE | | FALSE | | q3 | |
| GSE240446 | FALSE | | FALSE | | q3 | |
| GSE280004 | TRUE | | TRUE | | q3 | |
| GSE283746 | TRUE | | TRUE | | q3 | |
| GSE285335 | TRUE | | TRUE | | q3 | |
| GSE287751 | TRUE | TRUE | | | q3 | immune cells |
| GSE289752 | TRUE | | TRUE | | q3 | |
| GSE296830 | FALSE | | FALSE | | q3 | |
| GSE299820 | FALSE | | FALSE | | q3 | |
| GSE308682 | FALSE | | FALSE | | q3 | |
| GSE308745 | TRUE | | TRUE | | q3 | |
| GSE309688 | FALSE | | TRUE | | q3 | no a scRNASeq |
| GSE310382 | FALSE | | FALSE | | q3 | |
| GSE310625 | FALSE | | FALSE | | q3 | |
| GSE310729 | FALSE | | TRUE | | q3 | |
| GSE311112 | TRUE | | TRUE | | q3 | |
| GSE312336 | FALSE | | TRUE | | q3 | no control |
| GSE314342 | TRUE | | FALSE | | q3 | |
| GSE315138 | TRUE | TRUE | | | q3 | |
| GSE316149 | FALSE | | TRUE | | q3 | |

| | | | | | | |
|---|---|---|---|---|---|---|
| GSE316337 | TRUE | | TRUE | | q3 | |
| GSE316872 | TRUE | | TRUE | | q3 | |
| GSE318329 | FALSE | | FALSE | | q3 | |
| GSE318863 | TRUE | | FALSE | | q3 | mtx file inside the RAR |
| GSE318881 | FALSE | | TRUE | | q3 | |
| GSE318901 | FALSE | | FALSE | | q3 | |
| GSE319605 | FALSE | TRUE | | | q3 | |
| GSE319779 | FALSE | TRUE | | | q3 | |
| GSE319783 | FALSE | TRUE | | | q3 | |
| GSE202734 | TRUE | | TRUE | | q4 | |
| GSE214210 | TRUE | | TRUE | | q4 | |
| GSE236905 | FALSE | TRUE | | | q4 | |
| GSE244627 | TRUE | | TRUE | | q4 | |
| GSE250019 | TRUE | | TRUE | | q4 | |
| GSE254726 | TRUE | | TRUE | | q4 | |
| GSE254728 | TRUE | | TRUE | | q4 | |
| GSE266191 | TRUE | | TRUE | | q4 | |
| GSE268630 | TRUE | | TRUE | | q4 | |
| GSE272733 | TRUE | | TRUE | | q4 | |
| GSE279314 | TRUE | TRUE | TRUE | | q4 | |
| GSE284180 | TRUE | | TRUE | | q4 | |
| GSE285980 | TRUE | | TRUE | | q4 | |
| GSE288394 | TRUE | | TRUE | | q4 | |
| GSE289547 | TRUE | | TRUE | | q4 | |
| GSE290274 | TRUE | | TRUE | | q4 | |
| GSE290325 | TRUE | | TRUE | | q4 | |
| GSE300175 | TRUE | | TRUE | | q4 | |
| GSE302350 | TRUE | | TRUE | | q4 | |
| GSE302680 | TRUE | | TRUE | | q4 | |
| GSE304885 | TRUE | | TRUE | | q4 | |
| GSE307731 | TRUE | | TRUE | | q4 | |
| GSE308611 | TRUE | | TRUE | | q4 | |
| GSE311423 | TRUE | | TRUE | | q4 | |
| GSE312204 | TRUE | | TRUE | | q4 | |
| GSE313298 | TRUE | | TRUE | | q4 | |
| GSE313299 | TRUE | | TRUE | | q4 | |
| GSE313731 | TRUE | | TRUE | | q4 | |
| GSE314706 | TRUE | | TRUE | | q4 | |
| GSE314943 | TRUE | | TRUE | | q4 | |
| GSE316401 | TRUE | | TRUE | | q4 | |
| GSE316872 | TRUE | | TRUE | | q4 | |
| GSE317025 | TRUE | | TRUE | | q4 | |
| GSE124323 | FALSE | | FALSE | | q5 | |
| GSE124324 | FALSE | | FALSE | | q5 | |
| GSE151610 | FALSE | | FALSE | | q5 | |
| GSE160099 | FALSE | | FALSE | | q5 | |
| GSE175895 | FALSE | | FALSE | | q5 | |
| GSE212520 | FALSE | | FALSE | | q5 | |
| GSE212523 | FALSE | | FALSE | | q5 | |
| GSE213933 | FALSE | | FALSE | | q5 | |
| GSE221860 | FALSE | | FALSE | | q5 | |
| GSE221861 | FALSE | | FALSE | | q5 | |
| GSE235462 | FALSE | | FALSE | | q5 | |
| GSE249130 | FALSE | | FALSE | | q5 | |
| GSE254272 | FALSE | | FALSE | | q5 | |
| GSE270418 | FALSE | | FALSE | | q5 | |
| GSE285562 | FALSE | | FALSE | | q5 | |
| GSE286440 | FALSE | | FALSE | | q5 | |
| GSE288833 | FALSE | | FALSE | | q5 | |
| GSE291008 | FALSE | | FALSE | | q5 | |
| GSE297297 | FALSE | | FALSE | | q5 | |
| GSE306082 | FALSE | | FALSE | | q5 | |
| GSE309829 | FALSE | | FALSE | | q5 | |
| GSE311934 | FALSE | | FALSE | | q5 | |
| GSE313168 | FALSE | | FALSE | | q5 | |
| GSE315579 | FALSE | | FALSE | | q5 | |
| GSE327656 | FALSE | | FALSE | | q5 | |
| GSE327855 | FALSE | | FALSE | | q5 | |
| GSE128142 | TRUE | TRUE | | | q6 | |
| GSE137311 | TRUE | | | TRUE | q6 | |
| GSE147013 | FALSE | | FALSE | | q6 | fibroblasts |
| GSE181251 | TRUE | TRUE | TRUE | | q6 | |
| GSE181324 | FALSE | | FALSE | | q6 | |
| GSE196097 | TRUE | | TRUE | | q6 | |
| GSE196106 | TRUE | | TRUE | | q6 | |
| GSE269694 | FALSE | TRUE | | | q6 | not retina |
| GSE273668 | TRUE | | TRUE | | q6 | |
| GSE315262 | TRUE | | TRUE | | q6 | |

| | | | | | |
|---|---|---|---|---|---|
| GSE116106 | TRUE | | | TRUE | q7 |
| GSE135167 | TRUE | | TRUE | | q7 |
| GSE138002 | TRUE | | | TRUE | q7 |
| GSE147979 | TRUE | | | TRUE | q7 |
| GSE149040 | TRUE | | TRUE | | q7 |
| GSE149715 | TRUE | | | TRUE | q7 |
| GSE165784 | TRUE | | | TRUE | q7 |
| GSE169039 | TRUE | | TRUE | | q7 |
| GSE169097 | TRUE | | TRUE | | q7 |
| GSE169102 | TRUE | | TRUE | | q7 |
| GSE174215 | TRUE | | | TRUE | q7 |
| GSE179754 | FALSE | | | TRUE | q7 |
| GSE188639 | TRUE | | TRUE | | q7 |
| GSE192814 | TRUE | | TRUE | | q7 |
| GSE192882 | TRUE | | TRUE | | q7 |
| GSE199317 | TRUE | | | TRUE | q7 |
| GSE201254 | TRUE | | | TRUE | q7 |
| GSE222389 | TRUE | | TRUE | | q7 |
| GSE242941 | TRUE | | TRUE | | q7 |
| GSE247166 | TRUE | | TRUE | | q7 |
| GSE255520 | TRUE | | TRUE | | q7 |
| GSE263879 | TRUE | TRUE | | | q7 |
| GSE265774 | TRUE | | | TRUE | q7 |
| GSE275131 | TRUE | | TRUE | | q7 |
| GSE282444 | TRUE | | TRUE | | q7 |
| GSE285843 | TRUE | | TRUE | | q7 |
| GSE286217 | TRUE | | TRUE | | q7 |
| GSE286897 | TRUE | | TRUE | | q7 |
| GSE287253 | TRUE | | TRUE | | q7 |
| GSE287347 | TRUE | | TRUE | | q7 |
| GSE291654 | TRUE | | TRUE | | q7 |
| GSE292463 | TRUE | | TRUE | | q7 |
| GSE292834 | TRUE | | TRUE | | q7 |
| GSE294652 | TRUE | | TRUE | | q7 |
| GSE295601 | TRUE | | TRUE | | q7 |
| GSE295977 | TRUE | | TRUE | | q7 |
| GSE297627 | TRUE | | TRUE | | q7 |
| GSE299589 | TRUE | | TRUE | | q7 |
| GSE300588 | TRUE | | TRUE | | q7 |
| GSE300607 | TRUE | | TRUE | | q7 |
| GSE301651 | FALSE | | TRUE | | q7 |
| GSE304010 | TRUE | | TRUE | | q7 |
| GSE305067 | TRUE | | TRUE | | q7 |
| GSE306682 | TRUE | | TRUE | | q7 |
| GSE306737 | TRUE | | TRUE | | q7 |
| GSE307707 | TRUE | | TRUE | | q7 |
| GSE307731 | TRUE | | TRUE | | q7 |
| GSE308745 | TRUE | | TRUE | | q7 |
| GSE310490 | TRUE | | TRUE | | q7 |
| GSE310529 | TRUE | | TRUE | | q7 |
| GSE310923 | TRUE | | TRUE | | q7 |
| GSE310929 | TRUE | | TRUE | | q7 |
| GSE311208 | TRUE | | TRUE | | q7 |
| GSE312053 | TRUE | | TRUE | | q7 |
| GSE312336 | TRUE | | TRUE | | q7 |
| GSE312549 | TRUE | | TRUE | | q7 |
| GSE312862 | TRUE | | TRUE | | q7 |
| GSE313291 | TRUE | | TRUE | | q7 |
| GSE314071 | FALSE | TRUE | | | q7 |
| GSE314072 | TRUE | TRUE | | | q7 |
| GSE314399 | TRUE | | TRUE | | q7 |
| GSE314943 | TRUE | | TRUE | | q7 |
| GSE316301 | TRUE | | TRUE | | q7 |
| GSE316869 | TRUE | | TRUE | | q7 |
| GSE317019 | FALSE | | TRUE | | q7 |
| GSE317349 | FALSE | | TRUE | | q7 |
| GSE319369 | TRUE | | TRUE | | q7 |
| GSE319479 | TRUE | | TRUE | | q7 |
| GSE319557 | TRUE | | TRUE | | q7 |
| GSE132229 | TRUE | | TRUE | | q8 |
| GSE138002 | TRUE | | | TRUE | q8 |
| GSE147979 | FALSE | | | TRUE | q8 |
| GSE181251 | TRUE | TRUE | | | q8 |
| GSE199840 | TRUE | | TRUE | | q8 |
| GSE202886 | TRUE | | TRUE | | q8 |
| GSE230049 | TRUE | | TRUE | | q8 |
| GSE237202 | TRUE | | TRUE | | q8 |
| GSE237203 | TRUE | | TRUE | | q8 |
| GSE237204 | TRUE | | TRUE | | q8 |
| GSE237205 | TRUE | | TRUE | | q8 |
| GSE237206 | TRUE | | TRUE | | q8 |

| | | | | | |
|---|---|---|---|---|---|
| GSE237207 | TRUE | | TRUE | | q8 |
| GSE237208 | TRUE | | TRUE | | q8 |
| GSE237209 | TRUE | | TRUE | | q8 |
| GSE237210 | TRUE | | TRUE | | q8 |
| GSE237211 | TRUE | | TRUE | | q8 |
| GSE237212 | TRUE | | TRUE | | q8 |
| GSE237213 | TRUE | | TRUE | | q8 |
| GSE237214 | TRUE | | TRUE | | q8 |
| GSE237215 | FALSE | | TRUE | | q8 |
| GSE242229 | TRUE | | TRUE | | q8 |
| GSE245137 | TRUE | | TRUE | | q8 |
| GSE254587 | TRUE | | TRUE | | q8 |
| GSE268630 | TRUE | | TRUE | | q8 |
| GSE269337 | TRUE | | TRUE | | q8 |
| GSE281526 | TRUE | | TRUE | | q8 |
| GSE284729 | TRUE | | TRUE | | q8 |
| GSE289547 | TRUE | | TRUE | | q8 |
| GSE307302 | TRUE | | TRUE | | q8 |
| GSE308982 | TRUE | | TRUE | | q8 |
| GSE309445 | TRUE | | TRUE | | q8 |
| GSE312067 | TRUE | | TRUE | | q8 |
| GSE314867 | TRUE | | TRUE | | q8 |
| GSE135092 | FALSE | | TRUE | | q9 |
| GSE135133 | TRUE | | TRUE | | q9 |
| GSE137311 | TRUE | | | TRUE | q9 |
| GSE144300 | FALSE | | TRUE | | q9 |
| GSE147573 | FALSE | | FALSE | | q9 |
| GSE148077 | FALSE | | TRUE | | q9 |
| GSE155154 | TRUE | | TRUE | | q9 |
| GSE160306 | TRUE | | TRUE | | q9 |
| GSE160306 | FALSE | | FALSE | | q9 |
| GSE160308 | TRUE | | TRUE | | q9 |
| GSE164092 | FALSE | | TRUE | | q9 |
| GSE165743 | FALSE | | FALSE | | q9 |
| GSE175937 | TRUE | | TRUE | | q9 |
| GSE177039 | FALSE | | FALSE | | q9 |
| GSE181550 | FALSE | | TRUE | | q9 |
| GSE212189 | FALSE | | FALSE | | q9 |
| GSE220155 | FALSE | | TRUE | | q9 |
| GSE221042 | FALSE | | TRUE | | q9 |
| GSE226108 | TRUE | | TRUE | | q9 |
| GSE234447 | FALSE | | TRUE | | q9 |
| GSE247157 | TRUE | | TRUE | | q9 |
| GSE262534 | TRUE | TRUE | | | q9 |
| GSE265774 | TRUE | | TRUE | TRUE | q9 |
| GSE268630 | TRUE | | TRUE | | q9 |
| GSE271251 | FALSE | | TRUE | | q9 |
| GSE276851 | FALSE | | FALSE | | q9 |
| GSE276923 | FALSE | | FALSE | | q9 |
| GSE277326 | TRUE | | FALSE | | q9 |
| GSE277361 | FALSE | | TRUE | | q9 |
| GSE281526 | TRUE | | TRUE | | q9 |
| GSE282457 | FALSE | | TRUE | | q9 |
| GSE298665 | TRUE | TRUE | | | q9 |
| GSE310659 | FALSE | | TRUE | | q9 |
| GSE311369 | FALSE | | TRUE | | q9 |
| GSE311370 | FALSE | | TRUE | | q9 |
| GSE128142 | TRUE | TRUE | | | q10 |
| GSE133492 | TRUE | | FALSE | | q10 |
| GSE137311 | TRUE | | | TRUE | q10 |
| GSE164904 | FALSE | | FALSE | | q10 |
| GSE180157 | FALSE | | FALSE | | q10 |
| GSE181251 | TRUE | TRUE | TRUE | | q10 |
| GSE214175 | FALSE | | FALSE | | q10 |
| GSE241255 | TRUE | TRUE | | | q10 |
| GSE244374 | TRUE | | TRUE | | q10 |
| GSE254741 | TRUE | | TRUE | | q10 |
| GSE266083 | FALSE | | FALSE | | q10 |
| GSE269694 | TRUE | TRUE | | | q10 |
| GSE286289 | FALSE | | FALSE | | q10 |
| GSE294933 | TRUE | | FALSE | | q10 |
| GSE294966 | TRUE | TRUE | FALSE | | q10 |
| GSE296097 | TRUE | TRUE | | | q10 |
| GSE296953 | FALSE | TRUE | | | q10 |
| GSE298334 | FALSE | | FALSE | | q10 |
| GSE301937 | FALSE | TRUE | | | q10 |
| GSE301939 | TRUE | TRUE | | | q10 |
| GSE303721 | FALSE | | FALSE | | q10 |
| GSE306878 | TRUE | | FALSE | | q10 |
| GSE306960 | TRUE | | FALSE | | q10 |

| | | | | | |
|---|---|---|---|---|---|
| GSE307920 | FALSE | | FALSE | | q10 |
| GSE309944 | TRUE | | TRUE | | q10 |
| GSE310167 | TRUE | | FALSE | | q10 |
| GSE311787 | FALSE | | FALSE | | q10 |
| GSE315021 | FALSE | | FALSE | | q10 |
| GSE315354 | FALSE | | FALSE | | q10 |
| GSE315996 | FALSE | | FALSE | | q10 |
| GSE315997 | FALSE | | FALSE | | q10 |
| GSE316009 | FALSE | | FALSE | | q10 |
| GSE316339 | FALSE | | FALSE | | q10 |
| GSE318080 | FALSE | | FALSE | | q10 |
| GSE318663 | FALSE | | FALSE | | q10 |
| GSE80571 | FALSE | | FALSE | | q10 |
| GSE80686 | TRUE | | TRUE | | q10 |

Table S4: Sample counts relavent to Figure 3A-B

| labels | count |
|---|---|
| Retina | 33 |
| retina | 32 |
| retinal organoid | 9 |
| Whole blood | 5 |
| blood | 5 |
| Blood | 4 |
| Peripheral blood | 4 |
| fetal retina | 4 |
| RPE | 3 |
| Whole Blood | 3 |
| Blood plasma | 2 |
| Peripheral Blood Mononuclear Cells | 1 |

Table S5 - RAG evaluation metrics

| Evaluation category | Metric | Definition |
|---|---|---|
| Answer Quality | Answer Relevancy | Measures how directly and appropriately the generated answer addresses the user's query. |
| | Faithfulness | Measures whether claims in the generated answer are supported by the retrieved evidence without unsupported or contradictory statements. |
| Retrieval Context Quality | Contextual Relevancy | Measures how relevant and useful the retrieved context is for answering the query. |
| | Contextual Precision | Measures whether the most relevant retrieved contexts are prioritized and ranked ahead of less relevant contexts. |
| Grounding & Constraint Compliance | Contextual Recall | Measures whether the retrieved context captures the information needed to support the expected answer. |
| | Precision Grounding | Measures the extent to which specific claims in the generated answer are directly grounded in the retrieved evidence. |
| | Constraint Adherence | Measures whether the generated answer satisfies the explicit conditions and restrictions specified in the query. |
